# Insect egg size and shape evolve with ecology, not developmental rate

**DOI:** 10.1101/471946

**Authors:** Samuel H. Church, Seth Donoughe, Bruno A. S. de Medeiros, Cassandra G. Extavour

**Affiliations:** Department of Organismic and Evolutionary Biology, Harvard University, Cambridge, MA 02138, United States; Current address: Department of Cell and Molecular Biology, University of Chicago, Chicago, IL 60637, United States; Department of Molecular and Cellular Biology, Harvard University, Cambridge, MA 02138, United States

## Abstract

The evolution of organism size is hypothesized to be predicted by a combination of development, morphological constraints, and ecological pressures. However, tests of these predictions using phylogenetic methods have been limited by taxon sampling. To overcome this limitation, we generated a database of more than ten thousand observations of insect egg size and shape from the entomological literature and combined them with published genetic and novel life-history datasets. This enabled us to perform phylogenetic tests of long-standing predictions in size evolution across hexapods. Here we show that across eight orders of magnitude in egg volume variation, the relationship between egg shape and size itself evolves, such that predicted universal patterns of scaling do not adequately explain egg shape diversity. We test the hypothesized relationship between size and development, and show that egg size is not correlated with developmental rate across insects, and that for many insects egg size is not correlated with adult body size either. Finally, we show that the evolution of parasitism and aquatic oviposition both help to explain the diversification of egg size and shape across the insect evolutionary tree. Our study challenges assumptions about the evolutionary constraints on egg morphology, suggesting that where eggs are laid, rather than universal mathematical allometric constants, underlies egg size and shape evolution.

## 2 Body text

Size is a fundamental factor in many biological processes. Size may impact ecological interactions^8,9^, it scales with features of morphology and physiology^10^, and in many cases larger animals have higher fitness^11^. Previous studies have aimed to identify the macroevolutionary forces that explain observed size distributions^8,12,13^, but limited data on the phylogenetic distribution of size has precluded robust tests of these predicted forces^11,14^. Here we address this problem by assembling a dataset of insect egg phenotypes with sufficient taxon sampling to rigorously test evolutionary hypotheses in a phylogenetic framework, and use these data to elucidate the roles of allometric constraints, development, and ecology in size and shape evolution.

We have chosen insect eggs as a compelling system in which to test macroevolutionary hypotheses about the causes and consequences of size evolution^14^. Insect eggs are extraordinarily diverse^15^, yet their morphologies can be readily compared across distant lineages using quantitative traits. Changes in egg size are often assumed to be linked to changes in other aspects of organismal biology^16^, including adult body size^17,18^, features of adult anatomy^19,20^, and offspring fitness via maternal investment^21^. The egg is an independent ecological vessel that must withstand the physiological challenges of being laid in diverse microenvironments, including water, air, or internal to plants or animal hosts^22^. Furthermore, because the fertilized egg is the homologous, single-cell stage in the life-cycle of multicellular organisms, egg size diversity is relevant to both cell size and organism size evolution^15,21^

Three classes of hypotheses have historically been proposed to explain the evolution of egg size and shape: [1] geometric constraints due to the physical scaling of size and shape^19,20,23–25^, [2] the interaction between size and the rate of development^26–29^, and [3] the diversification of size and shape in response to ecological or life-history changes^17,19,22,30^. We use a large-scale phylogenetic approach to test all three of these hypotheses, and show that many presumed universal patterns in egg size, shape, and embryonic development are not supported across insects. Instead, our results show that models accounting for ecological change best explain extant egg morphological diversity.

Using custom bioinformatic tools, we assembled a database of 10,449 morphological descriptions of eggs from the published scientific literature, comprising 6,706 species, 526 families, and every extant insect and non-insect hexapod order (Figs. 1A, S1, S2)^31^. We combined this database with backbone hexapod phylogenies^1,32^ that we enriched to include taxa in the egg morphology database (Fig. S3), and used it to describe the distribution of egg shape and size (Fig. 1B). Our results showed that insect eggs span more than eight orders of magnitude in volume (Figs. 1C, S4), and revealed new candidates for the smallest and largest described insect eggs: respectively, a parasitoid wasp Platygaster vernalis^7^ (volume = 7 x 10^-7^ mm^3^), and an earth-boring beetle, *Bolboleaus hiaticollis*^2^ (volume = 5 x 10^2^ mm^3^) (Fig. 1C).

**Figure 1:**
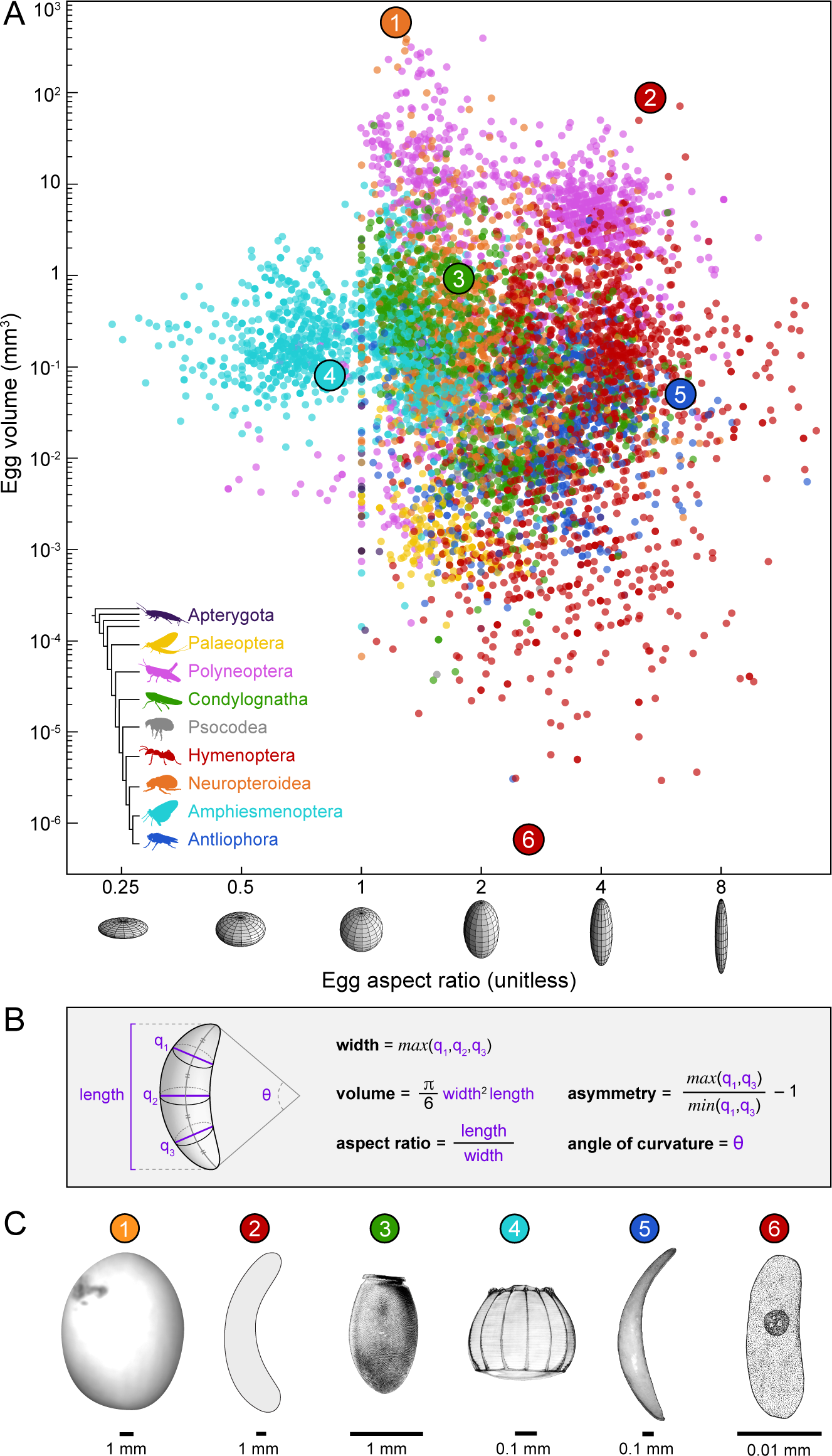
The shapes and sizes of hexapod eggs. **A**, The distribution of egg descriptions in a morphospace defined by egg volume (mm^3^) and aspect ratio (unitless). Both traits are plotted on a log scale. Points are colored according to the groups on the in the lower left, which illustrates relationships according to Misof et al. 2014^1^, one of the backbone phylogenies used in this study. Numbered points correspond to the six eggs shown in C, chosen as examples of specific or extreme sizes or shapes. There are 10,449 published egg descriptions in the database; the plot shows the 7,935 descriptions with both volume and aspect ratio data. **B**, Egg shape is described in the database with three parameters (aspect ratio, asymmetry, and angle of curvature) calculated from the measurements shown in purple (see text for details). **C**, Images of six example eggs, arranged from largest to smallest and oriented with the axis of rotational symmetry vertical: 1. Earth-borer beetle *Bolboleaus hiaticollis*^2^, 2. Tropical carpenter bee *Xylocopa latipes*^3^, 3. Kissing bug *Rhodnius robustus*^4^, 4. Many-banded daggerwing butterfly *Marpesia chiron*^5^, 5. Tephritid fruit fly *Anastrepha distincta*^6^, 6. Parasitoid wasp *Platygaster vernalis*^7^.

Plotting eggs by morphological traits revealed that some egg shapes have evolved only in certain clades (Fig. 1C, S5, S6). For example, oblate ellipsoid eggs (with an aspect ratio smaller than one) are found only in moths, butterflies, and stoneflies (Amphismenoptera: Lepidoptera and Polyneoptera: Plecoptera; image 4 in Fig. 1C). However, the most prominent pattern was that distantly related insects have converged on similar egg morphologies many times independently (Fig. 1C, S6). This high degree of morphological convergence allowed us to perform robust tests of phenotypic associations across independent evolutionary events.

### 2.1 Evolutionary patterns of egg shape and size

Two opposing hypotheses based on predicted geometric constraints have been proposed to explain the evolutionary relationship between propagule shape and size. *Hypothesis A:* When eggs evolve to be larger, they get wider (increases in egg size are associated with decreases in aspect ratio)^24,25^. This hypothesis predicts a reduction in relative surface area as size increases, which has been proposed as a solution to the presumed cost of making eggshell material^25^. *Hypothesis B:* When eggs evolve to be larger, they get longer (increases in egg size are associated with increases in aspect ratio)^19,20,25^,. This hypothesis predicts a reduction in relative cross sectional area as eggs get larger, which has been proposed as a solution to the need for eggs to pass through a narrow opening during oviposition^19,20^.

To test these hypotheses about the physical scaling of size and shape, we first modeled the individual evolutionary history of each morphological trait across insects. This allows us to first determine whether distributions of extant shape and size have been shaped by phylogenetic relationships, or if these traits are instead stochastically distributed across insects. For egg volume, aspect ratio, asymmetry, and angle of curvature (Fig. 1B), we considered four models of potential evolutionary change in egg size and shape, and asked which of these best fit our data: Brownian motion (BM), Brownian motion with evolutionary friction (Ornstein-Uhlenbeck, OU), Brownian motion with a decreasing rate of evolution (Early Burst, EB), and a non-phylogenetic model of stochastic motion (white noise, WN). We found strong evidence that models that account for covariance based on phylogeny fit our data better than a non-phylogenetic model (WN); in other words, insect egg morphology tends to be similar in closely related insects. For egg size and aspect ratio, an ‘Early-Burst’ (EB) model in which the rate of evolution decreases over time, best describes the observed distributions of these two traits. This change in rate is distributed non-uniformly over the insect phylogeny, with some clades evolving more rapidly than others (Figs. S8-10). In earlier studies EB models were rarely detected^33^; however our findings are consistent with recent work evaluating datasets that, like ours, comprise many taxa and orders of magnitude in morphological variation^34,35^. Having established the phylogenetic models that best describe egg trait evolution, we used these results to test hypotheses about the relationship between egg shape and size.

To test which aforementioned scaling relationship best describes insect egg evolution, we compared support for Hypotheses A and B using a phylogenetic generalized least-squares approach (PGLS) to determine the scaling exponent of egg length and width (the slope of the regression of log-length and log-width). A slope less than one would support Hypothesis A, while a slope greater than one would support Hypothesis B^36^. An alternative, Hypothesis C, is that egg shape remains the same as size changes, which would result in a slope near one (an isometric relationship). We found that Hypothesis B is best supported across all insects: larger eggs have higher aspect ratios than smaller eggs (0 < p-value < 0.005, slope = 0.76, Fig. 2B, Table S6), even when controlling for adult body size (Fig. S13, Table 8). However, the allometric relationship between size and shape evolves dynamically across the phylogeny, as has also been shown for metabolic scaling in mammals^37^. Among the large clades we tested, Hypothesis C could not be rejected for Neuropteroidea (p-value of alternative hypothesis test = 0.02; Fig. 2C, 2F, S11, S12, Table S7). Calculating the scaling relationship on subgroups of each of these major lineages revealed that many additional insect clades, including mayflies, crickets and grasshoppers, and shield bugs, also had eggs with isometric scaling of shape and size (Fig. S12). The marked differences in scaling exponents are evidence that throughout the course of insect diversification, egg evolution was not governed by a universal allometric scaling constant. Instead, evolutionary forces beyond the constraints of physical scaling (e.g. development or ecology) are required to explain egg morphological diversification.

**Figure 2:**
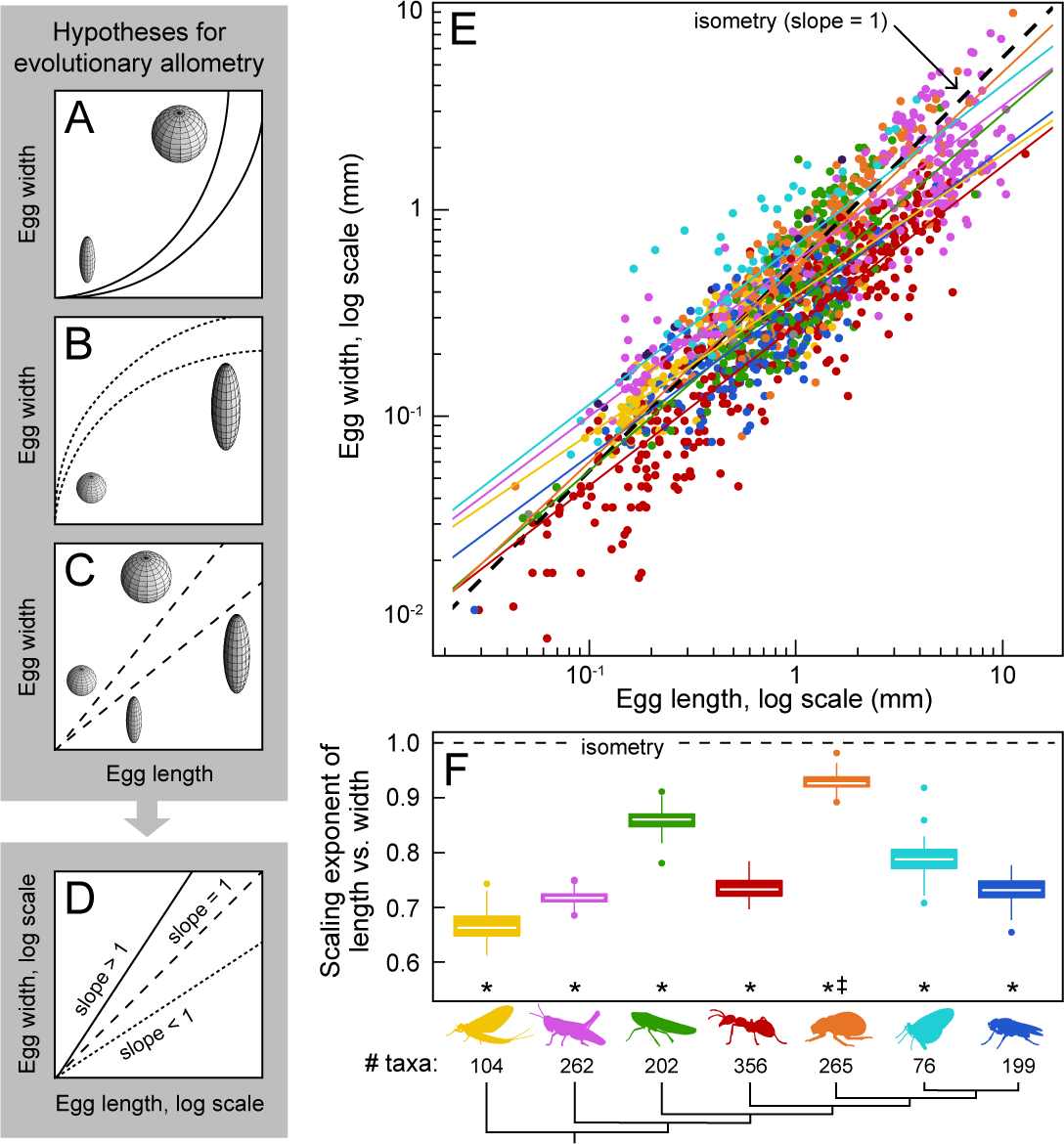
The allometric relationship of egg shape and size evolves across insect lineages. Historically hypothesized allometric relationships between egg size and shape: **A**, larger eggs are proportionally wider than smaller counterparts, (solid line); **B**, larger eggs are proportionally longer (dotted line); and **C**, insect egg shape and size scale isometrically (dashed line). Hypothetical eggs for each hypothesis are shown as schematics with length axis vertical. **D**, Each hypothesis (solid line = A, dotted line = B, dashed line = C) predicts a different value of the scaling exponent—that is, the slope of the regression between egg length and width in log-log space. **E**, The distribution of egg length and width in log-log space. The dashed black line represents a hypothetical 1:1 relationship (isometry, hypothesis C). Solid colored lines are the phylogenetic regressions for seven clades, and each colored point is a randomly selected representative egg for a genus. **F**, Distribution of the scaling exponents for the seven monophyletic insect clades included in this analysis, calculated over the posterior distribution of trees. We resampled tree topology and genus-level representatives from the egg morphology dataset 100 times, calculating a scaling exponent each time. White lines, boxes, bars, and dots represent median, 25-to-75th percentiles, 5-to-95th percentiles, and outliers, respectively. Asterisks indicate that the relationship between length and width is significant (p-value threshold <0.01, exact values in Table S6), and double-dagger indicates the relationship is not statistically distinguishable from isometry (p-value threshold >0.01, exact values in Table S7). In **E** and **F** the colors correspond to the clades shown in Fig. 1A.

### 2.2 Developmental traits and egg evolution

The egg is the starting material for embryogenesis, and the size of the hatchling is directly related to the size of the egg at fertilization^38^. It was previously reported that embryogenesis takes longer in species with larger eggs^29^, and that this relationship could influence size evolution^26–29^. This prediction is consistent with the observation that larger adult species have a lower metabolic rate than smaller species^39^. To test this prediction across our expanded egg dataset, we assembled embryological records from published studies, and found that indeed, simply comparing egg volume and duration of embryogenesis yields the previously reported positive relationship^29^ (Fig. S22). However, a linear regression that does not account for phylogenetic relationships is inappropriate for this analysis due to the covariance of traits on an evolutionary tree^40^. When we accounted for such potential phylogenetic covariance of data, we found that there is no significant relationship between egg size and duration of embryogenesis across insects, such that eggs of very different sizes can develop at a similar rate and vice versa (0.08 < p-value < 0.26; Fig. 3B, S21, Table S21). These results suggest that the often-invoked trade-off between size and development^26–29^ does not hold across insect species.

**Figure 3:**
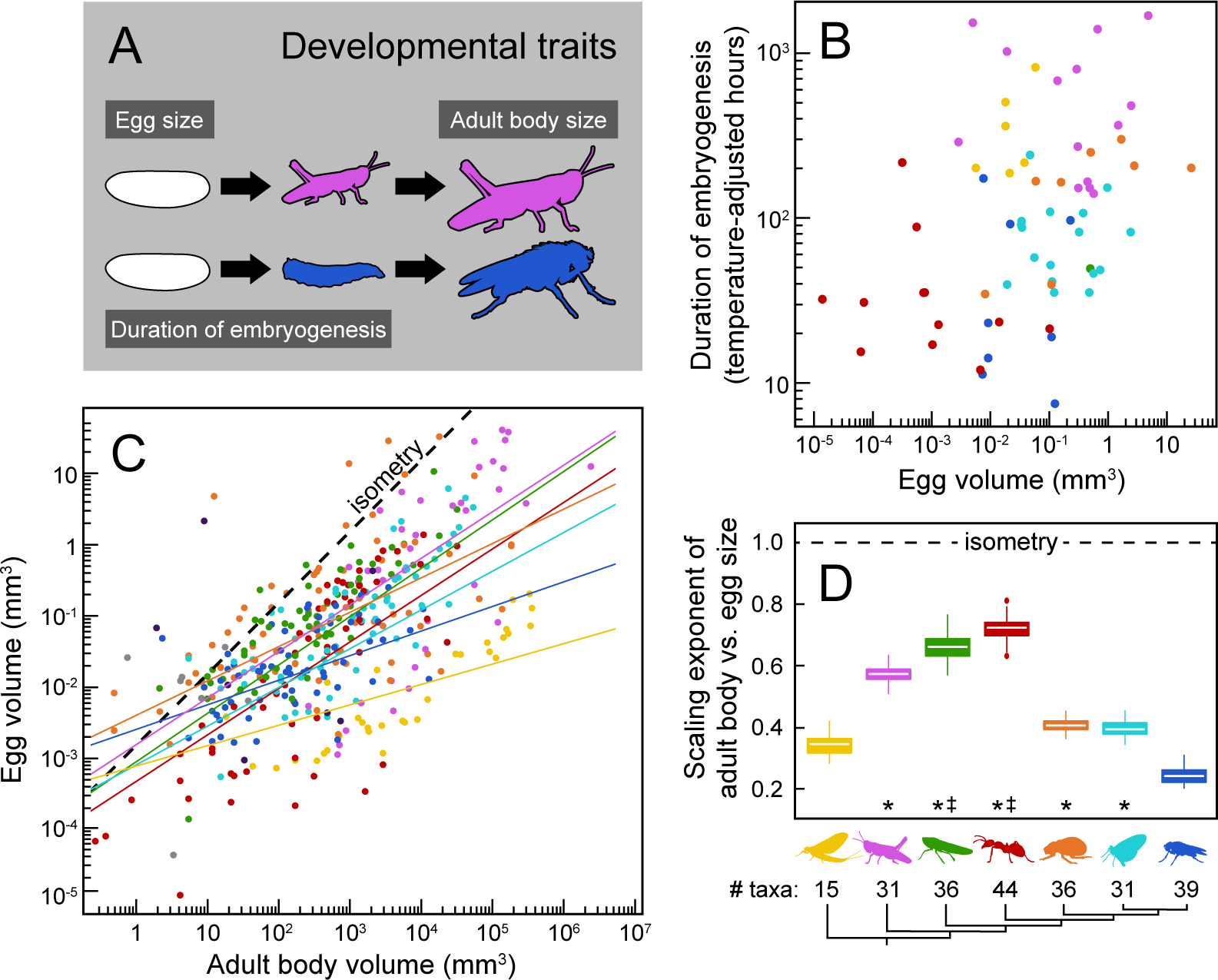
Rate of embryogenesis does not co-vary predictably with egg size. **A**, Mature eggs undergo embryonic development, hatch, and grow into adult insects. We define duration of embryogenesis as the time elapsed from egg-laying to the point at which the nymph (in Hemimetabola) or larva (in Holometabola) emerges from the egg. **B**, Duration of embryogenesis, adjusted for incubation temperature (hours, plotted on a log scale), compared to egg volume (mm^3^, plotted on a log scale). Each colored point represents an insect species for which duration of time-to-hatching has been reported (see Supplemental Information for sources), and egg morphological data are contained in our database^31^. When phylogenetic relationships are taken into account, there is no significant relationship between egg volume and duration of embryogenesis. **C**, Egg volume (mm^3^, plotted on a log scale) compared to adult body volume, calculated as body length cubed (mm^3^, plotted on a log scale). The dashed black line shows an example 1:1 relationship (isometry), solid colored lines are the phylogenetic regression for the seven clades included in this analysis, and colored points are family- and order-level averages for egg size and median adult insect size. **D**, The distributions of the allometric exponents for the seven monophyletic insect clades included in this analysis. Asterisks indicate that the relationship between length and width is significant (p-value threshold <0.01, exact values in Table S12), and double-dagger indicates the relationship is not statistically distinguishable from isometry (p-value threshold >0.01, exact values in Table S13). Colors and labels in B—E correspond to the clades shown in Fig. 1A.

We also tested the hypothesis that the size of the egg has a positive relationship with adult body size. Previous work reported this relationship in subsets of insects, and moreover suggested that smaller insects lay proportionally larger eggs for their bodies^18,38,41^. Such a relationship between egg size and body size would result in an allometric scaling exponent less than one. We combined our dataset of egg size with published adult body length data for insect families^42^, and found that this relationship was not generalizable across all insect lineages. For example, in flies and their relatives (Antliophora) and mayflies and odonates (Palaeoptera), egg size is not predicted by body size, meaning that insects of similar body size lay differently sized eggs (p-value, Antliophora = 0.04, Palaeoptera = 0.03; Fig. 3C, 3D, Tables S22). In thrips and true bugs (Condylognatha) and in bees, ants, and wasps (Hymenoptera), an isometric relationship between egg size and body size cannot be rejected (p-value of alternative hypothesis test, Hymenoptera = 0.02, Condylognatha = 0.02, Fig. S23, Table S23). In general, the predictive power of the relationship between body size and egg size is low: average egg volume can vary by up to four orders of magnitude among species with similar body size (Fig. 3C). This decoupling of both body size and duration of embryogenesis from egg size evolution suggests that egg diversification has not been universally constrained by development across insects.

### 2.3 Changes in size and shape are explained by oviposition ecology

Egg size and shape have been predicted to evolve in response to changes in life-history and ecology. Recent work in birds has highlighted one such relationship, suggesting that birds with increased flight capability have more elliptical and asymmetrical eggs^19^. We asked whether an analogous relationship exists between insect flight capability and egg shape. Unlike birds, insects have undergone many hundreds of evolutionary shifts to flightless and even wingless forms^43^. We focused on two clades in which the patterns of flight evolution have been extensively studied. Stick insects (Phasmatodea) have flightless and wingless species^44,45^ (Fig. S17), and many butterflies (Lepidoptera) show migratory behavior^46^, which could be considered a proxy for increased flight capability relative to non-migratory taxa (Fig. S17). We found that, in contrast to birds, evolutionary changes in flight ability in these two insect clades were not associated with changes in egg shape (OU model with multiple optima per regime; ΔAICc < 2, exact values in Tables S17, S18).

Like flight capacity, the microenvironment that eggs experience varies widely across insects, including being exposed to air, submerged or floating in water, or contained within a host animal^15^ (Fig. 4A). Each of these microenvironments places different demands on the egg, such as variable access to oxygen and water during development^22^. Preliminary studies in small groups of insects suggested that evolutionary changes in oviposition ecology and life history may drive the evolution of egg size and shape^17,30^. To test this prediction robustly across all insects, we compiled records on two specific oviposition ecology modes that have been extensively studied: oviposition within an animal host (which can be in the host adult body or in the host egg, called internal parasitic oviposition) and oviposition in or on water. For each mode we reconstructed ancestral changes along the insect phylogeny, and found that both aquatic and internal parasitic oviposition have been gained and lost multiple times independently (Fig. 4A, B, S15, S16). This extensive convergent evolution allowed us to perform a strong test of whether egg size and shape evolution is predicted by the evolution of oviposition ecology.

**Figure 4:**
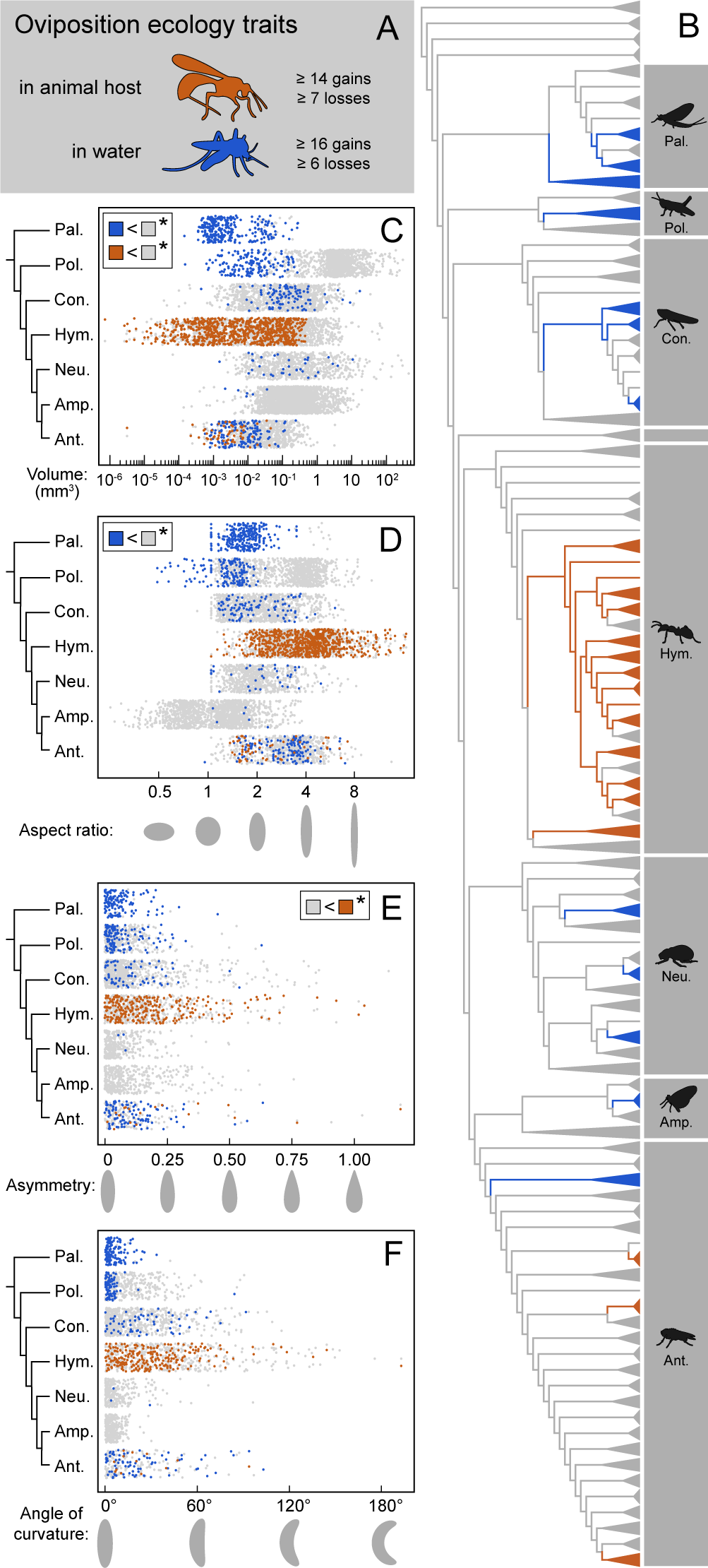
Shifts in oviposition ecology are associated with morphological changes in insect eggs. **A**, Laying eggs within an animal host (orange; e.g parasitoid wasps) and in water (blue; e.g. mosquitoes) are two modes of insect oviposition ecology. Other oviposition substrates (e.g. terrestrial) are shown in gray (see SI for further details on oviposition substrate classification). **B**, Ancestral state reconstruction of each oviposition ecology trait reveals that they have arisen multiple times in distantly related insect lineages (see Fig. S17 and S18 for full phylogeny). **C-F**, The distribution of four egg morphology parameters across insect clades, colored by oviposition ecology and plotted by phylogenetic group: **C**, volume (mm^3^, plotted on log scale); **D**, aspect ratio (unitless, plotted on log scale); **E**, asymmetry (unitless); and **F**, angle of curvature (degrees). The x-axes of **D-F** include theoretical egg silhouettes shown with the length axis vertical. Asterisks indicate that the model accounting for oviposition ecology (Ornstein-Uhlenbeck process with multiple regimes) fits the data better (ΔAICc >2, exact values in Table S14-S19) than a non-ecological model. Clade labels in **B-F** are abbreviations of clades shown in Fig. 1A.

We found that the evolution of new oviposition environments was linked to changes in egg size and shape. Models that accounted for shifts to new oviposition environments better explained egg size and shape distributions than models that did not (OU multistate model, ΔAICc > 2, exact values in Tables S14-S19). Specifically, shifts to aquatic oviposition were significantly associated with the evolution of smaller, wider eggs (Fig. 4C-D, Tables S11, S14), and shifts to internal parasitic oviposition were significantly associated with smaller, more asymmetric eggs (Figs. 4C, 4E, Table S11). Moreover, we note the smallest eggs in the dataset are from wasps with internal parasitic oviposition that develop polyembryonically (i.e. multiple embryos form from a single egg^47^; Fig. S7). Neither ecological change was associated significantly with consistent changes in the scaling relationship between size and shape (Fig. S18). These results were robust to uncertainty in how taxa had been classified for oviposition ecology; using broader ecological definitions, such as combining endo- and ectoparasites or semi-aquatic and riparian insects, did not change these results (Tables S12, S15, S13, S16). Taken together, these convergent evolutionary events are evidence that the microenvironment experienced by the egg following oviposition plays an important in role in the evolution of egg size and shape.

We have shown that insect eggs present an ideal example case for testing the predictability of macroevolutionary patterns in size and shape. By comparing egg size and shape across taxa, we find that prevalent assumptions about evolutionary trade-offs with developmental time, body size, or the cost of making egg shells do not hold across insects. Although we showed that time of development is not linked to egg size, we speculate that other features of development, such as cell number and distribution, may scale in predictable ways across eight orders of magnitude in egg size variation. We leveraged the power of the vast descriptive literature to overcome barriers common to macroevolutionary studies, establishing computational tools and methods that can be followed in future work. Finally, we provide evidence that the ecology of oviposition drives the evolution of egg size and shape.

## Supporting information

## 3 Acknowledgements

This work was supported by the National Science Foundation (NSF) under Grant No. IOS-1257217 to CGE, and NSF Graduate Research Training Fellowship No. DGE1745303 to SHC, and by a Jorge Paulo Lemann Fellowship to BdM from Harvard University. We thank the Extavour lab and Brian Farrell for discussion, and Casey Dunn, Dakota McCoy, Dan Rice, Elena Kramer, Jack Boyle, Leonora Bittleston, Mansi Srivastava, Milo Johnson, Peter Wilton, Richard Childers, and Sofia Prado-Irwin for suggestions on initial versions of this manuscript. We acknowledge the Ernst Mayr Library at the Museum of Comparative Zoology at Harvard, and specifically thank Mary Sears, for countless hours of support in gathering the references used in this study.

## 4 Author contributions

SHC and SD contributed equally to database generation, experimental design, data analysis, writing, and figure preparation. SHC wrote all code to perform statistical analyses. BdM performed the phylogenetic analyses. BdM and CGE contributed to experimental design, interpretation, and writing.

## 6 Methods

### 6.1 Data availability

The database of insect eggs is publicly available at Dryad https://datadryad.org/review?doi=doi:10.5061/dryad.pv40d2r and described in Church *et al*. 2018^1^. All code required to reproduce the analyses and figures shown here is available at https://github.com/shchurch/Insect_Egg_Evolution. The phylogenetic posterior distributions are provided as supplemental files ‘phylogeny_posterior_distribution_misof_back-bone.nxs’ and ‘phylogeny_posterior_distribution_rainford_backbone.nxs’.

### 6.2 Creating the insect egg database

A list of the 1,756 literature sources used to generate the egg database is provided as a supplemental file, ‘bibliography_egg_database.pdf’. A full description of the methods used to assemble the insect egg database have been published separately^1^. Egg descriptions were collected from published accounts of insect eggs using custom software to parse text from PDFs and measure published images (Fig. 1B), followed by manual verification. Each entry in the egg database includes a reference to an insect genus and, when reported, species name. Scientific names were validated using TaxReformer^1^, which relies on online taxonomic databases^2–6^.

### 6.3 Measuring egg features

Full trait definitions are described in the Supplementary Information and summarized briefly below. To resolve ambiguous cases and to measure published images, we used the definitions below.

#### Egg length

We defined egg length as the distance in millimeters (mm) of the axis of rotational symmetry.

#### Egg width

We defined egg width as the widest diameter (mm), measured perpendicular to the axis of rotational symmetry of the egg. For eggs described in published records as having both a width and breadth or depth (i.e. the egg is a flattened ellipsoid^7^), we defined width as the wider of the two diameters, and breadth as the diameter perpendicular to both the width and length.

#### Egg volume

Volume (mm^3^) was calculated using the equation for the volume of an ellipsoid: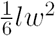, following previous workers^8,9^.

#### Egg aspect ratio

Aspect ratio was calculated as the ratio of length to width.

#### Egg asymmetry

Asymmetry was calculated as the ratio between the two egg diameters at the first and third quartile of the length axis. The first quartile was always defined as the larger of the two diameters, such that asymmetry is always between zero and one.

#### Angle of egg curvature

The angle of curvature was measured as the angle (degrees) of the arc created by the endpoints and midpoint of the length axis.

### 6.4 Phylogenetic methods

A genus-level phylogeny was built by combining mitochondrial 18S and 28S sequence data from the SILVA database^10–13^ with phylogenetic constraints from published higher-level insect phylogenies^14,15^. To account for phylogenetic uncertainty in comparative analyses, trees were estimated using a hierarchical approach^16,17^. Separate phylogenies for each insect order were inferred in a Bayesian framework using MrBayes v. 3.2.6^18^ and 100 randomly chosen post-burn-in trees for each order using the order-level backbone trees of Rainford et al.^15^ and Misof et al.^14^. See Supplemental Information, section “Sequence alignment and phylogenetic methods” for more details.

### 6.5 Annotating the egg database with developmental trait data

For developmental traits, a set of references were assembled from the embryological and ecological literature, and then used to compile data on interval between syncytial mitoses, time to cellularization, and duration of embryogenesis. Developmental rate observations were rescaled to approximate rates at a standardized temperature of 20°C following previous work^19^. For a full list of sources, methods used in this calculation, and further discussion of developmental trait definitions, see the Supplemental Information, section “Collecting developmental time data”.

### 6.6 Annotating the egg database with life-history trait data

For each of the ecological features of interest (internal parasitic oviposition, aquatic oviposition, flightlessness, and migratory behavior) taxonomic descriptions from the literature were matched to taxa in the insect egg database. For some taxonomic groups it was not possible to classify all members unambiguously. In these cases, the ecological state was coded “uncertain”, and the potential impact of this uncertainty on results was tested. For each trait the ancestral state reconstruction was estimated using an equal-rates model (R package corHMM^20^, function rayDISC, node.states=‘marginal’). For a full list of sources and methods used see the Supplemental Information, section “Evolutionary history of ecological traits”.

### 6.7 Data analysis and evolutionary model comparison

Egg length, width, volume, and aspect ratio were log10 transformed. Angle of curvature and asymmetry were square root transformed.

Models of evolution were compared using the R package geiger^21^. For each trait (egg length, width, volume, aspect ratio, asymmetry, and angle of curvature), the model fit of a Brownian Motion (BM), Ornstein-Uhlenbeck (OU), and Early-Burst (EB) model was compared against a null hypothesis of a white noise (WN) model that assumes no evolutionary correlation. See Supplemental Information, section ‘Evolutionary model fitting’ for details. The performance of the best fitting model was further analyzed by comparing expected values of parameters from simulations under the model to observed parameters, using the R package arbutus^22^.

The ancestral state of volume, aspect ratio, and angle of curvature were mapped on the summary phylogeny using the R package phytools^23^ (version 0.6-44, function contMap). Evolutionary rate regimes of volume, aspect ratio, and the angle of curvature were fit on the summary phylogeny using the program BAMM^24,25^ (version 2.5.0, R package BAMMtools verison 2.1.6, setBAMMpriors, prior for expected number of shifts set to 10, for 10,000,000 generations).

Following ancestral state reconstruction for ecological regimes (see above), for each trait (volume, aspect ratio, asymmetry, curvature) the fit of a Brownian-Motion model (BM), an Ornstein-Uhlenbeck model with a single optimum (OU1), and an Ornstein-Uhlenbeck model with an independent optimum for each ecological state (OUM) were compared using the R package OUwie^26^ (version 1.50).

All evolutionary regression analyses were performed using a phylogenetic generalized least squares (PGLS) approach in the R packages ape^27^ (version 5.0, correlation structure = corBrownian) and nlme^28^ (version 3.1-131.1). Given that the EB models best fit the data, we also tested a corBlomberg correlation structure, which invokes an accelerating-decelerating model of evolution, with the decelerating rate of trait change fixed at 1.3.

For comparisons performed at the genus level, each regression was repeated over 100 trees randomly drawn from the posterior distribution randomly selecting a representative egg database entry per genus. For comparisons performed at the family level, each regression was repeated 100 times calculating the family level average egg data from 50% of entries per family.

For phylogenetic regressions controlling for a third variable, we calculated the phylogenetic residuals of each variable against the dependent variable, and then calculated the phylogenetic regression of the residuals^29^. To test alternative hypotheses, new data were simulated using a fixed scaling exponent and the parameters of the best fitting model with the R package phylolm^30^ (version 2.5, function ‘rTrait’).

Allometric regressions were performed over all insect taxa as well as for seven monophyletic groups of insects individually (Palaeoptera, Polyneoptera, Condylognatha, Hymenoptera, Neuropteroidea, Amphiesmenoptera, Antliophora). In addition, the scaling exponent between egg length and width was calculated for each monophyleti group of taxa with more than 20 tips but fewer than 50.

### 6.8 Statistical Information

For evolutionary regressions and parametric bootstraps, a significance threshold of 0.01 was used. All p-values were rounded to the nearest hundredth. Exact values for all statistical comparisons are available in the figure legends and Supplemental Information. For evolutionary model comparisons, weighted AICc values were compared at a significance threshold of 2. Evolutionary regressions were performed 100 times each, taking into account phylogenetic and phenotypic uncertainty. For more details see Supplemental Information, section “Calculating allometric exponents using phylogenetic generalized least squares (PGLS)”.

