## Supplementary material for "Insect egg size and shape evolve with ecology, not developmental rate"

#### *Supplemental Information*

Samuel H. Church[*,1,†], Seth Donoughe[*,1,2], Bruno A. S. de Medeiros[1], Cassandra G. Extavour[1,3,†]

This document includes a more complete description of the diversity of insect eggs, ancestral state reconstructions, evolutionary model fitting results, and additional methodological details.

* Samuel H. Church and Seth Donoughe contributed equally to this work.
1 Department of Organismic and Evolutionary Biology, Harvard University, Cambridge, MA 02138, United States
2 *Current address:* Department of Cell and Molecular Biology, University of Chicago, Chicago, IL 60637, United States
3 Department of Molecular and Cellular Biology, Harvard University, Cambridge, MA 02138, United States

8 **Summary of Phylogenetic Generalized Least Squares (PGLS) results** **41**
**References** **43**

### 1 The insect egg database

A complete description of the methods used to compile the insect egg database is published in an accompanying article[1]. Here, we briefly summarize the procedures used to assemble it. Descriptions of insect eggs were assembled from the published entomological literature using custom bioinformatic software. A list of the 1,756 literature sources used to generate the egg database is provided as a supplemental file, 'bibliography_egg_database.pdf'. Each entry in the database includes an insect's genus name and, if it was available from the source publication, the species name. Every entry also includes text measurements of egg dimensions and/or a published image of an insect egg. Published images were subsequently measured using custom software to extract additional egg size and shape information. Taxonomic names were checked against databases for synonyms and matched to online sequence databases for building phylogenies, using the software TaxReformer[1]. The database has been statistically assessed for accuracy of measurement tools as well as potential sources of variation (e.g. intraspecific variation and variation across publications), and the results of those assessments are also included in the publication describing the database[1].

#### 1.1 Insect lineages represented in the egg database

The insect egg database includes 10,449 insect egg descriptions comprising all extant insect orders, listed in Figure S1. For many analyses and visualizations, these orders were categorized into the nine groups shown. With the exception of the paraphyletic Apterygota, each of these groups has been supported as monophyletic in recent phylogenetic analyses of insect evolution[2,3]. See Section 2 for further information on how phylogenetic relationships were inferred in this study.

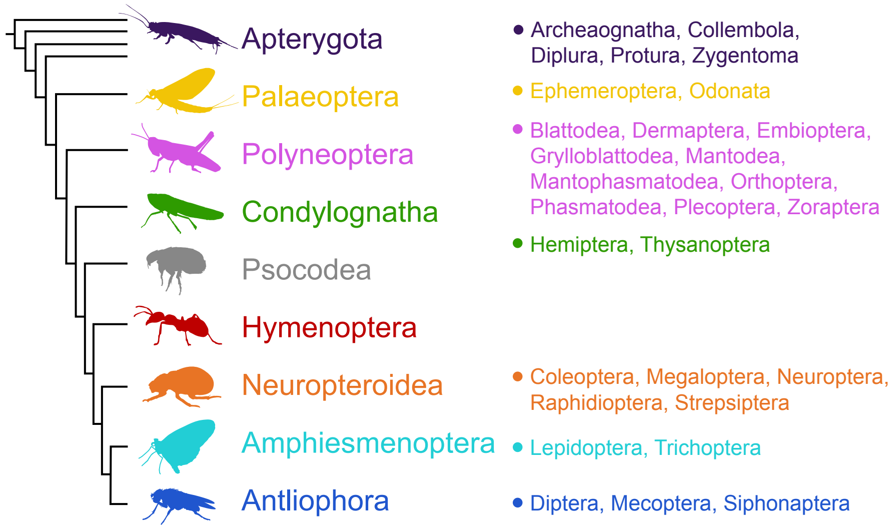

Figure S1: **Cladogram with constituent taxonomic orders.** The insect egg database includes egg descriptions from representatives of all extant insect orders, listed here. The relationships between insect clades are shown according to one of the two backbone phylogenies used to examine egg evolution in this study[2]. According to this backbone, the nine clades shown here represent monophyletic lineages with the exception of Apterygota, which is paraphyletic with respect to all other insects.

#### 1.2 Defining egg traits

The trait descriptions in this section are reproduced from the accompanying article that describes the insect egg database[1]. For each trait listed below we used the descriptions of egg length and width as presented in the original publications. Given that conventions vary across entomologists and insect taxonomic groups, we present the following definitions to resolve ambiguous cases and to serve as a suggestion for future egg descriptions.

*Egg:* The term *egg* is used to describe several successive developmental stages, including the mature oocyte, the zygote cell, and the developing embryo in its eggshell. For consistency we selected measurements that were recorded closest to the time of fertilization, when multiple descriptions were available within a single publication, given that in some insects it has been documented that the dimensions of the egg change over time (typically $<$20% change in length due to water exchange during embryonic development)[4–8]. In most insects the egg is oviposited outside the adult body, however in viviparous insects, eggs proceed through some or all of embryonic development within the body of the mother. The egg is often enveloped in a secreted eggshell called the chorion[8], which may have elaborations (e.g. dorsal appendages or opercula)[9]. We selected egg measurements that excluded chorionic elaborations over those that included them, as our goal was to measure the comparable cellular material across species.

*Length:* To resolve ambiguous cases, and when measuring egg features from images, we defined egg length as the distance in millimeters (mm) of the axis of rotational symmetry. This definition maximizes consistency with published descriptions of egg length. Under this definition, length is not always longer than width (as defined below). For some insect groups (e.g. Lepidoptera) the axis of rotational symmetry is sometimes referred to in the literature as *height*[10–12]. For published images with a scale bar, we measured both the straight and curved length of the egg (for those eggs that are curved), but for all analyses and figures, we used the straight length of the egg in order to maximize consistency with published records. Further details on how egg traits were measured from egg images are also available[1].

*Width* and *breadth:* To resolve ambiguous cases, and when measuring egg features from images, we defined width as the widest diameter (mm), measured perpendicular to the axis of rotational symmetry of the egg. For some insect groups this axis is referred to in the literature as diameter[10] or breadth[13]. For eggs described in published records as having a length, width, and breadth or depth (i.e., the egg is a flattened ellipsoid[14]), we considered *width* as the wider of the two diameters, and *breadth* as the diameter perpendicular to both width and length. For published images with a scale bar, we measured width as the widest of the three egg diameters at the first quartile, midpoint, and third quartile of the length axis. We did not measure breadth from published images.

*Volume:* Volume ($mm^3$) was calculated using the equation for the volume of an ellipsoid, following previous studies[15,16]. The formula is $\frac{1}{6}\pi lwb$, with $l$, $w$, and $b$ as length, width, and breadth, respectively. This simplifies to $\frac{1}{6}\pi lw^2$ when the egg is rotationally symmetric. For records in which the volume was reported but egg length and width were not, we used the reported volume. For all other entries, we recalculated volume from the measurements in the text and from measurements of images published with a scale bar.

*Aspect ratio:* We calculated aspect ratio as the ratio of length to width. An aspect ratio of one corresponds to a spherical egg. An aspect ratio less than one corresponds to an egg that is wider than long (oblate ellipsoid). An aspect ratio greater than one corresponds to an egg that is longer than it is wide (prolate ellipsoid). We note that egg *ellipticity* has been used to describe egg shape in birds[17,18] and that it can be calculated as the aspect ratio minus one. Analyses testing the sensitivity of our measurement software for egg images indicated that the variance in measured aspect ratio is highest for eggs with extremely high aspect ratios[1]. Therefore we excluded the eggs in the top 0.1 percentile of aspect ratio from subsequent analyses. We recorded the aspect ratio from images published with or without a scale bar, as aspect ratio is a scale-free attribute.

*Asymmetry:* We defined asymmetry as $\frac{\max(q_1,q_3)}{\min(q_1,q_3)} - 1$, where $q_1$ and $q_3$ are the egg diameters at the first and third quartile of the curved length axis. Therefore an egg with an asymmetry of zero has quartile diameters with equal length. Baker's $\lambda$ value, used to measure asymmetry in bird eggs[18], can be converted to the asymmetry parameter used in the present study (as shown in Fig. S5). Analyses testing the sensitivity of our image measuring software indicated that the variance is highest for eggs with extremely high values of asymmetry[1]. We therefore excluded the eggs in the top 0.1 percentile of asymmetry from subsequent analyses. Asymmetry was only recorded from published egg images.

*Angle of curvature:* We defined the angle of egg curvature as the angle of the arc created by the endpoints and midpoint of the length axis (measured in degrees). Analyses testing the sensitivity of our image measuring software indicated that the variance in curvature increases when the curvature and aspect ratio are low[1]. We therefore did not calculate curvature for eggs with an aspect ratio of one or less. Angle of curvature was only recorded from published egg images.

### 2 Estimating phylogenetic relationships

#### 2.1 Obtaining genetic data for genera in the egg database

While there are published order-level[2] and family-level[3] transcriptomic phylogenies for insects, to our knowledge there is no tree that includes all genera for which we assembled egg data. To address this, we produced a new tree by combining publicly available sequence data for the genera in the egg database with phylogenetic results from published insect evolutionary studies[2,3]. Sequence data for 18S and 28S ribosomal RNA were obtained from the SILVA database[19–22], a curated set of sequences for both the small and large ribosomal units. Sequences classified as Hexapoda in the SILVA database (release 128) were downloaded and associated with the corresponding National Center for Biotechnology Information (NCBI) ID using NCBI Entrez tools implemented in Biopython[23]. Each NCBI ID was then searched on the Open Tree of Life Taxonomy (OTT) to obtain the corresponding OTT ID. Finally, the dataset was curated by keeping only the longest sequence available for each species and genus in the egg database.

#### 2.2 Verifying sequence identity

To avoid inclusion of mislabeled or uninformative sequences, the data downloaded from the SILVA database were filtered using a phylogenetic criterion. First we created a reference dataset of ribosomal RNA sequences from the

Misof et al. (2014)[2] phylotranscriptomic dataset, as downloaded from the NCBI Sequence Read Archive (SRA). Raw reads from the SRA were filtered with Trimmomatic (version 0.32)[24] and mapped to the SILVA sequences to identify ribosomal RNA, using bowtie2 (version 2.2.9)[25]. Ribosomal RNA reads were assembled using Trinity, and the identity of the longest assembled contig was checked using BLAST in NCBI. For some taxa in the Misof et al. (2014) read archive we could not assemble rRNA sequences *de novo*, and in these cases we used reference sequences from the SILVA database which corresponded to the same genus in the Misof et al. (2014) tree.

Next, the reference sequences were aligned using MAFFT (version 7.245)[24] with the E-INS-I algorithm in Geneious (Biomatters) and trimmed manually. Reference alignments used for 18S and 28S are available at `https://github.com/shchurch/Insect_Egg_Evolution`, directory 'phylogeny'. Candidate SILVA sequences that matched an insect genus in the egg database by name were added to this reference alignment with MAFFT (option `--keeplength`, `--addlong`, and the default alignment algorithm). Aligned candidate SILVA sequences were placed on the Misof et al. (2014)[2] phylogeny using the evolutionary placement algorithm implemented in RAxML (version 8.2.9[24,26]). Candidate sequences that were placed in a different insect order on the phylogeny than the order reported in OTT were removed from the dataset.

#### 2.3 Estimating phylogenies for insect orders

Ribosomal RNA sequences that passed the filtering criteria were aligned using UPP (version 4.3.1)[27,28], which allowed us to align thousands of full and partial rRNA sequences. Bayesian clock models require an impractical computation time when applied to hundreds or thousands of sequences. Therefore, instead of performing a single phylogenetic analysis for all insects in our study, we divided the alignment into taxonomic orders and inferred a distribution of trees for each order individually (all orders included here have been recovered as monophyletic in previous studies[2]). For each order-level alignment, one outgroup sequence was included from every other order. The representative outgroup sequences were randomly selected from those taxa with both 18S and 28S sequences, or from taxa with 18S when both sequences were not available for a given order.

Order-level alignments were trimmed by removing regions at the alignment margins with less than 20% of the species included. Internal sites of the alignment that were represented by fewer than ten sequences were also removed, as were sequences with fewer than 100 total unambiguous sites. It has been shown that trimming of sites with excessive amounts of missing data speeds up computation and does not interfere with phylogenetic inference, while more aggressive criteria for trimming usually results in lower-quality trees[29]. After trimming, ribosomal genes were concatenated into a single dataset (see Table S2 for statistics on genera with sequence data included in this study).

For each alignment we generated a distribution of phylogenetic trees under a Bayesian framework with MrBayes (version 3.2.6[30]). Alignments were partitioned by gene, using a general time-reversal (GTR) model[31] with invariant sites and gamma rate variation applied to each partition. Molecular clock rates were allowed to vary according to the Independent Gamma rates model. The birth-death model was used for the tree topology prior, with speciation and extinction priors derived from previous inferences of diversification rate for insects based on the Rainford et al. (2014) tree[32].

Insect families that are present in the Rainford et al. (2014) study and are considered monophyletic on the Open

| Evolutionary placement | Misof et al. (2014)[2] | Open tree taxonomy | SILVA |
|---|---|---|---|
| Archaeognatha | Archaeognatha | Archaeognatha | Archaeognatha |
| Coleoptera | Coleoptera | Coleoptera | Coleoptera |
| Collembola | Collembola | Collembola | Collembola |
| Dermaptera | Dermaptera | Dermaptera | Dermaptera |
| **Dictyoptera** | **Blattodea, Isoptera** | **Blattodea** | **Blattodea, Isoptera** |
| Diplura | Diplura | Diplura | Diplura |
| Diptera | Diptera | Diptera | Diptera |
| Embioptera | Embioptera | Embioptera | Embioptera |
| Ephemeroptera | Ephemeroptera | Ephemeroptera | Ephemeroptera |
| Grylloblattodea | Grylloblattodea | Grylloblattodea | Grylloblattodea |
| Hemiptera | Hemiptera | Hemiptera | Hemiptera |
| Hymenoptera | Hymenoptera | Hymenoptera | Hymenoptera |
| Lepidoptera | Lepidoptera | Lepidoptera | Lepidoptera |
| Mantodea | Mantodea | Mantodea | Mantodea |
| Mantophasmatodea | Mantophasmatodea | Mantophasmatodea | Mantophasmatodea |
| Mecoptera | Mecoptera | Mecoptera | Mecoptera |
| Megaloptera | Megaloptera | Megaloptera | Megaloptera |
| Neuroptera | Neuroptera | Neuroptera | Neuroptera |
| Odonata | Odonata | Odonata | Odonata |
| Orthoptera | Orthoptera | Orthoptera | Orthoptera |
| Phasmatodea | Phasmatodea | Phasmatodea | Phasmatodea |
| Plecoptera | Plecoptera | Plecoptera | Plecoptera |
| Protura | Protura | Protura | Protura |
| **Psocodea** | **Psocodea** | **Phthiraptera, Psocoptera** | **Phthiraptera, Psocoptera** |
| Raphidioptera | Raphidioptera | Raphidioptera | Raphidioptera |
| Siphonaptera | Siphonaptera | Siphonaptera | Siphonaptera |
| Strepsiptera | Strepsiptera | Strepsiptera | Strepsiptera |
| Thysanoptera | Thysanoptera | Thysanoptera | Thysanoptera |
| Trichoptera | Trichoptera | Trichoptera | Trichoptera |
| Zoraptera | Zoraptera | Zoraptera | Zoraptera |
| **Zygentoma** | **Zygentoma** | **Zygentoma** | **Lepismatidae, Thysanura** |

Table S1: **Equivalence of order-level taxonomic concepts of main sources of data.** Differences in taxonomic names between data sources are shown in bold.

| Order | Genera with OTT IDs in egg database | Genera with 18S data | Genera with 28S data | Genera with any rRNA data |
|---|---|---|---|---|
| Archaeognatha | 2 | 2 | 2 | 2 |
| Coleoptera | 381 | 255 | 171 | 279 |
| Collembola | 8 | 7 | 6 | 7 |
| Dermaptera | 7 | 5 | 1 | 5 |
| Dictyoptera | 74 | 52 | 45 | 57 |
| Diplura | 3 | 3 | 3 | 3 |
| Diptera | 271 | 114 | 186 | 207 |
| Embioptera | 2 | 1 | 1 | 1 |
| Ephemeroptera | 76 | 62 | 42 | 62 |
| Grylloblattodea | 2 | 2 | 2 | 2 |
| Hemiptera | 431 | 185 | 139 | 215 |
| Hymenoptera | 635 | 341 | 338 | 394 |
| Lepidoptera | 1025 | 69 | 89 | 136 |
| Mantodea | 6 | 6 | 6 | 6 |
| Mantophasmatodea | 3 | 2 | 3 | 3 |
| Mecoptera | 7 | 3 | 3 | 3 |
| Megaloptera | 6 | 2 | 1 | 2 |
| Neuroptera | 32 | 20 | 2 | 20 |
| Odonata | 53 | 42 | 48 | 49 |
| Orthoptera | 184 | 89 | 66 | 93 |
| Phasmatodea | 113 | 45 | 58 | 61 |
| Plecoptera | 71 | 61 | 40 | 63 |
| Protura | 2 | 2 | 2 | 2 |
| Psocodea | 29 | 17 | 2 | 17 |
| Raphidioptera | 1 | 1 | 0 | 1 |
| Siphonaptera | 14 | 11 | 11 | 12 |
| Strepsiptera | 5 | 2 | 0 | 2 |
| Thysanoptera | 7 | 4 | 2 | 4 |
| Trichoptera | 23 | 13 | 3 | 14 |
| Zoraptera | 1 | 1 | 1 | 1 |
| Zygentoma | 2 | 2 | 2 | 2 |

Table S2: **Number of genera from egg dataset included in DNA sequence alignments.**

Tree of Life (OTL) were constrained to be monophyletic in our phylogenetic analysis. We used "soft constraints" for these relationships, meaning that taxa were allowed to be placed freely within an order if they belonged to families that were not considered monophyletic on the OTL or were not present in the Rainford et al. (2014) study. Insect orders, excluding the outgroup sequences, were also constrained to be monophyletic. We used the estimated divergence times from the Misof et al. study (2014) as the calibration time between orders, while nodes within each order were not time calibrated.

For each alignment we ran two metropolis-coupled Markov chains, with four chains per run, for at least 25 million generations, removing the first 25% of trees as burn-in. Convergence was assessed by the standard deviation of split frequencies between the two runs and by checking traces for each parameter in Tracer[33]. Convergence was achieved (<0.05 split frequencies) for all orders except Diptera and Hymenoptera (Table S3 and S4). Inspection of trace files showed a lack of long-term trends in the post-burn-in samples, indicating that the region of highest posterior probability was reached, but mixing was slow for these large trees. In subsequent analyses we used a random sample drawn from the posterior distribution to account for variation in relationships.

#### 2.4 Building the backbone phylogenies

We incorporated the results from published phylotranscriptomic studies of insects[2,3] into our analysis using phylogenetic backbones. For each order, a random sample of 100 trees from the posterior distribution was grafted onto one of two alternative backbone phylogenies, one from Misof et al. (2014) and one from Rainford et al. (2014), which differ in both the inferred relationships between orders (shown in Fig. S2) and the estimated divergence times. This grafting approach is similar to that used to infer other large-scale phylogenies, such as those for seed plants[34] and birds[35]. Maximum clade credibility trees (MCC) were also generated by grafting the MCC tree of each order to the corresponding backbone (Fig. S3). The final 100 trees drawn from the posterior distributions are available as the supplementary files 'phylogeny_posterior_distribution_misof_backbone.nxs' and 'phylogeny_posterior_distribution_rainford_backbone.nxs'.

Hereafter we refer to the resulting genus-level trees as the "Misof backbone tree" and the "Rainford backbone tree". All primary figures and tables shown in this study are based on the Misof backbone tree, with a comparison of the results between the backbones given in Table S22.

Rainford et al. (2014)[3] topology and divergence times were readily available from their supplementary data, but Misof et al. (2014)[2] provided divergence times only as a table, with a time tree included as a figure. We added the Misof backbone[2] divergence times as annotations to their respective nodes in a new cladogram. A tree file containing these annotations is included in the present study at `https://github.com/shchurch/Insect_Egg_Evolution` file 'fully_annotated_misof.nexml'.

Tree and alignment manipulations throughout the pipeline were done by custom bash and python scripts using Biopython[23] and Dendropy[36]. Some steps of this pipeline made use of GNU parallel[37].

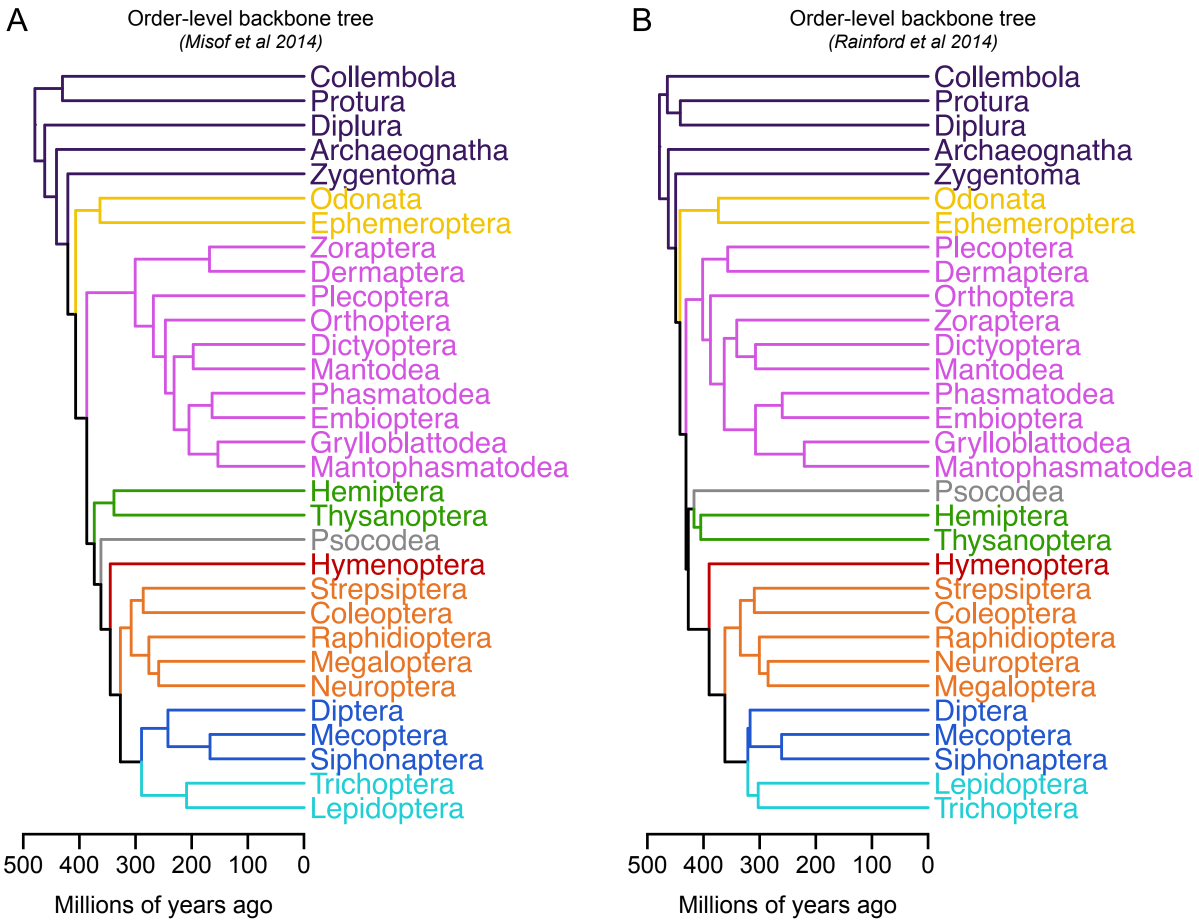

Figure S2: **Relationships between insect orders in two backbone phylogenies. A**, Order-level backbone tree from Misof et al. (2014)[2]. **B**, Order-level backbone tree from Rainford et al. (2014)[3].

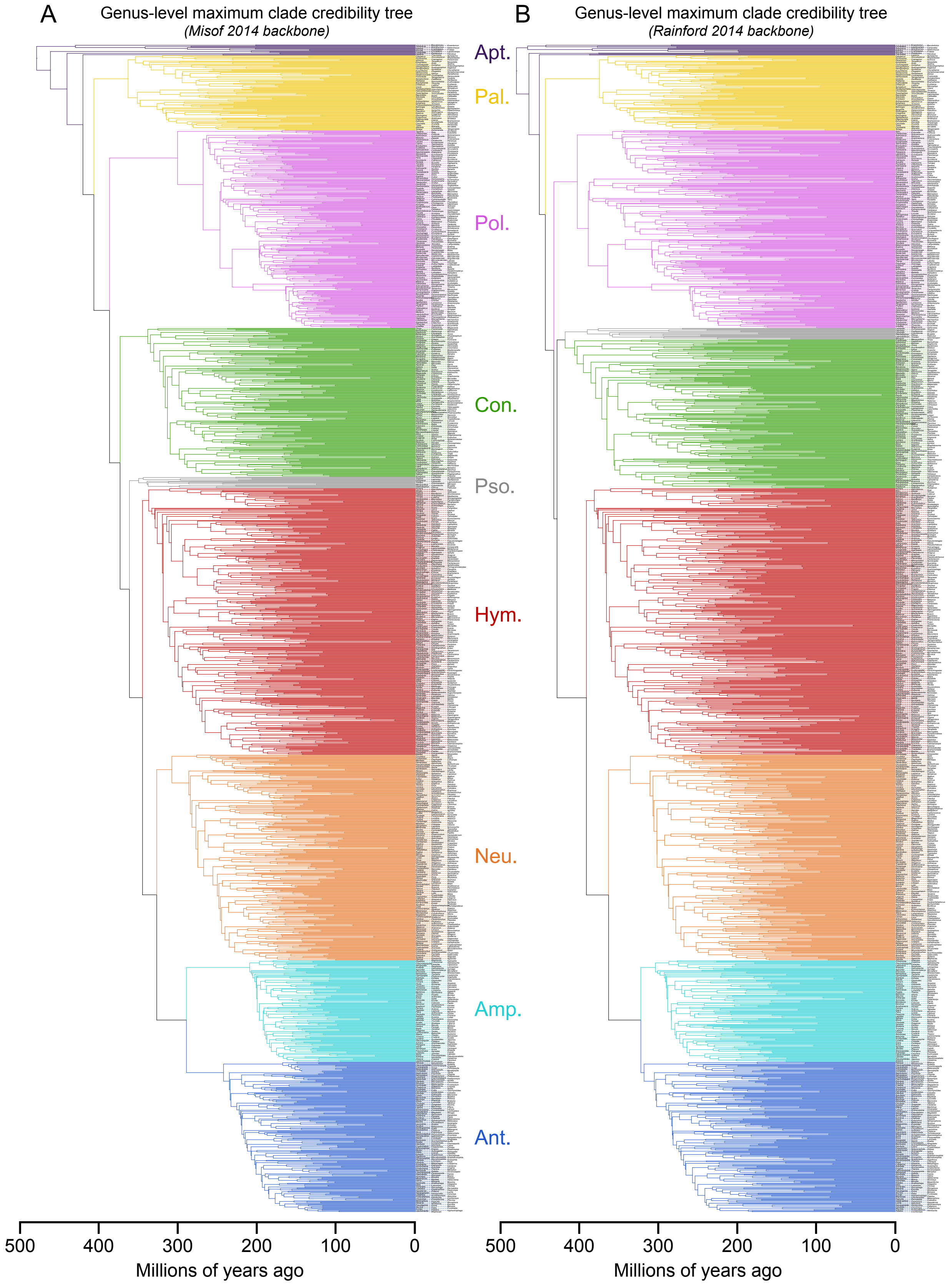

Figure S3: **Maximum clade credibility hexapod phylogenies. A**, Genus-level maximum clade credibility tree with the backbone tree from Misof et al. (2014) fixed as a constraint[2]. **B**, Genus-level maximum clade credibility tree with the backbone tree from Rainford et al. (2014) fixed as a constraint[3]. Colors correspond to the clades shown in Fig. S1

| Taxon | Post-burn-in samples | ESS (likelihood) | ESS (prior) | Standard deviation of split frequencies | Number of tips in ingroup |
|---|---|---|---|---|---|
| Archaeognatha | 9651 | 8060 | 1748 | 0.000000 | 2 |
| Coleoptera | 42800 | 115 | 338 | 0.035881 | 279 |
| Collembola | 13691 | 10381 | 1126 | 0.000000 | 7 |
| Dermaptera | 10818 | 8932 | 1001 | 0.000523 | 5 |
| Dictyoptera | 3936 | 1046 | 145 | 0.008308 | 57 |
| Diplura | 9781 | 4594 | 730 | 0.000000 | 3 |
| Diptera | 334040 | 9 | 122 | 0.106535 | 207 |
| Ephemeroptera | 16908 | 6216 | 460 | 0.009309 | 62 |
| Grylloblattodea | 10810 | 4882 | 997 | 0.000000 | 2 |
| Hemiptera | 45710 | 1793 | 133 | 0.030570 | 216 |
| Hymenoptera | 29350 | 1110 | 179 | 0.070895 | 394 |
| Lepidoptera | 75002 | 3464 | 379 | 0.007657 | 136 |
| Mantodea | 8186 | 6384 | 607 | 0.001289 | 6 |
| Mantophasmatodea | 8444 | 7116 | 916 | 0.000061 | 3 |
| Mecoptera | 8470 | 7324 | 1127 | 0.000000 | 3 |
| Megaloptera | 7000 | 7000 | 1178 | 0.000000 | 2 |
| Neuroptera | 6186 | 3526 | 435 | 0.006141 | 20 |
| Odonata | 4492 | 2065 | 63 | 0.004686 | 49 |
| Orthoptera | 2876 | 580 | 30 | 0.007835 | 93 |
| Phasmatodea | 4384 | 1358 | 85 | 0.009547 | 61 |
| Plecoptera | 9406 | 3002 | 222 | 0.003344 | 63 |
| Protura | 10720 | 10039 | 1346 | 0.000000 | 2 |
| Psocodea | 9876 | 7028 | 791 | 0.000035 | 17 |
| Siphonaptera | 6470 | 4478 | 264 | 0.002504 | 12 |
| Strepsiptera | 8738 | 7868 | 1563 | 0.000000 | 2 |
| Thysanoptera | 10050 | 9426 | 1428 | 0.000000 | 4 |
| Trichoptera | 7586 | 5074 | 675 | 0.000644 | 14 |
| Zygentoma | 13616 | 12581 | 2503 | 0.000000 | 2 |

Table S3: **Convergence statistics for phylogenetic analyses using Misof tree[2] as backbone.**

| Taxon | Post-burn-in samples | ESS (likelihood) | ESS (prior) | Standard deviation of split frequencies | Number of tips in ingroup |
|---|---|---|---|---|---|
| Archaeognatha | 8606 | 7797 | 1576 | 0.000000 | 2 |
| Coleoptera | 43136 | 535 | 218 | 0.052288 | 279 |
| Collembola | 8851 | 4629 | 538 | 0.000000 | 7 |
| Dermaptera | 10128 | 8026 | 879 | 0.000845 | 5 |
| Dictyoptera | 6980 | 2257 | 238 | 0.005348 | 57 |
| Diplura | 9398 | 8718 | 2003 | 0.000000 | 3 |
| Diptera | 32318 | 12 | 188 | 0.164839 | 207 |
| Ephemeroptera | 7646 | 2698 | 362 | 0.004788 | 62 |
| Grylloblattodea | 6344 | 5969 | 1578 | 0.000000 | 2 |
| Hemiptera | 41996 | 2691 | 263 | 0.011937 | 216 |
| Hymenoptera | 22260 | 83 | 60 | 0.058971 | 394 |
| Lepidoptera | 75002 | 4168 | 584 | 0.008016 | 136 |
| Mantodea | 6620 | 5096 | 811 | 0.000942 | 6 |
| Mantophasmatodea | 6380 | 5311 | 806 | 0.000497 | 3 |
| Mecoptera | 6248 | 4824 | 844 | 0.000000 | 3 |
| Megaloptera | 6926 | 5723 | 1242 | 0.000000 | 2 |
| Neuroptera | 12470 | 8112 | 1268 | 0.001025 | 20 |
| Odonata | 10322 | 4222 | 214 | 0.003670 | 49 |
| Orthoptera | 7424 | 1350 | 53 | 0.007249 | 93 |
| Phasmatodea | 10074 | 2499 | 223 | 0.008602 | 61 |
| Plecoptera | 4988 | 2154 | 87 | 0.003978 | 63 |
| Protura | 8076 | 6477 | 2138 | 0.000000 | 2 |
| Psocodea | 5406 | 3082 | 562 | 0.000017 | 17 |
| Siphonaptera | 5692 | 4250 | 569 | 0.000728 | 12 |
| Strepsiptera | 4234 | 3282 | 1179 | 0.000000 | 2 |
| Thysanoptera | 5486 | 5118 | 796 | 0.000000 | 4 |
| Trichoptera | 4908 | 3240 | 447 | 0.000576 | 14 |
| Zygentoma | 6126 | 5541 | 963 | 0.000000 | 2 |

Table S4: **Convergence statistics for phylogenetic analyses using Rainford tree[3] as backbone.**

### 3 Morphological diversity of insect eggs

#### 3.1 Distribution of egg traits within insect clades

To place the diversity of insect propagule sizes in context, we compared their distribution to a recently published study of eggs in birds[18], as well as to an estimated range of extant plant seed sizes (Fig. S4, panel A). We found that insect eggs range across eight orders of magnitude in volume, from $10^{-6}$ to $10^{2}$ $mm^3$. In comparison, bird eggs range across three orders of magnitude in volume, based on the largest (*Aepyornis sp.*, length 238 mm, width 164 mm) and smallest egg (*Hylocharis xantusii*, length 12.1 mm, width 8.0 mm) included in the Stoddard et al. (2017)[18] dataset. Angiosperm seed volumes range over more than 11 orders of magnitude, from the dust-seeds of orchids (*Paphiopedilum barbatum*, volume $5.69 * 10^{-5}$ $mm^3$)[38] to the giants seeds in palms (*Lodoicea maldivica*, length ~300 mm, width ~280 mm)[39]. Both birds and angiosperms are younger and less speciose than insects[2,40–42].

We also compared the distribution of shape parameters across insect groups. Egg aspect ratio is distributed heterogeneously across insect groups (Fig. S4 panel B). Insect eggs with an aspect ratio less than one (the egg is an oblate ellipsoid, that is, width is greater than length) have evolved and diversified in at least two main groups, Amphiesmenoptera (within Lepidoptera) and Polyneoptera (within Plecoptera). Across diverse groups of insects, some eggs were reported as exactly spherical (aspect ratio of 1). In the morphospace shown in Figure 1A, for example, these eggs form a conspicuous vertical alignment of datapoints for which aspect ratio equals exactly one. We attribute this pattern in the data to a tendency among researchers to describe near-spherical eggs as exactly spherical in cases when they did not measure length and width separately. Insect eggs vary with respect to aspect ratio to a much greater extent than bird eggs, affording an opportunity to test hypotheses about shape evolution across a greater diversity of possible shapes (Fig. S5).

Insect egg shapes vary considerably in the degree of asymmetry and the angle of curvature (Fig. S4, panels C and D). Like bird eggs, insect eggs range from completely symmetrical to highly asymmetrical, with extreme asymmetry found mainly in Hymenoptera and Condylognatha. Unlike in birds, insect eggs are often curved along the longitudinal axis of the egg. A high degree of curvature has evolved in Hymenoptera, Condylognatha, Antliophora, and Polyneoptera (specifically in the orders Hymenoptera, Hemiptera, Diptera, and Orthoptera).

#### 3.2 Insect egg morphospace

Recent work by Stoddard and colleagues showed that bird morphospace, defined by aspect ratio and asymmetry, was bounded such that no bird eggs are both asymmetrical and have an aspect ratio close to 1[18]. This is not true for insects (Fig. S5).

#### 3.3 Distribution of polyembryonic insects in egg morphospace

In polyembryonic insects, one egg develops into multiple embryos[43]. Observing that the smallest egg in the database is laid by a polyembryonic wasp, we collected records on polyembryony across insects and plotted their presence in

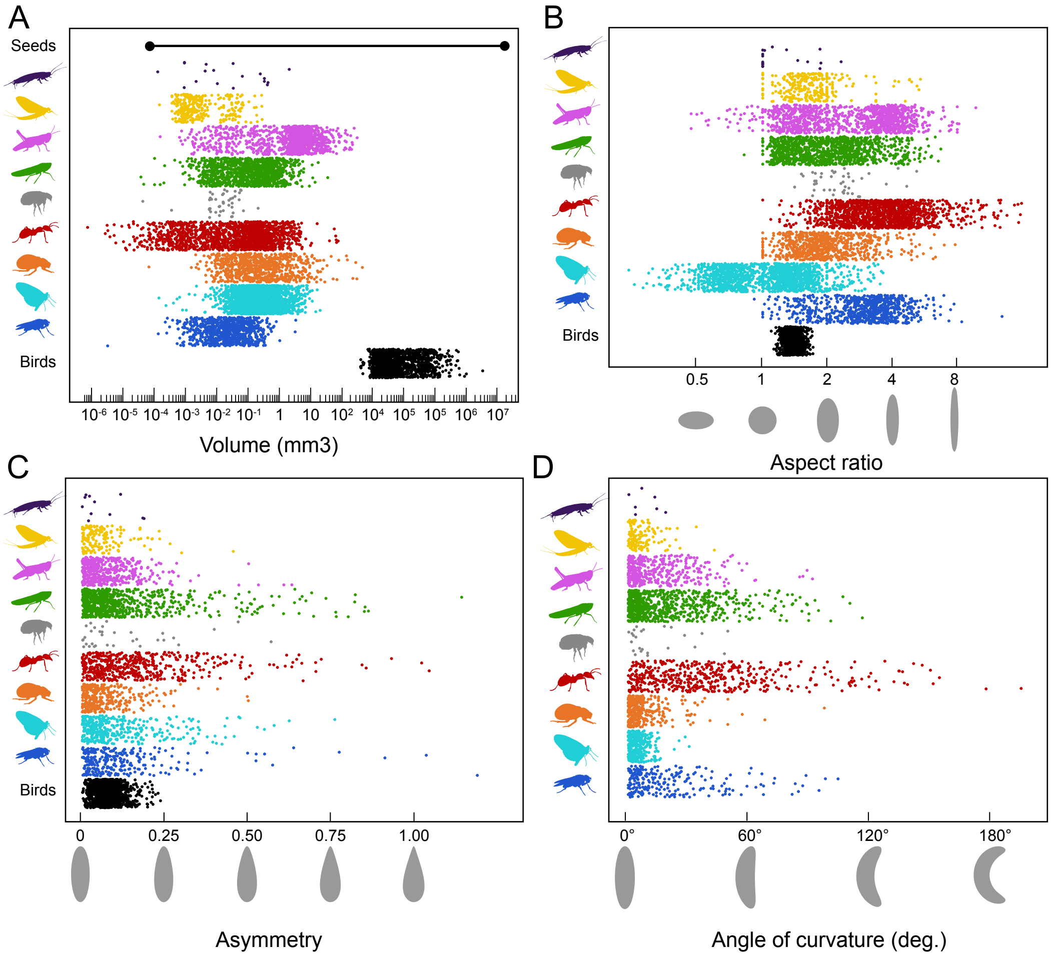

Figure S4: **Distributions of egg shape and size in insect groups.** In each panel egg traits are plotted by phylogenetic group on the y-axis (within a group, points are randomly spread vertically). All groups are monophyletic clades with the exception of Apterygota, which is paraphyletic with respect to all other insects. Colors correspond to the clades shown in Fig. S1. **A**, Egg volume ($mm^3$, log transformed) across insect clades and compared to the distribution of extant bird egg sizes[18] (bottom row) and the range of angiosperm seed sizes (top row). The lower bound of seed size is represented by the orchid *Acanthephippium sylhetense*[38]; the upper bound is represented by the palm *Lodoicea maldivica*[39]. **B**, Aspect ratio (unitless, log transformed) across insect clades and compared to the distribution of extant bird eggs[18]. **C**, Asymmetry (unitless) across insect clades and compared to the distribution of extant bird eggs[18]. **D**, Angle of curvature (degrees) across insect clades.

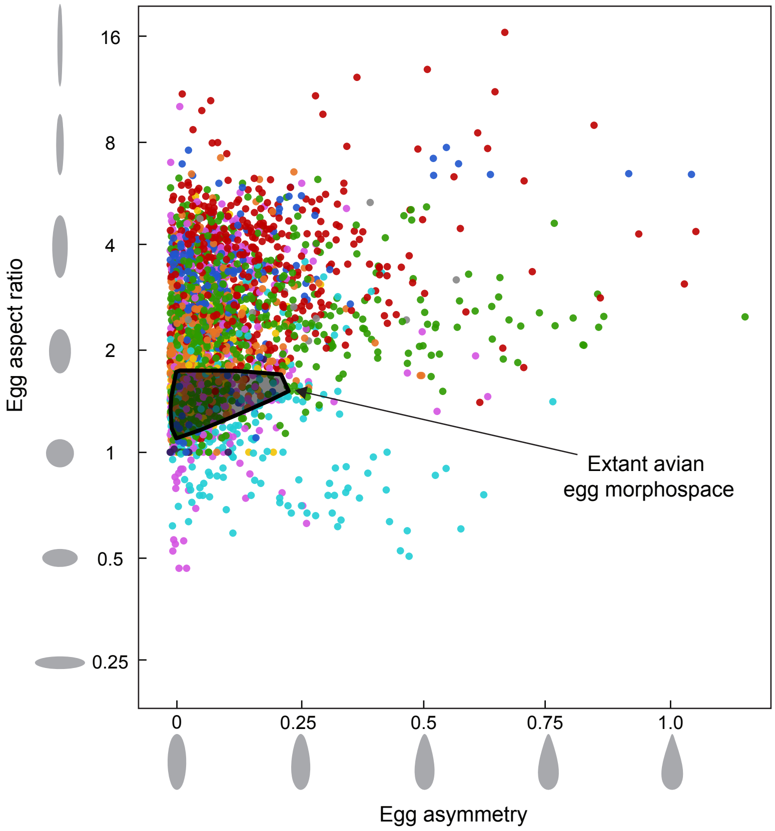

Figure S5: **Comparison of insect and bird egg morphospace occupancy.** The distribution of insect and avian eggs in the shape space defined by asymmetry and aspect ratio (plotted on a log scale). Both traits are unitless ratios. Points represent entries from the egg database, colored according to the clades defined in Fig. S1. The range of morphospace occupied by birds is shown in gray[18].

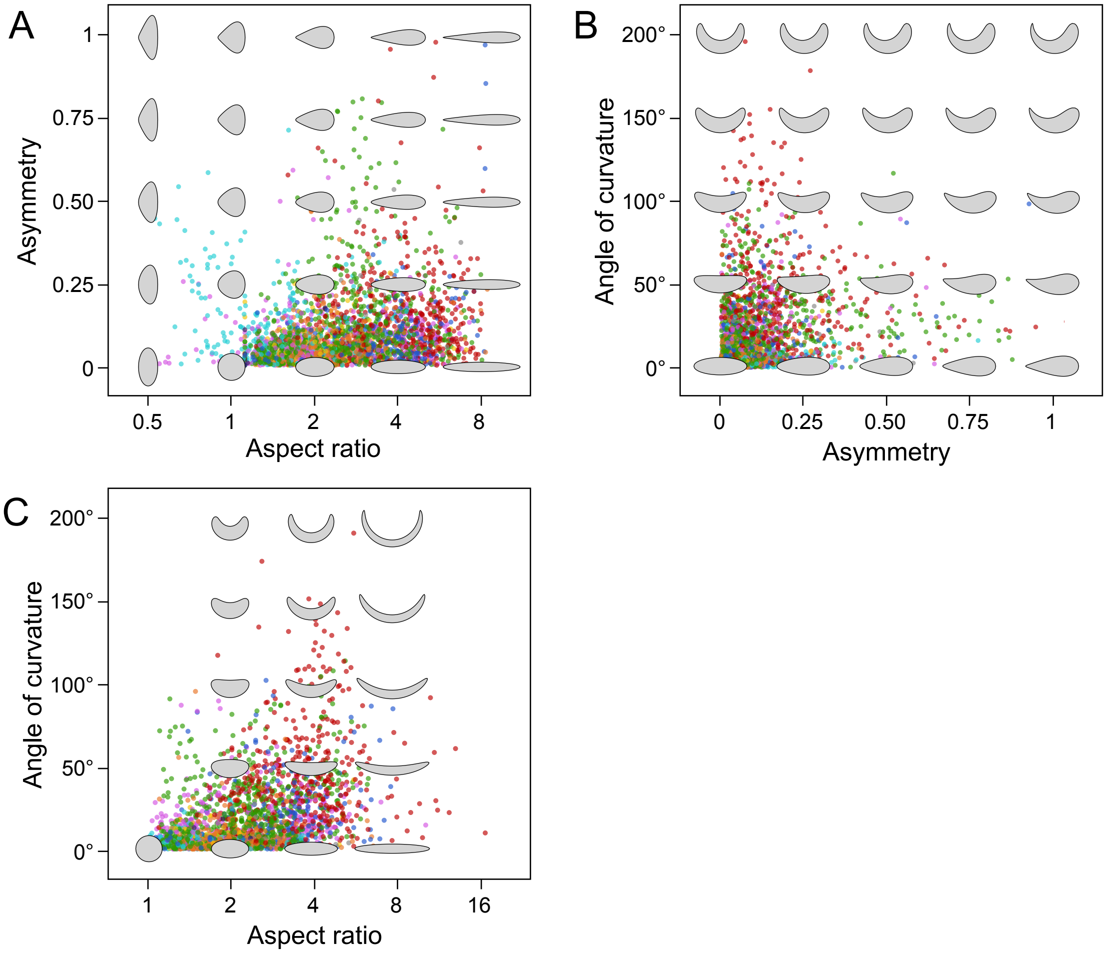

Figure S6: **Distributions of insects in egg morphospace.** The distribution of insect eggs in the shape space defined by **A** asymmetry and aspect ratio (log scale), **B** angle of curvature and asymmetry, and **C** angle of curvature and aspect ratio (log scale). Theoretical eggs are drawn as laterally oriented silhouettes in gray. The morphospace described by angle of curvature and aspect ratio is bounded at an aspect ratio of one, according to the definition of angle of curvature. See Section 1.2 for details.

insect egg morphospace[43–49]. Polyembryony has evolved at least five times in insects[43], four times in Hymenoptera and once in Strepsiptera. In those polyembryonic lineages for which we have egg shape and size data, we observe that all polyembryonic insects are among the smallest eggs (below $10^{-3}$ $mm^3$ in volume). We hypothesize that additional instances of polyembryony will be observed when detailed embryological studies are conducted on insect species that lay particularly small eggs.

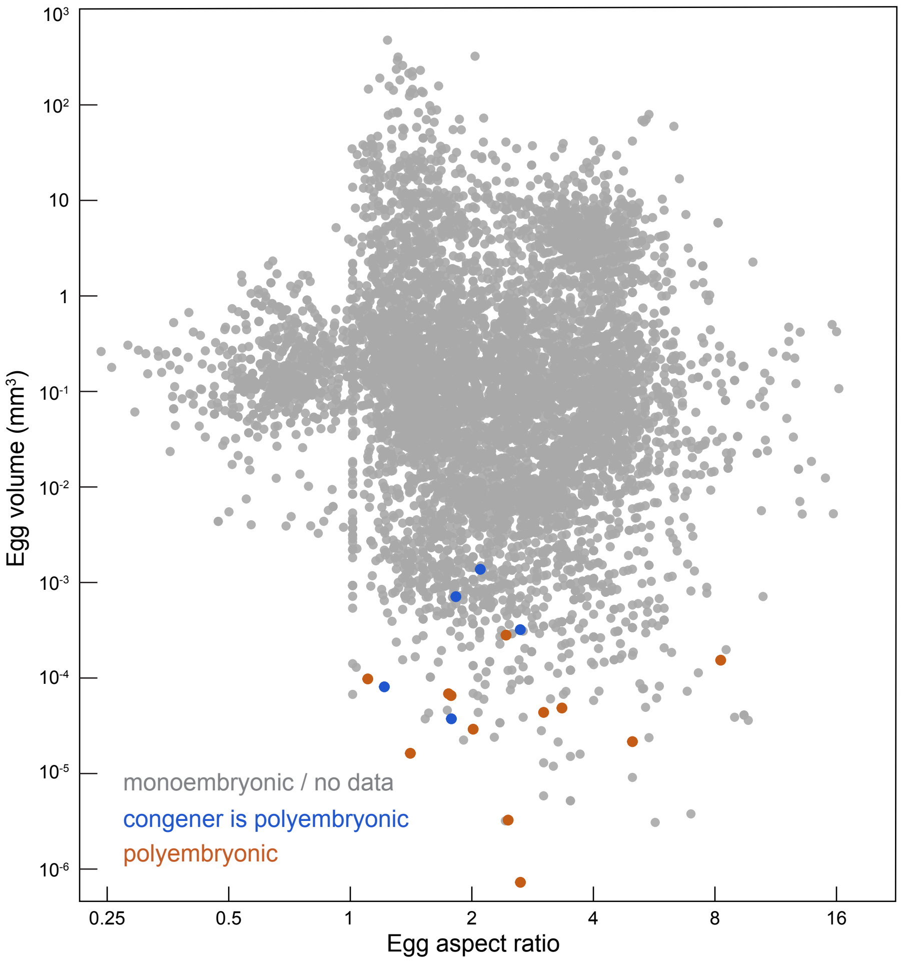

Figure S7: **Distribution of polyembryonic insects in egg morphospace.** The distribution of the eggs of polyembryonic insects (orange) and their congeners (blue) in the space defined by volume ($mm^3$) and aspect ratio (unitless). Both traits are plotted on a log scale. Gray points represent eggs of insects that develop monoembryonically or taxa for which no information on monoembryony vs. polyembryony was found.

### 4 Evolutionary history of egg traits

#### 4.1 Evolutionary model fitting

We compared models of evolution with respect to six parameters of egg size and shape using the R package geiger[50]. For each parameter we tested the fit of a Brownian motion model (BM), Ornstein-Uhlenbeck model (OU), Early-

Burst model (EB), and stochastic white-noise process (WN) using the Misof backbone[2] maximum clade credibility (MCC) phylogeny.

An Early-Burst model of evolution with a decreasing rate of evolution best explains ($\Delta$AICc > 2) the observed distributions of length, width, volume and aspect ratio (Table S5; $\alpha$ values for parameters as follows—log transformed, length: -0.005, width: -0.003, volume: -0.003, aspect ratio: 0.002). In contrast, for egg asymmetry and angle of curvature, OU and BM models are the best fit, respectively.

| | $\Delta$AICc, BM | $\Delta$AICc, OU | $\Delta$AICc, EB | $\Delta$AICc, WN |
|---|---|---|---|---|
| Volume | 51.29 | 53.30 | 0.00 | 1701.65 |
| Aspect Ratio | 15.38 | 17.39 | 0.00 | 1119.52 |
| Asymmetry | 33.96 | 0.00 | 35.99 | 82.30 |
| Curvature | 0.65 | 2.67 | 0.00 | 337.21 |
| Length | 72.04 | 74.05 | 0.00 | 1731.51 |
| Width | 28.75 | 30.76 | 0.00 | 1544.65 |
| Body Volume | 1.81 | 0.00 | 3.86 | 104.63 |
| Egg Volume - family | 0.00 | 2.03 | 2.05 | 163.10 |

Table S5: **Evolutionary model fitting results.** Comparing the fit ($\Delta$AICc) of evolutionary models, including Brownian Motion (BM), Ornstein-Uhlenbeck (OU), Early-Burst (EB), and stochastic white-noise process (WN).

In order to better test the fit of the Early-Burst model to our data, we performed a parametric bootstrap of the model using the R package arbutus[51] (Fig. S8). This package simulates 100 additional datasets using the optimized parameters of the model and compares six descriptive parameters from the observed dataset to the null distribution generated with simulation. The results of the six parameter comparisons are as follows:

1. **m.sig**: *mean of the squared contrasts*. The rate of evolution of both egg volume and aspect ratio is well estimated by the Early-Burst model (the observed value falls within the null distribution).

2. **c.var**: *coefficient of variation of the absolute value of the contrasts*. For both egg volume and aspect ratio there is additional rate heterogeneity, beyond the decreasing rate of evolution fit with the Early-Burst model, which is not well accounted for (the observed value falls well outside the null distribution).

3. **s.var**: *slope of a linear model fitted to the absolute value of the contrasts against their expected variances*. For both egg volume and aspect ratio, contrasts are smaller than expected based on their branch lengths, suggesting possible branch length error.

4. **s.asr**: *slope of a linear model fitted to the absolute value of the contrasts against the ancestral state at the corresponding node*. For egg volume there is no correlation between the rate of evolution and the state (larger eggs do not evolve faster). However, for aspect ratio, more elliptical eggs evolve faster, suggesting rate-state interactions.

5. **s.hgt**: *slope of a linear model fitted to the absolute value of the contrasts against node depth*. The Early-Burst model accounts well for the decreasing rate of evolution in the data.

6. **d.cdf**: *the D statistic from a Kolmogorov-Smirnov test comparing the distribution of contrasts to an expected normal distribution*. For both egg volume and aspect ratio the data do not fit a normal distribution of

contrasts well, suggesting there are likely non-Brownian motion based processes at play (e.g. jump-diffusion processes).

These results suggest that the Early-Burst model fits some aspects of the data well, specifically the overall rate of evolution and its deceleration over time. However, it also suggests a more complex evolutionary history than can be captured in this model alone, including additional rate heterogeneity, rate-state interactions, and possible jump-diffusion-like processes.

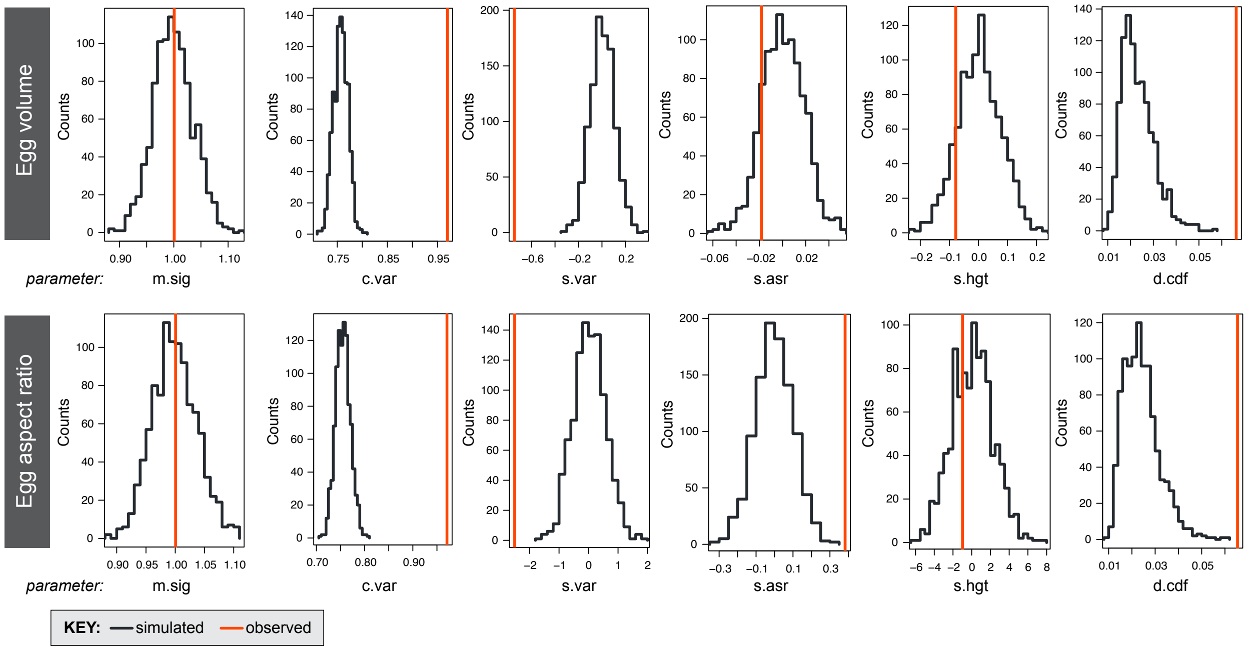

Figure S8: **Parametric bootstrap of the Early-Burst model for insect egg size and aspect ratio.** The results of a parametric bootstrap of the best fitting evolutionary model, the Early-Burst model, for egg volume ($mm^3$) and aspect ratio, calculated by the R package arbutus[51]. In each of the 6 panels, the observed statistic (red line) is compared to a null distribution generated from re-simulation (black distribution). See Section 4.1 for details on the interpretation of each parameter.

#### 4.2 Ancestral state reconstructions and evolutionary rate

Given the additional complexity in trait evolution suggested by the evolutionary model analyses, we explored the evolutionary history of egg size and shape further by reconstructing the ancestral state for the continuous traits egg volume, aspect ratio, asymmetry and the angle of curvature using the R package phytools[52] (version 0.6-44, function contMap). We also fit a rate regime map for each of these traits using the program BAMM in the R package BAMMtools (version 2.5.0) and setBAMMpriors (version 2.1.6). The prior for expected number of shifts was 10, with 10,000,000 generations. Consistent with the results of the model comparison, we observe that the rate of evolution for volume and aspect ratio generally decreases across insects, but that large shifts in rate have occurred multiple times. For example, there are dramatic increases in the rate of volume evolution in parasitoid Hymenoptera, and in the rate of aspect ratio evolution in Noctuoidea (Amphiesmenoptera: Lepidoptera).

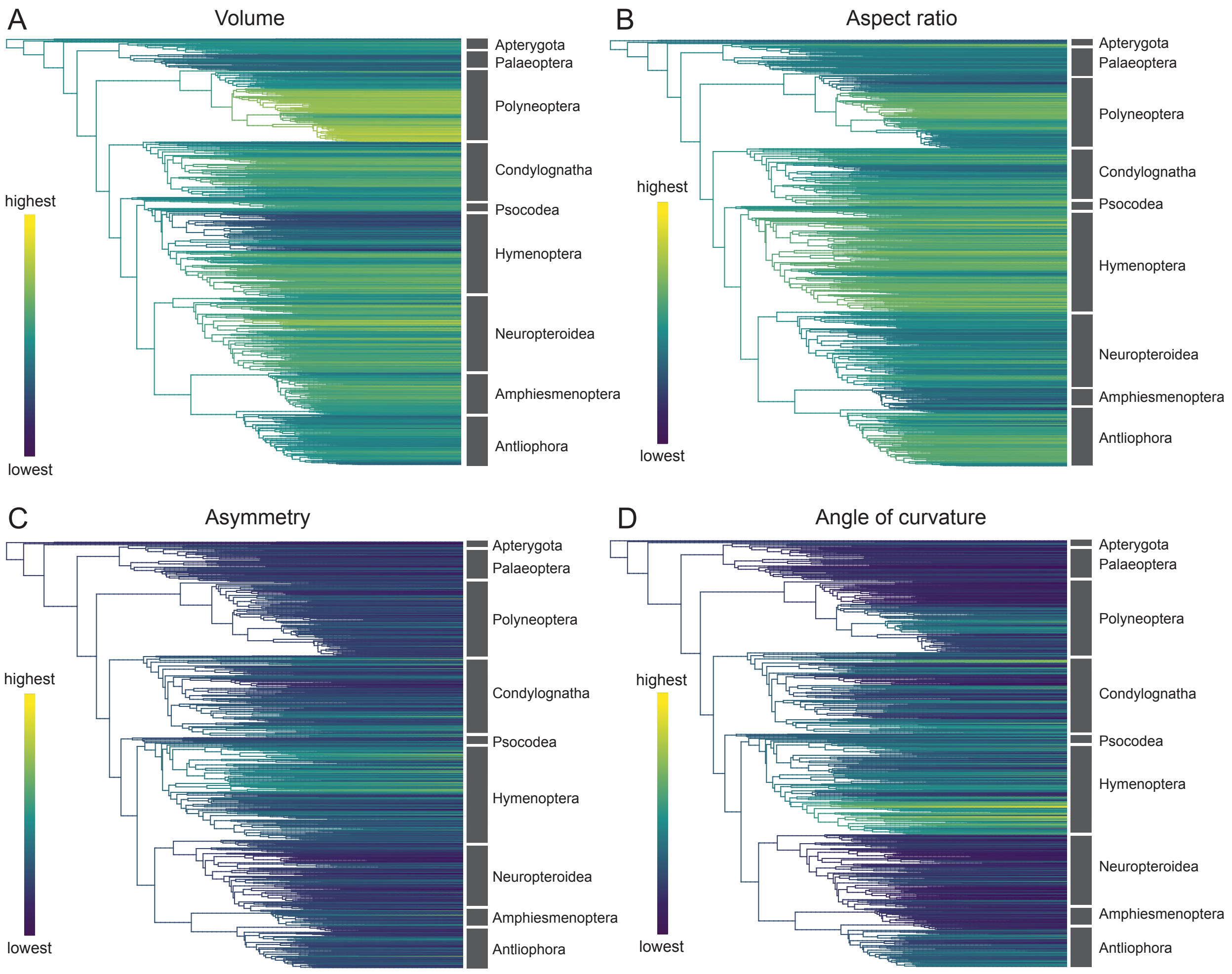

Figure S9: **Ancestral state reconstructions of egg morphological parameters.** Ancestral state reconstructions of **A** egg volume (mm$^3$; log scale), **B** aspect ratio (unitless; log scale), **C** asymmetry (unitless; square root scale), and **D** angle of curvature (degrees; square root scale). Low parameter values are shown in purple and high parameter values are shown in yellow.

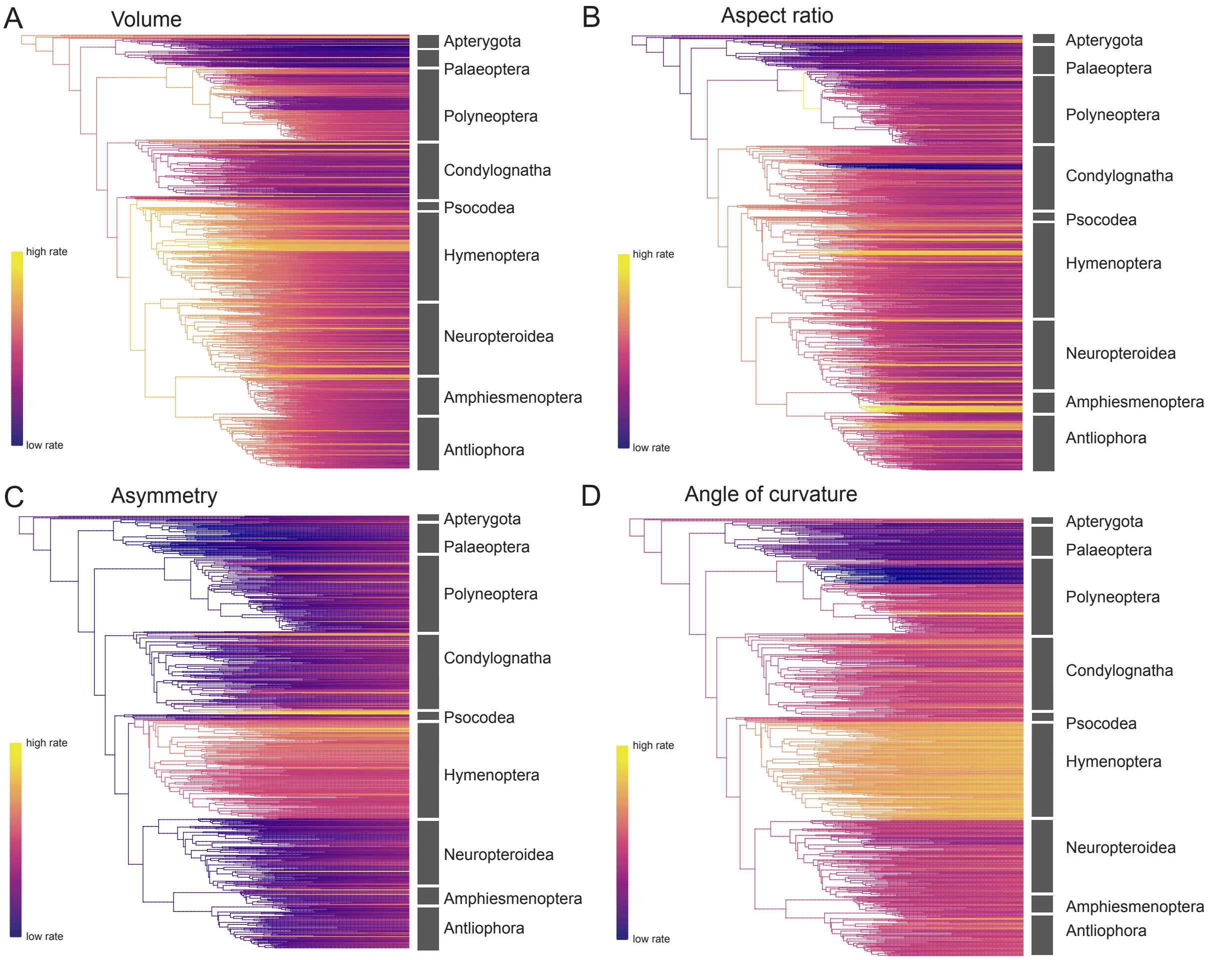

Figure S10: **Evolutionary rate regimes** Evolutionary rate regimes calculated with the software BAMM[53] of **A** egg volume ($mm^3$; log scale), **B** aspect ratio (unitless; log scale), **C** asymmetry (unitless; square root scale), and **D** angle of curvature (degrees; square root scale). Low rates are shown in purple and high rates are shown in yellow.

### 5 Allometric slopes of egg shape vary across insects

#### 5.1 Calculating allometric exponents using phylogenetic generalized least squares (PGLS)

Allometric relationships can be described using a power law, in which two traits $x$ and $y$ are related according to $y = bx^a$[54]. The scaling exponent $a$ can be estimated for a group of taxa as the slope of a regression between two continuous traits in log-log space, accounting for the non-independence of phylogenetically correlated data with a phylogenetic generalized least squares approach (PGLS)[55].

All PGLS comparisons were performed in R using the packages ape[56] (version 5.0) and nlme[57] (3.1-131.1). The principle findings of this paper were calculated using a Brownian-Motion based correlation structure (corBrownian). We also tested the robustness of results when using an Accelerating-Decelerating based correlation structure with a fixed decelerating rate of evolution (corBlomberg, g = 1.3) to approximate the Early-Burst model, which best describes egg size and aspect ratio evolution. For a comparison of PGLS results under these covariance matrices, see the summary in Section 8.

PGLS comparisons of egg size, shape, and developmental time were performed at the genus level over a posterior distribution of trees. The principle findings reported in this paper use the posterior distribution based on the Misof backbone phylogeny[2]. We also test the robustness of results to uncertainty in the backbone by using the posterior distribution based on the Rainford backbone[3]. For a comparison of PGLS results using these backbone phylogneis, see the summary in Section 8. For each iteration over the posterior distribution we selected a random representative entry per genus from the insect egg database. We therefore report the range of observed p-values, intercepts, and slopes (allometric exponents) accounting for both the phylogenetic and sampling uncertainty.

PGLS comparisons involving body size were performed at the family/order level using the published Rainford phylogeny[3]. To test the sensitivity of our results to sampling discrepancies between the egg database and published body size data, we downsampled the egg database by 50% and repeated each PGLS involving body size 100 times.

#### 5.2 Dynamic evolution of the allometry of egg shape and size

The results of a PGLS comparison between log-egg length and log-egg width show a significant allometric relationship with a slope less than one across insects. However, the scaling exponent of length vs. width varies considerably across insect lineages (Table S6, main text Fig. 2 and Fig. S12).

We compared our results to alternative hypotheses of size and shape evolution by simulating new egg length and width datasets under known models and analyzing them using the same methods. We tested two hypotheses: (1) that egg length and width have a 1:1 relationship (isometry), and (2) that egg length and width evolve independently. For each hypothesis we simulated data with the same parameters as the observed data (number and phylogenetic position of taxa, fitted evolutionary model parameters, EB model for both length and width) using the R package phylolm[58] (version 2.5; function 'rTrait'). The p-value of each hypothesis was calculated as the count of scaling exponents (the slope of the PGLS regression) that are more extreme than our test statistic, which is the median observed scaling exponent of each of the seven major insect groups analysed.

| | p-value | slope | intercept | sample size |
|---|---|---|---|---|
| Hexapoda | 0 - <0.005 | 0.77 - 0.80 | -0.28 - -0.25 | 1489 |
| Hymenoptera | 0 - <0.005 | 0.70 - 0.78 | -0.58 - -0.54 | 356 |
| Condylognatha | 0 - <0.005 | 0.78 - 0.91 | -0.42 - -0.39 | 202 |
| Antliophora | 0 - <0.005 | 0.65 - 0.78 | -0.47 - -0.43 | 199 |
| Neuropteroidea | 0 - <0.005 | 0.89 - 0.98 | -0.31 - -0.28 | 265 |
| Amphiesmenoptera | 0 - <0.005 | 0.71 - 0.92 | -0.20 - -0.13 | 76 |
| Polyneoptera | 0 - <0.005 | 0.69 - 0.75 | -0.28 - -0.25 | 262 |
| Palaeoptera | 0 - <0.005 | 0.61 - 0.74 | -0.45 - -0.38 | 104 |

Table S6: **Results of PGLS of egg length and width.**

Our results show that the first hypothesis, isometry, cannot be rejected for the lineage Neuropteroidea (beetles and relatives, p-value, isometry = 0.02, Fig. S11 and Table S7). The second hypothesis, that egg length and width evolve independently, can be rejected for all lineages (p-value, no relationship <0.01, out of 100 bootstraps, no values were greater than the test statistic)

| | test statistic | p-value, isometry | p-value, no relationship |
|---|---|---|---|
| Palaeoptera | 0.66 | <0.01 | <0.01 |
| Polyneoptera | 0.72 | <0.01 | <0.01 |
| Condylognatha | 0.86 | <0.01 | <0.01 |
| Hymenoptera | 0.73 | <0.01 | <0.01 |
| Neuropteroidea | 0.93 | 0.02 | <0.01 |
| Amphiesmenoptera | 0.79 | <0.01 | <0.01 |
| Antliophora | 0.73 | <0.01 | <0.01 |

Table S7: **Results of a parametric bootstrap of alternate hypotheses of egg shape and size evolution.** For parametric bootstraps, a p-value of <0.01 indicates that out of 100 bootstraps, no values were greater / less than the test statistic.

The seven lineages selected for comparison are large monophyletic lineages of insects, but it is also informative to estimate allometries for other clade divisions. To better represent the dynamic evolution of the scaling exponent, we broke down the insect phylogeny further. First, we identified the nodes in the phylogeny that had a sufficient number of descendant tips with morphological data to calculate the allometric exponent. We then identified the minimum number of unique nodes such that no node had more than 50 descendant tips. We repeated the PGLS comparison of log-egg length and log-egg width for each of these groups, and plotted the distributions of scaling exponents on the phylogeny (Fig. S12).

Our results show that additional subgroups of insects have a near-isometric relationship between egg size and shape, including lineages within Palaeoptera, Polyneoptera, and Hemiptera. Most lineages have an scaling exponent less than 1, supporting the prediction that larger eggs will tend to be proportionally longer than smaller eggs[18,59,60].

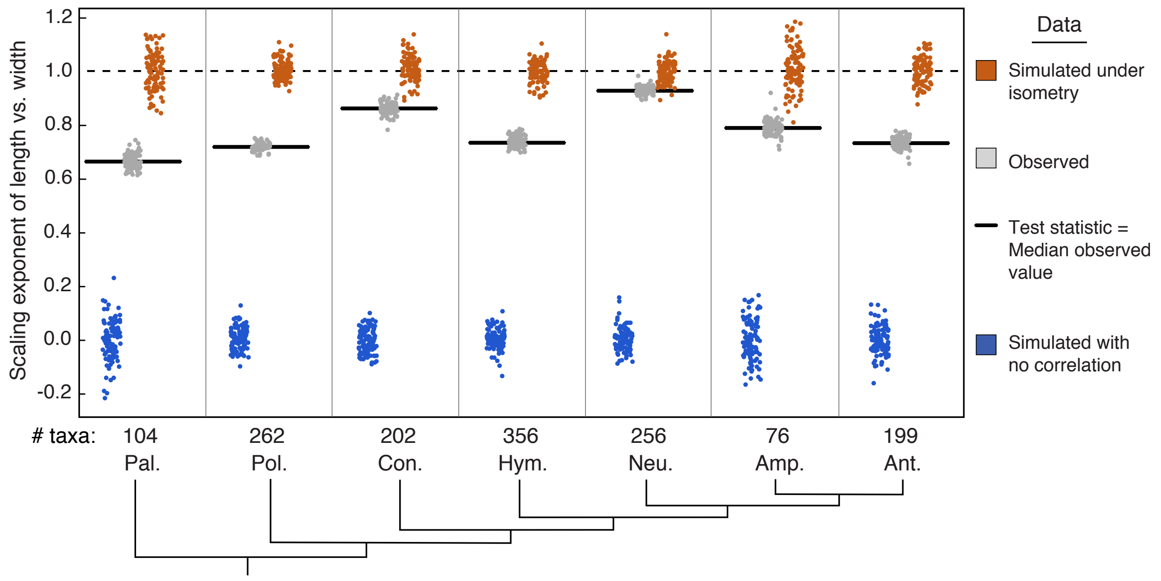

Figure S11: **Testing alternate hypotheses of egg size and aspect ratio evolution using a parametric bootstrap.** The distribution of the scaling exponent of length vs. width calculated from data simulated under alternate hypotheses compared to the observed distribution of scaling exponents (gray, test statistic = median value, black bar) for seven insect lineages. Alternate hypotheses include that egg shape and size are unrelated (slope = 0, blue), and that egg shape and size have an isometric relationship (slope = 1, orange).

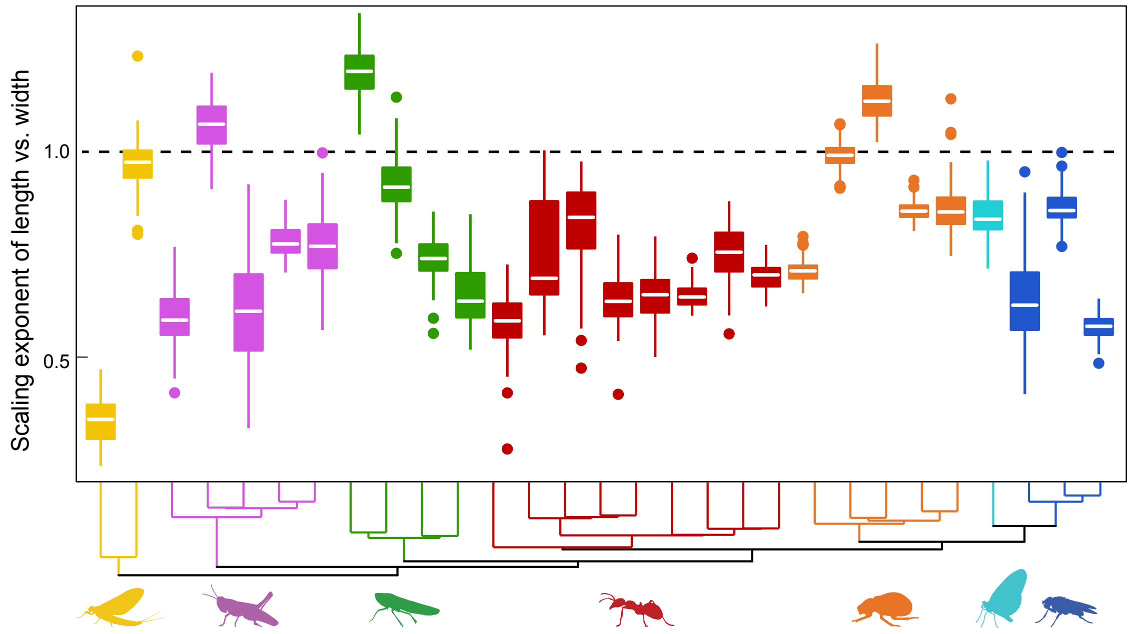

Figure S12: **Evolution of the relationship between egg size and aspect ratio across insect lineages.** The distribution of scaling exponents of length vs. width across all monophyletic clades in our dataset with more than 20 and fewer than 50 tips. The dashed black line represents a hypothetical 1:1 relationship (isometry). Colors correspond to the clades shown in Fig. S1.

#### 5.3 Accounting for body size in egg shape and size allometries

Given that hypotheses about the relationship between egg shape and size invoke egg scaling constraints within the insect body[18,59–61], we tested the effect of accounting for body size on our results. We matched the previously published[62] median body length (see Section 6.3 for details) to the average egg length and width for 417 insect families and 9 insect orders. We controlled for adult body size in the egg allometry comparison by calculating the phylogenetic residuals[63] of log-egg length and log-egg width against the log-adult body length.

Consistent with analyses that did not account for body size, in Neuropteroidea, egg length scales near-isometrically with width when accounting for body size, while in other insect clades larger eggs for a given body size are proportionally longer (Fig. S13 and Table S8). In the groups Palaeoptera and Condylognatha, the relationship between egg width and length is not significant. However these clades have the lowest sample size at the family-level, therefore our ability to detect relationships is weakest (Fig. S13B).

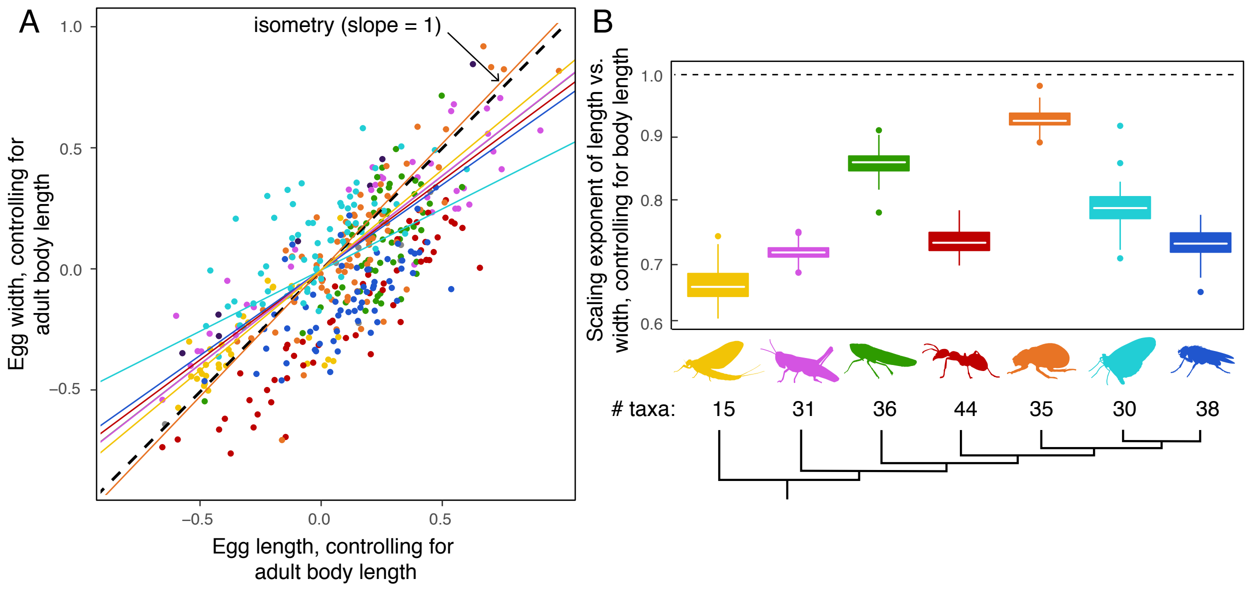

Figure S13: **Allometry of egg size and aspect ratio, controlling for adult body size. A**, PGLS regression of egg width (mm, log tranformed) and length (mm, log transformed), comparing the phylogenetic residuals of both traits against adult body length (mm, log transformed). The colored lines are the phylogenetic regression for each clade on the summary tree, colors correspond to the clades shown S1. Colored points are family or order-level averages. **B**, The distributions of scaling exponents of length vs. width, controlling for adult body length, for seven monophyletic insect clades. In both panels the dashed black line represents a hypothetical 1:1 relationship (isometry).

#### 5.4 Testing additional shape allometries

In addition to comparing the relationship between egg aspect ratio and egg size, we also tested the relationship between aspect ratio and two other features of egg shape: asymmetry and angle of curvature. We compared each shape parameter (square root transformed) to log of egg length, controlling for egg width using phylogenetic residuals. This allows us to ask whether eggs which are longer given their width (higher aspect ratio) are also more asymmetrical or more curved.

| | p-value | slope | intercept | sample size |
|---|---|---|---|---|
| Hexapoda | 0 - <0.005 | 0.67 - 0.75 | 0 | 236 |
| Hymenoptera | 0 - <0.005 | 0.69 - 0.90 | 0 | 44 |
| Condylognatha | 0 - 0.05 | 0.39 - 0.88 | 0 | 36 |
| Antliophora | 0 - <0.005 | 0.57 - 0.70 | 0 | 38 |
| Neuropteroidea | 0 - <0.005 | 0.91 - 1.07 | 0 | 35 |
| Amphiesmenoptera | 0 - <0.005 | 0.43 - 0.56 | 0 | 30 |
| Polyneoptera | 0 - <0.005 | 0.67 - 0.82 | 0 | 31 |
| Palaeoptera | 0.02 - 0.75 | 0.05 - 0.55 | 0 | 15 |

Table S8: **Results of PGLS of egg length and with, controlling for body size.**

Our results show that eggs with a higher aspect ratio (proportionally longer for their width) are not more asymmetrical than low aspect ratio counterparts (Fig. S14 and Table S9). Across insects, eggs with a higher aspect ratio tend to be more curved, though this relationship is likely driven by the lineages with very curved eggs (Hymenoptera, Condylognatha, and Antliophora; Fig. S14 and Table S10).

| | p-value | slope | intercept | sample size |
|---|---|---|---|---|
| Hexapoda | 0 - 0.17 | 0.05 - 0.18 | 0 | 800 |
| Hymenoptera | 0.05 - 1.00 | -0.15 - 0.18 | 0 | 177 |
| Condylognatha | 0 - 0.95 | -0.01 - 0.45 | 0 | 150 |
| Antliophora | 0 - 0.18 | 0.16 - 0.46 | 0 | 81 |
| Neuropteroidea | 0.08 - 0.84 | 0.02 - 0.15 | 0 | 141 |
| Amphiesmenoptera | 0.11 - 1.00 | -0.38 - 0.05 | 0 | 25 |
| Polyneoptera | 0.13 - 0.98 | -0.13 - 0.13 | 0 | 140 |
| Palaeoptera | 0.02 - 0.99 | -0.03 - 0.27 | 0 | 71 |

Table S9: **Results of PGLS of egg length and asymmetry, controlling for egg width.**

| | p-value | slope | intercept | sample size |
|---|---|---|---|---|
| Hexapoda | 0 - <0.005 | 0.41 - 0.58 | 0 | 785 |
| Hymenoptera | 0 - 0.01 | 0.40 - 0.77 | 0 | 177 |
| Condylognatha | 0 - 0.02 | 0.38 - 0.75 | 0 | 150 |
| Antliophora | 0.01 - 0.82 | 0.05 - 0.55 | 0 | 80 |
| Neuropteroidea | 0 - 0.11 | 0.22 - 0.52 | 0 | 141 |
| Amphiesmenoptera | 0.21 - 0.99 | -0.32 - 0.16 | 0 | 23 |
| Polyneoptera | 0 - 0.04 | 0.33 - 0.71 | 0 | 131 |
| Palaeoptera | 0.03 - 0.68 | 0.07 - 0.37 | 0 | 70 |

Table S10: **Results of PGLS of egg length and curvature, controlling for egg width**

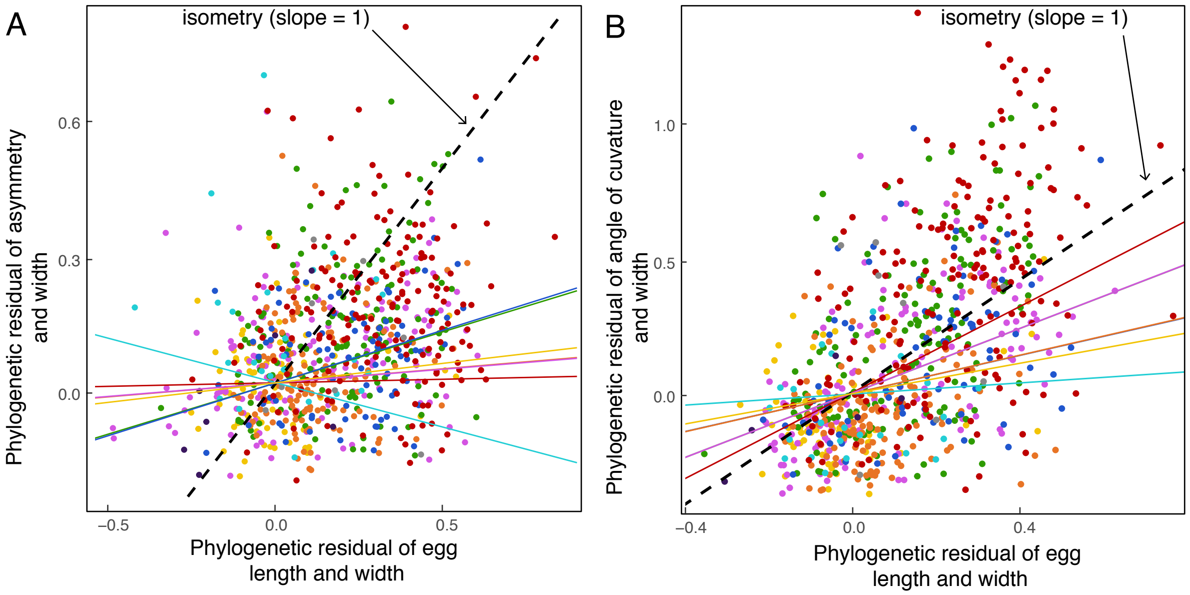

Figure S14: **Allometry of egg asymmetry, angle of curvature, and aspect ratio.** **A**, PGLS regression of egg asymmetry (unitless; square root transformed) and length (mm; log transformed), comparing the phylogenetic residuals of both traits against egg width (mm; log transformed). **B**, PGLS regression of egg curvature (degrees; square root transformed) and length (mm; log transformed), comparing the phylogenetic residuals of both traits against egg width (mm; log transformed). In both panels the dashed black line represents a hypothetical 1:1 relationship (isometry), colored lines are the phylogenetic regression for each clade, and colored points are representative eggs from each genus in the phylogeny. Colors correspond to the clades shown in Fig. S1.

### 6 Egg size and development

We tested the relationship between the evolution of egg morphology, embryonic development, and adult size. To compare traits across species, we collected descriptions of developmental times and adult size from the insect literature.

#### 6.1 Collecting developmental time data

We collected literature sources that described the development of insects and used them to assemble a dataset of three developmental traits. The developmental time data and corresponding original sources are available at `https://github.com/shchurch/Insect_Egg_Evolution`, file 'development.csv'. The developmental traits considered were as follows:

*Interval between syncytial mitoses:* Insects in most lineages that have been studied begin embryogenesis with a series of syncytial nucleus divisions (mitotic divisions with absent or incomplete cytokinesis)[64,65]. For sources that reported a single estimate of the time interval between mitotic divisions, we used that value (converted to hours). When a source reported multiple intervals, we used the mean duration of the reported mitotic intervals that occur before nuclei initially reach the periphery of the egg. We did not collect mitotic interval data from the species of polyembryonic insects that develop holoblastically.

*Time to cellularization:* This trait was included only for species with syncytial development. When sources reported a single time point, we used it (converted to hours). If a range was reported, we used the midpoint of that range.

*Duration of embryogenesis:* We define embryogenesis as the development that takes place prior to *hatching*, which is the point at which a mobile first instar insect (larva or nymph) exits the egg.

We only included data from sources that reported the temperature at which the embryo developed, as developmental rate varies with incubation temperature[66,67]. Moreover, data from many animals, including insects, are consistent with the hypothesis that the temperature-dependence of developmental rate is due the general temperature-dependence of reaction kinetics[68]. Thus, we followed the method of recent work on insect developmental rate[69] to re-scale all developmental times to a standardized temperature of 20 °C using the Boltzmann-Arrhenius equation with an Arrhenius temperature parameter set to 8000K. All temperature-adjusted developmental times were $\log_{10}$ transformed.

#### 6.2 Comparing egg size and developmental time

The three measures of developmental time described above were compared to egg volume using a PGLS regression over 100 trees randomly drawn from the posterior distribution. Only species present in both the development and the egg dataset were compared. For species with developmental records that had more than one egg description in the database, a random matching egg entry was chosen for each iteration over the posterior distribution of trees.

None of the three developmental parameters had a significant relationship to egg volume across the insect phylogeny (Fig. S15, Table S11). Furthermore, we observed that if phylogeny was not taken into account, we could recover a spurious relationship between egg volume and duration of embryogenesis (p-value = 0.002, adjusted $R^2$ = 0.122, Fig. S16; using a linear model on the same data included in the phylogenetic regression). Given that a previous study had reported a significant relationship between developmental time and egg size[69], we suggest that the results of that study were likely due to the artifact caused by failing to account for the phylogenetic non-independence of phenotypes.

| | p-value | slope | intercept | sample size |
|---|---|---|---|---|
| egg volume vs duration of embryogenesis | 0.07 - 0.28 | 0.07 - 0.11 | 2.29 - 2.37 | 46 |
| egg volume vs interval between pre-blastoderm mitoses | 0.30 - 0.83 | 0.02 - 0.12 | -0.14 - 0 | 16 |
| egg volume vs time to cellularization | 0.08 - 0.51 | 0.12 - 0.33 | 1.20 - 1.43 | 18 |

Table S11: **Results of PGLS of developmental time and egg size**

#### 6.3 Egg size and body size

We compared the predicted evolutionary relationship of egg size and body size[70,71] by matching the egg database to published records of insect body length. Rainford et al. (2016)[62] described the maximum and minimum adult body length for 764 insect families and 10 insect orders, of which 426 are represented in the insect egg database. From these we calculated the median body length for each family, and matched this to the average egg volume from the

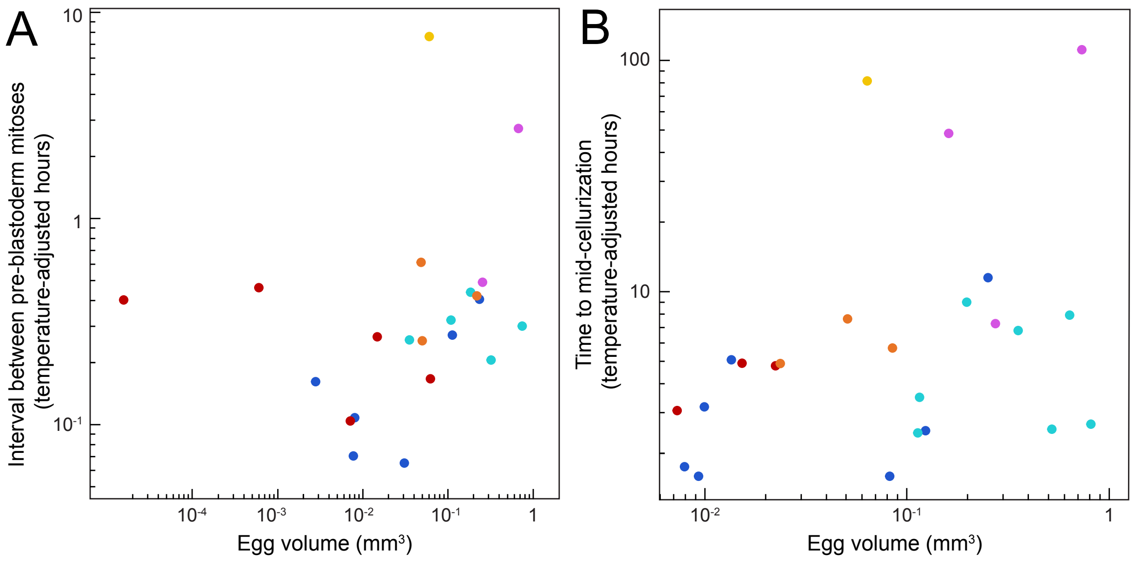

Figure S15: **Comparisons of egg size with additional measures of embryonic development time.** **A**, Embryonic time measured as the mean reported hours between mitoses in the pre-blastoderm stage (temperature adjusted[68]; log scale), compared to egg volume ($mm^3$; log scale). **B**, Embryonic time measured as the reported hours to the midpoint of cellularization (temperature adjusted[68]; log scale), compared to egg volume ($mm^3$; log scale). In both panels, each point represents an insect species for which both developmental and egg morphological data were available. Colors correspond to the clades shown in Fig. S1.

insect egg database for the same family. Given that the median body lengths reported in Rainford et al. (2016)[62] may have been drawn from a different subset of species per family than the average egg size from our egg database, we tested the impact of sampling by randomly reducing the number of entries in each family in the egg database by 50% and reanalyzing the data 100 times.

We compared the allometric relationship between egg size and body size with PGLS regression across the family-level phylogeny published by Rainford et al. (2014)[3]. Our results showed that across the insect phylogeny, smaller insects lay proportionally larger eggs (that is, there is a significant allometric relationship with a slope less than one; main text Fig. 4 and Table S12). Within the insect lineages Palaeoptera and Antliophora, however, body size does not predict egg size (that is, there is no statistically significant relationship between egg size and body size). These results are robust to downsampling the egg database for each family/order, indicating that they are not due to an artifact of sampling differences between the egg size and body size datasets.

To test our results against alternative hypotheses we simulated egg size and body size datasets under known evolutionary models and analyzed them using the same methods. We followed the same parametric bootstrap method as described in 5.2, using here the best fitting models (Table S5) for body size and egg size to simulate family-level egg volume data. Our results show that the first hypothesis, an isometric relationship between egg size and body size, cannot be rejected for the lineages Condylognatha and Hymenoptera (p-value, isometry = 0.02 for both lineages). The second hypothesis, that egg size and body size evolve independently, cannot be rejected for the lineages Palaeoptera and Antliophora (p-value, no relationship = 0.04 and 0.03 respectively).

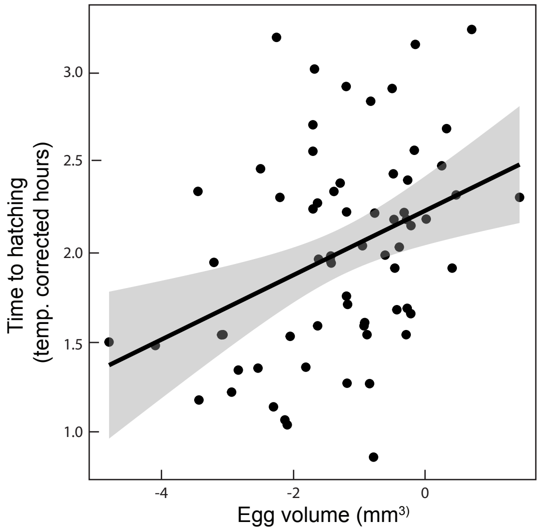

Figure S16: **Non-phylogenetic regression of developmental time and egg size.** For illustration, we show that a regression that failed to account for phylogeny would find a spurious significant relationship (p-value 0.002, adjusted $R^2$ = 0.122) between duration-of-embryogenesis (temperature adjusted[68]; log scale) and egg volume ($mm^3$; log scale). The black line is the fitted regression, with 95% confidence intervals shown in gray. Each point represents an insect species for which developmental and egg morphological data were available for a member of a genus that could be included in the insect phylogeny described in Section 2.

| | p-value | slope | intercept | sample size |
|---|---|---|---|---|
| Hexapoda | 0 - <0.005 | 0.43 - 0.48 | -3.14 - -2.82 | 238 |
| Hymenoptera | 0 - <0.005 | 0.63 - 0.81 | -4.17 - -3.56 | 44 |
| Condylognatha | 0 - <0.005 | 0.57 - 0.77 | -3.70 - -3.02 | 36 |
| Antliophora | 0.02 - 0.19 | 0.17 - 0.31 | -2.60 - -2.12 | 39 |
| Neuropteroidea | 0 - <0.005 | 0.36 - 0.45 | -2.60 - -2.26 | 36 |
| Amphiesmenoptera | 0 - <0.005 | 0.34 - 0.46 | -3.34 - -2.78 | 31 |
| Polyneoptera | 0 - <0.005 | 0.51 - 0.64 | -3.00 - -2.46 | 31 |
| Palaeoptera | 0 - 0.02 | 0.28 - 0.42 | -3.95 - -3.39 | 15 |

Table S12: **Results of PGLS of egg volume and adult body length cubed**

| | test statistic | p-value, isometry | p-value, no relationship |
|---|---|---|---|
| Palaeoptera | 0.35 | <0.01 | 0.04 |
| Polyneoptera | 0.57 | <0.01 | <0.01 |
| Condylognatha | 0.66 | 0.02 | <0.01 |
| Hymenoptera | 0.72 | 0.02 | <0.01 |
| Neuropteroidea | 0.41 | <0.01 | <0.01 |
| Amphiesmenoptera | 0.39 | <0.01 | <0.01 |
| Antliophora | 0.23 | <0.01 | 0.03 |

Table S13: **Results of a parametric bootstrap of egg size and body size.** For parametric bootstraps, a p-value of <0.01 indicates that out of 100 bootstraps, no values values were greater / less than the test statistic.

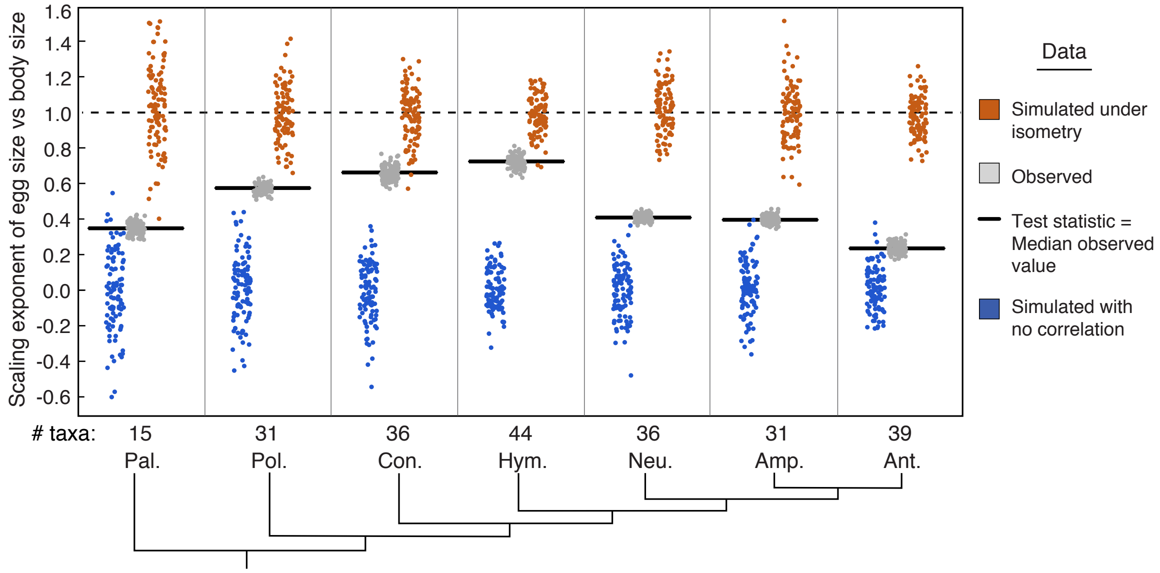

Figure S17: **Testing alternative hypotheses of egg size and adult body size evolution using a parametric bootstrap.** The distribution of scaling exponents of egg size vs. body size calculated from data simulated under alternate hypotheses compared to the observed distribution of the scaling exponent (gray, test statistic = median value, black bar), for seven insect lineages. Alternate hypotheses include that egg shape and adult body size are unrelated (slope = 0, blue), and that egg shape and adult body size have an isometric relationship (slope = 1, orange).

### 7 Evolutionary history of ecological traits

#### 7.1 Parasitoid and internal oviposition

We compiled a list of parasitoid insects from multiple published reviews[72–77]. The table of parasitoid insect taxa and the code used to perform ecological analyses is available at `https://github.com/shchurch/Insect_Egg_Evolution`, file 'ecology_table_parasitoid.csv'. We used this list to classify taxa in the insect egg database as non-parasitoid or parasitoid, including ecto- and endoparasitoids, and excluding kleptoparasitic and gall-forming insects. We further classified parasitoid taxa as laying eggs externally or internally to their hosts. Insects that were listed as strictly endoparasitic with no reference to eggs laid externally, and for which no additional information was available, were considered to lay eggs internally. Reviews of parasitism across insects differed in the taxonomic level described. For each source, we used the lowest recorded taxonomic level to annotate taxa in the egg database. For some clades it was not possible to classify all members unambiguously (e.g., the lowest description described the group has having "some parasitoids"). In order to test the impact of this uncertainty on our analyses we implemented both a "relaxed" classification system, in which taxa with ambiguous records were also coded as parasitoid / internal, and a "strict" classification system, in which only taxa that could be unambiguously defined as parasitoid / internal were coded as such.

We reconstructed the evolutionary history of both internal parasitic oviposition and ecto- or endoparasitoid habit (Fig. S18) on the Misof backbone MCC phylogeny[2] using an equal-rates model (R package corHMM[78], version 1.22, function rayDISC, `node.states=marginal`). Using the relaxed classification method, we recovered 22 evolutionary shifts to ecto- or endoparasitoid habit across Hymenoptera, Antliophora, and Neuropteroidea, with 14 shifts to internal oviposition. We also found evidence for 8 reversals from parasitoid habit and 7 reversals from internal oviposition. These numbers likely reflect a minimum bound as more changes may have occurred in groups that are not represented in the insect egg database and phylogeny.

#### 7.2 Aquatic insects and oviposition

We compiled a list of aquatic taxa in our database from multiple published reviews[77,79–97]. The table of aquatic insect taxa and the code used to perform ecological analyses is available at `https://github.com/shchurch/Insect_Egg_Evolution`, file 'ecology_table_aquatic.csv'. Taxa were first classified as broadly aquatic, including semi-aquatic or riparian, and excluding insects that lay eggs within aquatic plants (phytophilous) or overhanging water. We further classified aquatic insects as laying eggs in water or out of water. We used the same relaxed and strict classification methods as described for parasitoid insects.

Using the same methods described above, we reconstructed the evolutionary history of aquatic and semiaquatic insects and oviposition in water (Fig. S19). Using the relaxed classification method, we recovered 32 separate transitions to aquatic or semiaquatic larval habit, and 16 transitions to aquatic oviposition. We also recovered 5 reversals to non-aquatic, semiaquatic, or riparian habit and 6 reversals to non-aquatic oviposition. As described above, these numbers are likely a minimum of the number of possible transitions.

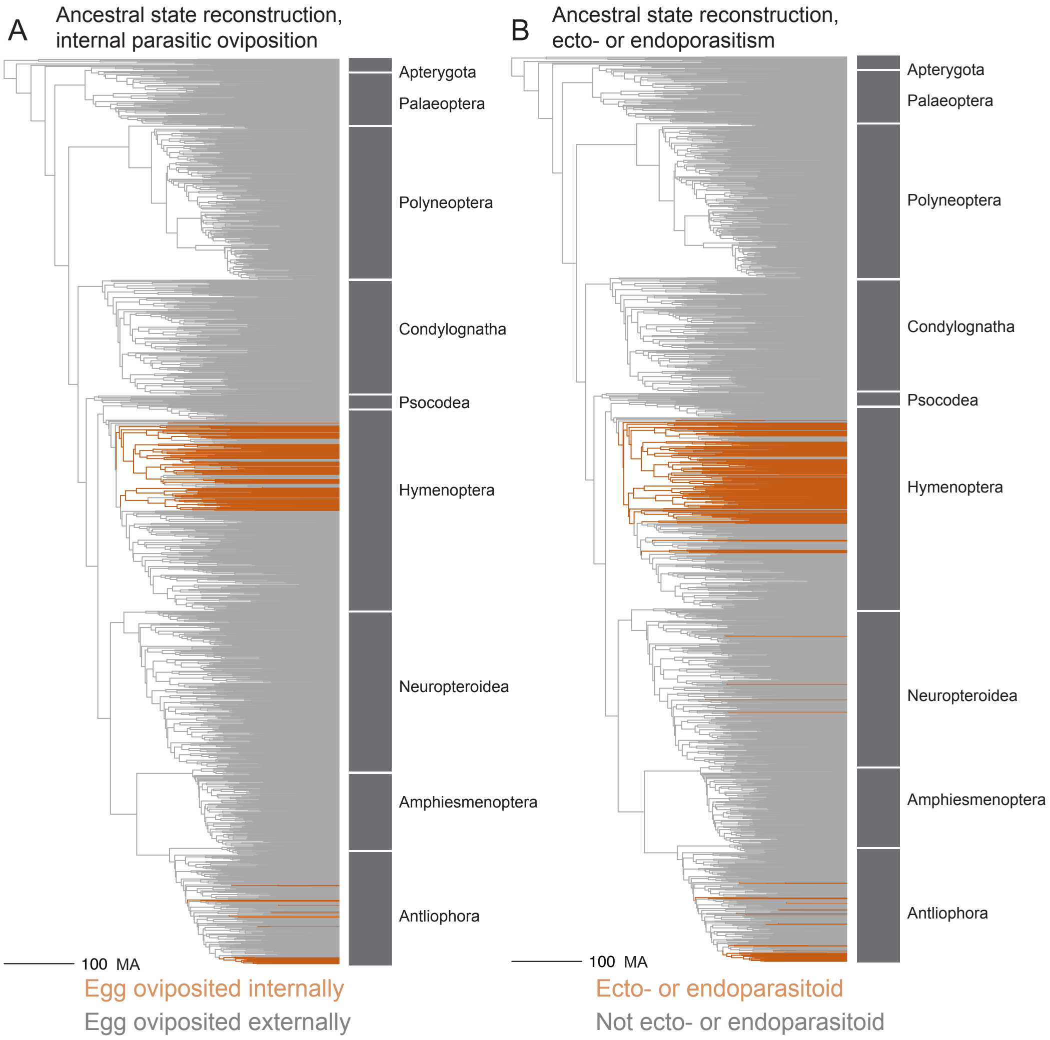

Figure S18: **Ancestral state reconstruction of parasitoid oviposition ecology and life-history.** Ancestral state reconstructions using the "relaxed" classifcation method of **A** oviposition within an animal host, and **B** endoparasitoid and ectoparasitoid life history. Lineages that descend from a node reconstructed with a more than 50% likelihood of the derived state (internal or parasitoid, respectively) are shown in orange. Scale bar represents 100 million years.

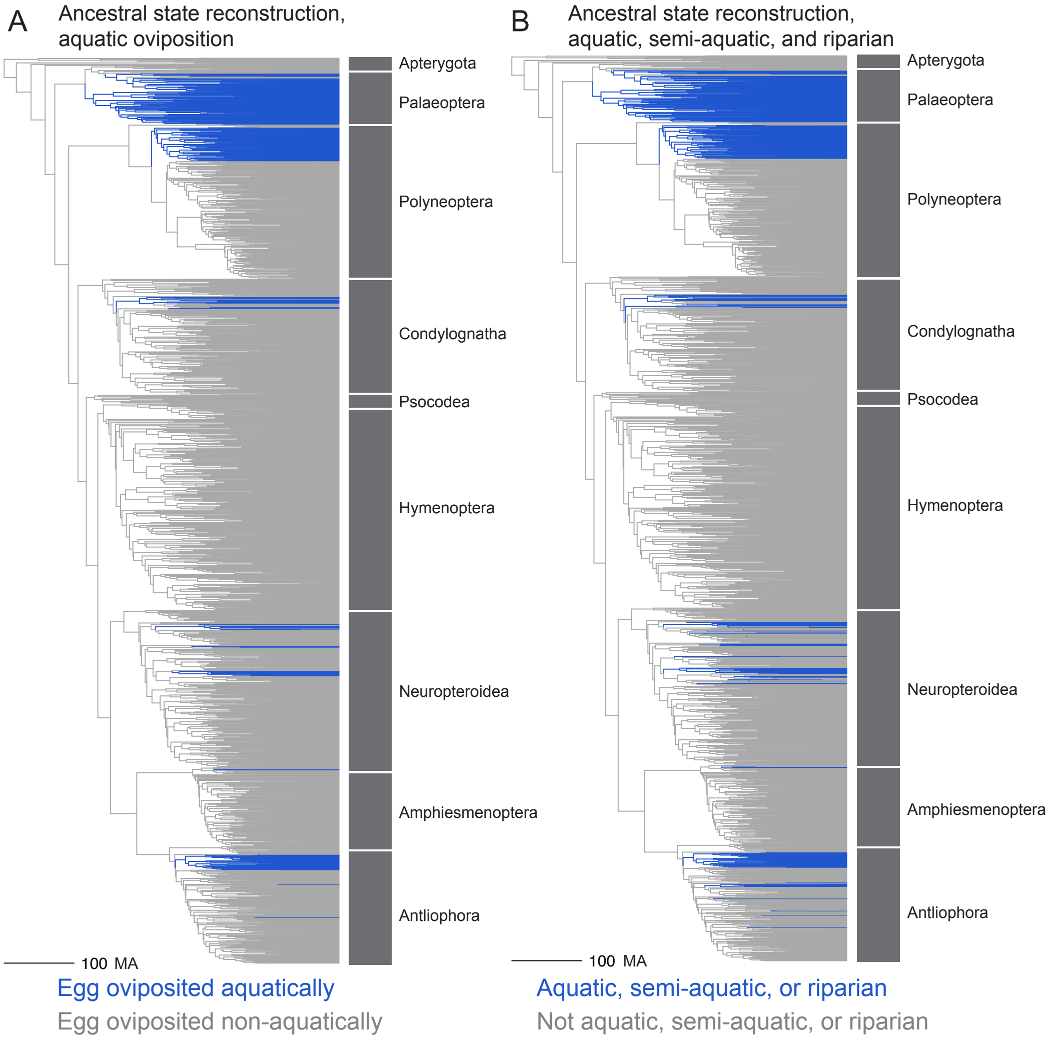

Figure S19: **Ancestral state reconstruction of aquatic oviposition ecology and life-history.** Ancestral state reconstructions using the "relaxed" classification method of **A** oviposition in water, and **B** aquatic, semi-aquatic, or riparian life history (excluding ovipositing in aquatic plants or overhanging water). Lineages which descend from a node reconstructed with a more than 50% likelihood of the derived state (aquatic) are shown in blue. Scale bar represents 100 million years.

##### 7.3 Migration, flight, and wingless insects

There have likely been thousands of evolutionary shifts to flightless and wingless forms in insects[98]. We analyzed flight ability in Phasmatodea and Lepidoptera, using different metrics for flight ability in each. For Phasmatodea, we used published reviews to classify stick insects as either capable of flying or flightless (the latter category including both wingless and partially winged species that are not capable of flying)[99,100]. Phasmatodea taxa that could not be reliably classified in our dataset were excluded from subsequent analyses. For Lepidoptera, we analyzed migratory behavior as a proxy for the capability of flying longer distances than non-migratory Lepidoptera. We used published reviews of migratory insects to identify taxa in our database known to exhibit long-distance migration[101–106]. The ancestral state reconstructions of these traits are shown in Fig. S20.

##### 7.4 Testing eco-evolutionary models of egg evolution

We used the ancestral state reconstructions of parasitoid (Section 7.1) and aquatic (Section 7.2) ecological states in a comparison of evolutionary models. For each egg morphological trait (volume, aspect ratio, asymmetry, and angle of curvature) we compared the fit of evolutionary models that account for ecological history (OU model with different optima for each ecological regime) to models which do not take ecology into account (BM, OU with a single optimum) using the R package OUwie[107] (version 1.50). We considered a significantly better fit at $\Delta$AICc > 2.

Our results show that models that account for internal parasitic oviposition fit the data best for both egg volume and asymmetry, but not for aspect ratio or curvature (Table S14). Insect lineages that oviposit in animal hosts typically have smaller eggs (OUM $\theta$ values for egg volume, log transformed, non-internal: -1.82, internal: -3.96) and more asymmetrical eggs (OUM $\theta$ values for egg asymmetry, sq. root transformed, non-internal: 0.24, internal: 0.40), than lineages that do not oviposit in animal hosts. These results are consistent when considering all endo- and ectoparasitoids, rather than only those that oviposit internally to their animal hosts (Table S15). These results are also robust to uncertainty in the classification system (Table S16).

| | Brownian Motion | OU - 1 Optimum | OU - Multiple Optima |
|---|---|---|---|
| volume | 29.14 | 31.15 | 0.00 |
| aspect ratio | 0.00 | 2.01 | 3.42 |
| curvature | 0.00 | 2.02 | 3.05 |
| asymmetry | 49.70 | 12.03 | 0.00 |

Table S14: **Comparing evolutionary models ($\Delta$AICc values) of egg morphology and internal parasitic oviposition using the "relaxed" classification method.**

With respect to aquatic oviposition, models accounting for this characteristic fit the data best for both egg volume and aspect ratio, but not asymmetry or curvature (Table S17). Insect lineages that oviposit in water typically have smaller eggs (OUM $\theta$ values for egg volume, log transformed, non-aquatic: -1.66, aquatic: -3.19) and more spherical eggs (OUM $\theta$ values for egg aspect ratio, log transformed, non-aquatic: 0.22, aquatic: -0.06), than lineages that do not oviposit in water. These results are consistent when considering aquatic and riparian insects (Table S18); they are also robust to uncertainty in the classification system (Table S19).

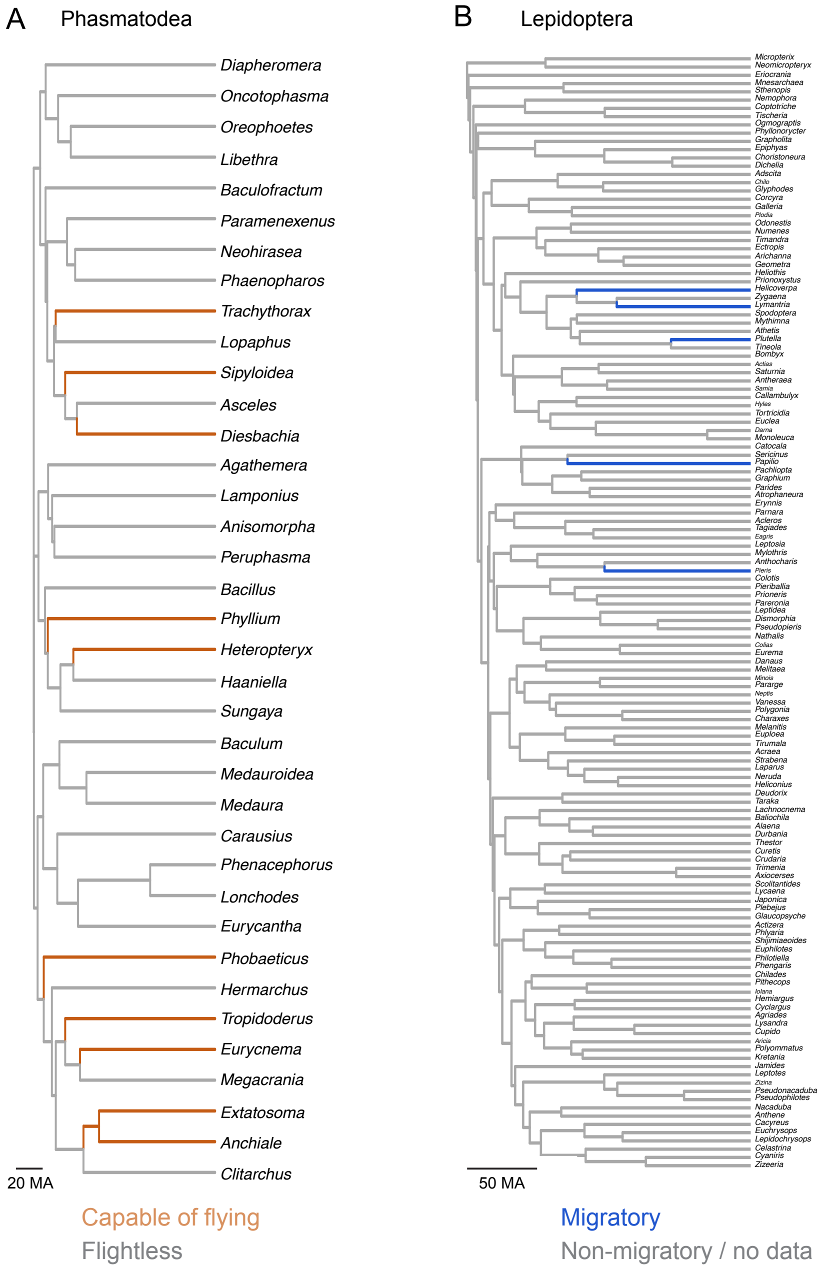

Figure S20: **Ancestral state reconstructions of flight capability in selected clades.** **A**, Ancestral state reconstruction of flightless (gray) vs capable of flying (orange) Phasmatodea. Scale bar represents 20 million years. **B**, Ancestral state reconstruction of migratory behavior (blue) in Lepidoptera. Lineages that descend from a node reconstructed with more than 50% likelihood of the derived state (capable of flying or migratory) are shown in color. Scale bar in million years.

| | Brownian Motion | OU - 1 Optimum | OU - Multiple Optima |
|---|---|---|---|
| volume | 14.56 | 16.57 | 0.00 |
| aspect ratio | 0.00 | 2.01 | 0.49 |
| curvature | 0.00 | 2.02 | 3.31 |
| asymmetry | 42.94 | 11.93 | 0.00 |

Table S15: **Comparing evolutionary models (ΔAICc values) of egg morphology and ecto- or endoparasitoid habit using the "relaxed" classification method.**

| | Brownian Motion | OU - 1 Optimum | OU - Multiple Optima |
|---|---|---|---|
| volume | 23.62 | 25.63 | 0.00 |
| aspect ratio | 0.33 | 2.34 | 0.00 |
| curvature | 0.00 | 2.02 | 3.36 |
| asymmetry | 42.92 | 3.12 | 0.00 |

Table S16: **Comparing evolutionary models (ΔAICc values) of egg morphology and internal parasitic oviposition using the "strict" classification method (ambiguous taxa coded as not exhibiting internal parasitic oviposition)**

| | Brownian Motion | OU - 1 Optimum | OU - Multiple Optima |
|---|---|---|---|
| volume | 21.42 | 23.43 | 0.00 |
| aspect ratio | 15.03 | 17.04 | 0.00 |
| curvature | 0.96 | 2.98 | 0.00 |
| asymmetry | 28.86 | 0.00 | 0.47 |

Table S17: **Comparing evolutionary models (ΔAICc values) of egg morphology and and aquatic oviposition using the "relaxed" classification method.**

| | Brownian Motion | OU - 1 Optimum | OU - Multiple Optima |
|---|---|---|---|
| volume | 23.26 | 25.26 | 0.00 |
| aspect ratio | 15.01 | 17.02 | 0.00 |
| curvature | 0.23 | 2.25 | 0.00 |
| asymmetry | 35.99 | 0.00 | 1.73 |

Table S18: **Comparing evolutionary models (ΔAICc values) of egg morphology and aquatic, semi-aquatic, and riparian habit using the "relaxed" classification method.**

| | Brownian Motion | OU - 1 Optimum | OU - Multiple Optima |
|---|---|---|---|
| volume | 26.94 | 28.95 | 0.00 |
| aspect ratio | 12.84 | 14.84 | 0.00 |
| curvature | 1.73 | 3.54 | 0.00 |
| asymmetry | 37.58 | 0.00 | 1.79 |

Table S19: **Comparing evolutionary models (ΔAICc values) of egg morphology and aquatic oviposition using the "strict" classification method (ambiguous taxa coded as not exhibiting aquatic oviposition).**

Within both Phasmatodea (Table S20) and Lepidoptera (Table S21), models that account for the evolutionary history of flight ability do not fit any egg morphological data significantly better than those that do not.

| | Brownian Motion | OU - 1 Optimum | OU - Multiple Optima |
|---|---|---|---|
| volume | 0.00 | 2.37 | 4.30 |
| aspect ratio | 0.00 | 2.04 | 4.23 |
| curvature | 0.00 | 2.49 | 5.55 |
| asymmetry | 0.00 | 0.11 | 3.26 |

Table S20: **Comparing evolutionary models ($\Delta$AICc values) of egg morphology and flightlessness in Phasmatodea.**

| | Brownian Motion | OU - 1 Optimum | OU - Multiple Optima |
|---|---|---|---|
| volume | 0.00 | 2.37 | 4.30 |
| aspect ratio | 0.00 | 2.04 | 4.23 |
| curvature | 0.00 | 2.49 | 5.55 |
| asymmetry | 0.00 | 0.11 | 3.26 |

Table S21: **Comparing evolutionary models ($\Delta$AICc values) of egg morphology and migratory behavior in Lepidoptera.**

#### 7.5 Allometry and ecology

To test for a possible interaction between ecology and the evolution of the allometric relationship between size and shape, we compared the scaling exponent of length vs. width (slope in log-log space) between groups that have converged upon the same ecological state. First, we identified the nodes where an ecological shift was most likely to have occured (the probability of an ecological state being different than the parent node was above 50%), and then further identified those nodes that had a sufficient number (threshold > 20) of descendant tips with egg morphological data to robustly calculate the scaling exponent. For internal oviposition there were two clades (both in Hymenoptera) that met these requirements, and for aquatic oviposition there were three (Ephemeroptera, Plecoptera, and a subset of Odonata).

We calculated the allometric exponent as the slope of the phylogenetic regression between log-egg length and log-egg width, over 100 trees randomly drawn from the posterior distribution. We compared the slopes to the slope of the paraphyletic group of insects with the ancestral state (non-internal or non-aquatic oviposition). If transitions to new oviposition ecologies were predictive of a change in the allometric relationship, clades with the derived ecology would show consistent shifts up or down relative the ancestral ecological state.

Our results show dynamic evolution of the allometric exponent, but no consistent directional shift across ecologically convergent clades (Fig. S21). Because the number of shifts with sufficient sample size is low, further exploration by expanding the number of described egg morphologies in other internal and aquatic lineages would strengthen the power of this comparison.

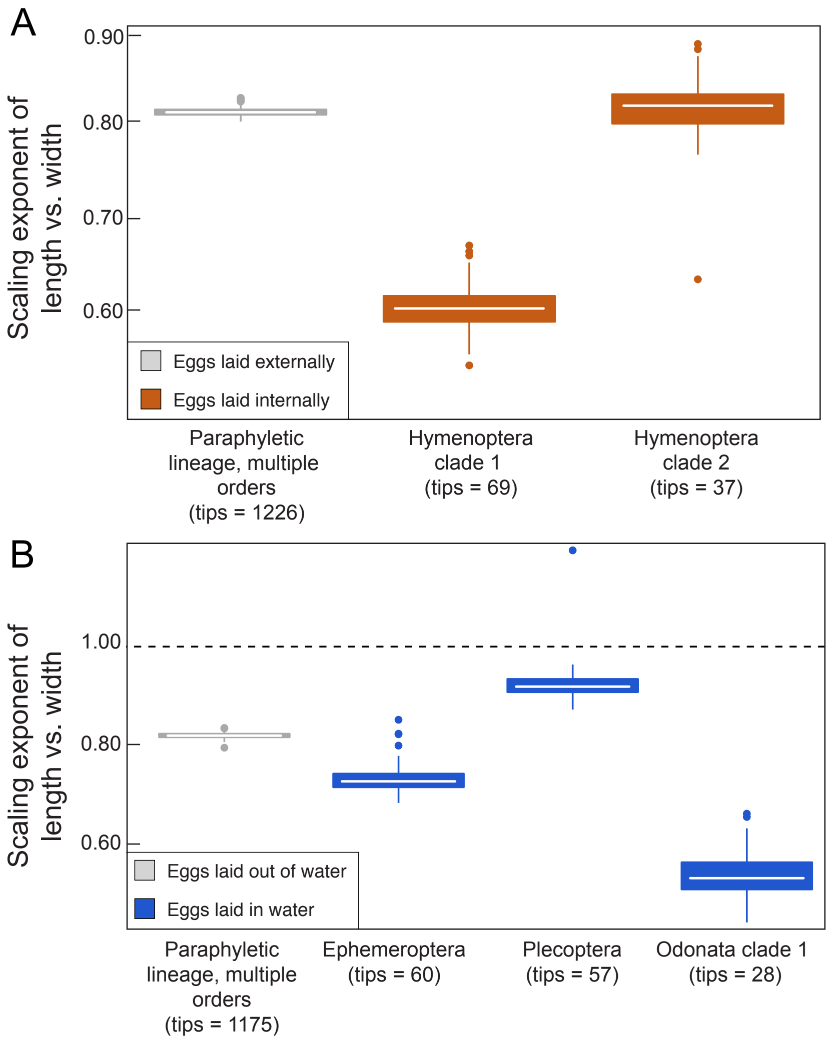

Figure S21: **Comparing allometric slope by ecological regimes.** The scaling exponent of egg length vs. width, comparing clades with convergent ecological regimes to paraphyletic lineages with the ancestral state. **A**, Comparing the allometric exponent between two independent lineages of internal oviposition, and the paraphyletic non-internal lineage (gray). **B**, Comparing the allometric exponent between three independent lineages of aquatic oviposition (blue), and the paraphyletic non-aquatic lineage (gray). The scaling exponents are calculated over a posterior distribution of trees and accounting for intrageneric variation. Only lineages with more than 20 genera with egg descriptions are included. See Fig. S18 and S19 for details on lineage composition. The dashed black line represents a hypothetical 1:1 relationship.

### 8 Summary of Phylogenetic Generalized Least Squares (PGLS) results

To test the robustness of our results to the phylogenetic backbone, we repeated each PGLS comparison using the Rainford et al. (2018)[3] backbone phylogeny (see Section 5.1 for more details). We found that no results were significantly different than those calculated using the Misof et al. (2018)[2] backbone phylogeny (Table S22).

We also repeated each PGLS comparison using a covariance matrix based on a decelerating rate of evolution, using the corBlomberg function in the R package nlme[57] (see Section 5.1 for more details). We found that no results were significantly different than those calculated using a Brownian-Motion based covariance matrix (Table S23).

| analysis | clade | Rain. p-value | Rain. slope | Misof p-value | Misof slope |
|---|---|---|---|---|---|
| egg volume vs duration of embryogenesis | Hexapoda | 0.02 - 0.10 | 0.10 - 0.14 | 0.07 - 0.28 | 0.07 - 0.11 |
| egg volume vs interval between pre-blastoderm mitoses | Hexapoda | 0.27 - 0.75 | 0.03 - 0.13 | 0.30 - 0.83 | 0.02 - 0.12 |
| egg volume vs time to cellularization | Hexapoda | 0.08 - 0.39 | 0.15 - 0.30 | 0.08 - 0.51 | 0.12 - 0.33 |
| egg length vs width | Hexapoda | 0 - <0.005 | 0.78 - 0.81 | 0 - <0.005 | 0.77 - 0.80 |
| egg length vs width | Hymenoptera | 0 - <0.005 | 0.71 - 0.80 | 0 - <0.005 | 0.70 - 0.78 |
| egg length vs width | Condylognatha | 0 - <0.005 | 0.79 - 0.91 | 0 - <0.005 | 0.78 - 0.91 |
| egg length vs width | Antliophora | 0 - <0.005 | 0.69 - 0.79 | 0 - <0.005 | 0.65 - 0.78 |
| egg length vs width | Neuropteroidea | 0 - <0.005 | 0.89 - 0.98 | 0 - <0.005 | 0.89 - 0.98 |
| egg length vs width | Amphiesmenoptera | 0 - <0.005 | 0.71 - 0.89 | 0 - <0.005 | 0.71 - 0.92 |
| egg length vs width | Polyneoptera | 0 - <0.005 | 0.71 - 0.79 | 0 - <0.005 | 0.69 - 0.75 |
| egg length vs width | Palaeoptera | 0 - <0.005 | 0.59 - 0.73 | 0 - <0.005 | 0.61 - 0.74 |
| egg length vs asymmetry, residuals to egg width | Hexapoda | 0 - 0.08 | 0.07 - 0.16 | 0 - 0.17 | 0.05 - 0.18 |
| egg length vs asymmetry, residuals to egg width | Hymenoptera | 0.06 - 0.98 | -0.09 - 0.18 | 0.05 - 1.00 | -0.15 - 0.18 |
| egg length vs asymmetry, residuals to egg width | Condylognatha | 0 - 0.79 | 0.03 - 0.41 | 0 - 0.95 | -0.01 - 0.45 |
| egg length vs asymmetry, residuals to egg width | Antliophora | 0 - 0.10 | 0.17 - 0.46 | 0 - 0.18 | 0.16 - 0.46 |
| egg length vs asymmetry, residuals to egg width | Neuropteroidea | 0.06 - 0.98 | 0 - 0.16 | 0.08 - 0.84 | 0.02 - 0.15 |
| egg length vs asymmetry, residuals to egg width | Amphiesmenoptera | 0.06 - 0.93 | -0.41 - 0.03 | 0.11 - 1.00 | -0.38 - 0.05 |
| egg length vs asymmetry, residuals to egg width | Polyneoptera | 0.10 - 0.99 | -0.08 - 0.16 | 0.13 - 0.98 | -0.13 - 0.13 |
| egg length vs asymmetry, residuals to egg width | Palaeoptera | 0.19 - 0.94 | -0.02 - 0.15 | 0.02 - 0.99 | -0.03 - 0.27 |
| egg length vs angle of curvature, residuals to egg width | Hexapoda | 0 - <0.005 | 0.34 - 0.56 | 0 - <0.005 | 0.41 - 0.58 |
| egg length vs angle of curvature, residuals to egg width | Hymenoptera | 0 - <0.005 | 0.46 - 0.79 | 0 - 0.01 | 0.40 - 0.77 |
| egg length vs angle of curvature, residuals to egg width | Condylognatha | 0 - 0.04 | 0.34 - 0.76 | 0 - 0.02 | 0.38 - 0.75 |
| egg length vs angle of curvature, residuals to egg width | Antliophora | 0 - 0.94 | -0.07 - 0.59 | 0.01 - 0.82 | 0.05 - 0.55 |
| egg length vs angle of curvature, residuals to egg width | Neuropteroidea | 0 - 0.19 | 0.17 - 0.56 | 0 - 0.11 | 0.22 - 0.52 |
| egg length vs angle of curvature, residuals to egg width | Amphiesmenoptera | 0.33 - 0.98 | -0.26 - 0.16 | 0.21 - 0.99 | -0.32 - 0.16 |
| egg length vs angle of curvature, residuals to egg width | Polyneoptera | 0 - 0.24 | 0.22 - 0.71 | 0 - 0.04 | 0.33 - 0.71 |
| egg length vs angle of curvature, residuals to egg width | Palaeoptera | 0.04 - 0.66 | 0.07 - 0.32 | 0.03 - 0.68 | 0.07 - 0.37 |

Table S22: **Results of all PGLS analysis using the Rainford backbone phylogeny**[3]

| analysis | clade | Blom. p-value | Blom. slope | Brown, p-value | Brown. slope |
|---|---|---|---|---|---|
| egg volume vs duration of embryogenesis | Hexapoda | 0.08 - 0.32 | 0.06 - 0.10 | 0.07 - 0.28 | 0.07 - 0.11 |
| egg volume vs interval between pre-blastoderm mitoses | Hexapoda | 0.35 - 0.90 | 0.01 - 0.11 | 0.30 - 0.83 | 0.02 - 0.12 |
| egg volume vs time to cellularization | Hexapoda | 0.07 - 0.45 | 0.13 - 0.30 | 0.08 - 0.51 | 0.12 - 0.33 |
| egg length vs width | Hexapoda | 0 - <0.005 | 0.76 - 0.82 | 0 - <0.005 | 0.77 - 0.80 |
| egg length vs width | Hymenoptera | 0 - <0.005 | 0.67 - 0.78 | 0 - <0.005 | 0.70 - 0.78 |
| egg length vs width | Condylognatha | 0 - <0.005 | 0.79 - 0.89 | 0 - <0.005 | 0.78 - 0.91 |
| egg length vs width | Antliophora | 0 - <0.005 | 0.60 - 0.81 | 0 - <0.005 | 0.65 - 0.78 |
| egg length vs width | Neuropteroidea | 0 - <0.005 | 0.90 - 1.00 | 0 - <0.005 | 0.89 - 0.98 |
| egg length vs width | Amphiesmenoptera | 0 - <0.005 | 0.70 - 0.88 | 0 - <0.005 | 0.71 - 0.92 |
| egg length vs width | Polyneoptera | 0 - <0.005 | 0.69 - 0.78 | 0 - <0.005 | 0.69 - 0.75 |
| egg length vs width | Palaeoptera | 0 - <0.005 | 0.60 - 1.03 | 0 - <0.005 | 0.61 - 0.74 |
| egg length vs asymmetry, residuals to egg width | Hexapoda | 0 - 0.08 | 0.07 - 0.17 | 0 - 0.17 | 0.05 - 0.18 |
| egg length vs asymmetry, residuals to egg width | Hymenoptera | 0.01 - 0.97 | -0.11 - 0.19 | 0.05 - 1.00 | -0.15 - 0.18 |
| egg length vs asymmetry, residuals to egg width | Condylognatha | 0 - 0.54 | -0.09 - 0.49 | 0 - 0.95 | -0.01 - 0.45 |
| egg length vs asymmetry, residuals to egg width | Antliophora | 0 - 0.52 | 0.08 - 0.43 | 0 - 0.18 | 0.16 - 0.46 |
| egg length vs asymmetry, residuals to egg width | Neuropteroidea | 0.03 - 0.96 | -0.04 - 0.20 | 0.08 - 0.84 | 0.02 - 0.15 |
| egg length vs asymmetry, residuals to egg width | Amphiesmenoptera | 0.13 - 0.91 | -0.43 - -0.03 | 0.11 - 1.00 | -0.38 - 0.05 |
| egg length vs asymmetry, residuals to egg width | Polyneoptera | 0.12 - 1.00 | -0.15 - 0.13 | 0.13 - 0.98 | -0.13 - 0.13 |
| egg length vs asymmetry, residuals to egg width | Palaeoptera | 0.01 - 0.99 | -0.01 - 0.31 | 0.02 - 0.99 | -0.03 - 0.27 |
| egg length vs angle of curvature, residuals to egg width | Hexapoda | 0 - <0.005 | 0.39 - 0.57 | 0 - <0.005 | 0.41 - 0.58 |
| egg length vs angle of curvature, residuals to egg width | Hymenoptera | 0 - 0.04 | 0.33 - 0.91 | 0 - 0.01 | 0.40 - 0.77 |
| egg length vs angle of curvature, residuals to egg width | Condylognatha | 0 - 0.04 | 0.34 - 0.91 | 0 - 0.02 | 0.38 - 0.75 |
| egg length vs angle of curvature, residuals to egg width | Antliophora | 0.01 - 0.93 | -0.06 - 0.57 | 0.01 - 0.82 | 0.05 - 0.55 |
| egg length vs angle of curvature, residuals to egg width | Neuropteroidea | 0 - 0.24 | 0.17 - 0.49 | 0 - 0.11 | 0.22 - 0.52 |
| egg length vs angle of curvature, residuals to egg width | Amphiesmenoptera | 0.21 - 1.00 | -0.34 - 0.24 | 0.21 - 0.99 | -0.32 - 0.16 |
| egg length vs angle of curvature, residuals to egg width | Polyneoptera | 0 - 0.55 | 0.10 - 0.71 | 0 - 0.04 | 0.33 - 0.71 |
| egg length vs angle of curvature, residuals to egg width | Palaeoptera | 0.10 - 0.89 | -0.09 - 0.29 | 0.03 - 0.68 | 0.07 - 0.37 |
| egg length vs width, residuals to body size | Hexapoda | 0 - <0.005 | 0.67 - 0.75 | 0 - <0.005 | 0.67 - 0.75 |
| egg length vs width, residuals to body size | Hymenoptera | 0 - <0.005 | 0.66 - 0.92 | 0 - <0.005 | 0.69 - 0.90 |
| egg length vs width, residuals to body size | Condylognatha | 0 - 0.11 | 0.32 - 0.87 | 0 - 0.05 | 0.39 - 0.88 |
| egg length vs width, residuals to body size | Antliophora | 0 - <0.005 | 0.58 - 0.71 | 0 - <0.005 | 0.57 - 0.70 |
| egg length vs width, residuals to body size | Neuropteroidea | 0 - <0.005 | 0.90 - 1.05 | 0 - <0.005 | 0.91 - 1.07 |
| egg length vs width, residuals to body size | Amphiesmenoptera | 0 - <0.005 | 0.40 - 0.56 | 0 - <0.005 | 0.43 - 0.56 |
| egg length vs width, residuals to body size | Polyneoptera | 0 - <0.005 | 0.69 - 0.85 | 0 - <0.005 | 0.67 - 0.82 |
| egg length vs width, residuals to body size | Palaeoptera | 0.03 - 0.81 | 0.05 - 0.51 | 0.02 - 0.75 | 0.05 - 0.55 |
| egg volume vs cubic body length | Hexapoda | 0 - <0.005 | 0.41 - 0.47 | 0 - <0.005 | 0.43 - 0.48 |
| egg volume vs cubic body length | Hymenoptera | 0 - <0.005 | 0.62 - 0.77 | 0 - <0.005 | 0.63 - 0.81 |
| egg volume vs cubic body length | Condylognatha | 0 - <0.005 | 0.54 - 0.74 | 0 - <0.005 | 0.57 - 0.77 |
| egg volume vs cubic body length | Antliophora | 0.02 - 0.22 | 0.15 - 0.29 | 0.02 - 0.19 | 0.17 - 0.31 |
| egg volume vs cubic body length | Neuropteroidea | 0 - <0.005 | 0.36 - 0.45 | 0 - <0.005 | 0.36 - 0.45 |
| egg volume vs cubic body length | Amphiesmenoptera | 0 - <0.005 | 0.33 - 0.44 | 0 - <0.005 | 0.34 - 0.46 |
| egg volume vs cubic body length | Polyneoptera | 0 - <0.005 | 0.48 - 0.60 | 0 - <0.005 | 0.51 - 0.64 |
| egg volume vs cubic body length | Palaeoptera | 0 - 0.09 | 0.22 - 0.40 | 0 - 0.02 | 0.28 - 0.42 |

Table S23: **Results of PGLS analysis using a Blomberg correlation structure with a fixed deceleration parameter at 1.3**
