## Supplementary material for "Insect egg size and shape evolve with ecology, not developmental rate"

### Egg Database Bibliography

*Note:* This document is a list of the 1,756 published sources that were used to generate the assembled dataset of insect egg traits. 'Diss.' indicates a PhD dissertation, whereas 'MA thesis' indicates a Master's thesis. For more information on the dataset, please see Church et al. 2018: "A database of egg size and shape from more than 6,700 insect species" *(preprint)*. It describes the criteria that were used to include sources, definitions of each trait, and details on the procedure that was used to collect data from each source.

* Samuel H. Church and Seth Donoughe contributed equally to this work.
1 Department of Organismic and Evolutionary Biology, Harvard University, Cambridge, MA 02138, United States
2 *Current address:* Department of Cell and Molecular Biology, University of Chicago, Chicago, IL 60637, United States
3 Department of Molecular and Cellular Biology, Harvard University, Cambridge, MA 02138, United States

Aiello, A. and M. Balcazar-lara. "The immature stages of Oxytenis modestia with comments of the larvae of Asthenidia and Homoeopteryx (Saturniidae: Oxyteninae)". *Journal of The Lepidopterists' Society* 51.1-1 (1997): 105–118.

Aiken, R. B. "Effects of temperature on incubation times and mortality rates of eggs of Dytiscus alaskanus (Coleoptera: Dytiscidae)". *Ecography* 9.2 (1986): 133–136.

Aina, J. O. "The life history of Riptortus dentipes F.(Alydidae, Heteroptera) a pest of growing cowpea pods". *Journal of Natural History* 9.5 (1975): 589–596.

Aitken, A. D. "A strain of small Oryzaephilus surinamensis (L.) (Coleoptera, Silvanidae) from the Far East". *Journal of Stored Products Research* 2.1 (1966): 45–55.

Ajidagba, P. O. A. "Embryonic development of the stable fly, Stomoxys calcitrans Linnaeus (Diptera: Muscidae): a light and electron microscopy study". MA thesis. Kansas State University, 1979.

Ajidagba, P., C. W. Pitts, and D. E. Bay. "Early embryogenesis in the stable fly (Diptera: Muscidae)". *Annals of the Entomological Society of America* 76.4 (1983): 616–623.

Al Bitar, L., S. N. Gorb, C. P. W. Zebitz, and D. Voigt. "Egg adhesion of the codling moth Cydia pomonella L. (Lepidoptera, Tortricidae) to various substrates: II. Fruit surfaces of different apple cultivars". *Arthropod-Plant Interactions* 6.3 (2014): 471–488.

Alba-Tercedor, J. and M. El Alami. "Description of the nymphs and eggs of Acentrella almohades sp. n. from Morocco and Southern Spain (Ephemeroptera: Baetidae)". *Aquatic Insects* 21.4 (1999): 241–247.

Alexander, B. and J. G. Rozen Jr. "Ovaries, ovarioles, and oocytes in parasitic bees (Hymenoptera: Apoidea)". *The Pan-Pacific Entomologist* 63.2 (1987): 155–164.

Alford, D. V. "The biology and immature stages of Syntretus splendidus (Marshall) (Hymenoptera: Braconidae, Euphorinae), a parasite of adult bumblebees". *Transactions of the Royal Entomological Society of London* 120.17 (1968): 375–393.

Allsopp, P. G. "Identification of false wireworms (Coleoptera: Tenebrionidae) from southern Queensland and northern New South Wales". *Australian Journal of Entomology* 18.3 (1980): 277–286.

Allsopp, P. G. and L. N. Robertson. "Biology, ecology and control of soldier flies Inopus spp (Diptera, Stratiomyidae) - a review". *Australian Journal of Zoology* 36.6 (1988): 627–648.

Almeida, F., L. Suesdek, M. T. Motoki, E. S. Bergo, and M. A. M. Sallum. "Morphometric comparisons of the scanning electron micrographs of the eggs of Anopheles (Nyssorhynchus) darlingi Root (Diptera: Culicidae)". *Acta Tropica* 139 (2014): 115–122.

Altahtawy, M. M., S. M. El-Sawaf, and F. F. Shalaby. "Studies on morphology and development of the immature stages of Microplitis rufiventris Kokujev (Hym., Bracon.)" *Zeitschrift für Angewandte Entomologie* 71.1-4 (1972): 134–140.

Altson, A. M. "On the method of oviposition and the egg of Lyctus brunneus, Steph." *Journal of the Linnean Society of London, Zoology* 35.234 (1923): 217–227.

Alves-dos-Santos, I., G. A. R. Melo, and J. G. Rozen Jr. "Biology and immature stages of the bee tribe Tetrapediini (Hymenoptera: Apidae)". *American Museum Novitates* 3377 (2002): 1–45.

Amos, W. B. and G. Salt. "An atlas of the development of eggs of an ichneumon wasp". *Journal of Entomology Series B, Taxonomy* 43.1 (1974): 11–18.

Amy, R. L. "The embryology of Habrobracon juglandis (Ashmead)". *Journal of Morphology* 109.2 (1961): 199–217.

Andaur-Arenas, D. and T. S. Olivares. "Ultraestructura de huevos en cinco especies de macrolepidópteros con una clave de los huevos de Copitarsia Hampson (Lepidoptera, Ditrysia)". *Agrociencia* 43.1 (2009): 49–59.

Anderson, D. T. "The embryology of Dacus tryoni (Frogg.)[Diptera, Trypetidae (= Tephritidae)], the queensland fruit-fly". *Development* 10.3 (1962): 248–292.

Anderson, D. T. and C. Lawson-Kerr. "The embryonic development of the marine caddis fly, Philanisus plebeius Walker (Trichoptera: Chathamidae)". *The Biological Bulletin* 153.1 (1977): 98–105.

Anderson, J. M. E. "Aquatic Hydrophilidae (Coleoptera). The biology of some Australian species with descriptions of immature stages reared in the laboratory". *Australian Journal of Entomology* 15.2 (1976): 219–228.

Anderson, N. H. and J. R. Bourne. “Bionomics of three species of glossosomatid caddis flies (Trichoptera: Glossosomatidae) in Oregon”. *Canadian Journal of Zoology* 52.3 (1974): 405–411.

Ando, H., R. Machida, and T. Nagashima. “Cleavage of the jumping bristletail, Pedetontus unimaculatus Machida: Hexapoda, Microcoryphia, Machilidae”. *Bulletin of Seisen Women’s Junior College* 8-9 (1990): 197–202.

Ando, H. “Old oocytes and newly laid eggs of scorpion-flies and hanging-flies (Mecoptera: Panorpidae and Bittacidae)”. *Science Reports of the Tokyo Kyoiku Daigaku* 15 (1973): 163–187.

Ando, H. and H. Kobayashi. “Description of early and middle developmental stages in embryos of the firefly, Luciola cruciata Motschulsky (Coleoptera: Lampyridae)”. *Sugadaira Kogen Biological Laboratory Research Report* 7 (1975): 1–11.

Ando, H. and R. Machida. “Relationship between Notoptera and Dermaptera, from the embryological standpoint”. *Recent Advances in Insect Embryology in Japan and Poland*. Tsukuba: Arthropodan Embryological Society of Japan, 1987. 151–157.

Ando, H. and J. Okada. “Embryology of the butterbur stem sawfly Aglaeostigma occipitosa (Malaise) as studied by external observation (Tenthredinidae, Hymenoptera)”. *Acta Hymenopterologica* 1 (1958): 55–62.

Ando, H. and M. Tanaka. “The formation of germ rudiment and embryonic membranes in the primitive moth, Endoclita excrescens Butler (Hepialidae, Monotrysia, Lepidoptera) and its phylogenetic significance”. *Proceedings of the Japanese Society of Systematic Zoology* 12 (1976): 52–55.

Ando, H. and M. Tanaka. “Early embryonic development of the primitive moths, Endoclyta signifer Walker and E. excrescens Butler (Lepidoptera: Hepialidae)”. *International Journal of Insect Morphology and Embryology* 9.1 (1980): 67–77.

Ando, Y. “Structure of the chorion in the false melon beetle, Atrachya menetriesi Faldermann (Coleoptera: Chrysomelidae)”. *Kontyû* 41.4 (1973): 405–412.

Ansari, M. A., H. Casteels, L. Tirry, and M. Moens. “Biology of Hoplia philanthus (Col., Scarabaeidae, Melolonthinae): a new and severe pest in Belgian turf”. *Environmental Entomology* 35.6 (2006): 1500–1507.

Antunes, F. F., A. d. O. Menezes Jr, M. Tavares, and G. R. P. Moreira. “Morfologia externa dos estágios imaturos de heliconíneos neotropicais: I. Eueides isabella dianasa (Hübner, 1806)”. *Revista Brasileira de Entomologia* 46.4 (2002): 601–610.

Aparecido Pereira, R., M. Manfrini Morais, L. Domingos Gioli, F. Santos Nascimento, M. A. Rossi, and L. Rolandi Bego. “Comparative morphology of reproductive and trophic eggs in Melipona bees (Apidae, Meliponini)”. *Brazilian Journal of Morphological Sciences* 23.3-4 (2006): 349–354.

Arbogast, R. T., J. H. Brower, and R. G. Strong. “External morphology of the eggs of Tinea pallescentella Stainton, Tinea occidentella Chambers, and Niditinea fuscella (L.) (Lepidoptera: Tineidae)”. *International Journal of Insect Morphology and Embryology* 18.5 (1989): 321–328.

Arbogast, R. T., M. Carthon, and J. R. Roberts Jr. “Developmental stages of Xylocoris flavipes (Hemiptera: Anthocoridae), a predator of stored-product insects”. *Annals of the Entomological Society of America* 64.5 (1971): 1131–1134.

Arbogast, R. T., G. L. Lecato, and R. Van Byrd. “External morphology of some eggs of stored-product moths (Lepidoptera Pyralidae, Gelechiidae, Tineidae)”. *International Journal of Insect Morphology and Embryology* 9.3 (1980): 165–177.

Arbogast, R. T. and R. Van Byrd. “External morphology of the egg of the pink scavenger caterpillar, Pyroderces rileyi (Wallsingham) (Lepidoptera: Cosmopterigidae)”. *International Journal of Insect Morphology and Embryology* 15.3 (1986): 165–169.

Arbogast, R. T. and R. Van Byrd. “External morphology of the eggs of the meal moth, Pyralis farinalis (L.), and the murky meal moth, Aglossa caprealis (Hübner) (Lepidoptera: Pyralidae)”. *International Journal of Insect Morphology and Embryology* 10.5-6 (1981): 419–423.

Archangelsky, M. “Description of the immature stages of three Nearctic species of the genus Berosus Leach (Coleoptera: Hydrophilidae)”. *Internationale Revue der gesamten Hydrobiologie und Hydrographie* 79.3 (1994): 357–372.

Archangelsky, M. “Description of the preimaginal stages of Dactylosternum cacti (Coleoptera: Hydrophilidae, Sphaeridiinae)”. *Insect Systematics & Evolution* 25.2 (1994): 121–128.

Archangelsky, M. “Larvae of Neotropical Berosus (Coleoptera, Hydrophilidae): B. aulus Orchymont, 1941 and B. auriceps Boheman, 1859”. *Tijdschrift voor Entomologie* 142.1-2 (1999): 1–8.

Archangelsky, M. and M. E. Durand. “Description of the immature stages and biology of Phaenonotum exstriatum (Say 1835) (Coleoptera: Hydrophilidae: Sphaeridiinae)”. *The Coleopterists' Bulletin* 46.3 (1992): 209–215.

Archangelsky, M. and M. E. Durand. “Description of the preimaginal stages of Derallus angustus Sharp, 1882 (Coleoptera: Hydrophilidae, Berosinae)”. *Aquatic Insects* 14.3 (1992): 169–178.

Archangelsky, M. and L. A. Fernandez. “Description of the preimaginal stages and biology of Phaenonotum (Hydroglobus) puncticolle Bruch (Coleoptera: Hydrophilidae)”. *Aquatic Insects* 16.1 (1994): 55–63.

Archangelsky, M. and M. Fikáček. “Descriptions of the egg case and larva of Anacaena and a review of the knowledge and relationships between larvae of Anacaenini (Coleoptera: Hydrophilidae: Hydrophilinae)”. *European Journal of Entomology* 101.4 (2004): 629–636.

Arcila, A. M., L. A. Gómez, and P. Ulloa-Chacón. “Immature development and colony growth of crazy ant Paratrechina fulva under laboratory conditions (Hymenoptera: Formicidae)”. *Sociobiology* 39.2 (2002): 307–322.

Arivoli, S., A. N. Upadhyay, and P. Venkatesan. “Scanning electron microscopy on eggshell and eclosion process of Tenagogonus fluviorum (Fabricius) (Hemiptera: Gerridae)”. *Advances in Biological Research* 5.6 (2011): 309–314.

Armstrong, J. S. “The breeding habits of the Corduliidae (Odonata) in the Taupo district of New Zealand”. *Transactions of the Royal Society of New Zealand* 85.2 (1958): 275–282.

Arthur, A. P. “The cleptoparasitic habits and the immature stages of Eurytoma pini Bugbee (Hymenoptera: Chalcidae), a parasite of the European pine shoot moth, Rhyacionia buoliana (Schiff.) (Lepidoptera: Olethreutidae)”. *The Canadian Entomologist* 93.8 (1961): 655–660.

Arthur, A. P. “Development, behaviour, and descriptions of immature stages of Spilochalcis side (Walk.) (Hymenoptera: Chalcididae)”. *The Canadian Entomologist* 90.10 (1958): 590–595.

Arthur, A. P. and P. G. Mason. “Life history and immature stages of the parasitoid Microplitis mediator (Hymenoptera: Braconidae), reared on the bertha armyworm Mamestra configurata (Lepidoptera: Noctuidae)”. *The Canadian Entomologist* 118.5 (1986): 487–491.

Arthur, A. P. and Y. M. Powell. “Descriptions of the immature stages and adult reproductive systems of Athrycia cinerea (Coq.) (Diptera: Tachinidae), a native parasitoid of Mamestra configurata (Walk.) (Lepidoptera: Noctuidae)”. *The Canadian Entomologist* 121.12 (1989): 1117–1123.

Asaba, H. and H. Ando. “Ovarian structures and oogenesis in Lepidocampa weberi Oudemans (Diplura: Campodeidae)”. *International Journal of Insect Morphology and Embryology* 7.5-6 (1978): 405–414.

Asano, M. “Morphology and biology of Nepachys japonicus (Kiesenwetter) (Coleoptera: Malachiidae: Attalini): The foetomorphic larva in Malachiidae, III”. *Japanese Journal of Systematic Entomology* 20.2 (2014): 193–200.

Ashby, D. G. and D. W. Wright. “The immature stages of the carrot fly”. *Ecological Entomology* 97.14 (1946): 355–379.

Ashe, J. S. “Studies of the life history and habits of Phanerota fasciata Say (Coleoptera: Staphylinidae: Aleocharinae) with notes on the mushroom as a habitat and descriptions of the immature stages”. *The Coleopterists' Bulletin* 35.1 (1981): 83–96.

Asís, J. D., S. F. Gayubo, and J. Tormos. “Notes on the natural history of Stizus perrisii ibericus Beaumont (Hymenoptera: Sphecidae)”. *Journal of Natural History* 25.5 (1991): 1331–1337.

Askew, R. R. and J. M. Ruse. “The biology of some Cecidomyiidae (Diptera) galling the leaves of birch (Betula) with special reference to their chalcidoid (Hymenoptera) parasites”. *Ecological Entomology* 126.2 (1974): 129–167.

Auten, M. “The early embryological development of Phormia regina: Diptera (Calliphoridae)”. *Annals of the Entomological Society of America* 27.3 (1934): 481–506.

Avelar, T. and M. T. Rocha Pité. “Egg size and number in Drosophila subobscura under semi-natural conditions”. *Evolutionary Biology* 3 (1989): 37–48.

Avilla, J., G. Viggiani, X. Diaz, and M. J. Sarasua. “Morphological and biological notes on Encarsia meritoria Gahan (Hymenoptera, Aphelinidae), a parasitoid of Trialeurodes vaporariorum (Westwood) (Homoptera, Aleyrodidae) new in Europe”. *Biocontrol Science and Technology* 1.4 (1991): 289–295.

Awad, H., H. Elelimy, A. H Omar, and A. A Meguid. “Some biological parameters and morphological descriptions study on the milkweed bug, Spilostethus Pandurus Scop., (Hemiptera: Lygaeidae)”. *Wulfenia* 20.5 (2013): 169–187.

Baehrecke, E. H. and M. R. Strand. “Embryonic morphology and growth of the polyembryonic parasitoid Copidosoma floridanum (Ashmead) (Hymenoptera: Encyrtidae)”. *International Journal of Insect Morphology and Embryology* 19.3-4 (1990): 165–175.

Baerends, G. P. and J. M. Baerends van Roon. “Embryological and ecological investigations on the development of the egg of Ammophila campestris Jur”. *Tijdschrift voor Entomologie* 92 (1949): 53–112.

Bahia, A. C., N. F. C. Secundino, J. C. Miranda, D. B. Prates, A. P. A. Souza, F. F. Fernandes, A. Barral, P. F. P. Pimenta, A. Barral, and P. F. P. Pimenta. “Ultrastructural comparison of external morphology of immature stages of Lutzomyia (Nyssomyia) intermedia and Lutzomyia (Nyssomyia) whitmani (Diptera: Psychodidae), vectors of cutaneous leishmaniasis, by scanning electron microscopy”. *Journal of Medical Entomology* 44.6 (2007): 903–914.

Baig, M. M., A. K. Dubey, and V. V. Ramamurthy. “Biology and morphology of life stages of three species of whiteflies (Hemiptera: Aleyrodidae) from India”. *The Pan-Pacific Entomologist* 91.2 (2015): 168–183.

Baker, G. T., S. D. Hight, and R. L. Brown. “External morphology of the egg of the native (Melitara prodenialis) and exotic (Cactoblastis cactorum) cactus moths (Lepidoptera: Pyralidae)”. *Proceedings of the Entomological Society of Washington* 114.4 (2012): 433–438.

Baker, G. T., A. Lawrence, R. Kuklinski, and J. Goddard. “Morphological and ultrastructural characteristics of the chorion of Cimex lectularius Linnaeus (Hemiptera: Cimicidae)”. *Proceedings of the Entomological Society of Washington* 115.4 (2013): 325–332.

Baker, G. T. and P. W. K. Ma. “Morphology and chorionic fine structure of the egg of Neurocolpus nubilus (Hemiptera: Miridae)”. *Transactions of the American Microscopical Society* 113.1 (1994): 80–85.

Baker, G. T. and W. K. Ma. “Chorionic structure of Dineutes horni Rbts. (Coleoptera: Gyrinidae)”. *Italian Journal of Zoology* 54.3 (1987): 209–212.

Baker, J. R. “Development and sexual dimorphism of larvae of the bee genus Coelioxys”. *Journal of the Kansas Entomological Society* 44.2 (1971): 225–235.

Baker, J. R. and H. H. Neunzig. “The egg masses, eggs, and first-Instar larvae of eastern North American Corydalidae”. *Annals of the Entomological Society of America* 61.5 (1968): 1181–1187.

Ballerio, A. “Unusual morphology in a new genus and species of Ceratocanthinae from New Guinea (Coleoptera: Scarabaeoidea: Hybosoridae)”. *The Coleopterists Bulletin* 63.1 (2009): 44–53.

Balter, R. S. “The microtopography of avian lice eggs”. *Medical Biology Illustrated* 18.3 (1968): 166–179.

Bambara, S. B. and H. H. Neunzig. “Descriptions of immature stages of the grape root borer, Vitacea polistiformis (Lepidoptera: Sesiidae)”. *Annals of the Entomological Society of America* 70.6 (1977): 871–875.

Bantock, C. R. “Experiments on chromosome elimination in the gall midge, Mayetiola destructor”. *Development* 24.2 (1970): 257–286.

Baran, T. “Immature stages and bionomy of Scythris bifissella [Hofmann, 1889] [Lepidoptera: Scythrididae]”. *Polskie Pismo Entomologiczne* 71.3 (2002): 195–209.

Baran, T. “Life history and description of the preimaginal stages of Scythris siccella (Zeller, 1839) (Lepidoptera: Scythrididae)”. *Entomologica Fennica* 14.4 (2003): 211–219.

Baranowski, R. M. “Notes on the biology of Ischnodemus oblongus and I. fulvipes with descriptions of the immature stages (Hemiptera: Lygaeidae)”. *Annals of the Entomological Society of America* 72.5 (1979): 655–658.

Baranowski, R. M. and J. A. Slater. “The Pachygronthinae (Hemiptera: Lygaeidae) of Trinidad with the description of a new species and notes on other sedge feeding lygaeids”. *The Florida Entomologist* 65.4 (1982): 492–506.

Barata, J. M. S. "Morphological aspects of triatominae eggs: II. Macroscopic and exochorial characteristics of ten species of the genus Rhodnius Stal, 1859 (Hemiptera-Reduviidae)". *Revista de Saúde Pública* 15.5 (1981): 490–542.

Barber, G. W. M. "Observations on the egg and newly hatched larva of the corn ear worm on corn silk". *Journal of Economic Entomology* 34.3 (1941): 451–456.

Barber, G. W. M. and W. M. O. Ellis. "Eggs of three Cercopidae". *Psyche* 29.1 (1922): 1–3.

Barbier, R. and G. Chauvin. "The aquatic egg of Nymphula nymphaeata (Lepidoptera: Pyralidae)". *Cell and Tissue Research* 149.4 (1974): 473–479.

Barbosa, E. P., L. A. Kaminski, and A. V. L. Freitas. "Immature stages of the butterfly Diaethria clymena janeira (Lepidoptera: Nymphalidae: Biblidinae)". *Zoologia* 27.5 (2010): 696–702.

Barnes, A. M. and F. J. Radovsky. "A new Tunga (Siphonaptera) from the nearctic region with description of all stages". *Journal of Medical Entomology* 6.1 (1969): 19–36.

Barnes, J. K. "Biology and Immature Stages of Compsobata univitta (Walker, 1849) (Diptera: Micropezidae: Calobatinae)". *Proceedings of the Entomological Society of Washington* 117.4 (2015): 421–434.

Bartell, D. P. and B. C. Pass. "Morphology, development, and behavior of the immature stages of the parasite Bathyplectes anurus (Hymenoptera: Ichneumonidae)". *The Canadian Entomologist* 112.5 (1980): 481–487.

Batelka, J. "New synonym and notes on the distribution of Metoecus javanus (Coleoptera: Ripiphoridae) Nové synonymum a poznámky k rozlíení druhu Metoecus javanus (Coleoptera: Ripiphoridae)". *Klapalekiana* 39.4 (2003): 199–203.

Batiz, M. F. R., A. M. M. de Remes Lenicov, and H. Hagedorn. "Description of the immature stages of the planthopper Lacertinella australis (Hemiptera: Delphacidae)". *Journal of Insect Science* 14.1 (2014): 1–9.

Baumann, R. W. "Studies on Utah stoneflies (Plecoptera)". *The Great Basin Naturalist* 33.2 (1973): 91–108.

Beaver, R. A. "The biology and immature stages of Entedon leucogramma (Ratzeburg) (Hymenoptera: Eulophidae), a parasite of bark beetles". *Proceedings of the Royal Entomological Society of London. Series A, General Entomology* 41.1-3 (1966): 37–41.

Beckage, N. E. and I. de Buron. "Extraembryonic membranes of the endoparasitic wasp Cotesia congregata: presence of a separate amnion and serosa". *The Journal of Parasitology* 80.3 (1994): 389–396.

Becnel, J. J. and S. W. Dunkle. "Evolution of micropyles in dragonfly eggs (Anisoptera)". *Odonatologica* 19.3 (1990): 235–241.

Bedding, R. A. "The immature stages of Rhinophorinae (Diptera: Calliphoridae) that parasitise British woodlice". *Transactions of the Royal Entomological Society of London* 125.1 (1973): 27–44.

Bedford, G. O. "Description and development of the eggs of two stick insects (Phasmatodea: Phasmatidae) from New Britain". *Austral Entomology* 15.4 (1977): 389–393.

Bedford, G. O. "The development of the egg of Didymuria cuilescens (Phasmatodea: Phasmatidae: Podacanthinae) embryology and determination of the stage at which first diapause occurs". *Australian Journal of Zoology* 18.2 (1970): 155–169.

Beeman, S. L. and D. M. Norris. "Embryogenesis of Xyleborus ferrugineus (Fabr.) (Coleoptera, Scolytidae). I. External morphogenesis of male and female embryos". *Journal of Morphology* 152.2 (1977): 177–219.

Beig, D., O. C. Bueno, R. A. da Cunha, and H. J. de Moraes. "Differences in quantity of food in worker and male brood cells ofScaptotrigona postica (Latr. 1807) (Hymenoptera, Apidae)". *Insectes Sociaux* 29.2 (1982): 189–194.

Belfiore, C., H. Barber-James, and E. Gaino. "The eggs of Afronurus Lestage, 1924 (Ephemeroptera: Heptageniidae): A cue for phylogenetic relationships". *Proceedings of the Xth International Conference on Ephemeroptera, XIV International Symposium on Plecoptera*. Magnolia Press, 2003. 113–116.

Beliavsky, G. "The study of Braula coeca". *Bee World* 10.6 (1929): 84–87.

Bellanger, Y. "A new stick insect of the genus Oncotophasma from Costa Rica (Phasmatodea, Diapheromeridae, Diapheromerinae)". *Bulletin de la Société Entomologique de France* 121.2 (2016): 141–148.

Bellanger, Y. and O. Conle. "A new stick insect from Costa Rica (Phasmatodea, Pseudophasmatidae, Xerosomatinae)". *Bulletin de la Société Entomologique de France* 118.4 (2013): 503–508.

Benedek, P. “On the Eurydema species in Hungary”. *Zeitschrift für Angewandte Entomologie* 61.1-4 (1968): 113–118.

Benítez-Mora, A. and T. S. Olivares. “Ultraestructura de los huevos de dos mariposas nocturnas de Chile: Ormiscodes socialis y Polythysana cinerascens (Lepidoptera: Saturniidae)”. *Revista de Biología Tropical* 54.4 (2006): 1085–1091.

Benito, M. J. S. and P. G. Sanz. “Immature stages of five species of the genus Exapion Bedel (Coleoptera: Brentidae, Apioninae) associated with the seeds of Genista (Tournfourt) and Cytisus L.(Fabaceae)”. *The Coleopterists' Bulletin* 53.1 (1999): 8–26.

Bennett, F. D. “Parasites of Ancylostomia stercorea (Zell.), (Pyralidae, Lepidoptera) a pod borer attacking pigeon pea in Trinidad”. *Bulletin of Entomological Research* 50.4 (1960): 737–757.

Benton, F. and D. Claugher. “The structure and surface properties of the eggshell of Megaselia imitatrix Borgmeier (Diptera, Phoridae) in relation to the respiration of the embryo”. *Physiological Entomology* 25.2 (2000): 133–140.

Berg, V. L. “The external morphology of the immature stages of the bee fly, Systoechus vulgaris Loew, (Diptera, Bombyliidae), a predator of grasshopper egg pods”. *The Canadian Entomologist* 72.9 (1940): 169–178.

Bernard, E. C. “Egg types and hatching of Eosentomon Berlese (Protura: Eosentomidae)”. *Transactions of the American Microscopical Society* 98.1 (1979): 123–126.

Bernhardt, J. L. “Color changes and development of eggs of rice stink bug (Hemiptera: Pentatomidae) in response to temperature”. *Annals of the Entomological Society of America* 102.4 (2009): 638–641.

Berrigan, D. “The allometry of egg size and number in insects”. *Oikos* 60.3 (1991): 313–321.

Berté, S. B. and G. Pritchard. “The life histories of Limnephilus externus Hagen, Anabolia bimaculata (Walker), and Nemotaulius hostilis (Hagen) (Trichoptera, Limnephilidae) in a pond in southern Alberta, Canada”. *Canadian Journal of Zoology* 64.10 (1986): 2348–2356.

Betz, O. and S. Fuhrmann. “Life history traits in different life forms of predaceous Stenus beetles (Coleoptera, Staphylinidae), living in waterside environments”. *Netherlands Journal of Zoology* 51.4 (2001): 371–393.

Bey-Bienko, G. “Descriptions of six new species of Palearctic Blattodea”. *Konowia* 14 (1935): 117–134.

Bianchi, F. M., V. C. Matesco, L. A. Campos, and J. Grazia. “External morphology of the egg and the first and fifth instars of Cyrtocoris egeris Packauskas & Schaefer (Hemiptera: Heteroptera: Pentatomidae: Cyrtocorinae)”. *Zootaxa* 2991 (2011): 29–34.

Biasotto, L. D., F. M. Bianchi, and L. A. Campos. “Morphology of immatures of Euschistus (Mitripus) grandis (Insecta: Hemiptera: Pentatomidae)”. *Zoologia* 30.3 (2013): 346–352.

Bicchierai, M. C. and E. Gaino. “Fine structure of the egg envelopes in Silo mediterraneus saturniae (Trichoptera, Goeridae)”. *Italian Journal of Zoology* 67.2 (2000): 141–146.

Bielenin, I. “Early stages of embryonic development in the weevil Polydrosus pterygomalis Boh. (Coleoptera: Curculionidae)”. *Polskie pismo Entomologiczne* 25.7 (1955): 93–113.

Biemont, J. C., G. Chauvin, and C. Hamon. “Ultrastructure and resistance to water loss in eggs of Acanthoscelides obtectus Say (Coleoptera: Bruchidae)”. *Journal of Insect Physiology* 27.10 (1981): 667–679.

Birchard, G. F. “Water vapor and oxygen exchange of praying mantis (Tenodera aridifolia sinensis) egg cases”. *Physiological Zoology* 64.4 (1991): 960–972.

Bittencourt, M. A. L. and E. Berti Filho. “Development of immature stages of Palmistichus elaeisis Delvare & LaSalle (Hymenoptera, Eulophidae) in Lepidoptera pupae”. *Revista Brasileira de Entomologia* 48.1 (2004): 65–68.

Blackburn, T. M. “Comparative and experimental studies of animal life history variation”. Diss. University of Oxford, 1990.

Bledsoe, L. W., R. V. Flanders, and C. R. Edwards. “Morphology and development of the immature stages of Pediobius foveolatus (Hymenoptera: Eulophidae)”. *Annals of the Entomological Society of America* 76.6 (1983): 953–957.

Blokhina, A. V., N. A. Rozhkova, and O. A. Timoshkin. “Phenological peculiarities and evolution of Baicalina bellicosa Mart. (Trichoptera, Apataniidae)–an endemic species of Lake Baikal”. *Hydrobiologia* 568.1 (2006): 103–106.

Blom, P. E. and W. H. Clark. “Phobetus desertus, a new melolonthine Scarabaeidae (Coleoptera) from the Central Desert of Baja California, Mexico”. *The Pan-Pacific Entomologist* 60.4 (1984): 304–312.

Blume, R. R. “Description of larva and notes on biology of Pseudocanthon perplexus (LeConte) (Coleoptera: Scarabaeidae)”. *The Coleopterists' Bulletin* 36.2 (1982): 250–254.

Bock, E. “Bildung und Differenzierung der Keimblätter bei Chrysopa perla (L.)” *Zoomorphology* 35.4 (1939): 615–700.

Bohart, G. E., W. P. Stephen, and R. K. Eppley. “The biology of Heterostylum robustum (Diptera: Bombyliidae), a parasite of the alkali bee”. *Annals of the Entomological Society of America* 53.3 (1960): 425–435.

Bohart, G. E. and N. N. Youssef. “Notes on the biology of Megachile (Megachiloides) umatillensis Mitchell (Hymenoptera: Megachilidae) and its parasites”. *Transactions of the Royal Entomological Society of London* 124.1 (1972): 1–19.

Boivin, G., C. Picard, and J. L. Auclair. “Preimaginal development of Anaphes n. sp. (Hymenoptera: Mymaridae), an egg parasitoid of the carrot weevil (Coleoptera: Curculionidae)”. *Biological Control* 3.3 (1993): 176–181.

Boivin, G. “Reproduction and immature development of egg parasitoids”. *Egg Parasitoids in Agroecosystems with Emphasis on Trichogramma*. Dordrecht: Springer Netherlands, 2009. 1–23.

Boivin, G. and M.-J. Gauvin. “Egg size affects larval performance in a coleopteran parasitoid”. *Ecological Entomology* 34.2 (2009): 240–245.

Boivin, G. and J.-P. Nénon. “Variability and inheritance of the chorionic tubercles on eggs of Listronotus oregonensis (Coleoptera: Curculionidae)”. *The Canadian Entomologist* 135.6 (2003): 765–774.

Bologna, M. A. and A. Di Giulio. “Egg and first instar larval morphology of Prionotolytta binotata (Péringuey, 1888), an endemic southern African species (Coleoptera: Meloidae)”. *African Entomology* 11.2 (2003): 213–219.

Bonduriansky, R. and R. J. Brooks. “Reproductive allocation and reproductive ecology of seven species of Diptera”. *Ecological Entomology* 24.4 (1999): 389–395.

Boring, C. A., B. J. Sharanowski, and M. J. Sharkey. “Maxfischeriinae: a new braconid subfamily (Hymenoptera) with highly specialized egg morphology”. *Systematic Entomology* 36.3 (2011): 529–548.

Boucek, Z. “A contribution to the biology of Eucharis adscendens (F.) (Hymenoptera)”. *Vestnik Ceskoslovenske Zoologicke Spolecoenosti* 20.1 (1956): 97–99.

Boudou-Saltet, P. “Oeuf, ponte et eclosion chez un Orthoptère cavernicole (Dolichopoda linderi Duf. Orth., Rhaph.)” *Le Bulletin de la Société d'Histoire Naturelle de Toulouse* 116 (1980): 44–51.

Bradley, E. L., M. W. MacGown, B. H. Ebel, and W. W. Neel. “Surface patterns of eggs of some cone-infesting Lepidoptera of Southeastern United States [Dioryctria, Eucosma and Cydia (= Laspeyresia), pests of Pinus spp.]” *Journal of the Georgia Entomological Society* 17.2 (1982): 255–259.

Bradshaw, W. E., C. M. Holzapfel, and T. O'Neill. “Egg size and reproductive allocation in the pitcherplant mosquito Wyeomyia smithii (Diptera: Culicidae)”. *Journal of Medical Entomology* 30.2 (1993): 384–390.

Bragg, P. E. “A description of the male and egg of Sipyloidea acutipennis (Bates, 1865) (Diapheromeridae: Necroscinae)”. *Phasmid Studies* 16.1 (2007): 1–5.

Bragg, P. E. “A new species of Lopaphus Westwood, described from Borneo (Insecta: Phasmida: Heteronemiidae: Necrosciinae)”. *Zoologische Mededelingen, Leiden* 69.9 (1995): 105–111.

Bragg, P. E. “A new subgenus of Orthomeria Kirby, 1904 and a new species from Danum Valley, Sabah”. *Phasmid Studies* 14.1 (2006): 12–19.

Bragg, P. E. “A redescription of Sosibia lysippus (Westwood, 1859)”. *Phasmid Studies* 15.1-2 (2007): 11–14.

Bragg, P. E. “A review and key to the genus Phenacephorus Brunner (Insecta: Phasmida: Heteronemiidae: Lonchodinae), including the description of two new species”. *Zoologische Mededelingen Leiden* 68.22 (1994): 231–248.

Bragg, P. E. “A review of the subfamily Korinninae (Phasmida: Pseudophasmatidae), with the description of a new species”. *Tijdschrift voor Entomologie* 138.1 (1995): 45–50.

Bragg, P. E. “New synonyms and new records of phasmids (Insecta: Phasmida) in Borneo”. *Raffles Bulletin of Zoology* 41.1 (1993): 31–46.

Bragg, P. E. "Notes on Necroscia affinis (Gray, 1835), Necroscia fragilis (Redtenbacher, 1908) and Necroscia pallida (Redtenbacher, 1908)". *Phasmid Studies* 17.1 (2008): 16–26.

Bragg, P. E. "The first description of the male and egg of Syringodes rubicundus (de Haan, 1842) (Phasmida: Diapheromeridae: Necrosciinae)". *Zoologische Mededelingen* 82 (2008): 255–260.

Braham, M. and T. Jardak. "Contribution a L'etude de la bio-ecologie du scolyte du pistachier Chaetoptelius vestitus muls & rey (Coleoptera, Scolytidae) dans les regions du centre et du sud tunisiens". *Revue Ezzaitouna* 13.1-2 (2012): 1–17.

Brauer, A. "Studies on the embryology of Bruchus quadrimaculatus, Fabr". *Annals of the Entomological Society of America* 18.3 (1925): 283–312.

Breland, O. P. "Podagrion mantis Ashmead and other parasites of praying mantid egg cases (Hym.: Chalcidoidea; Dipt.: Chloropidae)". *Annals of the Entomological Society of America* 34.1 (1941): 99–113.

Breland, O. P. and J. W. Dobson. "Specificity of mantid oothecae (Orthoptera: Mantidae)". *Annals of the Entomological Society of America* 40.4 (1947): 557–575.

Bresseel, J. and J. Constant. "The Picasso stick insect: A striking new species of Calvisia from Vietnam". *Belgian Journal of Entomology* 14 (2017): 1–18.

Bresseel, J. "Comments on the genus Neoclides Uvarov, 1940 with the description of a new species (Phasmatodea: Diapheromerinae: Necrosciinae)". *Polish Journal of Entomology* 82.1 (2013): 3–11.

Bresseel, J. and J. Constant. "Giant sticks from Vietnam and China, with three new taxa including the second longest insect known to date (Phasmatodea, Phasmatidae, Clitumninae, Pharnaciini)". *European Journal of Taxonomy* 104 (2014): 1–38.

Bridges, E. T. and B. C. Pass. "Biology of Draeculacephala mollipes (Homoptera: Cicadellidae)". *Annals of the Entomological Society of America* 63.3 (1970): 789–792.

Brittain, J. E., A. Lillehammer, and S. J. Saltveiti. "The effect of temperature on intraspecific variation in egg biology and nymphal size in the stonefly, Capnia atra (Plecoptera)". *Journal of Animal Ecology* 53 (1984): 161–169.

Britton, E. B. "On the larva of Sphaerius and the systematic position of the Sphaeriidae (Coleoptera)". *Australian Journal of Zoology* 14.6 (1966): 1193–1198.

Brock, P. D. "Studies on Australian stick-insects of the family Heteronemiidae, subfamily Lonchodinae, including the description of a new genus". *Journal of Orthoptera Research* 9 (2000): 51–55.

Brock, P. D. "Studies on the Australasian stick-insect genus Extatosoma Gray (Phasmida: Phasmatidae: Tropoderinae: Extatosomatini)". *Journal of Orthoptera Research* 10.2 (2001): 303–313.

Brock, P. D. "Studies on the stick-insect genus Eurycnema Audinet-Serville (Phasmida; Phasmatidae) with particular reference to Australian species". *Journal of Orthoptera Research* 7 (1998): 61–70.

Brock, P. D. "Taxonomic notes on giant southern African stick insects (Phasmida), including the description of a new Bactrododema species". *Annals of the Transvaal Museum* 41.1 (2004): 61–77.

Brock, P. D. "Three new species of South African stick insects (Phasmida)". *Journal of Orthoptera Research* 15.1 (2006): 37–44.

Brock, P. D. and N. Cliquennois. "A review of the genus Medaura Stal, 1875 (Phasmatida: Phasmatinae) including the description of a new species from Bangladesh". *Phasmid Studies* 9.1-2 (2000): 11–26.

Brock, P. D. and J. Hasenpusch. "Studies on the Australian stick-insect genus Onchestus Stål (Phasmida: Phasmatidae)". *Journal of Orthoptera Research* 14.1 (2005): 17–22.

Brock, P. D. and J. Hasenpusch. "Studies on the leaf insects (Phasmida: Phylliidae) of Australia". *Journal of Orthoptera Research* 11.2 (2002): 199–205.

Brock, P. D. and L. Lowe. "A Study of Stick-insects (Phasmida) from Kakadu National Park, Northern Territory, Australia". *Journal of Orthoptera Research* 7 (1998): 71–76.

Brock, P. D. and A. Shlagman. "The stick-insects (Phasmatodea) of Israel, including the description of a new species". *Israel Journal of Entomology* 28 (1994): 101–107.

Bronskill, J. F. "Embryogenesis of Mesoleius tenthredinis Morl.(Hymenoptera: Ichneumonidae)". *Canadian Journal of Zoology* 42.3 (1964): 439–453.

Bronskill, J. F. “Embryology of Pimpla turionellae (L.) (Hymenoptera: Ichneumonidae)”. *Canadian Journal of Zoology* 37.5 (1959): 655–688.

Brothers, D. J. “Biology and immature stages of Myrmosula parvula (Hymenoptera: Mutillidae)”. *Journal of the Kansas Entomological Society* 51.4 (1978): 698–710.

Brown, G. C., M. J. Sharkey, and D. W. Johnson. “Bionomics of Scymnus (Pullus) louisianae J. Chapin (Coleoptera: Coccinellidae) as a predator of the soybean aphid, Aphis glycines Matsumura (Homoptera: Aphididae)”. *Journal of Economic Entomology* 96.1 (2003): 21–24.

Brown, H. P. “Neocylloepus, a new genus from Texas and Central America (Coleoptera: Dryopoidea: Elmidae)”. *The Coleopterists’ Bulletin* 24.1 (1970): 1–29.

Brown, H. P. “The life history of Climacia areolaris (Hagen), a Neuropterous’ parasite’of fresh water sponges”. *American Midland Naturalist* 47.1 (1952): 130–160.

Brown, H. P. and C. M. Murvosh. “Lutrochus arizonicus new species, with notes on ecology and behavior (Coleoptera, Dryopoidea, Limnichidae)”. *Annals of the Entomological Society of America* 63.4 (1970): 1030–1035.

Brown, K. W. and J. T. Doyen. “Review of the genus Microschatia (Solier) (Tenebrionidae: Coleoptera)”. *Journal of the New York Entomological Society* 99.4 (1991): 539–582.

Brown, V. K. “The biology and development of Brachygaster minutus Olivier (Hymenoptera: Evaniidae), a parasite of the oothecae of Ectobius spp.(Dictyoptera: Blattidae)”. *Journal of Natural History* 7.6 (1973): 665–674.

Bruder, K. W. and A. P. Gupta. “Biology of the pavement ant, Tetramorium caespitum (Hymenoptera: Formicidae)”. *Annals of the Entomological Society of America* 65.2 (1972): 358–367.

Brust, M. L., W. W. Hoback, and C. B. Knisley. “Biology, habitat preference, and larval description of Cicindela cursitans LeConte (Coleoptera: Carabidae: Cicindelinae)”. *The Coleopterists Bulletin* 59.3 (2005): 379–390.

Bryson, H. R. and G. F. Dillon. “Observations on the morphology of the corn seed beetle (Agonoderus pallipes Fab., Carabidae)”. *Annals of the Entomological Society of America* 34.1 (1941): 43–50.

Bubala, M. “Ecology of Hylecoetus dermestoides Linnaeus (Coleoptera: Lymexylidae) in North East Scotland”. MA thesis. University of Edinburgh, 1988.

Buck, M. “A new family and genus of acalypterate flies from the Neotropical region, with a phylogenetic analysis of Carnoidea family relationships (Diptera, Schizophora)”. *Systematic Entomology* 31.3 (2006): 377–404.

Budrienė, A., E. Budrys, and Ž. Nevronytė. “Sexual size dimorphism in the ontogeny of the solitary predatory wasp Symmorphus allobrogus (Hymenoptera: Vespidae)”. *Comptes Rendus Biologies* 336.2 (2013): 57–64.

Bull, A. L. “Stages of living embryos in the jewel wasp Mormoniella (Nasonia) vitripennis (Walker) (Hymenoptera: Pteromalidae)”. *International Journal of Insect Morphology and Embryology* 11.1 (1982): 1–23.

Bundy, C. S. and J. E. McPherson. “Life history and laboratory rearing of Corimelaena incognita (Hemiptera: Heteroptera: Thyreocoridae), with descriptions of immature stages”. *Annals of the Entomological Society of America* 102.6 (2009): 1068–1076.

Bundy, C. S. and J. E. McPherson. “Life history and laboratory rearing of Corimelaena obscura (Heteroptera: Thyreocoridae) with descriptions of immature stages”. *Annals of the Entomological Society of America* 90.1 (1997): 20–27.

Bundy, C. S. and J. E. McPherson. “Life history and laboratory rearing of Mecidea minor (Hemiptera: Heteroptera: Pentatomidae), with descriptions of immature stages”. *Annals of the Entomological Society of America* 104.4 (2011): 605–612.

Bundy, C. S. and R. M. McPherson. “Morphological examination of stink bug (Heteroptera: Pentatomidae) eggs on cotton and soybeans, with a key to genera”. *Annals of the Entomological Society of America* 93.3 (2000): 616–624.

Büning, J. “Reductions and new inventions dominate oogenesis of Strepsiptera (Insecta)”. *International Journal of Insect Morphology and Embryology* 27.1 (1998): 3–8.

Burkhart, C. N., W. Gunning, and C. G. Burkhart. “Scanning electron microscopic examination of the egg of the pubic louse (Anoplura: Pthirus pubis)”. *International Journal of Dermatology* 39.3 (2000): 201–202.

Burns, A. N. and A. Neboiss. “Two new species of Plecoptera from Victoria”. *Memoirs of the National Museum of Victoria* 212 (1957): 91–242.

Buschman, L. L. “Biology of the firefly Pyractomena lucifera (Coleoptera: Lampyridae)”. *The Florida Entomologist* 67.4 (1984): 529–542.

Bushing, R. W. “A synoptic list of the parasites of Scolytidae (Coleoptera) in North America north of Mexico”. *The Canadian Entomologist* 97.5 (1965): 449–492.

Bushland, R. C. “A study of the sculpture of the chorion of the eggs of eighteen South Dakota grasshoppers (Acridiidae)”. MA thesis. South Dakota State College of Agriculture and Mechanic Arts, 1934.

Butt, F. H. “Embryology of Sciara (Sciaridae: Diptera)”. *Annals of the Entomological Society of America* 27.4 (1934): 565–579.

Butt, F. H. “Embryology of the milkweed bug: Oncopeltus Fasciatus (Hemiptera)”. *Cornell University Agricultural Experiment Station Memoir* 283 (1949): 1–43.

Byers, G. W. “The life history of Panorpa nuptialis (Mecoptera: Panorpidae)”. *Annals of the Entomological Society of America* 56.2 (1963): 142–149.

Byrne, M. “The immature stages of Philonthus sanamus Tottenham (Coleoptera: Staphylinidae)”. *African Entomology* 1.2 (1993): 229–234.

Cabrera, N., A. J. Sosa, J. Dorado, and M. Julien. “Systena nitentula (Coleoptera: Chrysomelidae), a flea beetle injurious to Alternanthera philoxeroides (Amaranthaceae): redescription, biology, and distribution”. *Annals of the Entomological Society of America* 98.5 (2005): 643–652.

Çakici, Ö. and G. Ergen. “External egg morphology of Melanogryllus desertus (Pallas, 1771) (Orthoptera: Gryllidae)”. *Biharean Biologist* 6.2 (2012): 122–125.

Caldas, B.-H. C., L. R. Redaelli, and L. M. G. Diefenbach. “Description of immature stages of Corecoris dentiventris Berg (Hemiptera: Coreidae)”. *Anais da Sociedade Entomológica do Brasil* 27.3 (1998): 405–412.

Calvert, P. D., J. H. Tsai, and S. W. Wilson. “Delphacodes nigrifacies (Homoptera: Delphacidae): Field biology, laboratory rearing and descriptions of immature stages”. *The Florida Entomologist* 70.1 (1987): 129–134.

Calvo, D. and J. M. A. Molina. “Fecundity-body size relationship and other reproductive aspects of Streblote panda (Lepidoptera: Lasiocampidae)”. *Annals of the Entomological Society of America* 98.2 (2005): 191–196.

Calvo, D. and J. M. Molina. “Morphological aspects of developmental stages of Streblote panda (Lepidoptera: Lasiocampidae)”. *Annales de la Société Entomologique de France* 44.1 (2008): 37–46.

Cameron, A. E. “Bionomics of the Tabanidae (Diptera) of the Canadian prairie”. *Bulletin of Entomological Research* 17.1 (1926): 1–42.

Campos, L. A., J. Grazia, T. d. A. Garbelotto, F. M. Bianchi, and N. C. Lanzarini. “A new South American species of Banasa Stål (Hemiptera: Heteroptera: Pentatomidae: Pentatominae): from egg to adult”. *Zootaxa* 2559 (2010): 47–57.

Candan, S. and Z. Suludere. “Scanning electron microscope studies of the eggs of Psacasta exanthematica Scopoli, 1763 (Hemiptera: Heteroptera: Scutelleridae)”. *Polish Journal of Entomology* 72 (2003): 241–247.

Candan, S., Z. Suludere, A. Hasbenli, N. Çagiran, R. Lavigne, and A. Scarbrough. “Ultrastructure of the chorion of Dioctria flavipennis meigen, 1820 (Diptera: Asilidae: Stenopogoninae) compared with those of fourteen asilid speciesfrom the mid-Atlantic region of North America”. *Proceedings of the Entomological Society of Washington* 106.4 (2004): 811–825.

Candan, S. “Piezodorus îituratus (F.) (Heteroptera: Pentatomidae) yumurtalannin dis morfolojisi”. *Türkiye Entomoloji Dergisi* 22.4 (1998): 307–313.

Candan, S., D. Durak, Z. Suludere, and Y. Kalender. “External morphology of the eggs of Coreus marginatus (Linneaus, 1758) (Heteroptera: Coreidae)”. *Türkiye Entomoloji Dergisi* 27.3 (2003): 163–170.

Candan, S. and Z. Suludere. “Apodiphus amygdali (Germar, 1817) (Heteroptera: Pentatomidae) yumurtalarının yüzey morfolojisi”. *Türkiye Entomoloji Dergisi* 34.1 (2010): 67–74.

Candan, S. and Z. Suludere. “Chorion morphology of eggs of Aelia albovittata Fieber, 1868 and Aelia rostrata Boheman, 1852 (Heteroptera: Pentatomidae)”. *Journal of the Entomological Research Society* 8.1 (2006): 61–71.

Candan, S. and Z. Suludere. “External morphology of eggs of Carpocoris pudicus (Poda, 1761) (Heteroptera, Pentatomidae)”. *Journal of the Entomological Research Society* 1.2 (1999): 21–26.

Candan, S., Z. Suludere, F. Acikgöz, and A. Hasbenli. “Chorion morphology of eggs of the North American stink bug Euschistus variolarius (Palisot de Beauvois, 1817) (Heteroptera: Pentatomidae): A scanning electron microscopy study”. *Entomological News* 116.3 (2005): 177–182.

Candan, S., Z. Suludere, and F. Bayrakdar. “Surface morphology of eggs of Euproctis chrysorrhoea (Linnaeus, 1758)”. *Acta Zoologica* 89.2 (2008): 133–136.

Candan, S., Z. Suludere, and D. Durak. “Ultrastructure of the eggs chorion of Ceraleptus obtusus (Brullé, 1839) (Heteroptera: Coreidae)”. *Ohio Journal of Science* 105 (2005): 138–141.

Candan, S., Z. Suludere, and M. Erbey. “Morphology of eggs and spermatheca of Odontotarsus purpureolineatus (Heteroptera, Scutelleridae)”. *Biologia* 62.6 (2007): 763–769.

Candan, S., Z. Suludere, M. Erbey, and F. S. Yilmaz. “Morphology of spermatheca and eggs of Coptosoma putoni Montandon, 1898 (Hemiptera: Plataspidae)”. *Turkish Journal of Entomology* 36.3 (2012): 321–333.

Candan, S., Z. Suludere, and M. Güllü. “Description of spermatheca and eggs of Eurygaster austriaca (Schrank, 1778) (Heteroptera: Scutelleridae), based on optical and scanning electron microscopy”. *Turkish Journal of Zoology* 35.5 (2011): 653–662.

Candan, S., Z. Suludere, Y. Kalender, and O. Eryilmaz. “Ultrastructure of the chorion of Echthistus cognatus (Loew, 1849) (Diptera, Asilidae)”. *Ohio Journal of Science* 104.4 (2004): 93–96.

Candan, S., Z. Suludere, H. Koç, and N. Kuyucu. “External morphology of eggs of Tipula (Lunatipula) decolor, Tipula (Lunatipula) dedecor, and Tipula (Acutipula) latifurca (Diptera: Tipulidae)”. *Annals of the Entomological Society of America* 98.3 (2005): 346–350.

Canterbury, L. E. and S. E. Neff. “Eggs of Sialis (Sialidae: Megaloptera) in eastern North America”. *The Canadian Entomologist* 112.4 (1980): 409–419.

Cantrell, B. K. “The immature stages of some Australian Sarcophaginae (Diptera: Sarcophagidae)”. *Australian Journal of Entomology* 20.3 (1981): 237–248.

Capinera, J. L. “Striped blister beetle, Epicauta vittata (Fabricius) (Coleoptera: Meloidae)”. *University of Florida IFAS Extension* EENY-280 (2003): 1–4.

Carcupino, M. and A. Lucchi. “Eggshell fine structure of Bradysia aprica (Winnertz) (Diptera: Sciaridae)”. *International Journal of Insect Morphology and Embryology* 24.1 (1995): 109–117.

Cardozo-De-Almeida, M., S. Castro-De-Souza, M. L. R. De Oliveira, S. A. S. De Almeida, T. C. M. Gonçalves, and J. R. Dos Santos-Mallet. “Ultrastructure and morphometry of eggs of Triatoma rubrovaria (Blanchard, 1843), Triatoma carcavalloi Juberg, Rocha & Lent, 1998 and Triatoma circummaculata (Stål, 1859) (Hemiptera-Reduviidae-Triatominae)”. *Zootaxa* 3750.4 (2013): 348–356.

Cargnus, E., F. Pavan, N. Mori, and M. Martini. “Identification and phenology of Hyalesthes obsoletus (Hemiptera: Auchenorrhyncha: Cixiidae) nymphal instars”. *Bulletin of Entomological Research* 102.5 (2012): 504–514.

Carignan, S., G. Boivin, and R. K. Stewart. “Developmental biology and morphology of Peristenus digoneutis Loan (Hymenoptera: Braconidae: Euphorinae)”. *Biological Control* 5.4 (1995): 553–560.

Carl, M. “Die Präimaginalstadien der Tenebrionidae. Teil 3: Beschreibung der Larven und Eier von sechs Arten aus Nordafrika und Lanzarote (Coleóptera: Tenebrionidae)”. *Koleopterologische Rundschau* 66 (1996): 199–214.

Caron, E., C. S. Ribeiro-Costa, and A. M. Linzmeier. “The egg morphology of some species of Sennius Bridwell (Coleoptera: Chrysomelidae: Bruchinae) based on scanning electron micrographs”. *Zootaxa* 556 (2004): 1–10.

Carrière, J. and O. Bürger. *Die Entwicklungsgeschichte der Mauerbiene (Chalicodoma muraria, Fabr.) im Ei*. Leipzig: Halle: Ehrhardt Karras, 1897.

Carriere, Y., S. Masaki, and D. A. Roff. “The coadaptation of female morphology and offspring size: a comparative analysis in crickets”. *Oecologia* 110.2 (1997): 197–204.

Carrillo, R. and M. E. Fresard. “Absorción de agua en huevos de Hylamorpha elegans Burm. (Coleoptera: Scarabaeidae) efecto de la temperatura”. *Agro Sur* 38.3 (2010): 194–198.

Carroll, L. E. and R. A. Wharton. “Morphology of the immature stages of Anastrepha ludens (Diptera: Tephritidae)”. *Annals of the Entomological Society of America* 82.2 (1989): 201–214.

Cary, P. R. L. "The biology of the weta Zealandosandrus gracilis:(Orthoptera: Stenopelmatidae) from the Cass region". MA thesis. University of Canterbury, 1981.

Casañas-Arango, A. D., E. E. Trujillo, R. D. Friesen, and A. M. R. Hernandez. "Field biology of Zapriothrica sp. Wheeler (Dipt., Drosophilidae), a pest of Passiflora spp. of high elevation possessing long tubular flowers". *Journal of Applied Entomology* 120.1-5 (1996): 111–114.

Casañas-Arango, A. D., E. E. Trujillo, A. M. Hernandez, and G. Taniguchi. "Field biology of Cyanotricha necyria Felder (Lep., Dioptidae), a pest of Passiflora spp., in southern Colombia's and Ecuador's Andean region". *Journal of Applied Entomology* 109.1-5 (1990): 93–97.

Cassani, J. R., D. H. Habeck, and D. L. Matthews. "Life history and immature stages of a plume moth Sphenarches anisodactylus (Lepidoptera: Pterophoridae) in Florida". *The Florida Entomologist* 73.2 (1990): 257–266.

Castro, D. d. C., A. Cicchino, M. A. de Hamity, and F. Ortiz. "A new species of Phtheiropoios Eichler, 1940 (Phthiraptera: Amblycera: Gyropidae) from Argentina, with a key to the males collected from Ctenomys (Mammalia: Rodentia) from South America". *Entomological News* 118.4 (2007): 377–384.

Caxambú, M. G. and L. M. de Almeida. "Description of the immature stages and redescription of Lamprosoma azureum Germar (Chrysomelidae, Lamprosomatinae)". *Revista Brasileira de Zoologia* 16.Supl. 1 (1999): 243–256.

Cedeño, P. E. and R. W. Flowers. "Heilipodus unifasciatus (Champion) (Coleoptera: Curculionidae: Molytinae: Hylobiini) attacking plantations of Ochroma pyramidale (Cavanilles Ex Lamarck) Urban (Malvaceae) in Ecuador". *The Coleopterists Bulletin* 66.4 (2012): 344–346.

Celary, W. "Biology of the solitary ground-nesting bee Melitta leporina (Panzer, 1799) (Hymenoptera: Apoidea: Melittidae)". *Journal of the Kansas Entomological Society* 79.2 (2006): 136–145.

Cervantes-Peredo, L. and G. Ortega-León. "Description of a new species of Neoadoxoplatys and immature stages of Neoadoxoplatys saileri Kormilev (Heteroptera: Pentatomidae) associated with bamboo". *Neotropical Entomology* 43.3 (2014): 236–244.

Cervantes, L. P. and I. Pacheco R. "Biology and description of a new species of Cholula (Heteroptera: Rhyparochromidae: Myodochini) associated with a fig in México". *Journal of the New York Entomological Society* 111.1 (2003): 41–47.

Chaboo, C. S., T. C. Nguyen, P. Jolivet, J. A. Santiago-Blay, and M. Schmitt. "Immatures of Hemisphaerota palmarum (Boheman), with discussion of the caudal processes and shield architecture in the tribe Hemisphaerotini (Chrysomelidae, Cassidinae)". *New developments in the biology of Chrysomelidae*. 2004. 171–184.

Chandler, A. E. F. "A preliminary key to the eggs of some of the commoner aphidophagous Syrphidae (Diptera) occurring in Britain". *Transactions of the Royal Entomological Society of London* 120.8 (1968): 199–217.

Chandrapatya, A. and G. T. Baker. "Morphology of the chorion of Sternocera aequisignata (Coleoptera: Buprestidae)". *Thai Journal of Agricultural Science* 32.4 (1999): 481–486.

Chang, C.-C., W.-C. Lee, C. E. Cook, G.-W. Lin, and T. Chang. "Germ-plasm specification and germline development in the parthenogenetic pea aphid Acyrthosiphon pisum: Vasa and Nanos as markers". *International Journal of Developmental Biology* 50.4 (2004): 413–421.

Chapman, R. F. "Egg pods from grasshoppers collected in southern Ghana". *Journal of the Entomological Society of Southern Africa* 24.2 (1961): 259–284.

Chapman, R. F. and I. A. D. Robertson. "The egg pods of some tropical African grasshoppers". *Journal of the Entomological Society of South Africa* 21.1 (1958): 85–112.

Chaudhuri, P. K. and A. Mazumdar. "On the biology of Halictophagus australensis Perkins, 1905 from India (Strepsiptera, Halictophagidae)". *Deutsche Entomologische Zeitschrift* 47.2 (2000): 203–215.

Chauvin, G. and R. Barbier. "Perméabilité et ultrastructures des oeufs de deux Lépidoptères Tineidae: Monopis rusticella et Trichophaga tapetzella". *Journal of Insect Physiology* 18.8 (1972): 1447–1462.

Chauvin, G., C. Hamon, M. Vancassel, and G. Vannier. "The eggs of Forficula auricularia L.(Dermaptera, Forficulidae): ultrastructure and resistance to low and high temperatures". *Canadian Journal of Zoology* 69.11 (1991): 2873–2878.

Chauvin, J. T. and G. Chauvin. "Formation des reliefs externes de l'oeuf de Micropteryx calthella L.(Lepidoptera: Micropterigidae)". *Canadian Journal of Zoology* 58.5 (1980): 761–766.

Cheke, R. A., L. D. C. Fishpool, and J. M. Ritchie. "An ecological study of the egg-pods of Oedaleus senegalensis (Krauss) (Orthoptera: Acrididae)". *Journal of Natural History* 14.3 (1980): 363–371.

Chen, H., D. Yang, D. Gu, S. G. Compton, and Y. Peng. "Secondary galling: a novel feeding strategy among 'non-pollinating'fig wasps from Ficus curtipes". *Ecological Entomology* 38.4 (2013): 381–389.

Chen, R. P.-Y., C.-L. Shih, G.-W. Lin, C. E. Cook, T.-Y. Huang, and C.-C. Chang. "Developmental expression of Apnanos during oogenesis and embryogenesis in the parthenogenetic pea aphid Acyrthosiphon pisum". *International Journal of Developmental Biology* 53.1 (2003): 169–176.

Chen, X. and A.-P. Liang. "Laboratory rearing of Callitettix versicolor (Hemiptera: Cicadomorpha: Cercopidae), with descriptions of the immature stages". *Annals of the Entomological Society of America* 105.5 (2012): 664–670.

Cheng, L. and R. L. Pitman. "Mass oviposition and egg development of the ocean-skater Halobates sobrinus (Heteroptera: Gerridae)". *Pacific Science* 56.4 (2002): 441–447.

Chernyakhovskii, M. E. "New and little known egg-pods of acridids (Orthoptera, Acrididae) of the fauna of Russia and adjacent countries". *Entomological Review* 86.6 (2006): 635–637.

Chernyakhovskii, M. E. "On the studies of Acridid egg pods (Acridoidea)". *Zoologicheskii Zhurnal* 66 (1987): 832–839.

Chiquetto-Machado, P. I. "Redescription of the Brazilian stick insect Pseudophasma cambridgei Kirby (Phasmatodea: Pseudophasmatidae), with first description of the female and egg". *Austral Entomology* (2017).

Cho, H. W. and J. E. Lee. "Gonioctena koryeoensis (Coleoptera: Chrysomelidae: Chrysomelinae), a new species from Korea, with a description of immature stages". *Zootaxa* 2438 (2010): 52–60.

Christensen, P. J. H. "Embryologische und zytologische Studien iiber die erste und friihe Eientwickelung bei Orgyia antiqua Linnt (Fam. Lymantridae, Lepidoptera)". *Videnskabelige Meddelelser Naturhistorisk Forening i København* 106 (1943): 1–223.

Christensen, P. J. H. *The embryonic development of cochlidion Limacodes Hufn. (Fam. Cochlididae, Lepidoptera); a study on living dated eggs*. Copenhagen: Ejnar Munksgaard, 1953.

Ciach, M. and J. Michalcewicz. "Egg morphology of Rosalia alpina (Linnaeus, 1758) (Coleoptera: Cerambycidae) from southern Poland". *Entomological News* 120.1 (2009): 61–64.

Cicchino, A. C. and D. d. C. Castro. "On Gyropus parvus parvus (Ewing, 1924) and Phtheiropoios rionegrensis sp. n.(Phthiraptera, Amblycera, Gyropidae), parasitic on Ctenomys haigi Thomas, 1919 (Mammalia, Rodentia, Ctenomyidae)". *Iheringia, Série Zoologia* 77 (1994): 3–14.

Clancy, D. W. "Insect parasites of the Chrysopidae (Neuroptera)". Diss. University of California, 1946.

Claridge, M. F. and R. R. Askew. "Sibling species in the Eurytoma rosae group (Hym., Eurytomidae)". *Entomophaga* 5.2 (1960): 141–153.

Clark, J. T. "The eggs of stick insects (Phasmida): a review with descriptions of the eggs of eleven species". *Systematic Entomology* 1.2 (1976): 95–105.

Clark, J. T. "The spruce bud midge, Rhabdophaga swainei Felt (Cecidomyiidae: Diptera)". *The Canadian Entomologist* 84.3 (1952): 87–89.

Clark, R. C. and N. R. Brown. "Studies of Predators of the Balsam Woolly Aphid, Adelges piceae (Ratz.) (Homoptera: Adelgidae), VII. Laricobius rubidus Lec. (Coleoptera: Derodontidae), a Predator of Pineus strobi (Htg.) (Homoptera: Adelgidae)". *The Canadian Entomologist* 92.3 (1960): 237–240.

Clausen, C. P. "The egg-larval host relationship among the parasitic Hymenoptera". *Bollettino del Laboratorio di Zoologia Generale e Agraria della R. Scuola Superiore d'Agricoltura in Portici* 33 (1954): 119–133.

Claypole, A. M. "The embryology of the apterygota". *Zoological Bulletin* 2.2 (1898): 69–76.

Cliquennois, N. and P. D. Brock. "Phasmids of Mauritius: Mauritiophasma n. gen., Monoiognosis n. gen., Epicharmus Stål 1875 and discussion on their remarkable eggs (Phasmatodea)". *Journal of Orthoptera Research* 13.1 (2004): 1–13.

Cobben, R. H. “Evolutionary trends in Heteroptera. Part I, Eggs, architecture of the shell, gross embryology and eclosion”. *Agricultural Research Reports* 707 (1968): 1–475.

Cobblah, M. A. and J. D. Hollander. “Specific differences in immature stages, oviposition sites and hatching patterns in two rice pests, Leptocorisa oratorius (Fabricius) and L. acuta (Thunberg) (Heteroptera: Alydidae)”. *International Journal of Tropical Insect Science* 13.1 (1992): 1–6.

Cogley, T. P. and J. R. Anderson. “Ultrastructure and function of the attachment organ of Gasterophilus eggs (Diptera: Gasterophilidae)”. *International Journal of Insect Morphology and Embryology* 12.1 (1983): 13–23.

Cogley, T. P., J. R. Anderson, and J. Weintraub. “Ultrastructure and function of the attachment organ of warble fly eggs (Diptera: Oestridae: Hypodermatinae)”. *International Journal of Insect Morphology and Embryology* 10.1 (1981): 7–18.

Cogley, T. P. “Key to the eggs of the equid stomach bot flies Gasterophilus Leach 1817 (Diptera: Gasterophilidae) utilizing scanning electron microscopy”. *Systematic Entomology* 16.2 (1991): 125–133.

Cogley, T. P. “Morphology of the eggs of the rhinoceros bot flies Gyrostigma conjungens and G. pavesii (Diptera: Gasterophilidae)”. *International Journal of Insect Morphology and Embryology* 19.5 (1990): 323–326.

Cohen, S., M. T. Greenwood, and J. A. Fowler. “The louse Trinoton anserinum (Amblycera: Phthiraptera), an intermediate host of Sarconema eurycerca (Filarioidea: Nematoda), a heartworm of swans”. *Medical and Veterinary Entomology* 5.1 (1991): 101–110.

Collier, T., S. Kelly, and M. Hunter. “Egg size, intrinsic competition, and lethal interference in the parasitoids Encarsia pergandiella and Encarsia formosa”. *Biological Control* 23.3 (2002): 254–261.

Colwell, D. D., C. R. Baird, B. Lee, and K. Milton. “Scanning electron microscopy and comparative morphometrics of eggs from six bot fly species (Diptera: Oestridae)”. *Journal of Medical Entomology* 36.6 (1999): 803–810.

Common, I. F. B. and E. D. Edwards. “The early stages of Gastridiota adoxima (Turner) (Lepidoptera: Bombycoidea) and its family placement”. *Australian Journal of Entomology* 30.2 (1991): 187–192.

Common, I. F. B. and N. McFarland. “A new subfamily for Munychryia Walker and Gephyroneura Turner (Lepidoptera: Anthelidae) and the description of a new species from Western Australia”. *Australian Journal of Entomology* 9.1 (1970): 11–22.

Compton, S. and A. B. Ware. “Ants disperse the elaisosome-bearing eggs of an African stick insect”. *Psyche* 98.2-3 (1991): 207–214.

Conci, C. and L. Tamanini. “Iconography of eggs of Italian Psylloidea (Insecta, Homoptera)”. *Atti dell'Academia Roveretana degli Agiati, Serie VII B* 10 (2000): 5–32.

Conci, C. and L. Tamanini. “Seven species of psylloidea new for Italy (Homoptera)”. *Annali del Museo Civico di Rovereto* 4 (1988): 307–320.

Condrashoff, S. F. “Description and morphology of the immature stages of three closely related species of Contarinia Rond. (Diptera: Cecidomyiidae) from galls on Douglas-fir needles”. *The Canadian Entomologist* 93.10 (1961): 833–851.

Conle, O. V. and F. H. Hennemann. “Studies of Neotropical Phasmatodea XII: Pseudophasma lakini sp. n.-a new stick insect from eastern Ecuador (Phasmatodea: Pseudophasmatidae: Pseudophasmatinae)”. *Polish Journal of Entomology* 81.1 (2012): 3–10.

Conle, O. V. and F. H. Hennemann. “Studies on neotropical Phasmatodea I: A remarkable new species of Peruphasma Conle & Hennemann, 2002 from Northern Peru (Phasmatodea: Pseudophasmatidae: Pseudophasmatinae)”. *Zootaxa* 1068.1 (2005): 59–68.

Conle, O. V., F. H. Hennemann, and P. Fontana. “Studies on neotropical Phasmatodea V: Notes on certain species of Pseudosermyle Caudell, 1903, with the descriptions of three new species from Mexico (Phasmatodea: Diapheromeridae: Diapheromerinae: Diapheromerini)”. *Zootaxa* 1496 (2007): 31–51.

Conle, O. V., F. H. Hennemann, H. Käch, and B. Kneubuhler. “Studies on neotropical Phasmatodea IX: Oreophoetes topoense n. sp.-Diapheromeridae: Diapheromerinae: Oreophoetini”. *Journal of Orthoptera Research* 18.2 (2009): 145–152.

Conle, O. V., F. H. Hennemann, H. Käch, and B. Kneubühler. "Studies on Neotropical Phasmatodea IX: Oreophoetes topoense n. sp.-a New Colorful Walking-Stick from Central Ecuador (Phasmatodea: Diapheromeridae: Diapheromerinae: Oreophoetini)". *Journal of Orthoptera Research* 18.1 (2009): 145–152.

Conle, O. V., F. H. Hennemann, and D. E. Perez-Gelabert. "Studies on neotropical Phasmatodea II: Revision of the genus Malacomorpha Rehn, 1906, with the descriptions of seven new species (Phasmatodea: Pseudophasmatidae: Pseudophasmatinae)". *Zootaxa* 1748 (2008): 1–64.

Conle, O. V., F. H. Hennemann, and D. E. Perez-Gelabert. "Studies on Neotropical Phasmatodea XV: A remarkable new stick insect from highly montane habitats of Hispaniola (Pseudophasmatidae: Xerosomatinae: Hesperophasmatini)". *Novitates Caribaea* 7 (2014): 28–36.

Conlong, D. E., D. Y. Graham, and H. Hastings. "Notes on the natural host surveys and laboratory rearing of Goniozus natalensis Gordh (Hymenoptera: Bethylidae), a parasitoid of Eldana saccharina Walker(Lepidoptera: Pyralidae) larvae from Cyperus papyrus L. in Southern Africa". *Journal of the Entomological Society of Southern Africa* 51.1 (1988): 115–127.

Cônsoli, F. L., E. W. Kitajima, and J. R. P. Parra. "Ultrastructure of the natural and factitious host eggs of Trichogramma galloi Zucchi and Trichogramma pretiosum Riley (Hymenoptera: Trichogrammatidae)". *International Journal of Insect Morphology and Embryology* 28.3 (1999): 211–231.

Cônsoli, F. L., C. T. Wuellner, S. B. Vinson, and L. E. Gilbert. "Immature development of Pseudacteon tricuspis (Diptera: Phoridae), an endoparasitoid of the red imported fire ant (Hymenoptera: Formicidae)". *Annals of the Entomological Society of America* 94.1 (2001): 97–109.

Constant, B., S. Grenier, and G. Bonnot. "Analysis of some morphological and biochemical characteristics of the egg of the predaceous bug Macrolophus caliginosus (Het.: Miridae) during embryogenesis". *Entomophaga* 39.2 (1994): 189–198.

Cooper, K. W. "A southern California Boreus, B. notoperates n. sp. I. Comparative morphology and systematics. (Mecoptera: Boreidae)". *Psyche* 79.4 (1972): 269–283.

Cooper, K. W. "Egg gigantism, oviposition, and genital anatomy: their bearing on the biology and phylogenetic position of Orussus (Hymenoptera: Siricoidea)". *Proceedings of the Rochester Academy of Science* 10 (1953): 38–68.

Cooper, K. W. "Ruptor ovi, the number of moults in development, and method of exit from masoned nests. Biology of Eumenine wasps, VII". *Psyche* 73.4 (1966): 238–250.

Cooper, W. R. and D. W. Spurgeon. "Oviposition behaviors and ontogenetic embryonic characteristics of the western tarnished plant bug, Lygus hesperus". *Journal of Insect Science* 12.36 (2012): 1–11.

Corbet, S. A. "Gomphids from Cameroon, West Africa (Anisoptera: Gomphidae)". *Odonatologica* 6.2 (1977): 55–68.

Cordo, H. A., C. J. De Loach, R. Ferrer, and J. Briano. "Bionomics of Carmenta haematica (Ureta) (Lepidoptera: Sesiidae) which attacks snakeweeds (Gutierrezia spp) in Argentina". *Biological Control* 5.1 (1995): 11–24.

Cornelis, M., E. Quiran, and M. C. Coscaron. "The scentless plant bug, Liorhyssus hyalinus (Fabricius) (Hemiptera: Heteroptera: Rhopalidae): Description of immature stages and notes on its life history". *Zootaxa* 3525 (2012): 83–88.

Coshan, P. F. "The biology of Coleophora serratella (L.) (Lepidoptera: Coleophoridae)". *Transactions of the Royal Entomological Society of London* 126.2 (1974): 169–188.

Costello, S. L., P. D. Pratt, M. B. Rayachhetry, and T. D. Center. "Morphology and life history characteristics of Podisus mucronatus (Heteroptera: Pentatomidae)". *The Florida Entomologist* 85.2 (2002): 344–350.

Couri, M. S. "Immature stages of Fannia pusio (Wiedemann, 1830) (Diptera, Fanniidae)". *Revista Brasileira de Biologia* 52.1 (1992): 83–91.

Coville, R. E. "Biological observations on Trypoxylon (Trypargilum) orizabense Richards in Arizona (Hymenoptera: Sphecidae)". *Journal of the Kansas Entomological Society* 52.3 (1979): 613–620.

Cox, M. L. and D. M. Windsor. "The first instar larva of Aulacoscelis appendiculata n. sp. (Coleoptera: Chrysomelidae: Aulacoscelinae) and its value in the placement of the Aulacoscelinae". *Journal of Natural History* 33.7 (1999): 1049–1087.

Craig Jr, G. B. and W. R. Horsfall. "Eggs of floodwater mosquitoes. VII. Species of Aedes common in the southeastern United States (Diptera: Culicidae)". *Annals of the Entomological Society of America* 53.1 (1960): 11–18.

Craig, D. A. "The eggs and embryology of some New Zealand Blepharoceridae (Diptera, Nematocera) with reference to the embryology of other Nematocera". *Transactions of the Royal Society of New Zealand* 8.18 (1967): 191–206.

Cronin, J. T. and D. R. Strong. "Biology of Anagrus delicatus (Hymenoptera: Mymaridae), an egg parasitoid of Prokelisia marginata (Homoptera: Delphacidae)". *Annals of the Entomological Society of America* 83.4 (1990): 846–854.

Cruz-Landim, d. C. and M. A. Cruz-Höfling. "Cytochemical and ultrastructural studies on eggs from workers and queen of Trigona". *Brazilian Journal of Medical and Biological Research* 4.1-2 (1971): 19–25.

Cruz, M. and I. Martinez. "Data on nesting and preimaginal development in two Mexican species of Ataenius Harold, 1867 (Coleoptera, Scarabaeoidea, Aphodiidae; Eupariinae)". *Folia Entomologica Mexicana* 41.1 (2002): 1–5.

Cruz, Y. P. "Development of the polyembryonic parasite Copidosomopsis tanytmemus (Hymenoptera: Encyrtidae)". *Annals of the Entomological Society of America* 79.1 (1986): 121–127.

Cuccodoro, G. and I. Löbl. "Revision of the Palaearctic rove beetles of the genus Megarthrus Curtis (Coleoptera: Staphylinidae: Proteininae)". *Journal of Natural History* 31.9 (1997): 1347–1415.

Cullen, M. J. "The biology of giant water bugs (Hemiptera: Belostomatidae) in Trinidad". *Proceedings of the Royal Entomological Society of London. Series A, General Entomology* 44.7-9 (1969): 123–136.

Culliney, T. W. "Site of opposition and description of eggs of Sophonia rufofascia (Homoptera: Cicadellidae: Nirvaninae), a polyphagous pest in Hawai'i." *Proceedings of the Hawaiian Entomological Society* 33 (1998): 67–73.

Cumming, M. E. P. "Notes on the life history and seasonal development of the pine needle scale, Phenacaspis pinifoliae (Fitch) (Diaspididae: Homoptera)". *The Canadian Entomologist* 85.9 (1953): 347–352.

Da Rosa, J. A., J. M. S. Barata, J. L. F. Santos, and M. Cilense. "Egg morphology of Triatoma circummaculata and Triatoma rubrovaria (Hemiptera, Reduviidae)". *Revista de Saúde pública* 34.5 (2000): 538–542.

Da Silva, D. S., R. Dell'Erba, L. A. Kaminski, and G. R. P. Moreira. "External morphology of the immature stages of neotropical heliconians: V. Agraulis vanillae maculosa (Lepidoptera, Nymphalidae, Heliconiinae)". *Iheringia, Série Zoologia* 96.2 (2006): 219–228.

Dahlan, A. N. and G. Gordh. "Development of Trichogramma australicum Girault (Hymenoptera: Trichogrammatidae) in eggs of Helicoverpa armigera (Hiibner) (Lepidoptera: Noctuidae) and in artificial diet". *Australian Journal of Entomology* 37.3 (1998): 254–264.

Dahlan, A. N. and G. Gordh. "Development of Trichogramma australicum Girault (Hymenoptera: Trichogrammatidae) on Helicoverpa armigera (Hubner) eggs (Lepidoptera: Noctuidae)". *Australian Journal of Entomology* 35.4 (1996): 337–344.

Dahms, E. C. "A review of the biology of species in the genus Melittobia (Hymenoptera: Eulophidae) with interpretations and additions using observations on Melittobia australica". *Memoirs of the Queensland Museum* 21.2 (1984): 337–360.

Dallai, R., D. Mercati, M. Gottardo, A. T. Dossey, R. Machida, Y. Mashimo, and R. G. Beutel. "The male and female reproductive systems of Zorotypus hubbardi Caudell, 1918 (Zoraptera)". *Arthropod Structure & Development* 41.4 (2012): 337–359.

Darling, D. C. and T. D. Miller. "Life history and larval morphology of Chrysolampus (Hymenoptera: Chalcidoidea: Chrysolampinae) in western North America". *Canadian Journal of Zoology* 69.8 (1991): 2168–2177.

Darling, D. C. and H. Roberts. "Life history and larval morphology of Monacon (Hymenoptera: Perilampidae), parasitoids of ambrosia beetles (Coleoptera: Platypodidae)". *Canadian Journal of Zoology* 77.11 (1999): 1768–1782.

Das, G. M. “Preliminary studies on the biology of Orasema assectator Kerrich (Hym., Eucharitidae), parasitic on Pheidole and causing damage to leaves of tea in Assam”. *Bulletin of Entomological Research* 54.3 (1963): 373–378.

Datta, R. K. and P. K. Mukherjee. “Life history of Tricholyga bombycis (Diptera: Tachinidae), a parasite of Bombyx mori (Lepidoptera: Bombicidae)”. *Annals of the Entomological Society of America* 71.5 (1978): 767–770.

Davidovavilimova, J. “The eggs of 2 coptosoma species, with a review of the eggs of the Plataspidae (Heteroptera)”. *Acta Entomologica Bohemoslovaca* 84.4 (1987): 254–260.

Davies, D. M. and B. V. Peterson. “Observations on the mating, feeding, ovarian development, and oviposition of adult black flies (Simuliidae, Diptera)”. *Canadian Journal of Zoology* 34.6 (1956): 615–655.

Davis, C. C. “A comparative study of larval embryogenesis in the mosquito Culex fatigans wiedemann (Diptera: Culicidae) and the sheep-fly Lucilia sericata meigen (Diptera: Calliphoridae)”. *Australian Journal of Zoology* 15.3 (1967): 547–579.

Davis, C. C. “A study of the hatching process in aquatic invertebrates. XVIII. Eclosion in Helicopsyche borealis (Hagen) (Trichoptera, Helicopsychidae). XIX. Hatching in Psephenus herricki (De Kay) (Coleoptera, Psephenidae)”. *American Midland Naturalist* 74.2 (1965): 443–450.

Davis, C. C. “A study of the hatching process in aquatic invertebrates. XXV. Hatching in the blackfly, Simulium (probably venustum) (Diptera, Simuliidae)”. *Canadian Journal of Zoology* 49.3 (1971): 333–336.

Davis, D. R., O. Karsholt, N. P. Kristensen, and E. S. Nielsen. “Revision of the genus Ogygioses (Palaeosetidae)”. *Invertebrate Systematics* 9.6 (1995): 1231–1263.

Davis, D. R., D. A. Quintero, R. A. T. Cambra, and A. Aiello. “Biology of a new Panamanian bagworm moth (Lepidoptera: Psychidae) with predatory larvae, and eggs individually wrapped in setal cases”. *Annals of the Entomological Society of America* 101.4 (2008): 689–702.

Davis, R. B., J. Javoiš, J. Pienaar, E. Õunap, and T. Tammaru. “Disentangling determinants of egg size in the Geometridae (Lepidoptera) using an advanced phylogenetic comparative method”. *Journal of Evolutionary Biology* 25.1 (2012): 210–219.

De Alencar, A. P. P. and A. C. R. Leite. “Ultrastructure of the egg of Muscina stabulans and Synthesiomyia nudiseta (Diptera: Muscidae)”. *Memórias do Instituto Oswaldo Cruz* 87.4 (1992): 463–466.

De Almeida, D. N., R. da Silva Oliveira, B. G. Brazil, and M. J. Soares. “Patterns of exochorion ornaments on eggs of seven South American species of Lutzomyia sand flies (Diptera: Psychodidae)”. *Journal of Medical Entomology* 41.5 (2004): 819–825.

De Coninck, E. and R. Coessens. “Life cycle and reproductive pattern of Acrotrichis intermedia (Coleoptera: Ptiliidae) in experimental conditions”. *Journal of Natural History* 15.6 (1981): 1047–1055.

De Fátima Ribeiro, M. and P. de Souza Santos Filho. “Size variation in eggs laid by normal-sized and miniature queens of Plebeia remota (Holmberg) (Hymenoptera: Apidae: Meliponini)”. *Sociobiology* 61.4 (2014): 483–489.

De Figueroa, J. M. T. and J. A. Palomino-Morales. “Eggs and clutches of Sialis nigripes Pictet, 1865 (Megaloptera , Sialidae)”. *Boletín de la Asociación Española de Entomología* 25 (2001): 175–181.

De Figueroa, J. M. T., J. M. Luzón-Ortega, and A. Sánchez-Ortega. “Imaginal biology of the stonefly Hemimelaena flaviventris (Pictet, 1841) (Plecoptera: Perlodidae)”. *Annales Zoologici Fennici* 35.4 (1998): 225–230.

De Figueroa, J. M. T. and T. Derka. “Egg description of Isoptena serricornis (Plecoptera: Chloroperlidae)”. *Entomological Problems* 33.1-2 (2003): 55–57.

De Figueroa, J. M. T. and A. Sánchez-Pérez. “Huevos y puestas de algunas especies de plecópteros (Insecta, Plecoptera) de Sierra Nevada (Granada, España)”. *Zoologica Baetica* 10 (1999): 161–184.

De Loach, C. J. and R. L. Rabb. “Life history of Winthemia manducae (Diptera: Tachinidae), a parasite of the tobacco hornworm”. *Annals of the Entomological Society of America* 64.2 (1971): 399–409.

De Luca, V. and R. Viscuso. “Prime fasi dello sviluppo dell’uovo fecondato di Eyprepocnemis plorans (Charp.) (Orth. acrid.)” *Redia* 1972.3 (1972): 239–249.

De Mello, F., J. Jurberg, and J. Grazia. “Morphological study of the eggs and nymphs of Triatoma dimidiata (Latreille, 1811) observed by light and scanning electron microscopy (Hemiptera: Reduviidae: Triatominae)”. *Memórias do Instituto Oswaldo Cruz* 104.8 (2009): 1072–1082.

De Remes Lenicov, A. M. M., M. E. Brentassi, and A. V. Toledo. “Description of the immature stages of Delphacodes kuscheli Fennah (Hemiptera: Delphacidae), vector of “Mal de Río Cuarto virus” on maize in Argentina”. *Studies on Neotropical Fauna and Environment* 43.1 (2008): 25–33.

De Remes Lenicov, A. M. M., M. C. Hernández, M. E. Brentassi, and B. Defea. “Descriptions of immatures of the South American plant hopper, Taosa (C.) longula”. *Journal of Insect Science* 12.1 (2012): 1–11.

De Sá, V. G. M., J. C. Zanuncio, M. A. Soares, C. S. Rosa, and J. E. Serrao. “Morphology and postdepositional dynamics of eggs of the predator Podisus distinctus (Stål) (Heteroptera: Pentatomidae: Asopinae)”. *Zootaxa* 3641.3 (2013): 282–288.

De Saint Phalle, B. and W. Sullivan. “Incomplete sister chromatid separation is the mechanism of programmed chromosome elimination during early Sciara coprophila embryogenesis”. *Development* 122.12 (1996): 3775–3784.

De Van Kamp, T. and F. H. Hennemann. “A tiny new species of leaf insect (Phasmatodea, Phylliidae) from New Guinea”. *Zootaxa* 3869.4 (2014): 397–408.

Deep, D. S. and H. S. Rose. “Study on the external morphology of the eggs of maize borer, chilo partellus (swinhoe)”. *Journal of Entomology and Zoology Studies* 4.2 (2014): 187–189.

Deepak, D. B., V. C. Minatai, R. H. Izhar, M. M. Hemraj, and P. T. Rani. “The egg of Uzi fly, Exorista sorbillans (? E. Bombycis Louis) (Diptera: Tachinidae)”. *International Journal of Zoology and Research* 5.4 (2015): 19–26.

Degrange, C. “Recherches sur la reproduction des Ephéméroptères”. *Travaux du Laboratoire de Pisciculture de l’Université de Grenoble* 51 (1960): 7–193.

Dell’Erba, R., L. A. Kaminski, and G. R. P. Moreira. “The egg stage of Heliconiini (Lepidoptera, Nymphalidae) from Rio Grande do Sul, Brazil”. *Iheringia, Série Zoologia* 95.1 (2005): 29–46.

Dellapé, P. M. “Redescription of Paromius procerulus (Berg) (new combination) (Heteroptera: Rhyparochromidae: Myodochini), and description of eggs and immature stages”. *Zootaxa* 1070 (2005): 49–60.

Dennis, D. S. and R. J. Lavigne. “Ethology of Efferia varipes with comments on species coexistence (Diptera: Asilidae)”. *Journal of the Kansas Entomological Society* 49.1 (1976): 48–62.

Dennis, D. S. and R. J. Lavigne. “Ethology of machzmus callidus with incidental observations on M. occzdentalis in Wyoming (Diptera: Asilidae)’s2”. *The Pan-Pacific Entomologist* 55.3 (1979): 208–221.

Dennis, D. S., R. J. Lavigne, and S. W. Bullington. “Ethology of Efferia cressoni with a review of the comparative ethology of the genus (Diptera: Asilidae)”. *Proceedings of the Entomological Society of Washington* 88.1 (1986): 42–55.

Deobhakta, S. R. “Preliminary notes on the early embryonic development of Mylabris pustulata Thunb.(Coleoptera)”. *Agra University Journal of Research* 2 (1953): 125–134.

Deutsch, W. G. “Oviposition of Hydropsychidae (Trichoptera) in a large river”. *Canadian Journal of Zoology* 62.10 (1984): 1988–1994.

Dhiman, S. C. and S. C. Goel. “Diaeretiella rapae (M’Intosh) (Hymenoptera: Aphidiidae) a potential biocontrol agent of mustard aphid Lipaphis erysimi (Kalt.)” *Advances in Indian Entomology: Productivity and Health (a Silver Jubilee, Supplement No. 3, Volume II; Insect and Environment)*. Muzaffarnagar: Uttar Pradesh Zoological Society, 2006. 101–109.

Di Giulio, A., M. A. Bologna, and J. D. Pinto. “Larval morphology of the Meloe subgenus Mesomeloe: inferences on its phylogenetic position and a first instar larval key to the Meloe subgenera (Coleoptera, Meloidae)”. *Italian Journal of Zoology* 69.4 (2002): 339–344.

Dias, F. M. S., E. Carneiro, M. M. Casagrande, and O. H. H. Mielke. “Biology and external morphology of immature stages of the butterfly, Diaethria candrena candrena”. *Journal of Insect Science* 12.9 (2012): 1–11.

Dias, F. M. S., M. M. Casagrande, and O. H. H. Mielke. “External morphology and ultra-structure of eggs and first instar of Prepona laertes laertes (Hübner, [1811]), with notes on host plant use and taxonomy”. *Journal of Insect Science* 11.100 (2011): 1–10.

Dias, M. "Immature stages of Citheronia (Citheronula) armata armata Rothschild, 1907 (Lepidoptera, Attacidae). Estagios imaturos de Citheronia (Citheronula) armata armata Rothschild, 1907 (Lepidoptera, Attacidae)". *Revista Brasileira de Entomologia* 25.4 (1981): 295–300.

Dietemann, V. and C. Peeters. "Queen influence on the shift from trophic to reproductive eggs laid by workers of the ponerine ant Pachycondyla apicalis". *Insectes Sociaux* 47.3 (2000): 223–228.

Dimaté, F. A. R., J. C. M. Poderoso, J. E. Serrão, S. Candan, and J. C. Zanuncio. "Comparative Morphology of Eggs of the Predators Brontocoris tabidus and Supputius cincticeps (Heteroptera: Pentatomidae)". *Annals of the Entomological Society of America* 107.6 (2014): 1126–1129.

Diniz, I. R. "Life history and immature stages of Chlamydastis platyspora (Elachistidae)". *Journal of the Lephlojiterists' Sooìetif* 58.2 (2004): 75–79.

Dolinskaya, I. V. "Comparative morphology on the egg chorion characters of some Noctuidae (Lepidoptera)". *Zootaxa* 4085.3 (2016): 374–392.

Dolinskaya, I. V. "Egg morphology of some Noctuidae (Lepidoptera)". *Vestnik Zoologii* 48.4 (2014): 353–364.

Dolinskaya, I. V. "Egg morphology of some Nolidae and Erebidae (Lepidoptera, Noctuoidea)". *Vestnik Zoologii* 48.6 (2014): 553–561.

Dolinskaya, I. V. "Key to the species of Ukrainian Notodontid moths (Lepidoptera, Notodontidae) on the egg characters". *Vestnik Zoologii* 50.6 (2016): 517–532.

Dolinskaya, I. V. "The chorionic sculpture in eggs of some Hadeninae (Lepidoptera, Noctuidae) from Ukraine". *Ukrainska Entomofaunistyka* 1.3 (2010): 2–32.

Dolinskaya, I. V. "The chorionic sculpture of the eggs of some Xyleninae (Lepidoptera, Noctuidae)". *Vestnik Zoologii* 45.1 (2011): 41–56.

Dolinskaya, I. V. and Y. Geryak. "The chorionic sculpture of the eggs of some Noctuinae (Lepidoptera, Noctuidae) from Ukraine". *Vestnik Zoologii* 44.5 (2010): 421–432.

Dolinskaya, I. V. and I. G. Pljushch. "Surface structure of the eggshell of some Vapourer moths (Lepidoptera, Lymantriidae)". *Lambillionea* 99.4 (1999): 489–502.

Dolinskaya, I. V., I. G. Pljushch, and I. G. Pljushch. "External morphology of the eggs of some lappet-moths (Lepidoptera, Lasiocampidae)". *Vestnik Zoologii* 34.3 (2000): 49–60.

Dolinskaya, I. V. and M. G. Ponomarenko. "The chorionic sculpture in eggs of some Noctuidae (Lepidoptera)". *Vestnik Zoologii* 47.5 (2013): 33–41.

Dolling, W. R. "A revision of the neotropical genus Vilga Stål (Hemiptera: Coreidae)". *Systematic Entomology* 2.1 (1977): 27–44.

Domínguez, E. and M. Gabriela Cuezzo. "Ephemeroptera egg chorion characters: a test of their importance in assessing phylogenetic relationships". *Journal of Morphology* 253.2 (2002): 148–165.

Donoughe, S. and C. G. Extavour. "Embryonic development of the cricket Gryllus bimaculatus". *Developmental Biology* 411.1 (2016): 140–156.

Döring, E. *Zur morphologie der schmetterlingseier*. Berlin: Akademie-Verlag, 1955.

Dos Santos, C. M., J. Jurberg, C. Galvão, J. A. da Rosa, W. Cjúnior, J. Barata, and M. T. Obara. "Comparative descriptions of eggs from three species of Rhodnius (Hemiptera: Reduviidae: Triatominae)". *Memórias do Instituto Oswaldo Cruz* 104.7 (2009): 1012–1018.

Al-Dosary, M. M., A. M. Al-Bekairi, and E. B. Moursy. "Morphology of the egg shell and the developing embryo of the Red Palm Weevil, Rhynchophorus ferrugineus (Oliver)". *Saudi Journal of Biological Sciences* 17.2 (2010): 177–183.

Dosdall, L. M. and M. A. McFarlane. "Morphology of the pre-imaginal life stages of the cabbage seedpod weevil, Ceutorhynchus obstrictus (Marsham) (Coleoptera: Curculionidae)". *The Coleopterists Bulletin* 58.1 (2004): 45–52.

Dossi, F. C. A., H. Conte, and A. A. Zacaro. "Histochemical characterization of the embryonic stages in Diatraea saccharalis (Lepidoptera: Crambidae)". *Annals of the Entomological Society of America* 99.6 (2006): 1206–1212.

Downey, J. C. and A. C. Allyn. “Chorionic sculpturing in eggs of Lycaenidae. Part I”. *Bulletin of the Allyn Museum* 61 (1981): 1–29.

Downey, J. C. and A. C. Allyn. “Chorionic sculpturing in eggs of Lycaenidae. Part II”. *Bulletin of the Allyn Museum* 84 (1984): 1–44.

Downey, J. C. and A. C. Allyn. “Eggs of Riodinidae”. *Journal of the Lepidopterists' Society* 34.2 (1980): 133–145.

Du Bois, A. M. “La détermination de l'ébauche embryonnaire chez Sialis lutaria L. (Megaloptera)”. *Revue Suisse de Zoologie* 45.1 (1938): 1–90.

Du, X., C. Yue, and B. Hua. “Embryonic development of the scorpionfly Panorpa emarginata Cheng with special reference to external morphology (Mecoptera: Panorpidae)”. *Journal of Morphology* 270.8 (2009): 984–995.

Duckett, C. N. and Z. Swigoňová. “Description of immature stages of Alagoasa januaria Bechyné (Coleoptera: Chrysomelidae)”. *Journal of the New York Entomological Society* 110.1 (2002): 115–126.

Duffy, E. A. J. *A monograph of the immature stages of African timber beetles (Cerambycidae)*. London: The British Museum (Natural History), 1957.

Duffy, E. A. J. *A monograph of the immature stages of Australasian timber beetles (Gerambycidae)*. London: The British Museum (Natural History), 1963.

Duffy, E. A. J. *A monograph of the immature stages of British and imported timber beetles (Cerambycidae)*. London: The British Museum (Natural History), 1953.

Duffy, E. A. J. *A monograph of the immature stages of Neotropical timber beetles (Cerambycidae)*. London: The British Museum (Natural History), 1960.

Dunlap-Pianka, H. L. “Ovarian dynamics in Heliconius butterflies: correlations among daily oviposition rates, egg weights, and quantitative aspects of oögenesis”. *Journal of Insect Physiology* 25.9 (1979): 741–749.

DuPraw, E. J. “The honeybee embryo”. *Methods in Developmental Biology*. New York: Thomas Y Cromwell, 1967. 183–217.

Dutra, V. S., B. Ronchi-Teles, G. J. Steck, and J. G. Silva. “Egg morphology of Anastrepha spp.(Diptera: Tephritidae) in the fraterculus group using scanning electron microscopy”. *Annals of the Entomological Society of America* 104.1 (2011): 16–24.

Dutrillaux, A. M., D. Pluot-Sigwalt, and B. Dutrillaux. “(Ovo-) viviparity in the darkling beetle, Alegoria castelnaui (Tenebrioninae: Ulomini), from Guadeloupe”. *European Journal of Entomology* 107.4 (2010): 481–485.

Dyar, H. G. “The life histories of the New York slug-caterpillars–XX”. *Journal of the New York Entomological Society* 22.3 (1914): 223–229.

Dybas, H. S. “Polymorphism in featherwing beetles, with a revision of the genus Ptinellodes (Coleoptera: Ptiliidae)”. *Annals of the Entomological Society of America* 71.5 (1978): 695–714.

Eayment, T. “Biology of a new Halictine bee”. *Arbeiten über Physiologische und angewandte Entomologie aus Berlin-Dahlem* 4.1 (1937): 30–60.

Eben, A. and M. E. Barbercheck. “Sculpturing of the eggshell of some Mexican Galerucinae (Coleoptera: Chrysomelidae)”. *The Coleopterists' Bulletin* 51.1 (1997): 80–85.

Edgerly, J. S., C. A. Szumik, and C. N. McCreedy. “On new characters of the eggs of Embioptera with the description of a new species of Saussurembia (Anisembiidae)”. *Systematic Entomology* 32.2 (2007): 387–395.

Edwards, J. G. “Observations on the Biology of Amphizoidae”. *The Coleopterists Bulletin* 8.1 (1954): 19–24.

Edwards, M. G. “Digestive enzymes of vine weevil (otiorhynchus sulcatus) as potential targets for insect control strategies”. Diss. University of Durham, 2002.

Egwuatu, R. I. and T. Ajibola Taylor. “Studies on the biology of Acanthomia tomentosicollis (Stål) (Hemiptera: Coreidae) in the field and insectary”. *Bulletin of Entomological Research* 67.2 (1977): 249–257.

Eickwort, G. C. “Aspects of the nesting biology and descriptions of immature stages of Perdita octomaculata and P. halictoides (Hymenoptera: Andrenidae)”. *Journal of the Kansas Entomological Society* 50.4 (1977): 577–599.

Ellertson, F. E. and P. O. Ritcher. “Biology of rain beetles, Pleocoma spp, associated with fruit trees in Wasco and Hood River counties”. *Oregon Agricultural Experiment Station Bullletin* 44 (1959): 1–42.

Elliott, K. L. and B. Stay. “Juvenile hormone synthesis as related to egg development in neotenic reproductives of the termite Reticulitermes flavipes, with observations on urates in the fat body”. *General and Comparative Endocrinology* 152.1 (2007): 102–110.

Ellis, J. D. and C. M. Z. Nalen. “Bee louse, bee fly, Braulid, Braula coeca Nitzsch (Insecta: Diptera: Braulidae)”. *Department of Entomology and Nematology, UF/IFAS Extension. Original Publication* EENY472 (2010): 1–3.

Embree, D. G. “The External Morphology of the Immature Stages of the Beech Leaf Tier, Psilocorsis faginella (Chamb) (Lepidoptera: Oecophoridae), with Notes on Its Biology in Nova Scotia”. *The Canadian Entomologist* 90.3 (1958): 166–174.

Endris, R. G., D. G. Young, and P. V. Perkins. “Ultrastructural comparison of egg surface morphology of five Lutzomyia species (Diptera: Psychodidae)”. *Journal of Medical Entomology* 24.4 (1987): 412–415.

Eppley, R. K. “Studies on Heterostylum robustum (Osten Sacken) (Diptera: Bombyliidae), a parasite of Nomia melanderi”. MA thesis. Oregon State University, 1963.

Epstein, M. E. *Revision and phylogeny of the limacodid-group families, with evolutionary studies on slug caterpillars (Lepidoptera: Zygaenoidea)*. Washington, D. C.: Smithsonian Institution Press, 1996.

Erzinçlioğlu, Y. Z. “Immature stages of British Calliphora and Cynomya, with a re-evaluation of the taxonomic characters of larval Calliphoridae (Diptera)”. *Journal of Natural History* 19.1 (1985): 69–96.

Erzinçlioğlu, Y. Z. “Studies on the morphology and taxonomy of the immature stages of Calliphoridae, with analysis of phylogenetic relationships within the family, and between it and other groups in the Cyclorrhapha (Diptera)”. Diss. University of Durham, 1984.

Erzinçlioğlu, Y. Z. “The value of chorionic structure and size in the diagnosis of blowfly eggs”. *Medical and Veterinary Entomology* 3.3 (1989): 281–285.

Eskafi, F. M. and E. F. Legner. “Descriptions of immature stages of the cynipid Hexacola sp. near websteri (Eucoilinae: Hymenoptera), a larval–pupal parasite of hippelates eye gnats (Diptera: Chloropidae)”. *The Canadian Entomologist* 106.10 (1974): 1043–1048.

Esmaili, M. “Four species of leaf miners attacking deciduous fruit trees in Iran’s central province”. *Zeitschrift für Angewandte Entomologie* 69.1-4 (1971): 407–415.

Espadaler, X. and S. Rey. “Biological constraints and colony founding in the polygynous invasive ant Lasius neglectus (Hymenoptera, Formicidae)”. *Insectes Sociaux* 48.2 (2001): 159–164.

Esselbaugh, C. O. “A study of the eggs of the Pentatomidae (Hemiptera)”. *Annals of the Entomological Society of America* 39.4 (1946): 667–691.

Evans, D. “The life history and immature stages of Synergus pacificus McCracken and Egbert (Hymenoptera: Cynipidae)”. *The Canadian Entomologist* 97.2 (1965): 185–188.

Evans, H. E. “A solitary wasp that preys upon lacewings (Hymenoptera: Sphecidae; Neuroptera: Chrysopidae)”. *Psyche* 85.1 (1978): 81–84.

Evans, H. E. “Observations on the nesting behavior of three species of the genus Crabro (Hymenoptera, Sphecidae)”. *Journal of the New York Entomological Society* 68.3 (1960): 123–134.

Evans, H. E., F. E. Kurczewski, and J. Alcock. “Observations on the nesting behaviour of seven species of Crabro (Hymenoptera, Sphecidae)”. *Journal of Natural History* 14.6 (1980): 865–882.

Evans, H. E. and R. W. Matthews. “Observations on the nesting behavior of Trachypus petiolatus (Spinola) in Colombia and Argentina (Hymenoptera: Sphecidae: Philanthini)”. *Journal of the Kansas Entomological Society* 46.2 (1973): 165–175.

Evans, H. E., R. W. Matthews, and W. Pulawski. “Notes on the nests and prey of four Australian species of Tachysphex Kohl, with description of a new species (Hymenoptera: Sphecidae)”. *Australian Journal of Entomology* 15.4 (1977): 441–445.

Evgeny, A. B. “Discover of the American Green-striped Forest Looper, Melanolophia imitata (Walker) Lepidoptera; Geometridae) in Korea”. *Korean Journal of Applied Entomology* 40.1 (2001): 1–4.

Eyles, A. C. “Key to the genera of Mirinae (Hemiptera: Miridae) in New Zealand and descriptions of new taxa”. *New Zealand Journal of Zoology* 28.2 (2001): 197–221.

Eyles, A. C. “New genera and species of the Lygus-Complex (Hemiptera: Miridae) in the New Zealand subregion compared with subgenera (now genera) studied by Leston (1952) and Niastama Reuter”. *New Zealand Journal of Zoology* 26.4 (1999): 303–354.

Eyles, A. C. “Revision of New Zealand Orthotylinae (Insecta: Hemiptera: Miridae)”. *New Zealand Journal of Zoology* 32.3 (2005): 181–215.

Eyles, A. C. “Variation in the adult and immature stages of Nysius huttoni White (Heteroptera: Lygaeidae) with a note on the validity of the genus Brachynysius Usinger”. *Transactions of the Royal Entomological Society of London* 112.4 (1960): 53–72.

Eyles, A. C. and R. T. Schuh. “Revision of New Zealand Bryocorinae and Phylinae (Insecta: Hemiptera: Miridae)”. *New Zealand Journal of Zoology* 30.3 (2003): 263–325.

Ezquiaga, M. C. and M. Lareschi. “Surface ultrastructure of the eggs of Malacopsylla grossiventris and Phthiropsylla agenoris (Siphonaptera: Malacopsyllidae)”. *Journal of Parasitology* 98.5 (2012): 1029–1031.

Falamarzi, S., A. Puetz, M. Heidari, and H. Nasserzadeh. “Confirmed occurrence of Hydroscapha granulum in Iran, with notes on its biology (Coleoptera: Myxophaga: Hydroscaphidae)”. *Acta Entomologica Musei Nationalis Pragae* 50.1 (2010): 97–106.

Fasoranti, J. O. “The life history and habits of a Ceanothus leaf miner, Tischeria immaculata (Lepidoptera: Tischeriidae)”. *The Canadian Entomologist* 116.11 (1984): 1441–1448.

Fateryga, A. V. and A. V. Amolin. “Nesting and biology of Jucancistrocerus caspicus (Hymenoptera, Vespidae, Eumeninae)”. *Entomological Review* 94.1 (2014): 73–78.

Faust, L. F. “Natural history and flash repertoire of the synchronous firefly Photinus carolinus (Coleoptera: Lampyridae) in the Great Smoky Mountains National Park”. *The Florida Entomologist* 93.2 (2010): 208–217.

Fausto, A. M., M. D. Feliciangeli, M. Maroli, and M. Mazzini. “Ootaxonomic investigation of five Lutzomyia species (Diptera, Psychodidae) from Venezuela”. *Memórias do Instituto Oswaldo Cruz* 96.2 (2001): 197–204.

Fausto, A. M., M. Maroli, and M. Mazzini. “Ootaxonomy and eggshell ultrastructure of Phlebotomus sandflies”. *Medical and Veterinary Entomology* 6.3 (1992): 201–208.

Fausto, A. M., M. Mazzini, M. Maroli, and M. J. Mutinga. “Scanning electron microscopical study of the eggshell of three species of Sergentomyia (Diptera: Psychodidae)”. *International Journal of Tropical Insect Science* 14.4 (1993): 483–488.

Fehrenbach, H. “Fine structure of the eggshells of four primitive moths: Hepialus hecta (L.), Wiseana umbraculata (Guénée) (Hepialidae), Mnesarchaea fusilella Walker and M. acuta Philp. (Mnesarchaeidae) (Lepidoptera, Exoporia)”. *International Journal of Insect Morphology and Embryology* 18.5 (1989): 261–274.

Fehrenbach, H., V. Dittrich, and D. Zissler. “Eggshell fine structure of three lepidopteran pests: Cydia pomonella (L.) (Tortricidae), Heliothis virescens (Fabr.), and Spodoptera littoralis (Boisd.) (Noctuidae)”. *International Journal of Insect Morphology and Embryology* 16.3-4 (1987): 201–219.

Feliciangeli, M. D., O. C. Castejon, and J. Limongi. “Egg surface ultrastructure of eight New World phlebotomine sand fly species (Diptera: Psychodidae)”. *Journal of Medical Entomology* 30.4 (1993): 651–656.

Ferguson, D. C. “A new genus of winter moths (Geometridae) from eastern California and western Nevada”. *The Journal of the Lepidopterists Society* 48.1 (1994): 8–23.

Fernandes, J. A. M. and J. Grazia. “Estudo dos estágios imaturos de Leptoglossus zonatus (Dallas, 1852) (Heteroptera-Coreidae)”. *Anais da Sociedade Entomológica do Brasil* 21.2 (1992): 180–188.

Fernandez, L. A. and R. E. Campos. “Description of immature stages of Berosus alternans Brullé (Coleoptera: Hydrophilidae)”. *Transactions of the American Entomological Society* 128.2/3 (2002): 255–263.

Fernando, W. “The early embryology of a viviparous psocid”. *Quarterly Journal of Microscopical Science* 77.305 (1934): 99–119.

Ferrar, P. *A guide to the breeding habits and immature stages of Diptera Cyclorrhapha*. Copenhagen: Scandinavian Science Press, 1987.

Ferrar, P. “The immature stages of dung-breeding Muscoid flies in Australia, with notes on the species, and keys to larvae and puparia”. *Australian Journal of Zoology* 27.73 (1979): 1–106.

Ferro, C., E. Cárdenas, D. Corredor, A. Morales, and L. E. Munstermann. "Life cycle and fecundity analysis of Lutzomyia shannoni (Dyar) (Diptera: Psychodidae)". *Memórias do Instituto Oswaldo Cruz* 93.2 (1998): 195–199.

Fischer, M., J. Tormos, X. Pardo, and J. D. Asís. "Description of Adults, Preimaginal Phases, and the Venom Apparatus of a New Species of Aspilota Förster from Spain, with Comments on and Discussion of Immature Stages of the Alysiinae (Hymenoptera: Braconidae)". *Zoological Studies* 47.3 (2007): 247–257.

Fish, W. A. "Embryology of Lucilia sericata Meigen (Diptera: Calliphoridae) Part I. Cell cleavage and early embryonic development". *Annals of the Entomological Society of America* 40.1 (1947): 15–28.

Fisher, R. M. and B. J. Sampson. "Morphological specializations of the bumble bee social parasite Psithyrus ashtoni (Cresson) (Hymenoptera: Apidae)". *The Canadian Entomologist* 124.1 (1992): 69–77.

Fitzpatrick, S. M. and J. T. Troubridge. "Fecundity, number of diapause eggs, and egg size of successive generations of the blackheaded fireworm (Lepidoptera: Tortricidae) on cranberries". *Environmental Entomology* 22.4 (1993): 818–823.

Fletcher, M. J. "Egg types and oviposition behaviour in some fulgoroid leafhoppers (Homoptera, Fulgoroidea)". *Australian Entomological Magazine* 6.1 (1979): 13–18.

Flosi, J. W. "The population biology of the giant water bug Belostoma flumineum Say (Hemiptera: Belostomatidae)". Diss. Iowa State University, 1980.

Flowers, R. W., W. D. Shepard, and R. Mera. "A new species of Lepicerus (Coleoptera: Lepiceridae) from Ecuador". *Zootaxa* 2639 (2010): 35–39.

Földvári, M. "Taxonomic and faunistic studies of big-headed flies (Diptera: Pipunculidae)". Diss. University of Szeged, 2004.

Fombong, A. T., F. Haas, P. N. Ndegwa, and L. W. Irungu. "Life history of Oplostomus haroldi (Coleoptera: Scarabaeidae) under laboratory conditions and a description of its third instar larva". *International Journal of Tropical Insect Science* 32.1 (2012): 56–63.

Foote, B. A. "Biology and immature stages of Coenia curvicauda (Diptera: Ephydridae)". *Journal of the New York Entomological Society* 98.1 (1990): 93–102.

Foote, B. A., S. E. Neff, and C. O. Berg. "Biology and immature stages of Atrichomelina pubera (Diptera: Sciomyzidae)". *Annals of the Entomological Society of America* 53.2 (1960): 192–199.

Forister, M. L., J. A. Fordyce, C. C. Nice, Z. Gompert, and A. M. Shapiro. "Egg morphology varies among populations and habitats along a suture zone in the Lycaeides idas-melissa species complex (Lepidoptera: Lycaenidae)". *Annals of the Entomological Society of America* 99.5 (2006): 933–937.

Forrester, J. A., N. J. Vandenberg, and J. V. Mchugh. "Redescription of Anovia circumclusa (Gorham) (Coleoptera: Coccinellidae: Noviini), with first description of the egg, larva, and pupa, and notes on adult intraspecific elytral pattern variation". *Zootaxa* 2112 (2009): 25–40.

Forrester, J. A. "Sacred systematics: the Noviini of the world (Coleoptera: Coccinellidae)". Diss. University of Georgia, 2008.

Forteath, G. N. R. and A. W. Osborn. "Biology, ecology and voltinism of the Australian spongillafly Sisyra pedderensis Smithers (Neuroptera: Sisyridae)". *Papers and Proceedings of the Royal Society of Tasmania* 146 (2012): 25–36.

Fox, C. W. "The influence of egg size on offspring performance in the seed beetle, Callosobruchus maculatus". *Oikos* 71 (1994): 321–325.

Fox, C. W., M. S. Thakar, and T. A. Mousseau. "Egg size plasticity in a seed beetle: an adaptive maternal effect". *American Naturalist* 149.1 (1997): 149–163.

Fox, C., K. I. M. Waddell, J. Des Lauries, and T. Mousseau. "Seed beetle survivorship, growth and egg size plasticity in a paloverde hybrid zone". *Ecological Entomology* 22.4 (1997): 416–424.

Fox, E. G. P., D. R. Solis, C. M. de Jesus, O. C. Bueno, A. T. Yabuki, and M. L. Rossi. "On the immature stages of the crazy ant Paratrechina longicornis (Latreille 1802) (Hymenoptera: Formicidae)". *Zootaxa* 1503 (2007): 1–11.

Fox, E. G. P., D. R. Solis, M. L. Rossi, J. H. C. Delabie, R. F. De Souza, and O. C. Bueno. "Comparative immature morphology of Brazilian fire ants (Hymenoptera: Formicidae: Solenopsis)". *Psyche* 2012 (2011): 1–10.

Fox, E. G. P., D. R. Solis, M. L. Rossi, R. Eizemberg, L. P. Taveira, and S. Bressan-Nascimento. "The preimaginal stages of the ensign wasp Evania appendigaster (Hymenoptera, Evaniidae), a cockroach egg predator". *Invertebrate Biology* 131.2 (2012): 133–143.
Frank, J. H. and H. Nadel. "Life cycle and behaviour of Charoxus spinifer and Charoxus major (Coleoptera: Staphylinidae: Aleocharinae), predators of fig wasps (Hymenoptera: Agaonidae)". *Journal of Natural History* 46.9-10 (2012): 621–635.
Fraulob, M., R. G. Beutel, R. Machida, and H. Pohl. "The embryonic development of Stylops ovinae (Strepsiptera, Stylopidae) with emphasis on external morphology". *Arthropod Structure & Development* 44.1 (2015): 42–68.
Freitas, A. V. L. and K. S. Brown Jr. "Immature stages of Vila emilia". *Tropical Lepidoptera Research* 18 (2008): 74–77.
Freitas, A. V. L., K. S. Brown Jr, and L. D. Otero. "Juvenile stages of Cybdelis, a key genus uniting the diverse". *Tropical Lepidoptera* 8.1 (1997): 29–34.
Frías, D. "Morphology of immature stages in the neotropical nonfrugivorous Tephritinae fruit fly species Rachiptera limbata Bigot (Diptera: Tephritidae) on Baccharis linearis (R. et Pav.) (Asteraceae)". *Neotropical Entomology* 37.5 (2008): 536–545.
Frost, S. W. "Hosts and eggs of Blepharida dorothea (Coleoptera: Chrysomelidae)". *The Florida Entomologist* 56.2 (1973): 120–122.
Fujita, M. and R. Machida. "Embryonic development of Eucorydia yasumatsui Asahina, with special reference to external morphology (Insecta: Blattodea, Corydiidae)". *Journal of Morphology* 278.11 (2017): 1469–1489.
Fujiwara, N. and H. Kobayashi. "Embryogenesis of the Leather Winged Beetle, Athemus suturellus Motschulsky (Coleoptera, Cantharidae)". *Recent Advances in Insect Embryology in Japan and Poland*. Tsukuba: The Arthropodan Embryological Society of Japan, 1987. 195–206.
Fujiwara, Y., T. Takahashi, T. Yoshioka, and F. Nakasuji. "Changes in egg size of the diamondback moth Plutella xylostella (Lepidoptera: Yponomeutidae) treated with fenvalerate at sublethal doses and viability of the eggs". *Applied Entomology and Zoology* 37.1 (2002): 103–109.
Furlan, L. "The biology of Agriotes sordidus Illiger (Col., Elateridae)". *Journal of Applied Entomology* 128.9-10 (2004): 696–706.
Furlan, L. "The biology of Agriotes ustulatus Schäller (Col., Elateridae). I. Adults and oviposition". *Journal of Applied Entomology* 120.1-5 (1996): 269–274.
Furneaux, P. J. S., C. R. James, and S. A. Potter. "The egg shell of the house cricket (Acheta domesticus): an electron-microscope study". *Journal of Cell Science* 5.1 (1969): 227–249.
Furniss, M. M. and S. J. Kegley. "Observations on the biology of Dryocoetes betulae (Coleoptera: Curculionidae) in paper birch in northern Idaho". *Environmental Entomology* 35.4 (2006): 907–911.
Furukawa, E. and K. Kaneko. "Studies on phorid flies (Phoridae, Diptera) in Japan. IV. Scanning electron microscopic observations of eggs of two Megaselia". *Japanese Journal of Sanitary Zoology* 31.1 (1981): 78–81.
Fuseini, B. A. and R. Kumar. "Biology and immature stages of cotton stainers (Heteroptera: Pyrrhocoridae) found in Ghana". *Biological Journal of the Linnean Society* 7.2 (1975): 83–111.
Gaino, E. and M. Mazzini. "Scanning electron microscope study of the eggs of some Habrophlebia and Habroleptoides species (Ephemeroptera, Leptophlebiidae)". *Proceedings of the Fourth International Conference on Ephemeroptera*. Ceské Budějovice: Institute of Entomology, Czechoslovak Academy of Sciences, 1984.
Gaino, E. and E. Bongiovanni. "Scanning electron microscopy of the eggs of Palingenia longicauda (Olivier) (Ephemeroptera: Palingeniidae)". *International Journal of Insect Morphology and Embryology* 22.1 (1993): 41–48.
Gaino, E. and J. Flannagan. "Fine external morphology of the eggs of Ephoron album (Say) and Ephoron shigae (Takahashi) (Ephemeroptera, Polymitarcyidae)". *The Canadian Entomologist* 127.4 (1995): 527–533.
Gaino, E. and M. Mazzini. "Fine structure of the chorionic projections of the egg of Rhithrogena kimminsi Thomas (Ephemeroptera: Heptageniidae) and their role in egg adhesion". *International Journal of Insect Morphology and Embryology* 17.2 (1988): 113–120.

Gaino, E., M. Mazzini, and M. Sartori. "Comparative analysis of the chorionic pattern in Habroleptoides species (Ephemeroptera, Leptophlebiidae)". *Italian Journal of Zoology* 60.2 (1993): 155–162.

Gaino, E. and M. Rebora. "Synthesis and function of the fibrous layers covering the eggs of Siphlonurus lacustris (Ephemeroptera, Siphlonuridae)". *Acta Zoologica* 82.1 (2001): 41–48.

Gaino, E., M. Sartori, and M. Rebora. "Chorionic fine structure of eggs from some species of Proboscidoplocia (Ephemeroptera, Ephemeroidea)". *Italian Journal of Zoology* 68.1 (2001): 1–8.

Gallard, L. "Notes on the life-history of the large yellow lacewing, Nymphes myrmeleonides". *Australian Naturalist* 9 (1935): 118–119.

Gallego, K. R., J. M. Lerma, C. G. Echeverri, and J. W. Brown. "Description of the early stages of Eccopsis galapagana Razowski & Landry (Tortricidae), a defoliator of Prosopis juliflora (SW.) DC. (Fabaceae) in Colombia". *The Journal of the Lepidopterists' Society* 66.3 (2012): 156–164.

Galvão, C., F. M. McAloon, D. S. Rocha, C. W. Schaefer, J. Patterson, and J. Jurberg. "Description of eggs and nymphs of Linshcosteus karupus (Hemiptera: Reduviidae: Triatominae)". *Annals of the Entomological Society of America* 98.6 (2005): 861–872.

Gambrell, F. L. and L. A. Jahn. "The embryology of the black fly, Simulium pictipes Hagen". *Annals of the Entomological Society of America* 26.4 (1933): 641–671.

Gangrade, G. A. "The biology and morphology of immature stages of Euderus agromyzae Gangrade (Eulophidae: Hymenoptera)". *Indian Journal of Entomology* 24 (1962): 265–273.

Ganguly, A., C. Malakar, H. Anand, S. Das, A. Das, and P. Haldar. "Scanning electron microscopy of egg-surface sculpturing of two common Indian short-horn grasshoppers (Orthoptera, Acrididae)". *Journal of Orthoptera Research* 17.1 (2008): 97–100.

Ganho, N. G. and R. C. Marinoni. "Algumas características da reprodução e ontogênese de Epilachna paenulata (Germar)(Coleoptera, Coccinellidae, Epilachninae)". *Revista Brasileira de Zoologia* 17.2 (2000): 445–454.

Garbiec, A., J. Kubrakiewicz, M. Mazurkiewicz-Kania, B. Simiczyjew, and I. Jedrzejowska. "Asymmetry in structure of the eggshell in Osmylus fulvicephalus (Neuroptera: Osmylidae): an exceptional case of breaking symmetry during neuropteran oogenesis". *Protoplasma* 253.4 (2015): 1033–1042.

García-Barros, E. "Egg size in butterflies (Lepidoptera: Papilionoidea and Hesperiidae): a summary of data". *Journal of Research on the Lepidoptera* 35 (2000): 90–136.

García-Barros, E. and J. Martin. "The eggs of European satyrine butterflies (Nymphalidae): external morphology and its use in systematics". *Zoological Journal of the Linnean Society* 115.1 (1995): 73–115.

Gardiner, P. "The morphology and biology of Ernobius mollis L. (Coleoptera—Anobiidae)". *Transactions of the Royal Entomological Society of London* 104.1 (1953): 1–24.

Garófalo, C. A., E. Camillo, and J. C. Serrano. "Reproductive aspects of Meloetyphlus fuscatus a meloid beetle cleptoparasite of the bee Eulaema nigrita (Hymenoptera, Apidae, Euglossini)". *Apidologie* 42.3 (2011): 337–348.

Garófalo, C. A. and J. G. Rozen Jr. "Parasitic behavior of Exaerete smaragdina with descriptions of its mature oocyte and larval instars (Hymenoptera: Apidae: Euglossini)". *American Museum Novitates* 3349 (2001): 1–28.

Garrevoet, T. and W. Garrevoet. "Bembecia lingenhoelei, a new Clearwing moth from Tajikistan (Lepidoptera: Sesiidae)". *Phegea* 39.2 (2011): 73–79.

Gates, M., J. Mena Correa, J. Sivinski, R. Ramírez-Romero, G. Córdova-García, and M. Aluja. "Description of the immature stages of Eurytoma sivinskii Gates and Grissell (Hymenoptera: Eurytomidae), an ectoparasitoid of Anastrepha (Diptera: Tephritidae) pupae in Mexico". *Entomological News* 119.4 (2008): 354–360.

Gathalkar, G. B., D. D. Barsagade, and A. Sen. "Biology and Development of Xanthopimpla pedator (Hymenoptera: Ichneumonidae): Pupal Endoparasitoid of Antheraea mylitta (Lepidoptera: Saturniidae)". *Annals of the Entomological Society of America* 110.6 (2017): 544–550.

Gautam, S. G., G. P. Opit, D. Margosan, D. Hoffmann, J. S. Tebbets, and S. Walse. "Comparative egg morphology and chorionic ultrastructure of key stored-product insect pests". *Annals of the Entomological Society of America* 108.1 (2015): 43–56.

Gautam, S. G., G. P. Opit, D. Margosan, J. S. Tebbets, and S. Walse. “Egg morphology of key stored-product insect pests of the United States”. *Annals of the Entomological Society of America* 107.1 (2014): 1–10.

Gauvin, M.-J., G. Boivin, and J.-P. Nénon. “Hydropy and ultrastructure of egg envelopes in Aleochara bilineata (Coleoptera, Staphylinidae)”. *Zoomorphology* 120.3 (2001): 171–175.

Geertsema, H. “Micropyle and chorion structure of the eggs of some South African hepialoid moths (Lepidoptera: Prototheoridae: Hepialidae): short communication”. *African Entomology* 12.1 (2004): 147–153.

Genung, W. G., R. E. Woodruff, and E. E. Grissell. “Languria erythrocephalus: Host plants, immature stages, parasites, and habits (Coleoptera: Languriidae)”. *The Florida Entomologist* 63.2 (1980): 206–210.

Gepp, J. “An illustrated review of egg morphology in the families of Neuroptera (Insecta: Neuropteroidea)”. *Advances in Neuropterology: Proceedings of the Third International Symposium on Neuropterology*. Pretoria, R.S.A: Directorate of Agricultural Development, 1990. 131–149.

Gerard, P. J. “Biology and morphology of immature stages of Centrodora scolylbopae (Hymenoptera: Aphelinidae)”. *New Zealand Entomologist* 12.1 (1989): 24–29.

Gerard, P. J. and L. D. Ruf. “Development and biology of the immature stages of Anthrenocerus australis Hope (Coleoptera: Dermestidae)”. *Journal of Stored Products Research* 33.4 (1997): 347–357.

Gerberg, E. J. “A revision of the New World species of powder-post beetles belonging to the family Lyctidae”. *Technical Bulletin of the United States Department of Agriculture* 1157 (1957): 1–55.

Gerwel, C. “Rozwój zarodkowy porphyrophora polonica Ckll. (Coccidae)”. *The Poznan Society of Friends of Science Department of Mathematical and Natural Sciences Biological Section* 12.3 (1950): 1–33.

Ghosh, C. C. “Life history of Helicominus dicus Walk”. *Journal of the Bombay Natural History Society* 22 (1913): 643–648.

Ghosh, K. N. and J. Mukhopadhyay. “A comparison of chorionic sculpturing of four Indian phlebotomine sandflies (Diptera: Psychodidae) by scanning electron microscopy”. *Parasite* 3.1 (1996): 61–67.

Giannotti, E. “Biology of the wasp Polistes (epicnemius) cinerascens Saussure (Hymenoptera: Vespidae)”. *Anais da Sociedade Entomológica do Brasil* 26.1 (1997): 61–67.

Gibbs, G. W. “A new species of tusked weta from the Raukumara Range, North Island, New Zealand (Orthoptera: Anostostomatidae: Motuweta)”. *New Zealand Journal of Zoology* 29.4 (2002): 293–301.

Gibbs, G. W. “Four new species of giant weta, Deinacrida (Orthoptera: Anostostomatidae: Deinacridinae) from New Zealand”. *Journal of the Royal Society of New Zealand* 29.4 (1999): 307–324.

Gibbs, G. W. “Presidential Address: Some notes on the biology and status of the Mnesarchaeidae (Lepidoptera)”. *New Zealand Entomologist* 7.1 (1979): 2–9.

Gilgado, J. D. and V. M. Ortuño. “Biological notes and description of egg and first instar larva of Carabus (Oreocarabus) ghilianii La Ferté-Sénectère 1847 (Coleoptera: Carabidae)”. *Annales de la Société Entomologique de France* 47.3-4 (2011): 444–456.

Giorgi, F. and J. H. Nordin. “Structure of yolk granules in oocytes and eggs of Blattella germanica and their interaction with vitellophages and endosymbiotic bacteria during granule degradation”. *Journal of Insect Physiology* 40.12 (1994): 1077–1092.

Gittelman, S. H. “Descriptions of immature and adult stages of Martarega hondurensis Bare (Hemiptera: Notonectidae)”. *Journal of the Kansas Entomological Society* 47.2 (1974): 145–155.

Gnaneswaran, R. and H. N. P. Wijayagunasekara. “Biology of Cyllodes bifacies walker (Coleoptera: Cucujoidea: Nitidulidae), a pest of oyster mushroom (Pleurotus ostreatus) in Sri Lanka”. *Tropical Agricultural Research* 8 (1996): 377–390.

Gobin, B., C. Peeters, and J. Billen. “Production of trophic eggs by virgin workers in the ponerine antGnamptogenys menadensis”. *Physiological Entomology* 23.4 (1998): 329–336.

Godunko, R. J. and M. Klonowska-Olejnik. “Ecdyonurus ntgrescens (Klapálek, 1908) (Ephemeroptera: Heptagenidae)-Neotype Designation, Taxonomical and Nomenclature Notes”. *Annales Zoologici* 58.4 (2008): 799–817.

González-Acuña, D., C. Briceño, A. Cicchino, S. M. Funk, and J. Jiménez. "First records of Trichodectes canis (Insecta: Phthiraptera: Trichodectidae) from Darwin's fox, Pseudalopex fulvipes (Mammalia: Carnivora: Canidae)". *European Journal of Wildlife Research* 53.1 (2007): 76–79.

Gonzalez, V. H., A. Mejia, and C. Rasmussen. "Ecology and nesting behavior of Bombus atratus Franklin in Andean highlands (Hymenoptera: Apidae)". *Journal of Hymenoptera Research* 13.2 (2004): 28–36.

Gorder, N. K. N. and J. W. Mertins. "Life history of the parsnip webworm, Depressaria pastinacella (Lepidoptera: Oecophoridae), in central Iowa". *Annals of the Entomological Society of America* 77.5 (1984): 568–573.

Gordh, G. and R. E. Medved. "Biological notes on Goniozus pakmanus Gordh (Hymenoptera: Bethylidae), a parasite of pink bollworm, Pectinophora gossypiella (Saunders) (Lepidoptera: Gelechiidae)". *Journal of the Kansas Entomological Society* 59.4 (1986): 723–734.

Gordinier, H. C. "Biology of Diastrophus nebulosus (Hymenoptera: Cynipidae) and its parasitoid/inquiline complex in galls on Rubus flagellaris (Rosaceae)". *Great Lakes Entomologist* 36.3/4 (2003): 129–151.

Görg, I. "Untersuchungen am Keim von Hierodula (Rhombodera) crassa GIGLIO TOS, ein Beitrag zur Embryologie der Mantiden". *Deutsche Entomologische Zeitschrift* 6 (1959): 390–450.

Goss, R. J. "The early embryology of the book louse, Liposcelis divergens Badonnel (Psocoptera; Liposcelidae)". *Journal of Morphology* 91.1 (1952): 135–167.

Gottardo, M. "A new species of Korinnis Günther from the Philippines (Phasmatodea: Prisopodidae: Korinninae)". *Zootaxa* 1917 (2008): 61–64.

Gottardo, M. and P. Heller. "An enigmatic new stick insect from the Philippine Islands (Insecta: Phasmatodea)". *Comptes Rendus Biologies* 335.9 (2012): 594–601.

Gower, A. M. "A study of Limnephilus lunatus Curtis (Trichoptera: Limnephilidae) with reference to its life cycle in watercress beds". *Transactions of the Royal Entomological Society of London* 119.10 (1967): 283–302.

Gradinarov, D., Y. Petrova, E. Tashevaterzieva, and A. V. Frolov. "Biology of the blind geobiont scarab beetle genus Chaetonyx Schaum, 1862 (Scarabaeidae: Orphninae) with new distribution records of Ch. robustus Schaum, 1862 from Bulgaria". *ZooNotes* 1.81 (2015): 1–14.

Gray, B. "The immature stages of Hylurdrectonus araucariae Schedl and H. piniarius Schedl (Coleoptera: Scolytidae: Hylesininae)". *Journal of Entomology Series B, Taxonomy* 42.1 (1973): 49–58.

Greenberg, B. and D. Singh. "Species identification of calliphorid (Diptera) eggs". *Journal of Medical Entomology* 32.1 (1995): 21–26.

Greenberg, B. and J. D. Wells. "Forensic use of Megaselia abdita and M. scalaris (Phoridae: Diptera): case studies, development rates, and egg structure". *Journal of Medical Entomology* 35.3 (1998): 205–209.

Greven, H., M. Mielewczik, and H. Hammer. "Röntgenmikroanalytische Untersuchung des Chorions der Gespenstschrecke Pharnacia westwoodi (Phasmatodea)". *Entomologie Heute* 17 (2005): 39–45.

Grochowska, M. "Remarks on the morphology and biology of Cleigastra apicalis (Meigen, 1826) (Diptera, Scathophagidae)". *Acta Zoologica* 87.4 (2006): 247–252.

Gross, J. B. and R. B. Howland. "The early embryology of Prodenia eridania". *Annals of the Entomological Society of America* 33.1 (1940): 56–76.

Grossniklaus-Bürgin, C., T. Wyler, R. Pfister-Wilhelm, and B. Lanzrein. "Biology and morphology of the parasitoid Chelonus inanitus (Braconidae, Hymenoptera) and effects on the development of its host Spodoptera littoralis (Noctuidae, Lepidoptera)". *Invertebrate Reproduction & Development* 25.2 (1994): 143–158.

Grzywacz, A. and T. Pape. "Egg morphology of Mydaea lateritia (Rondani, 1866) (Diptera: Muscidae)". *Entomologica Fennica* 21.3 (2010): 187–192.

Grzywacz, A., K. Szpila, and T. Pape. "Egg morphology of nine species of pollenia robineau-desvoidy, 1830 (Diptera: Calliphoridae)". *Microscopy Research and Technique* 75.7 (2012): 955–967.

Guglielmino, A. and E. G. Virla. "Postembryonic development and biology of Psammotettix alienus (Dahlbom) (Homoptera Cicadellidae) under laboratory conditions". *Journal of Experimental Biology* 29.1 (1997): 65–80.

Guglielmino, A., A. R. Taddei, and M. Carcupino. “Fine structure of the eggshell of Ommatissus binotatus Fieber (Homoptera, Auchenorrhyncha, Tropiduchidae)”. *International Journal of Insect Morphology and Embryology* 26.2 (1997): 85–89.

Gumovsky, A. V. “A taxonomic revision, biology and morphology of immature stages of the Entedon sparetus species group (Hymenoptera: Eulophidae), egg-larval endoparasitoids of weevils (Coleoptera: Curculionidae)”. *Bulletin of Entomological Research* 97.2 (2007): 139–166.

Gumovsky, A. V. and M. M. Ramadan. “Biology, immature and adult morphology, and molecular characterization of a new species of the genus Entedon (Hymenoptera: Eulophidae) associated with the invasive pest Specularius impressithorax (Coleoptera: Chrysomelidae, Bruchinae) on Erythrina plants”. *Bulletin of Entomological Research* 101.6 (2011): 715–739.

Gumovsky, A. V., S. A. Simutnik, and A. V. Prokhorov. “Life-history review of Oobius zahaikevitshi Trjapitzin, 1963 (Hymenoptera: Encyrtidae), an egg parasitoid of jewel beetles (Coleoptera: Buprestidae)”. *Russian Entomological Journal* 22.3 (2013): 181–188.

Gumovsky, A. V. “Parasitism of Entedon costalis (Hymenoptera: Eulophidae) in Glocianus punctiger (Coleoptera: Curculionidae): an example of intentional discovery of the parasitoid-host association”. *Zootaxa* 2008 (1964): 40–68.

Gumovsky, A., L. Rusina, and L. Firman. “Bionomics and morphological and molecular characterization of Elasmus schmitti and Baryscapus elasmi (Hymenoptera: Chalcidoidea, Eulophidae), parasitoids associated with a paper wasp, Polistes dominulus (Vespoidea, Vespidae)”. *Entomological Science* 10.1 (2007): 21–34.

Guppy, J. C. “Observations on the biology of Plagiognathus chrysanthemi (Hemiptera: Miridae), a pest of birdsfoot trefoil in Ontario”. *Annals of the Entomological Society of America* 56.6 (1963): 804–809.

Guppy, J. C. and F. Meloche. “Life history and description of the immature stages of Dacnusa dryas (Nixon) (Hymenoptera: Braconidae), a European parasite of the alfalfa blotch leafminer (Diptera: Agromyzidae) in eastern Canada”. *The Canadian Entomologist* 119.3 (1987): 281–285.

Gupta, N., V. Khan, S. Kumar, S. Saxena, A. Rashmi, and A. K. Saxena. “Eggshell morphology of selected Indian bird lice (Phthiraptera: Amblycera and Ischnocera)”. *Entomological News* 120.3 (2009): 327–336.

Gurney, A. B. “A synopsis of the Order Zoraptera, with notes on the biology of Zorotypus hubbardi Caudell. Una sinopsis del Orden Zoraptera, con apuntes sobre la biología de Zorotypus hubbardi Caudell”. *Proceedings of the Entomological Society of Washington* 40.3 (1938): 57–87.

Gustafson, J. F. “Biological observations on Timema californica (Phasmoidea: Phasmidae)”. *Annals of the Entomological Society of America* 59.1 (1966): 59–61.

Haas, G. E. and R. J. Dicke. “On Cuterebra horripilum Clark (Diptera: Cuterebridae) parasitizing cottontail rabbits in Wisconsin”. *The Journal of Parasitology* 44.5 (1958): 527–540.

Habeck, D. H. “Description of immature stages of the Chinese rose beetle, Adoretus sinicus Burmeister (Coleoptera: Scarabaeidae)”. *Proceedings of the Hawaiian Entomological Society* 18.2 (1963): 251–258.

Hafez, M., M. F. S. Tawfik, and A. A. Ibrahim. “The immature stages of Chelonus inanitus (L.), a parasite of the Cotton Leafworm, Spodoptera littoralis (Boisd.), in Egypt (Hym., Braconidae)”. *Deutsche Entomologische Zeitschrift* 27.1-3 (1980): 29–38.

Haga, K. “Oogenesis and embryogenesis of the idolothripine thrips, Bactrothrips brevitubus (Thysanoptera, Phlaeothripidae)”. *Recent Advances in Insect Embryology in Japan*. ISEBU Co.: Tsukuba Science City, 1985. 45–106.

Hagan, H. R. “Observations on the embryonic development of the mantid Paratenodera sinensis”. *Journal of Morphology* 30.1 (1917): 223–243.

Hagan, H. R. “The embryogeny of the polyctenid, Hesperoctenes fumarius Westwood, with reference to viviparity in insects”. *Journal of Morphology* 51.1 (1931): 1–117.

Hager, B. J. and F. E. Kurczewski. “Nesting behavior of Ammophila harti (Fernald) (Hymenoptera: Sphecidae)”. *American Midland Naturalist* 116.1 (1986): 7–24.

Hale, H. M. “Notes on eggs, habits and migration of some Australian aquatic bugs (Corixidae and Notonectidae)”. *The South Australian Naturalist* 5.4 (1924): 133–135.

Hamilton, R. W. “New life cycle data for two western North American weevils (Coleoptera: Rhynchitidae), with a summary of North American rhynchitid biology”. *The Coleopterists' Bulletin* 48.4 (1994): 331–343.

Hammer, O. “Biological and ecological investigations on flies associated with pasturing cattle and their excrement”. *Videnskabelige Meddelelser Naturhistorisk Forening i København* 105 (1941): 1–257.

Hamon, A. B., P. L. Lambdin, and M. Kosztarab. “Eggs and wax secretion of Kermes kingi”. *Annals of the Entomological Society of America* 68.6 (1975): 1077–1078.

Hancock, J. L. *The Tettigidae of North America*. Chicago: Published by special grant of Mrs. F. G. Logan, 1902.

Hanley, R. S. and K. Setsuda. “Immature stages of Oxyporus japonicus Sharp (Coleoptera: Staphylinidae: Oxyporinae), with notes on patterns of host use”. *The Pan-Pacific Entomologist* 75.2 (1999): 94–102.

Hanley, R. S. and M. A. Goodrich. “Natural history, development and immature stages of Oxyporus stygicus Say (Coleoptera: Staphylinidae: Oxyporinae)”. *The Coleopterists' Bulletin* 48.3 (1994): 213–225.

Hansell, M. H. “Brood development in the subsocial wasp Parischnogaster mellyi (Saussure) (Stenogastrinae, Hymenoptera)”. *Insectes Sociaux* 29.1 (1982): 3–14.

Hansen, M. “Observations on the immature stages of Georissidae (Coleoptera: Hydrophiloidea), with remarks on the evolution of the hydrophiloid egg cocoon”. *Invertebrate Systematics* 14.6 (2000): 907–916.

Harber, P. A. and J. A. Mutchmor. “The early embryonic development of Culiseta inornata (Diptera: Culicidae)”. *Annals of the Entomological Society of America* 63.6 (1970): 1609–1614.

Hardy, R. J. “Some aspects of the biology and behaviour of Adoryphorus couloni (Burmeister) (Coleoptera: Scarabaeidae: Dynastinae)”. *Australian Journal of Entomology* 20.1 (1981): 67–74.

Haridass, E. T. “Ultrastructure of the eggs of Reduviidae: I. Eggs of Piratinae (Insecta-Heteroptera)”. *Proceedings of the Indian Academy of Sciences: Animal Sciences* 94.5 (1985): 533–545.

Haridass, E. T. “Ultrastructure of the eggs of Reduviidae: II eggs of Harpactorinae (Insecta: Heteroptera)”. *Proceedings of the Indian Academy of Sciences: Animal Sciences* 95.2 (1986): 237–246.

Haridass, E. T. “Ultrastructure of the eggs of Reduviidae: III. Eggs of Triatominae and Echtrichodiinae (Insecta-Heteroptera)”. *Proceedings of the Indian Academy of Sciences: Animal Sciences* 95.4 (1986): 447–456.

Haridass, E. T. “Ultrastructure of the eggs of Reduviidae: IV. Eggs of Rhaphidosomatinae (Insecta–Heteroptera)”. *Proceedings of the Indian Academy of Sciences: Animal Sciences* 97.1 (1988): 49–54.

Harlan, D. P. and W. R. Enns. “Surface-printing of grasshopper eggs for identification”. *Annals of the Entomological Society of America* 59.5 (1966): 1018–1020.

Harley, K. L. S. and R. C. Kassulke. “Apion antiquum (Curculionoidea: Apionidae) for biological control of the weed Emex australis”. *Australian Journal of Entomology* 14.3 (1975): 271–276.

Harris, A. C. “Nesting behaviour, life history and description of the mature larva of the beetle predator, Podagritus parrotti Leclercq (Hymenoptera: Sphecidae: Crabroninae)”. *Journal of the Royal Society of New Zealand* 28.4 (1998): 591–604.

Harris, A. C. “Pompilidae (Insecta: Hymenoptera)”. *Fauna of New Zealand* 12 (1987): 1–160.

Hartley, J. C. “The egg of Tetrix (Tetrigidae, Orthoptera), with a discussion on the probable significance of the anterior horn”. *Quarterly Journal of Microscopical Science* 3.62 (1962): 253–259.

Hartley, J. C. “The respiratory system of the egg-shell of Homorocoryphus nitidulus vicinus (Orthoptera, Tettigoniidae)”. *Journal of Experimental Biology* 55 (1971): 165–176.

Hartley, J. C. “The shell of acridid eggs”. *Quarterly Journal of Microscopical Science* 3.58 (1961): 249–255.

Hartley, J. C. “The structure and function of the egg-shell of Deraeocoris ruber L. (Heteroptera, Miridae)”. *Journal of Insect Physiology* 11.1 (1965): 103–109.

Hartley, J. C. “The structure of the eggs of the British Tettigoniidae (Orthoptera)”. *Proceedings of the Royal Entomological Society of London. Series A, General Entomology* 39.7-9 (1964): 111–117.

Hasbenli, A., Z. Suludere, S. Candan, and F. Bayrakdar. “Chorionic structure of the eggs of five Laphriinae species (Diptera: Asilidae) from Turkey”. *Journal of the Entomological Research Society* 10.3 (2008): 47–60.

Hasenpusch, J., D. A. Lane, and M. S. Moulds. “The life history of the hawk moth Macroglossum insipida papuanum Rothschild and Jordan, 1903 (Lepidoptera: Sphingidae)”. *Australian Entomologist* 39.2 (2012): 79–86.

Hasenpusch, J. and P. D. Brock. “Studies on the Australian stick insect genus Ctenomorpha Gray (Phasmida: Phasmatidae: Phasmatinae), with the description of a new large species”. *Zootaxa* 1282.1 (2006): 1–15.

Hassan, A. I. “The biology of some British Delphacidae (Homopt.) and their parasites with special reference to the Strepsiptera”. *Transactions of the Royal Entomological Society of London* 89.9 (1939): 345–384.

Havelka, J., V. Landa Jr, and V. Landa. “Embryogenesis of Aphidoletes aphidimyza (Diptera: Cecidomyiidae): Morphological markers for staging of living embryos”. *European Journal of Entomology* 104.1 (2007): 81–87.

Haverfield, L. E. “A note on the mating ritual and biology of Eleodes hispilabris connexa (Coleoptera: Tenebrionidae)”. *Journal of the Kansas Entomological Society* 38.4 (1965): 389–391.

Hayashi, F. “Life history variation in a dobsonfly, Protohermes grandis (Megaloptera: Corydalidae): effects of prey availability and temperature”. *Freshwater Biology* 19.2 (1988): 205–216.

Hayashi, F. and H. Suzuki. “Fireflies with or without prespermatophores: Evolutionary origins and life-history consequences”. *Entomological Science* 6.1 (2003): 3–10.

Hayes, W. P. “A comparative study of the history of certain phytophagous Scarabaeid beetles”. *Kansas Technical Bulletin* 16 (1925): 3–133.

Headrick, D. H. and R. D. Goeden. “Description of the immature stages of Paracantha gentilis (Diptera: Tephritidae)”. *Annals of the Entomological Society of America* 83.2 (1990): 220–229.

Hedlin, A. F. “The Life History and Habits of a Midge, Contarinia oregonensis Foote (Diptera: Cecidomyiidae) in Douglas-fir Cones”. *The Canadian Entomologist* 93.11 (1961): 952–967.

Hegazi, E. M. and E. Führer. “Instars of Microplitis rufiventris [Hym.: Braconidae] and their relative developmental speed under different photoperiods”. *Entomophaga* 30.3 (1985): 231–243.

Heidemann, O. “Some remarks on the eggs of North American species of Hemiptera-Heteroptera”. *Proceedings of the Entomological Society of Washington* 13.3 (1911): 128–140.

El-Helaly, M. S., A. Y. El-Shazli, and F. H. El-Gayar. “Morphological studies on immature stages of Bemisia tabaci, Gennadius (Homoptera, Aleyrodidae)”. *Zeitschrift für Angewandte Entomologie* 68.1-4 (1971): 403–408.

Hemenway, R. and W. H. Whitcomb. “The life history of Disonycha glabrata (Coleoptera: Chrysomelidae)”. *Journal of the Kansas Entomological Society* 41.2 (1968): 174–178.

Heming, B. S. “Origin and fate of germ cells in male and female embryos of Haplothrips verbasci (Osborn) (Insecta, Thysanoptera, Phlaeothripidae)”. *Journal of Morphology* 160.3 (1979): 323–343.

Hennemann, F. H. “Notes on the genera Andropromachus Carl, 1913 and Spinohirasea Zompro, 2001”. *Phasmid Studies* 15.1 (2007): 15–26.

Hennemann, F. H. “PSG 28, Eurycnema herculeana (Charpentier)”. *Phasmid Studies* 34.2 (1992): 34–37.

Hennemann, F. H. and O. V. Conle. “Papuacocelus papuanus n. gen., n. sp.—a new Eurycanthinae from Papua New Guinea, with notes on the genus Dryococelus Gurney, 1947 and description of the egg (Phasmatodea: Phasmatidae: Eurycanthinae)”. *Zootaxa* 1375 (2006): 31–49.

Hennemann, F. H. and O. V. Conle. “Studies on neotropical Phasmatodea VII. Descriptions of a new genus and four new species of Diapheromerinae from Peru and Bolivia (Phasmatodea: ”Anareolatae”: Diapheromeridae)”. *Mitteilungen der Münchner Entomologischen Gesellschaft* 97 Suppl. (2007): 89–112.

Hennemann, F. H. and O. V. Conle. “Studies on Neotropical Phasmatodea X: Redescriptions of Aplopocranidium Zompro, 2004 and Jeremia Redtenbacher, 1908, with a Survey of the Tribe Cladomorphini Brunner v. Wattenwyl, 1893 and Keys to the Genera (Insecta: Phasmatodea: “Anareolatae”: Cladomorph)”. *Journal of Orthoptera Research* 19.1 (2010): 101–113.

Hennemann, F. H. and O. V. Conle. “Studies on neotropical Phasmatodea XIII: the genus Paracalynda Zompro, 2001 with notes on Eusermyleformia Bradler, 2009 (Insecta: Phasmatodea: Diapheromerinae: Diapheromerini)”. *Journal of Orthoptera Research* 21.1 (2012): 57–64.

Hennemann, F. H. and O. V. Conle. "Studies on Neotropical Phasmatodea XIV: Revisions of the Central American Genera Hypocyrtus Redtenbacher, 1908 and Rhynchacris Redtenbacher, 1908 (Phasmatodea:"Anareolatae": Xerosomatinae: Hesperophasmatini)". *Journal of Orthoptera Research* 21.1 (2012): 65–89.
Hennemann, F. H. and O. V. Conle. "Studies on New Guinean giant stick-insects of the tribe Stephanacridini Günther, 1953, with the descriptions of a new genus and three new species of Stephanacris Redtenbacher, 1908 (Phasmatodea:"Anareolatae")". *Zootaxa* 1283 (2006): 1–24.
Hennemann, F. H. and O. V. Conle. "The genus Paracyphocrania Redtenbacher, 1908 (Phasmatodea: Phasmatinae: Phasmatini)". *Zoologische Mededelingen* 80.4 (2006): 91–101.
Hennemann, F. H., O. V. Conle, P. D. Brock, and F. Seow-Choen. "Revision of the Oriental subfamily Heteropteryginae Kirby, 1896, with a re-arrangement of the family Heteropterygidae and the descriptions of five new species of Haaniella Kirby, 1904. (Phasmatodea: Areolatae: Heteropterygidae)". *Zootaxa* 4159.1 (2016): 1–219.
Hennemann, F. H., O. V. Conle, and D. E. Perez-Gelabert. "Studies on Neotropical Phasmatodea XVI: Revision of Haplopodini Günther, 1953 (rev. stat.), with notes on the subfamily Cladomorphinae Bradley & Galil, 1977 and the descriptions of a new tribe, four new genera and nine new species (Phasmatodea:"Anareolatae": Phasmatidae: Cladomorphinae)". *Zootaxa* 4128.1 (2016): 1–211.
Hennemann, F. H., O. V. Conle, and S. Suzuki. "A study of the members of the tribe Phasmatini Gray, 1835, that occur within the boundaries of Wallacea (Phasmatodea: Phasmatidae: Phasmatinae:"Lanceocercata")". *Zootaxa* 4008.1 (2015): 1–74.
Henry, C. S. "An evolutionary and geographical overview of Repagula (abortive eggs) in the Ascalaphidae (Neuroptera)". *Proceedings of the Entomological Society of Washington* 80.1 (1978): 75–86.
Henry, C. S. "Eggs and Rapagula of Ululodes and Ascaloptynx (Neuroptera: Ascalaphidae): A Comparative Study". *Psyche* 79.1-2 (1972): 1–22.
Henry, C. S. "The egg, repagulum, and larva of Byas albistigma (Neuroptera: Ascalaphidae): morphology, behaviour and phylogenetic significance". *Systematic Entomology* 3.1 (1978): 9–18.
Hentz, M. G. and G. S. Nuessly. "Morphology and biology of Diomus terminatus (Coleoptera: Coccinellidae), a predator of Sipha flava (Homoptera: Aphidae)". *The Florida Entomologist* 85.1 (2002): 276–278.
Heraty, J. M. and D. C. Darling. "A new genus and species of Perilampidae (Hymenoptera: Chalcidoidea) with uncertain placement in the family". *Journal of the Entomological Society of Ontario* 138 (2007): 33–47.
Heraty, J. M. and K. N. Barber. "Biology of Obeza floridana (Ashmead) and Pseudochalcura gibbosa (Provancher) (Hymenoptera: Eucharitidae)". *Proceedings of the Entomological Society of Washington* 92.2 (1990): 248–258.
Heraty, J. M. and D. C. Darling. "Comparative morphology of the planidial larvae of Eucharitidae and Perilampidae (Hymenoptera: Chalcidoidea)". *Systematic Entomology* 9.3 (1984): 309–328.
Heraty, J., D. Hawks, J. S. Kostecki, and A. Carmichael. "Phylogeny and behaviour of the Gollumiellinae, a new subfamily of the ant-parasitic Eucharitidae (Hymenoptera: Chalcidoidea)". *Systematic Entomology* 29.4 (2004): 544–559.
Hernández-Mejía, B. C., A. Flores-Gallardo, and J. Llorente. "Comparación Morfológica del Corion de Especies de los Géneros Pieriballia, Itaballia, y Perrhybris (Lepidoptera: Pieridae: Pierinae), y sus Implicaciones Filogenéticas". *Southwestern Entomologist* 38.2 (2013): 275–291.
Hernández-Mejía, B. C., A. Flores-Gallardo, and J. Llorente-Bousquets. "Morfología del corion en especies de los géneros Ascia y Ganyra y su comparación con otros géneros próximos de Pierinae (Lepidoptera: Pieridae)". *Southwestern Entomologist* 39.1 (2014): 119–134.
Hernández-Mejía, B. C., A. Flores-Gallardo, and J. Llorente-Bousquets. "Morfología del Corion en la Subfamilia Coliadinae (Lepidoptera: Pieridae) Chorionic Morphology in the Coliadinae Subfamily (Lepidoptera: Pieridae)". *Southwestern Entomologist* 39.4 (2014): 853–886.
Hernández-Mejía, C., A. Flores-Gallardo, and J. Llorente-Bousquets. "Morfología del Corion en Leptophobia (Lepidoptera: Pieridae) e Importancia Taxonómica". *Southwestern Entomologist* 40.2 (2015): 351–368.
Hernández, J. M. "Estudio de los caracteres del huevo en diversos Cerambycidae ibéricos y su interés taxonómico (Coleoptera)". *Graellsia* 47 (1991): 49–59.

Herring, J. L. “The genus Halobates (Hemiptera: Gerridae)”. *Pacific Insects* 3.2-3 (1961): 223–305.

Herth, W. and K. Sander. “Mode and timing of body pattern formation (regionalization) in the early embryonic development of cyclorrhaphic dipterans (Protophormia, Drosophila)”. *Development Genes and Evolution* 172.1 (1973): 1–27.

Hesami, S., H. Seyedoleslami, and R. Ebadi. “Biology of Anagrus atomus (Hymenoptera: Mymaridae), an egg parasitoid of the grape leafhopper Arboridia kermanshah (Homoptera: Cicadellidae)”. *Entomological Science* 7.3 (2004): 271–276.

Hess, W. N. “Notes on the biology of some common Lampyridae”. *The Biological Bulletin* 38.2 (1920): 39–76.

Hill, C. C. and W. T. Emery. “The biology of Platygaster herrickii, a parasite of the Hessian fly”. *Journal of Agricultural Research* 55.19 (1937): 199–213.

Hill, L. “Eggs of some tasmanian noctuidae (Lepidoptera)”. *Australian Entomological Magazine* 9.4 (1982): 49–59.

Hinton, H. E. “The fine structure and biology of the egg-shell of the wheat bulb fly, Leptohylemyia coarctata”. *Quarterly Journal of Microscopical Science* 3.62 (1962): 243–251.

Hinton, H. E. “The structure and function of the egg-shell in the Nepidae (Hemiptera)”. *Journal of Insect Physiology* 7.3 (1961): 224–257.

Hinton, H. E. “The structure of the shell and respiratory system of the eggs of Helopeltis and related genera (Hemiptera, Miridae)”. *Proceedings of the Zoological Society of London* 139.3 (1962): 483–488.

Ho, G. W. C. “Necroscia perplexus (Redtenbacher, 1908) comb. nov.(Phasmatodea: Diapheromeridae: Necrosciinae), a new species to China”. *Phasmid Studies* 18 (2013): 19–23.

Ho, G. W.-C. “Remarks on the genus Necroscia (Phasmatodea, Diapheromeridae, Necrosciinae) from China, with two new records, one new synonym, and one new combination and key to the species”. *Journal of Orthoptera Research* 22.1 (2013): 21–27.

Ho, G. W.-C., X.-Y. Liu, J. Bresseel, and J. Constant. “Brockphasma spinifemoralis gen. et spec. nov.: a new phasmid genus and new species of Neohiraseini (Phasmida: Necrosciinae) from Vietnam”. *Zootaxa* 3826.1 (2014): 282–290.

Ho, J.-Z., P.-H. Chiang, C.-H. Wu, and P.-S. Yang. “Life cycle of the aquatic firefly Luciola ficta (Coleoptera: Lampyridae)”. *Journal of Asia-Pacific Entomology* 13.3 (2010): 189–196.

Ho, K., O. M. Dunin-Borkowski, and M. Akam. “Cellularization in locust embryos occurs before blastoderm formation”. *Development* 124.14 (1997): 2761–2768.

Hokyo, N., K. Kiritani, F. Nakasuji, and M. Shiga. “Comparative biology of the two scelionid egg parasites of Nezara viridula L. (Hemiptera: Pentatomidae)”. *Applied Entomology and Zoology* 1.2 (1966): 94–102.

Holder, M. W. and S. W. Wilson. “Stages of the planthopper Prokelisia crocea (Van Duzee) (Homoptera: Delphacidae)”. *Journal of the New York Entomological Society* 100.3 (1992): 491–497.

Honan, P. “Notes on the biology, captive management and conservation status of the Lord Howe Island Stick Insect (Dryococelus australis) (Phasmatodea)”. *Journal of Insect Conservation* 12.3 (2008): 399–413.

Hood, G. R. and J. R. Ott. “Generational shape shifting: changes in egg shape and size between sexual and asexual generations of a cyclically parthenogenic gall former”. *Entomologia Experimentalis et Applicata* 141.1 (2011): 88–96.

Horak, M., M. F. Day, C. Barlow, E. D. Edwards, Y. N. Su, and S. L. Cameron. “Systematics and biology of the iconic Australian scribbly gum moths Ogmograptis Meyrick (Lepidoptera: Bucculatricidae) and their unique insectplant interaction”. *Invertebrate Systematics* 26.4 (2012): 357–398.

Hori, K. and T. Hanada. “Biology of Lygus disponsi Linnavuroi (Hemiptera, Miridae) in Obihiro”. *Research Bulletin of Obihiro University* 6.2 (1970): 304–317.

Horsburgh, R. L. and D. Asquith. “The eggs and oviposition sites of Diaphnidia capitata (Hemiptera: Miridae) on apple trees”. *The Canadian Entomologist* 102.10 (1970): 1316–1319.

Horsburgh, R. L. and D. Asquith. “The eggs and oviposition sites of Hyaliodes vitripennis on apple trees (Miridae: Hemiptera)”. *The Canadian Entomologist* 100.2 (1968): 199–201.

Horsfall, W. R. and G. B. Craig. “Eggs of Floodwater Mosquitoes IV. Species of Aedes Common in Illinois (Diptera: Culicidae)”. *Annals of the Entomological Society of America* 49.4 (1956): 368–374.
Horsfall, W. R., R. C. Miles, and J. T. Sokatch. “Eggs of floodwater mosquitoes. I. Species of Psorophora (Diptera: Culicidae)”. *Annals of the Entomological Society of America* 45.4 (1952): 618–624.
Horsfall, W. R. and F. R. Voorhees. “Eggs of Floodwater Mosquitoes XIV. Northern Aedes (Diptera: Culicidae)”. *Annals of the Entomological Society of America* 65.1 (1972): 123–126.
Horsfall, W. R., F. R. Voorhees, and E. W. Cupp. “Eggs of floodwater mosquitoes. XIII. Chorionic sculpturing”. *Annals of the Entomological Society of America* 63.6 (1970): 1709–1716.
Hosseinie, S. O. “Comparative studies on the developmental stages of three species of Tropisternus (Coleoptera: Hydrophilidae)”. *Internationale Revue der gesamten Hydrobiologie und Hydrographie* 61.6 (1976): 847–857.
Houghton, D. C. and K. W. Stewart. “Immature life stage descriptions and distribution of Culoptila cantha (Ross) (Trichoptera: Glossosomatidae)”. *Proceedings of the Entomological Society of Washington* 100.3 (1998): 511–520.
Houston, T. F. “Brood cells, life-cycle stages and development of some earth-borer beetles in the genera Bolborhachium, Blackburnium and Bolboleaus (Coleoptera: Geotrupidae), with notes on captive rearing and a discussion of larval diet”. *Austral Entomology* 55 (2015): 49–62.
Houston, T. F. “Egg gigantism in some Australian earth-borer beetles (Coleoptera: Geotrupidae: Bolboceratinae) and its apparent association with reduction or elimination of larval feeding”. *Australian Journal of Entomology* 50.2 (2011): 164–173.
Houston, T. F. “Observations of the biology and immature stages of the sandgroper Cylindraustralia kochii (Saussure), with notes on some congeners (Orthoptera: Cylindrachetidae)”. *Records of the Western Australian Museum* 23.3 (2007): 219–234.
Howard, G. W. “Immature stages and affinities of the southern lechwe warblefly, Strobiloestrus vanzyli Zumpt (Diptera: Oestridae)”. *Journal of Natural History* 14.5 (1980): 669–683.
Howard, R. D. and D. H. Kistner. “The eggs of Trichopsenius depressus and T. forsteri”. *Sociobiology* 3.2 (1978): 99–106.
Howden, H. F. “Larval and adult characters of Frickius Germain, its relationship to the Geotrupini, and a phytogeny of some major taxa in the Scarabaeoidea (Insecta: Coleoptera)”. *Canadian Journal of Zoology* 60.11 (1982): 2713–2724.
Howden, H., A. Howden, and G. Holloway. “Digging down under: Australian Bolboceratini, their habits and a list of species (Coleoptera: Scarabaeoidea: Geotrupidae)”. *Zootaxa* 1499 (2007): 47–59.
Hsiung, C.-C. “The Identity of Japanese Megacrania Kaup (Phasmatodea: Phasmatidae)”. *Journal of Orthoptera Research* 22.1 (2013): 67–68.
Hu, G. Y. and J. H. Frank. “Biology of Neohypnus pusillus (Sachse) (Coleoptera: Staphylinidae) and its predation on immature horn flies in the laboratory”. *The Coleopterists' Bulletin* 49.1 (1995): 43–52.
Hu, G. Y. and J. H. Frank. “Structural comparison of the chorion surface of five Philonthus species (Coleoptera: Staphylinidae)”. *Proceedings of the Entomological Society of Washington* 97.3 (1995): 582–598.
Hu, J., P. Wang, and W. Zhang. “Two types of embryos with different functions are generated in the polyembryonic wasp Macrocentrus cingulum (Hymenoptera: Braconidae)”. *Arthropod Structure & Development* 44.6 (2015): 677–687.
Hu, J., X. Yu, W. Fu, and W. Zhang. “A Helix pomatia lectin binding protein on the extraembryonic membrane of the polyembryonic wasp Macrocentrus cingulum protects embryos from being encapsulated by hemocytes of host Ostrinia furnaclis”. *Developmental & Comparative Immunology* 32.4 (2008): 356–364.
Huang, F., M. Shi, X.-X. Chen, G.-Y. Ye, and J.-H. He. “External morphology and development of immature stages of Diadegma semiclausum (Hymenoptera: Ichneumonidae), an important endoparasitoid of Plutella xylostella (Lepidoptera: Plutellidae)”. *Annals of the Entomological Society of America* 102.3 (2009): 532–538.
Huang, Y. S.-F. and P. D. Brock. “A new species of Phasmotaenia Navas (Phasmida: Phasmatidae) from Taiwan”. *Journal of Orthoptera Research* 10.1 (2001): 9–14.

Huang, D.-y., A. Nel, O. Zompro, and A. Waller. "Mantophasmatodea now in the Jurassic". *Naturwissenschaften* 95.10 (2008): 947–952.

Hughes, L. and M. Westoby. "Capitula on stick insect eggs and elaiosomes on seeds: convergent adaptations for burial by ants". *Functional Ecology* 6.6 (1992): 642–648.

Huie, L. H. "XV.—The formation of the germ-band in the egg of the holly Tortrix moth, Eudemis nævana (Hb.)" *Proceedings of the Royal Society of Edinburgh* 38 (1919): 154–165.

Hungerford, H. B. "Concerning the egg of Polystoechotes punctatus Fabr. (Neuroptera)". *Bulletin of the Brooklyn Entomological Society* 26.1 (1931): 22–23.

Hungerford, H. B. "Notes on the eggs of Corixidae". *Bulletin of the Brooklyn Entomological Society* 18.1 (1923): 13–16.

Hungerford, H. B. "The eggs of Corixidae (Hemiptera)". *Journal of the Kansas Entomological Society* 21.4 (1948): 141–146.

Husain, M. A. and M. L. Roonwal. "Studies on Schistocerca gregaria Forsk. I. The micropyle in Schistocerca gregaria Forsk. and some other Acrididae". *Indian Journal of Agricultural Science* 3.4 (1933): 639–645.

Hutcheson, J. "Notes of the ecology of Ectopsis ferrugalis (Curculionidae)". *New Zealand Entomologist* 14.1 (1991): 41–44.

Hwang, W.-S., R. Hanley, and K.-J. Ahn. "Immature Stages of Oxyporus germanus Sharp (Coleoptera: Staphylinidae: Oxyporinae)". *Journal of the Kansas Entomological Society* 75.3 (2002): 214–221.

Hynes, H. B. N. "Observations on the adults and eggs of Australian Plecoptera". *Australian Journal of Zoology Supplementary Series* 29 (1974): 37–52.

Hynes, H. B. N. "The neoperlinae of the ethiopian region (Plecoptera, Perlidae)". *Transactions of the Royal Entomological Society of London* 103.3 (1952): 85–108.

Hynes, H. B. N. "The taxonomy and ecology of the nymphs of British Plecoptera with notes on the adults and eggs". *Transactions of the Royal Entomological Society of London* 91.10 (1941): 459–557.

Ikeda, H., T. Kagaya, K. Kubota, and T. Abe. "Evolutionary relationships among food habit, loss of flight, and reproductive traits: life-history evolution in the Silphinae (Coleoptera: Silphidae)". *Evolution* 62.8 (2008): 2065–2079.

Ikeda, Y. and R. Machida. "Embryogenesis of the dipluran Lepidocampa weberi Oudemans (Hexapoda: Diplura, Campodeidae): formation of dorsal organ and related phenomena". *Journal of Morphology* 249.3 (2001): 242–251.

Imhof, J. E. and S. M. Smith. "Oviposition behaviour, egg-masses and hatching response of the eggs of five Nearctic species of Simulium (Diptera: Simuliidae)". *Bulletin of Entomological Research* 69.3 (1979): 405–425.

Infante, F., J. Valdez, D. I. Penagos, and J. F. Barrera. "Description of the life stages of Cephalonomia stephanoderis (Hymenoptera: Bethylidae), a parasitoid of Hypothenemus hampei (Coleoptera: Scolytidae)". *Vedalia* 1.1 (1994): 13–18.

Irvin, N. A. and M. S. Hoddle. "Egg maturation, oosorption, and wing wear in Gonatocerus ashmeadi (Hymenoptera: Mymaridae), an egg parasitoid of the glassy-winged sharpshooter, Homalodisca vitripennis (Hemiptera: Cicadellidae)". *Biological Control* 48.2 (2009): 125–132.

Irwin, M. E. and B. R. Stuckenberg. "A description of the female, egg and first-instar larva of Tongamya miranda, with notes on oviposition and the habitat of the species (Diptera: Apioceridae)". *Annals of the Natal Museum* 21.2 (1972): 439–453.

Isidoro, N. and A. Lucchi. "Eggshell fine morphology of Allocontarinia sorghicola (Coq.)(Diptera: Cecidomyiidae)". *Entomologica* 24 (2016): 127–138.

Ivanova-Kasas, O. M. "Biologija i embrionalnoje razvitije Eurytoma aciculata Ratz (Hymenoptera, Eurytomidae)". *Entomologiceskoe Obozrenie* 37.1 (1958): 5–23.

Ivanova-Kasas, O. M. "Die embryonale Entwicklung der Blattwespe Pontania capreae L.(Hymenoptera, Tenthredinidae)". *Zoologische Jahrbücher, Abteilung für Anatomie und Ontogenie der Tiere* 77 (1959): 193–228.

Ivanova-Kasas, O. M. "Polyembryony in insects". *Developmental Systems: Insects*. London: Academic Press, 1972. 243–271.

Ivey, R. K., J. C. Bailey, B. P. Stark, and D. L. Lentz. "A preliminary report of egg chorion features in dragonflies (Anisoptera)". *Odonatologica* 17.4 (1988): 393–399.

Iwabuchi, K. "Early Embryonic Development of a Polyembryonic Wasp, Litomastix maculata Ishii, in vivo and in vitro". *Applied Entomology and Zoology* 26.4 (1991): 563–570.

Iwaki, M. and W. Choochote. "Scanning electron microscopy of eggs of Mansonia uniformis, Ma. indiana, Ma. annulifera, and Ma. annulata (Diptera: Culicidae)". *Journal of Medical Entomology* 28.3 (1991): 334–339.

Iwan, D. and S. Becvar. "Description of the early stages of Anomalipus plebejus plebejulus (Coleoptera: Tenebrionidae) from Zimbabwe with notes on the classification of the Opatrinae". *European Journal of Entomology* 97.3 (2000): 403–412.

Iwata, K. "Bionomics of non-social wasps in Thailand". *Nature and Life in Southeast Asia* 3 (1964): 323–383.

Iwata, K. "Egg giantism in subsocial Hymenoptera, with ethological discussion on tropical bamboo carpenter bees". *Nature Life Southeast Asia* 3 (1964): 399–435.

Iwata, K. "Large-sized eggs in Curculionoidea (Coleoptera)". *Research Bulletin of Hyogo Agricultural College* 7 (1966): 43–45.

Iwata, K. "Ovarian eggs in Scarabaeoidea (Coleoptera)". *Seibutsu Kenkyu* 10 (1966): 1–3.

Iwata, K. "The comparative anatomy of the ovary in Hymenoptera. (Records on 64 species of Aculeata in Thailand, with descriptions of ovarian eggs)". *Mushi* 38 (1965): 101–109.

Iwata, K. "The comparative anatomy of the ovary in Hymenoptera. Part I. Aculeata". *Mushi* 29 (1955): 17–34.

Iwata, K. "The comparative anatomy of the ovary in Hymenoptera. Part II. Symphyta". *Mushi* 31 (1958): 47–60.

Iwata, K. "The comparative anatomy of the ovary in Hymenoptera. Part IV: Proctotrupoidea and Agriotypidae (Ichneumonoidea) with descriptions of ovarian eggs". *Kontyû* 27.1 (1959): 18–21.

Iwata, K. "The comparative anatomy of the ovary in Hymenoptera. Part V. Ichneumonidae". *Acta Hymenopterologica* 1.2 (1960): 115–169.

Iwata, K. "The comparative anatomy of the ovary in Hymenoptera. Supplement of Aculeata with descriptions of ovarian eggs of certain species". *Acta Hymenopterologica* 1.2 (1960): 205–211.

Iwata, K. "The comparative anatomy of the ovary in Hymenoptera. VI. Chalcidoidea with descriptions of ovarian eggs". *Acta Hymenopterologica* 1.4 (1962): 383–391.

Iwata, K. and S. H. F. Sakagami. "Gigantism and dwarfism in bee eggs in relation to the mode of life, with notes on the number of ovarioles". *Japanese Journal of Ecology* 16.1 (1966): 4–16.

Jacas, J.-A., J. E. Peña, and R. E. Duncan. "Morphology and development of immature stages of Fidiobia dominica (Hymenoptera: Platygastridae: Sceliotrachelinae)". *Annals of the Entomological Society of America* 100.3 (2007): 413–417.

Jacas, J.-A., J. E. Peña, and R. E. Duncan. "Morphology and development of the immature stages of Brachyufens osborni (Hymenoptera: Trichogrammatidae), an egg parasitoid of broad-nosed weevil species (Coleoptera: Curculionidae)". *Annals of the Entomological Society of America* 102.1 (2009): 112–118.

Jackson, C. G., J. S. Delph, and E. G. Neemann. "Development, longevity and fecundity of Chelonus blackburni [Hym.: Braconidae] as a parasite of Pectinophora gossypiella [Lep.: Gelechiidae]". *Entomophaga* 23.1 (1978): 35–42.

Jackson, D. J. "Egg-laying and egg-hatching in Agabus bipustulatus L., with notes on oviposition in other species of Agabus (Coleoptera: Dytiscidae)". *Transactions of the Royal Entomological Society of London* 110.3 (1958): 53–80.

Jackson, D. J. "Observations on the biology of Caraphractus cinctus Walker (Hymenoptera: Mymaridae), a parasitoid of the eggs of Dytiscidae". *Transactions of the Royal Entomological Society of London* 110.17 (1958): 533–554.

Jackson, D. J. "Observations on the biology of Caraphractus cinctus Walker (Hymenoptera: Mymaridae), a parasitoid of the eggs of Dytiscidae (Coleoptera). 2. Immature stages and seasonal history with a review of mymarid larvae". *Parasitology* 51.3-4 (1961): 269–294.

Jackson, D. J. “Observations on the life-history of Mestocharis bimacularis (Dalman) (Hym. Eulophidae), a parasitoid of the eggs of Dytiscidae”. *Opuscula Entomologica, Lund* 29 (1964): 81–97.

Jacobus, L. M. and W. P. McCafferty. “Revision of Ephemerellidae genera (Ephemeroptera)”. *Transactions of the American Entomological Society* 134.1 (2008): 185–274.

Jacobus, L. M., R. L. Newell, and W. P. McCafferty. “First adult and egg descriptions of Caudatella edmundsi (Ephemeroptera: Ephemerellidae) from Montana (USA), with habitat observations”. *Entomological News* 117.2 (2006): 175–180.

Jansen-G, S. and C. E. Sarmiento. “A new species of high mountain Andean fig wasp (Hymenoptera: Agaonidae) with a detailed description of its life cycle”. *Symbiosis* 45.1 (2008): 135–141.

Jarjees, E. A. and D. J. Merritt. “Development of Trichogramma australicum Girault (Hymenoptera: Trichogrammatidae) in Helicoverpa (Lepidoptera: Noctuidae) host eggs”. *Australian Journal of Entomology* 41.4 (2002): 310–315.

Javahery, M. “Development of eggs in some true bugs (Hemiptera–Heteroptera). Part I. Pentatomoidea”. *The Canadian Entomologist* 126.2 (1994): 401–433.

Javahery, M. “Natural history of Reduvius personatus Linnaeus (Hemiptera: Heteroptera: Reduviidae) in North America”. *Munis Entomology & Zoology* 8.2 (2013): 685–703.

Jennings, D. T. “Life history and habits of the southwestern pine tip moth, Rhyacionia neomexicana (Dyar) (Lepidoptera: Olethreutidae)”. *Annals of the Entomological Society of America* 68.3 (1975): 597–606.

Jensen, D. D. “Notes on the life history and ecology of blossom midge, Contarinia lycopersici Felt (Diptera: Cecidomyiidae)”. *Proceedings of the Hawaiian Entomological Society* 14.1 (1950): 91–100.

Jerez, V. and R. Briones. “Mylassa crassicollis (Blanchard, 1851) (Coleoptera: Chrysomelidae: Cryptocephalinae): Biology and Description of Immature Stages”. *The Coleopterists Bulletin* 64.1 (2010): 31–38.

Jervis, M. A. “Studies on oviposition behaviour and larval development in species of Chalarus (Diptera, Pipunculidae), parasites of typhlocybine leafhoppers (Homoptera, Cicadellidae)”. *Journal of Natural History* 14.6 (1980): 759–768.

Jia, L.-Y., J.-H. Xiao, L.-M. Niu, G.-C. Ma, Y.-G. Fu, D. W. Dunn, and D.-W. Huang. “Delimitation and description of the immature stages of a pollinating fig wasp, Ceratosolen solmsi marchali Mayr (Hymenoptera: Agaonidae)”. *Bulletin of Entomological Research* 104.2 (2014): 164–175.

Jiang, G. and Z. Zheng. “Morphological descriptions of the eggs of five species of Chinese catantopidae”. *Zoological Research* 15.1 (1993): 29–32.

Jintsu, Y., T. Uchifune, and R. Machida. “Egg Membranes of a Web-spinner, Aposthonia japonica (Okajima ) (Insecta: Embioptera)”. *Proceedings of Arthropodan Embryological Society of Japan* 42 (2007): 1–5.

Jintsu, Y., T. Uchifune, and R. Machida. “Structural features of eggs of the basal phasmatodean Timema monikensis Vickery and Sandoval, 1998 (Insecta: Phasmatodea: Timematidae)”. *Arthropod Systematics and Phylogeny* 68.1 (2010): 71–78.

Johannsen, O. A. “Some phases in the embryonic development of Diacrisia virginica Fabr. (Lepidoptera)”. *Journal of Morphology* 48.2 (1929): 493–541.

Johnson, C. D., J. Romero, and E. Raimúndez-Urrutia. “Ecology of Amblycerus crassipunctatus Ribeiro-Costa (Coleoptera: Bruchidae) in seeds of Humiriaceae, a new host family for bruchids, with an ecological comparision to other species of Amblycerus”. *The Coleopterists Bulletin* 55.1 (2001): 37–48.

Johnson, J. B., T. D. Miller, J. M. Heraty, and F. W. Merickel. “Observations on the biology of two species of Orasema (Hymenoptera: Eucharitidae)”. *Proceedings of the Entomological Society of Washington* 88.3 (1986): 542–549.

Johnson, L. K. “Reproductive behavior of Claeoderes bivittata (Coleoptera: Brentidae)”. *Psyche* 90.1-2 (1983): 135–150.

Johnson, N. E. “Contarinia washingtonensis (Diptera: Cecidomyiidae), New Species Infesting the Cones of Douglas-Fir.” *Annals of the Entomological Society of America* 56.1 (1963): 94–103.

José, L., S. Carrillo, and L. E. Caltagirone. "Observations on the Biology of Solierella peckhami, S. blaisdelli (Sphecidae), and two species of Chrysididae (Hymenoptera)". *Annals of the Entomological Society of America* 63.3 (1970): 672–681.

Juliano, S. A. "The effects of body size on mating and reproduction in Brachinus lateralis (Coleoptera: Carabidae)". *Ecological Entomology* 10.3 (1985): 271–280.

Jung, E. "Untersuchungen am Ei des Speisebohnenkäfers Bruchidius obtectus Say (Coleoptera)". *Development Genes and Evolution* 157.4 (1966): 320–392.

Junkum, A., A. Jitpakdi, N. Komalamisra, N. Jariyapan, P. Somboon, P. A. Bates, and W. Choochote. "Comparative morphometry and morphology of Anopheles aconitus Form B and C eggs under scanning electron microscope". *Revista do Instituto de Medicina Tropical de São Paulo* 46.5 (2004): 257–262.

Jupeng, L. and X. Ruihua. "Studies of eggs of Chinese acridoids: Morphological descriptions of the eggs of acridoids from Changbai mountains". *Sinozoologia* 5.5 (1987): 41–45.

Jura, C., A. Krzysztofowicz, and E. Kisiel. "Embryonic development of Tetrodontophora bielanensis (Collembola): descriptive, with scanning electron micrographs". *Recent Advances in Insect Embryology in Japan and Poland*. Tsukuba: The Arthropodan Embryological Society of Japan, 1987. 77–124.

Kádár, F., P. J. Fazekas, M. Sárospataki, and G. L. Lövei. "Seasonal dynamics, age structure and reproduction of four Carabus species (Coleoptera: Carabidae) living in forested landscapes in Hungary". *Acta Zoologica Academiae Scientiarum Hungaricae* 61.1 (2015): 57–72.

Kadowaki, K., R. A. B. Leschen, and J. R. Beggs. "Spore consumption and life history of Zearagytodes maculifer (Broun) (Coleoptera: Leiodidae) on Ganoderma, its fungal host". *New Zealand Journal of Ecology* 35.1 (2011): 61–68.

Kaib, M., M. Hacker, and R. Brandl. "Egg-laying in monogynous and polygynous colonies of the termite Macrotermes michaelseni (Isoptera, Macrotermitidae)". *Insectes Sociaux* 48.3 (2001): 231–237.

Kaiser, J. and D. F. Went. "Early embryonic development of the dipteran insect Heteropeza pygmaea in the presence of cytoskeleton-affecting drugs". *Development Genes and Evolution* 196.6 (1987): 356–366.

Kalender, Z. S. S. C. Y. "Chorionic sculpturing in eggs of six species of Eurydema (Heteroptera, Pentatomidae): A scanning electron microscope investigation". *Journal of the Entomological Research Society* 1.2 (1999): 27–56.

Kambysellis, M. P. and W. B. Heed. "Studies of oogenesis in natural populations of Drosophilidae. I. Relation of ovarian development and ecological habitats of the Hawaiian species". *American Naturalist* 105.941 (1971): 31–49.

Kaminski, L. A. and F. S. Carvalho-Filho. "Life History of Aricoris propitia (Lepidoptera: Riodinidae) A Myrmecophilous Butterfly Obligately Associated with Fire Ants". *Psyche* 2012 (2012): 1–10.

Kaminski, L. A., D. Rodrigues, and A. V. L. Freitas. "Immature stages of Parrhasius polibetes (Lepidoptera: Lycaenidae): host plants, tending ants, natural enemies and morphology". *Journal of Natural History* 46.11–12 (2012): 645–667.

Kaminski, L. A., M. Tavares, V. G. Ferro, and G. R. P. Moreira. "Morfologia externa dos estágios imaturos de heliconíneos neotropicais. 111. Heliconius erato phyllis (Fabricius) (Lepidoptera, Nymphalidae, Heliconiinae)". *Revista Brasileira de Zoologia* 19.4 (2002): 977–993.

Kamiya, A. and H. Ando. "External morphogenesis of the embryo of Ascalaphus ramburi (Neuroptera, Ascalaphidae)". *Recent Advances in Insect Embryology in Japan*. ISEBU Co.: Tsukuba Science City, 1985. 203–214.

Kan, E. and Y. Waku. "Analysis of oviposition preference in the webbing clothes moth, Tineola bisselliella Hum. (Lepidoptera: Tineidae)". *Applied Entomology and Zoology* 20.3 (1985): 322–330.

Kaoru, S. and M. Ryuichiro. "Embryonic Development of Occasjapyx japonicus (Enderlein): Notable Features (Hexapoda: Diplura, Dicellurata)". *Proceedings of Arthropodan Embryological Society of Japan* 44 (2009): 13–18.

Kasule, F. K. "Egg size increases with maternal age in the cotton stainer bugs Dysdercus fasciatus and D. cardinalis (Hemiptera: Pyrrhocoridae)". *Ecological Entomology* 16.3 (1991): 345–349.

Katiyar, K. N. "Ecology of oviposition and the structure of egg-pods and eggs in some Indian Acrididae". *Records of the Indian Museum* 55 (1957): 29–68.

Katzav-Gozansky, T., V. Soroker, J. Kamer, C. M. Schulz, W. Francke, and A. Hefetz. "Ultrastructural and chemical characterization of egg surface of honeybee worker and queen-laid eggs". *Chemoecology* 13.3 (2003): 129–134.

Kaufmann, T. and P. Stansly. "Bionomics of Neoheterocerus pallidus Say (Coleoptera: Heteroceridae) in Oklahoma". *Journal of the Kansas Entomological Society* 52.3 (1979): 565–577.

Kaupp, A., R. Guggenheim, and P. Nagel. "The chorion as a subject of phylogenetic research in Paussinae and other Carabidae (Coleoptera: Adephaga)". *Entomologica Basiliensia* 22 (2000): 149–154.

Kaupp, A., R. Guggenheim, and P. Nagel. "Egg-shell structure of Paussinae and other Carabidae, with notes on its phylogenetic relevance (Coleoptera)". *Natural History and Applied Ecology of Carabid Beetles, Proceedings of the 9th International Carabidologist Meeting, Cosenza, Italy*. Moscow: Pensoft Publishers, 2000. 111–115.

Kawaguchi, Y., Y. Banno, K. Koga, T. Kawarabata, and H. Doira. "Surface ultrastructure of the eggs of Malacopsylla grossiventris and Phthiropsylla agenoris (Siphonaptera: Malacopsyllidae)". *Applied Entomology and Zoology* 31.3 (1996): 407–415.

Kawaguchi, Y., M. Ichida, T. Kusakabe, and K. Koga. "Chorion morphology of the Eri-silkworm, Samia cynthia ricini (Donovan) (Lepidoptera: Saturniidae)". *Applied Entomology and Zoology* 35.4 (2000): 427–434.

Kawakami, T. "Development of the immature stages of Ascogaster reticulatus Watanabe (Hymenoptera: Braconidae), an egg-larval parasitoid of the smaller tea tortrix moth, Adoxophyes sp. (Lepidoptera: Tortricidae)". *Applied Entomology and Zoology* 20.4 (1985): 380–386.

Kawanishi, C. Y. "Embryonic development of the drywood termite, Cryptotermes brevis". *Technical Bulletin of the Hawaii Agricultural Experiment Station, University of Hawaii* 95 (1975): 1–36.

Keffer, S. L., S. J. Taylor, and J. E. McPherson. "Laboratory rearing and descriptions of immature stages of Curicta scorpio (Heteroptera: Nepidae)". *Annals of the Entomological Society of America* 87.1 (1994): 17–26.

Keiper, J. B. "Biology and immature stages of coexisting Hydroptilidae (Trichoptera) from Northeastern Ohio lakes". *Annals of the Entomological Society of America* 95.5 (2002): 608–616.

Keiper, J. B. and B. A. Foote. "Biology and larval feeding habits of coexisting Hydroptilidae (Trichoptera) from a small woodland stream in northeastern Ohio". *Annals of the Entomological Society of America* 93.2 (2000): 225–234.

Kekeunou, S., M. V. Anyeng, E. Konyal, B. Bapfubusa, C. F. B. Bilong, et al. "Morphology, Development and Reproduction of Zonocerus variegatus (L.) (Pyrgomorphidae) Feeding on Vernonia amygdalina (Asteraceae) and Manihot esculenta (Euphorbiaceae) in the Laboratory". *Pakistan Journal of Zoology* 46.6 (2014): 1529–1536.

Kelly, G. M. and E. Huebner. "Embryonic development of the hemipteran insect Rhodnius prolixus". *Journal of Morphology* 199.2 (1989): 175–196.

Kennedy, T. F., G. O. Evans, and A. M. Feeney. "Studies on the biology of Tachyporus hypnorum F. (Col. Staphylinidae), associated with cereal fields in Ireland". *Irish Journal of Agricultural Research* 25.1 (1986): 81–95.

Kershaw, J. C. "The Formation of the Ootheca of a Chinese Mantis, Hierodula Saussurii". *Psyche* 17.4 (1900): 136–141.

Kessel, E. L. "The embryology of fleas". *Smithsonian Miscellaneous Collections* 98.3 (1939): 1–78.

Kim, D. S. and J. E. Lee. "Immature Stages of Nephrotoma virgata (Diptera, Tipulidae) from Korea". *Korean Journal of Applied Entomology* 44.1 (2005): 1–4.

Kim, D. S. and J. E. Lee. "Immature stages of Tipula patagiata (Diptera, Tipulidae) from Korea". *Korean Journal of Applied Entomology* 43.4 (2004): 263–266.

Kim, D.-S. and J.-E. Lee. "Immature stages of Tipula (Yamatotipula) latemarginata (Diptera, Tipulidae) from Korea". *Animal Systematics, Evolution and Diversity* 18.2 (2002): 213–217.

Kim, I.-K., J.-D. Park, S.-C. Shin, and I.-K. Park. "Prolonged embryonic stage and synchronized life-history of Platygaster robiniae (Hymenoptera: Platygastridae), a parasitoid of Obolodiplosis robiniae (Diptera: Cecidomyiidae)". *Biological Control* 57.1 (2011): 24–30.

Kirk, W. D. J. "Egg-hatching in thrips (Insecta: Thysanoptera)". *Journal of Zoology* 207.2 (1985): 181–190.

Kirollos, J. Y. and E. T. Hibbs. “Viability of Empoasca fabae (Homoptera: Cicadellidae) eggs in sterile media with added monosaccharides, amino acids, or phorate”. *Annals of the Entomological Society of America* 64.1 (1971): 32–36.

Kishimoto, T. and H. Ando. “External features of the developing embryo of the stonefly, Kamimuria tibialis (Pictet) (Plecoptera, Perlidae)”. *Journal of Morphology* 183.3 (1985): 311–326.

Kitching, R. L. “The immature stages of Sextius virescens (Fairmaire) (Homopter A: Membracidae)”. *Australian Journal of Entomology* 13.1 (1974): 55–60.

Klass, K.-D., O. Zompro, N. P. Kristensen, and J. Adis. “Mantophasmatodea: a new insect order with extant members in the Afrotropics”. *Science* 296.5572 (2002): 1456–1459.

Klonowska-Olejnik, M., T. Jazdzewska, and E. Gaino. “Scanning electron microscopy study of the eggs of some rare mayfly (Ephemeroptera) species: Ametropus fragilis, Isonychia ignota and Neoephemera maxima”. *Research Update on Ephemeroptera & Plecoptera*. Perugia: University of Perugia, 2003. 147–462.

Klots, A. B. “Life History Notes on Lagoa laceyi (Barnes & McDunnough) (Lepidoptera: Megalopygidae)”. *Journal of the New York Entomological Society* 74.3 (1966): 140–142.

Knabke, J. J. and A. A. Grigarick. “Biology of the African earwig, Euborellia cincticollis (Gerstaecker) in California and comparative notes on Euborellia annulipes (Lucas)”. *Hilgardia* 41.7 (1971): 157–194.

Knight, A. W., A. V. Nebeker, and A. R. Gaufin. “Description of the eggs of common Plecoptera of Western United States”. *Entomological News* 76.4 (1965): 105–111.

Knutson, L. V. “Biology and immature stages of malacophagous flies: Antichaeta analis, A. atriseta, A. brevipennis, and A. obliviosa (Diptera: Sciomyzidae)”. *Transactions of the American Entomological Society (1890-)* 92.1 (1966): 67–101.

Kobayashi, S., R. Usui, K. Nomoto, M. Ushirokita, T. Denda, and M. Izawa. “Does egg dispersal occur via the ocean in the stick insect Megacrania tsudai (Phasmida: Phasmatidae)?” *Ecological Research* 29.6 (2014): 1025–1032.

Kobayashi, T. “Developmental stages of Brachynema and its allied genus of Japan (Pentatomidae): The developmental stages of some species of the Japanese Pentatomoidea XV”. *Kontyû* 33.3 (1965): 304–309.

Kobayashi, T. “Developmental stages of Glaucias and its allied genera of Japan (Hemiptera: Pentatomidae) (The developmental stages of some species of the Japanese Pentatomoidea, XVIII)”. *Transactions of the Shikoku Entomological Society* 20.3-4 (1994): 197–205.

Kobayashi, T. “Developmental stages of Urochela and an allied genus of Japan (Hemiptera: Urostylidae)”. *Transactions of the Shikoku Entomological Society* 8.3 (1965): 94–104.

Kobayashi, T. “The development stages of some species of the Japanese Pentatomoidea (Hemiptera): VII. Developmental stages of Nezara and its applied genera (Pentatomoidae s. st.)” *Japanese Journal of Applied Entomology and Zoology* 3 (1959): 221–231.

Kobayashi, T. “The developmental stages of six species of Japanese Pentatomoidea (Hemiptera)”. *Scientific Reports of Matsuyama Agricultural College* 11 (1953): 73–79.

Kobayashi, T. “The developmental stages of some species of the Japanese Pentatomoidea (Hemiptera), III”. *Transactions of the Shikoku Entomological Society* 4.4 (1954): 63–68.

Kobayashi, T. “The developmental stages of some species of the Japanese Pentatomoidea (Hemiptera), IV”. *Transactions of the Shikoku Entomological Society* 4.5-6 (1955): 79–82.

Kobayashi, T. “The developmental stages of some species of the Japanese Pentatomoidea (Hemiptera), V”. *Transactions of the Shikoku Entomological Society* 4 (1956): 120–130.

Kobayashi, T. “The developmental stages of some species of the Japanese Pentatomoidea (Hemiptera): XVI. Homalogonia and an allied genus of Japan (Pentatomidae)”. *Japanese Society of Applied Entomology and Zoology* 2.1 (1967): 1–8.

Kobayashi, T. “The developmental stages of some species of the Japanese Pentatomoidea (Hemiptera). IX. Developmental stages of Lagynotomus, Aelia, and their allied genera (Pentatomidae s. str.)” *Japanese Journal of Applied Entomology and Zoology* 3 (1960): 11–19.

Kobayashi, T. “The developmental stages of some species of the Japanese Pentatomoidea (Hemiptera). X. Developmental stages of Eysarcoris and its allied genera”. *Japanese Journal of Applied Entomology and Zoology* 4 (1960): 83–95.

Kobayashi, T. “The developmental stages of some species of the Japanese Pentatomoidea (Hemiptera). XI. Developmental stages of Scotinophara (Pentatomidae)”. *Japanese Journal of Applied Entomology and Zoology* 7.1 (1963): 70–78.

Kobayashi, T. “The developmental stages of some species of the Japanese Pentatomoidea (Hemiptera). XIV. Developmental stages of Graphosoma and its allied genera of Japan (Pentatomidae)”. *Japanese Journal of Applied Entomology and Zoology* 9.1 (1965): 34–41.

Kobayashi, Y. and H. Ando. “Phylogenetic relationships among the lepidopteran and trichopteran suborders (Insecta) from the embryological standpoint”. *Journal of Zoological Systematics and Evolutionary Research* 26.3 (1988): 186–210.

Kobayashi, Y. and G. W. Gibbs. “The Early Embryonic-Development of the Mnesarchaeid Moth, Mnesarchaea-Fusilella Walker (Lepidoptera, Mnesarchaeidae), and Its Phylogenetic Significance”. *Australian Journal of Zoology* 43.5 (1995): 479–488.

Kobayashi, Y. and K. Miya. “Structure of egg cortex relating to presumptive embryonic and extraembryonic regions in silkworm, Bombyx mori (Bombycidae: Lepidoptera)”. *Recent Advances in Insect Embryology in Japan and Poland*. Tsukuba: The Arthropodan Embryological Society of Japan, 1987. 181–194.

Kobayashi, Y. “Embryogenesis of the fairy moth, Nemophora albiantennella Issiki (Lepidoptera, Adelidae), with special emphasis on its phylogenetic implications”. *International Journal of Insect Morphology and Embryology* 27.3 (1998): 157–166.

Kobayashi, Y. and H. Ando. “Early embryonic development and external features of developing embryos of the caddisfly, Nemotaulius admorsus (Trichoptera: Limnephilidae)”. *Journal of Morphology* 203.1 (1990): 69–85.

Kobayashi, Y. and H. Ando. “The embryonic development of the primitive moth, Neomicropteryx nipponensis Issiki (Lepidoptera, Micropterygidae): morphogenesis of the embryo by external observation”. *Journal of Morphology* 169.1 (1981): 49–59.

Kobayashi, Y., K. Niikura, Y. Oosawa, and Y. Takami. “Embryonic development of Carabus insulicola (Insecta, Coleoptera, Carabidae) with special reference to external morphology and tangible evidence for the subcoxal theory”. *Journal of Morphology* 274.12 (2013): 1323–1352.

Kobayashi, Y., H. Suzuki, and N. Ohba. “Embryogenesis of the glowworm Rhagophthalmus ohbai Wittmer (Insecta: Coleoptera, Rhagophthalmidae), with emphasis on the germ rudiment formation”. *Journal of Morphology* 253.1 (2002): 1–9.

Kocarek, P. “A case of viviparity in a tropical non-parasitizing earwig (Dermaptera Spongiphoridae)”. *Tropical Zoology* 22 (2009): 237–241.

Koedam, D., P. H. Velthausz, M. R. Dohmen, and M. J. Sommeijer. “Morphology of reproductive and trophic eggs and their controlled release by workers in Trigona (Tetragonisca) angustula llliger (Apidae, Meliponinae)”. *Physiological Entomology* 21.4 (1996): 289–296.

Kojima, J. “Immatures of hover wasps (Hymenoptera, Vespidae, Stenogastrinae)”. *Kontyû* 58.3 (1990): 506–522.

Komatsu, S. and Y. Kobayashi. “Embryonic development of a whirligig beetle, Dineutus mellyi, with special reference to external morphology (insecta: Coleoptera, Gyrinidae)”. *Journal of Morphology* 273.5 (2012): 541–560.

Konečná, H. and H. Šefrová. “Morphology, Biology and Control Possibilities of Two Argyresthia species – A. thuiella and A. trifasciata (Lepidoptera: Argyresthiidae)”. *Acta Universitatis Agriculturae et Silviculturae Mendelianae Brunensis* 62.3 (2014): 529–538.

Konopko, S. A. “Description of the Immature Stages of Sigara (Aphelosigara) tucma”. *Journal of Insect Science* 14.1 (2014): 1–9.

Konopko, S. A. “Description of the immature stages of Sigara (Tropocorixa) santiagiensis (Hungerford, 1928) (Insecta: Heteroptera: Corixidae)”. *Journal of Natural History* 47.29-30 (2013): 1959–1982.

Konopko, S. A. “Description of the immature stages of Sigara (Tropocorixa) schadei (Hungerford) (Hemiptera: Heteroptera: Corixidae)”. *Zootaxa* 3487 (2012): 41–57.

Konopko, S. A. “Immature stages of the genus Ectemnostega Enderlein (Hemiptera: Heteroptera: Corixidae), with an identification key to instars and redescription of the nymphs of E. (Ectemnostega) quadrata (Signoret)”. *Studies on Neotropical Fauna and Environment* 48.1 (2013): 40–55.

Konopko, S. A. and S. A. N. A. Mazzucconi. “The immature stages of the genus Trepobates Uhler (Hemiptera: Heteroptera: Gerridae), with an identification key to instars and the description of the nymphs of T. taylori (Kirkaldy)”. *Zootaxa* 2733 (2011): 1–15.

Konopko, S. A., S. A. N. A. Mazzucconi, and A. O. Bachmann. “Description of the immature stages of Trichocorixa mendozana Jaczewski (Hemiptera: Heteroptera: Corixidae)”. *Zootaxa* 3060 (2011): 47–61.

Konopko[1], S. A., S. A. N. A. Mazzucconi, and M. L. Ruf. “Studies on the chorionic structure of the eggs of Corixoidea (Hemiptera: Heteroptera) with scanning electron microscopy”. *Zootaxa* 3737.3 (2013): 223–240.

Koppenhöfer, A. M. “Observations on the bionomics of Thyreocephalus interocularis (Eppelsheim) (Col., Staphylinidae), a predator of the banana weevil”. *Journal of Applied Entomology* 117.1-5 (1994): 388–392.

Korboot, K. “Observations on the life histories of the stick insects Acrophylla tessellata Gray and Extatosoma tiaratum Macleay”. *University of Queensland Papers, Department of Entomology* 1.11 (1961): 161–169.

Kormondy, E. J. *The systematics of Tetragoneuria, based on ecological, life history, and morphological evidence (Odonata: Corduliidae)*. Ann Arbor: Museum of Zoology, University of Michigan, 1959.

Kornhauser, S. I. “The sexual characteristics of the membracid, Thelia bimaculata (Fabr.). I. External changes induced by Aphelopus theliae (Gahan)”. *Journal of Morphology* 32.3 (1919): 531–636.

Korycinska, A. “A description of the eggs of seven species of Noctuidae (Lepidoptera) commonly transported by plant trade to the UK, and their separation using stereomicroscopy and scanning electron microscopy”. *Tijdschrift voor Entomologie* 155.1 (2012): 15–28.

Koss, R. W. and G. F. Edmunds. “Ephemeroptera eggs and their contribution to phylogenetic studies of the order”. *Zoological Journal of the Linnean Society* 55.4 (1974): 267–349.

Kovarik, P. W. “Development of Epierus divisus Marseul (Coleoptera: Histeridae)”. *The Coleopterists' Bulletin* 49.3 (1995): 253–260.

Kozo, M. I. Y. “Description of the eggs of common Plecoptera of western United States”. *Recent Advances in Insect Embryology in Japan and Poland*. Tsukuba: The Arthropodan Embryological Society of Japan, 1987. 125–149.

Kraus, W. F., M. J. Gonzales, and S. L. Vehrencamp. “Egg development and an evaluation of some of the costs and benefits for paternal care in the belostomatid, Abedus indentatus (Heteroptera: Belostomatidae)”. *Journal of the Kansas Entomological Society* 62.4 (1989): 548–562.

Krause, J. B. and M. T. Ryan. “The stages of development in the embryology of the horned passalus beetle, Popilius disjunctus Illiger”. *Annals of the Entomological Society of America* 46.1 (1953): 1–20.

Krysan, J. L. “The early embryology of Diabrotica undecimpunctata howardi (Coleoptera: Chrysomelidae)”. *Journal of Morphology* 149.1 (1976): 121–137.

Kučerová, Z. “Stored product psocids (Psocoptera): External morphology of eggs”. *European Journal of Entomology* 99 (2002): 491–503.

Kučerová, Z., J. Hromádková, and V. Stejskal. “External egg morphology of common stored-product pests from the families Anobiidae (Ptininae) and Dermestidae (Coleoptera)”. *10th International Working Conference on Stored Product Protection*. Berlin: Julius Kühn-Institut, 2010. 135–138.

Kučerová, Z. and M. Jokeš. “External morphology of eggs of the synanthrophic psocid Dorypteryx domestica (Psocoptera, Psyllipsocidae)”. *Deutsche Entomologische Zeitschrift* 49.1 (2002): 165–169.

Kučerová, Z., Z.-H. Li, I. Kalinović, Q.-Q. Yang, J. Hromádková, and C. Lienhard. “The external morphology of females, males and eggs of a Liposcelis silvarum (Insecta: Psocodea: Liposcelididae) strain with unusually developed compound eyes, visualised with scanning electron microscopy”. *Italian Journal of Zoology* 79.3 (2012): 402–409.

Kučerová, Z. and V. Stejskal. “Differences in egg morphology of the stored-grain pests Rhyzopertha dominica and Prostephanus truncatus (Coleoptera: Bostrichidae)”. *Journal of Stored Products Research* 44.1 (2008): 103–105.
Kučerová, Z. and V. Stejskal. “Comparative egg morphology of silvanid and laemophloeid beetles (Coleoptera) occurring in stored products”. *Journal of Stored Products Research* 38.3 (2002): 219–227.
Kučerová, Z. and V. Stejskal. “External egg morphology of two stored-product anobiids, Stegobium paniceum and Lasioderma serricorne (Coleoptera: Anobiidae)”. *Journal of Stored Products Research* 46.3 (2010): 202–205.
Kula, E. “Sculpture of exochorion in some eggs of Syrphidae (Diptera) (Pt. 2)”. *Acta Universitatis Agriculturae et Silviculturae Mendelianae Brunensis* 58.1-4 (1989): 99–126.
Kumar, A., V. C. Kapoor, and P. Laska. “Immature stages of some aphidophagous syrphid flies of India (Insecta, Diptera, Syrphidae)”. *Zoologica Scripta* 16.1 (1987): 83–88.
Kumar, R. “Studies on the biology, immature stages and relative groowth of some Australian bugs of the superfamily Coreoidea (Hemiptera: Heteroptera)”. *Australian Journal of Zoology* 14.5 (1966): 895–991.
Kumar, R. and V. V. Ramamurthy. “Morphology and bionomics of Phycodes radiata Ochsenheimer (Lepidoptera: Brachodidae) from New Delhi, India”. *Tijdschrift voor Entomologie* 153.1 (2010): 15–24.
Kumar, R., A. Kumar, and E. Mey. “Egg microtopography of chicken bug louse, Stenocrotaphus gigas (Taschenberg) (Phthiraptera: Ischnocera, Goniodidae)”. *Rudolstädter Naturhistorische Schriften* 16 (2010): 111–115.
Kumar, V. and C. K. Kamble. “Scanning electron microscope study on the egg chorion of silkmoth, Antheraea assamensis Helf. (Lepidoptera: Saturniidae)”. *Animal Biology* 58.2 (2008): 235–244.
Kumar, V., B. K. Kariappa, A. M. Badu, and S. B. Dandin. “Surface ultrastructure of the egg chorion of eri silkworm, Samia ricini (Donovan) (Lepidopotera: Saturniidae)”. *Journal of Entomology* 4.2 (2007): 68–81.
Kumar, V., M. N. Morrison, A. M. Babu, and V. Thiagarajan. “Egg shell architecture of the stink bug, Eocanthecona furcellata (Wolff.): Ultrastructure of micropylar processes and egg burster”. *International Journal of Tropical Insect Science* 22.1 (2002): 67–73.
Kumar, V., S. Rajadurai, A. M. Babu, and B. K. Kariappa. “Eggshell Fine Structure of Amata passalis F.(Lepidoptera: Amatidae), a Pest of Mulberry”. *International Journal of Tropical Insect Science* 23.4 (2003): 325–330.
Kumbhar, S. M., A. B. Mamlayya, S. J. Patil, and G. P. Bhawane. “Biology of Chiloloba orientalis”. *Journal of Insect Science* 12.1 (2012): 1–15.
Kuniata, L. S. and G. R. Young. “The biology of Lepidiota reuleauxi Brenske (Coleoptera: Scarabaeidae), a pest of sugarcane in Papua New Guinea”. *Australian Journal of Entomology* 31.4 (1992): 339–343.
Kurczewski, F. E. “A review of nesting behavior in the Tachysphex pompiliformis group, with observations on five species (Hymenoptera: Sphecidae)”. *Journal of the Kansas Entomological Society* 60.1 (1987): 118–126.
Kurczewski, F. E. “Behavioral Observations on Some Tachytini and Larrini (Hymenoptera: Sphecidae)”. *Journal of the Kansas Entomological Society* 49.3 (1976): 327–332.
Kurczewski, F. E. “Comparative nesting behavior of Episyron quinquenotatus (Hymenoptera: Pompilidae) in the northeastern United States”. *Northeastern Naturalist* 8.4 (2001): 403–426.
Kurczewski, F. E. “Observations on the nesting behavior of Auplopus caerulescens subcorticalis and other Auplopodini (Hymenoptera: Pompilidae)”. *Great Lakes Entomologist* 22.2 (1989): 71–74.
Kurczewski, F. E. “Observations on the nesting behaviors of spider-wasps in southern Florida (Hymenoptera: Pompilidae)”. *The Florida Entomologist* 64.3 (1981): 424–437.
Kurczewski, F. E. and R. E. Acciavatti. “A review of the nesting behaviors of the nearctic species of Crabro, including observations on C. advenus and C. latipes (Hymenoptera: Sphecidae)”. *Journal of the New York Entomological Society* 76.3 (1968): 196–212.
Kurczewski, F. E. and N. B. Elliott. “Nesting behavior and ecology of Tachysphex pechumani Krombein (Hymenoptera: Sphecidae)”. *Journal of the Kansas Entomological Society* 51.4 (1978): 765–780.
Kurczewski, F. E. and E. J. Kurczewski. “Host records for some North American Pompilidae (Hymenoptera). Third Supplement. Tribe Pompilini”. *Journal of the Kansas Entomological Society* 46.1 (1973): 65–81.
Kurczewski, F. E. and R. C. Miller. “Observations on the nesting of three species of Cerceris (Hymenoptera: Sphecidae)”. *The Florida Entomologist* 67.1 (1984): 146–155.

Kurczewski, F. E. and M. G. Spofford. "Observations on the behaviors of some Scoliidae and Pompilidae (Hymenoptera) in Florida". *The Florida Entomologist* 69.4 (1986): 636–644.

Küthe, H.-W. "Das Differenzierungszentrum als selbstregulierendes Faktorensystem für den Aufbau der Keimanlage im Ei von Dermestes frischi (Coleoptera)". *Wilhelm Roux'Archiv für Entwicklungsmechanik der Organismen* 157.3 (1966): 212–302.

Kyneb, A. and S. Toft. "Quality of two aphid species (Rhopalosiphum padi and Sitobion avenae) as food for the generalist predator Tachyporus hypnorum (Col., Staphylinidae)". *Journal of Applied Entomology* 128.9-10 (2004): 658–663.

Lachaise, D., L. Tsacas, and G. Couturier. "The Drosophilidae associated with tropical African figs". *Evolution* 36.1 (1982): 141–151.

Lachmann, A. D. "Sexual receptivity and post-emergence ovarian development in females of Coproica vagans (Diptera: Sphaeroceridae)". *Physiological Entomology* 23.4 (1998): 360–368.

Laing, D. R. and L. E. Caltagirone. "Biology of Habrobracon lineatellae (Hymenoptera: Braconidae)". *The Canadian Entomologist* 101.2 (1969): 135–142.

Lal, K. B. "The biology of Scottish Psyllidae". *Transactions of the Royal Entomological Society of London* 82.2 (1934): 363–385.

Lal, K. "Some aspects of the embryology of Tetrastichus pyrillae Craw. (Eulophidae: Hymenoptera)". *Proceedings of the Indian Academy of Sciences - Section B* 64.1 (1966): 38–44.

Lalonde, R. G. "Egg size variation does not affect offspring performance under intraspecific competition in Nasonia vitripennis, a gregarious parasitoid". *Journal of Animal Ecology* 74.4 (2005): 630–635.

Lamb, R. J. and S. M. Smith. "Comparison of egg size and related life-history characteristics for two predaceous tree-hole mosquitoes (Toxorhynchites)". *Canadian Journal of Zoology* 58.11 (1980): 2065–2070.

Landis, D. A., C. E. Sorenson, and E. D. Cashatt. "Biology of Clydonopteron sacculana (Lepidoptera: Pyralidae) in North Carolina, with description of the egg stage". *Annals of the Entomological Society of America* 85.5 (1992): 596–604.

Lansbury, I. "A revision of the genus Paranisops Hale (Heteroptera: Notonectidae)". *Proceedings of the Royal Entomological Society of London. Series B, Taxonomy* 33.11-12 (1964): 181–188.

Lantsov, V. I. "The ecology, biology and larval instars of the North Caucasian population (Lake Maliy Tambukan) of Tipula subcunctans Alexander, 1921 (Diptera: Tipulidae)". *Zoosymposia* 3.1 (2009): 115–129.

Larink, O. "Zur Entwicklungsgeschichte von Petrobius brevistylis (Thysanura, Insecta)". *Helgoländer Wissenschaftliche Meeresuntersuchungen* 19.1 (1969): 111–155.

Larink, O. and S. M. Biliński. "Fine structure of the egg envelopes of one proturan and two collembolan genera (Apterygota)". *International Journal of Insect Morphology and Embryology* 18.1 (1989): 39–45.

Lassmann, G. W. P. "The early embryological development of Melophagus ovinus L., with special reference to the development of the germ cells". *Annals of the Entomological Society of America* 29.3 (1936): 397–414.

Laudonia, S. and G. Viggiani. "Observations on the developmental stages of Edovum puttleri Grissell (Hymenoptera: Eulophidae), an egg-parasitoid of Colorado potato beetle". *Bollettino del Laboratorio di Entomologia Agraria Filippo Silvestri Portici* 43 (1986): 97–103.

Laudonia, S. and G. Viggiani. "Osservazioni sugli stadi preimmaginali di Cales noacki Howard (Hymenoptera: Aphelinidae)". *Bollettino del Laboratorio di Entomologia Agraria Filippo Silvestri Portici* 43 (1986): 22–28.

Lauterer, P. "Notes on the distribution and egg shape of several European psyllid species (Homoptera, Psylloidea)". *Acta Musei Moraviae* 82 (1998): 157–161.

Lavigne, R. J. "Notes on the distribution and ethology of Efferia bicaudata (Diptera: Asilidae), with a description of the eggs". *Annals of the Entomological Society of America* 57.3 (1964): 341–344.

Lavigne, R. J. and S. W. Bullington. "Ethology of Laphria fernaldi (Back) (Diptera: Asilidae) in southeast Wyoming". *Proceedings of the Entomological Society of Washington* 86.2 (1984): 326–336.

Lavigne, R. J. and D. S. Dennis. "Ethology of Efferia frewingi (Diptera: Asilidae)". *Annals of the Entomological Society of America* 68.6 (1975): 992–996.

Lavigne, R. J., D. S. Dennis, et al. "Ethology of Proctacanthella leucopogon in Mexico (Diptera: Asilidae)". *Proceedings of the Entomological Society of Washington* 82.2 (1980): 260–268.

Lawson, F. A. "Egg and larval case formation by Pachybrachis bivittatus". *Annals of the Entomological Society of America* 69.5 (1976): 942–944.

Lawson, F. A. "Structural Features of Cochroach Egg Capsules IV. The Oötheca of Parcoblatta Uhleriana.(Orthoptera: Blattidae)". *Journal of the Kansas Entomological Society* 27.1 (1954): 14–20.

Lawson, F. A. "Structural Features of Cockroach Egg Capsules: II. The Ootheca of Cariblatta Lutea Lutea (Orthoptera: Blattidae)". *Journal of the Kansas Entomological Society* 52.5 (1954): 296–300.

Lawson, F. A. "Structural features of cockroach egg capsules. V. The ootheca of Lamproblatta albipalpus Hebard (Orthoptera: Blattidae)". *Journal of the Kansas Entomological Society* 40.4 (1967): 601–607.

Lawson, F. A. "Structural features of cockroach egg capsules. VII. The ootheca of Cariblatta plagia Rehn & Hebard (Orthoptera: Blattidae)". *Journal of the Kansas Entomological Society* 48.2 (1975): 169–174.

LeBlanc, D. A. and C. R. Lacroix. "Developmental potential of galls induced by Diplolepis rosaefolii (Hymenoptera: Cynipidae) on the leaves of Rosa virginiana and the influence of Periclistus species on the Diplolepis rosaefolii galls". *International Journal of Plant Sciences* 162.1 (2001): 29–46.

Lebouvier, M., G. Chauvin, and C. Hamon. "L'oeuf de Myrmeleotettix maculatus Thunb. (Orthoptera: Acrididae): absorption d'eau et structure fine des enveloppes". *International Journal of Insect Morphology and Embryology* 14.2 (1985): 91–103.

LeCato, G. L. and B. R. Flaherty. "Description of eggs of selected species of stored-product insects (Coleoptera and Lepidoptera)". *Journal of the Kansas Entomological Society* 47.3 (1974): 308–317.

Lee, C.-F., S. Hisamatsu, and P.-S. Yang. "Morphology and ontogeny of immature stages of Helotidae based on descriptions of Helota thoracica Ritsema and H. gemmata Gorham (Insecta: Coleoptera: Cucujoidea)". *Zoological Studies* 46.6 (2007): 760–769.

Lee, G.-E., J. Hayden, and A. Y. Kawahara. "External egg morphology of the Hawaiian dancing moth, Dryadaula terpsichorella (Lepidoptera: Tineidae)". *Journal of Natural History* 48.15-16 (2014): 969–974.

Lee, J. E. "Immature stages of Pyrrhalta humeralis (Chen) and Galeruca vicina Solsky from Japan (Coleoptera, Chrysomelidae)". *Esakia* 1 (1990): 81–91.

Lee, J. E. "Morphological studies on the immature stages of two Japanese species of the genus Galerucella (Coleoptera, Chrysomelidae)". *Japanese Journal of Entomology* 58.2 (1990): 425–439.

Lees, A. H. "Some observations on the egg of Psylla mali". *Annals of Applied Biology* 2.4 (1916): 251–257.

Lefkovitch, L. P. and J. E. Currie. "Some morphological, biological and genetical differences between Cryptolestes pusillus fuscusssp. n. and C. pusillus pusillus (Schönherr) (Coleoptera, Cucujidae)". *Journal of Stored Products Research* 3.4 (1967): 311–320.

Leiby, R. W. and C. C. Hill. "The polyembryonic development of Platygaster vernalis". *Journal of Agricultural Research* 28.8 (1924): 829–839.

Leite, A. C. R. and P. Williams. "Morphological observations on the egg and first instar larva of Metacutereba apicalis (Diptera: Cuterebidae)". *Memórias do Instituto Oswaldo Cruz* 84.1 (1989): 123–130.

Leite, L. A. R., A. V. L. Freitas, E. P. Barbosa, M. M. Casagrande, and O. H. H. Mielke. "Immature stages of nine species of genus Dynamine Hübner, [1819]: morphology and natural history (Lepidoptera: Nymphalidae: Biblidinae)". *SHILAP Revista de Lepidopterología* 42.165 (2014): 27–55.

Leite, L. A. R., M. M. Casagrande, O. H. H. Mielke, and A. V. L. Freitas. "Immature stages of the Neotropical butterfly, Dynamine agacles agacles". *Journal of Insect Science* 12.37 (2012): 1–12.

Leite, L. A. R., F. M. S. Dias, E. Carneiro, M. M. Casagrande, and O. H. H. Mielke. "Immature stages of the Neotropical cracker butterfly, Hamadryas epinome". *Journal of Insect Science* 12.74 (2012): 1–12.

Leonardi, M. S., E. A. Crespo, J. A. Raga, and M. Fernández. "Scanning electron microscopy of Antarctophthirus microchir (Phthiraptera: Anoplura: Echinophthiriidae): Studying morphological adaptations to aquatic life". *Micron* 43.9 (2012): 929–936.

Leong, T. M. "Oviposition and hatching in the praying mantis, Hierodula patellifera (Serville) in Singapore (Mantodea: Mantidae: Paramantinae)". *Nature in Singapore* 2 (2009): 55–61.

Leong, T. M. and S. C. Teo. "Records of the praying mantis, Theopropus elegans (Westwood) (Mantodea: Hymenopodidae: Hymenopodinae) in Singapore, with notes on oviposition and hatching". *Nature in Singapore* 1 (2008): 211–214.

Leprince, D. J. and L. D. Foil. "Relationships among body size, blood meal size, egg volume, and egg production of Tabanus fuscicostatus (Diptera: Tabanidae)". *Journal of Medical Entomology* 30.5 (1993): 865–871.

LeSage, L., V. L. Stiefel, P. H. A. Jolivet, and M. L. Cox. "Biology and immature stages of the North American clytrines Anomoea laticlavia (Forster) and A. flavokansiensis Moldenke". *Chrysomelidae Biology* 3 (1996): 217–238.

LeSage, L. "Egg, larva, and pupa of Lexiphanes saponatus (Coleoptera: Chrysomelidae: Cryptocephalinae)". *The Canadian Entomologist* 116.4 (1984): 537–548.

LeSage, L. "Immature stages of Canadian Neochlamisus Karren (Coleoptera: Chrysomelidae)". *The Canadian Entomologist* 116.3 (1984): 383–409.

LeSage, L. "The eggs and larvae of Cryptocephalus quadruplex Newman and C. venustus Fabricius, with a key to the known immature stages of the Nearctic genera of Cryptocephaline leaf beetles (Coleoptera: Chrysomelidae)". *The Canadian Entomologist* 118.2 (1986): 97–111.

LeSage, L. "The eggs and larvae of Pachybrachis peccans and P. bivittatus, with a key to the known immature stages of the Nearctic genera of Cryptocephalinae (Coleoptera: Chrysomelidae)". *The Canadian Entomologist* 117.2 (1985): 203–220.

LeSage, L. "The immature stages of Exema canadensis Pierce (Coleoptera: Chrysomelidae)". *The Coleopterists' Bulletin* 36.2 (1982): 318–327.

Leschen, R. A. B. and C. E. Carlton. "Immature stages of Endomychus biguttatus Say (Coleoptera: Endomychidae) with observations on the alimentary canal". *Journal of the Kansas Entomological Society* 61.3 (1988): 321–327.

Leschen, R. A. B. and R. T. Allen. "Immature stages, life histories and feeding mechanisms of three Oxyporus spp. (Coleoptera: Staphylinidae: Oxyporinae)". *The Coleopterists' Bulletin* 42.4 (1988): 321–333.

Leston, D. "Notes on the Ethiopian Pentatomoidea (Hem.): XVIII, the eggs of three Nigerian shielbugs, with a tentative summary of egg forms in Pentatomoidea". *Entomologist's Monthly Magazine* 91 (1955): 33–36.

Łętowski, J., K. Pawlęga, R. Ścibior, and K. Rojek. "The morphology of the preimaginal stages of Squamapion elongatum (Germar, 1817) (Coleoptera, Curculionoidea, Apionidae) and notes on its biology". *ZooKeys* 519 (2015): 101–115.

Lévesque, C., J. G. Pilon, and J. Dubé. "Observations sur les oocytes de quelques Coleopteres Carabidae du Quebec". *Annals of the Entomological Society of Quebec* 25 (1980): 3–9.

Lewandowski, M., A. Sznyk, and A. Bednarek. "Biology and morphometry of Lycoriella ingenua (Diptera: Sciaridae)". *Biological Letters* 41.1 (2004): 41–50.

Lewis, W. J. "Life history and anatomy of Microplitis croceipes (Hymenoptera: Braconidae), a parasite of Heliothis spp. (Lepidoptera: Noctuidae)". *Annals of the Entomological Society of America* 63.1 (1970): 67–70.

Lewis, W. J. and S. B. Vinson. "Egg and larval development of Cardiochiles nigriceps". *Annals of the Entomological Society of America* 61.3 (1968): 561–565.

Liang, A.-P. "A new genus of Tropiduchidae (Hemiptera: Fulgoroidea) from China and Vietnam, with description of eggs". *The Florida Entomologist* 86.3 (2003): 361–369.

Liang, A.-P. and G.-M. Jiang. "Two new species of Tambinia Stål (Hemiptera: Tropiduchidae) from China, Laos and Vietnam, with description of eggs". *Journal of the Kansas Entomological Society* 76.3 (2003): 509–517.

Liebherr, J. K. "The unity of characters: ecological and morphological specialisation in larvae of Hawaiian platynine Carabidae (Coleoptera)". *Invertebrate Systematics* 14.6 (2000): 931–940.

Liles, M. P. "A study of the life history of the forked fungus beetle, Bolitotherus cornutus (Panzer) (Coleoptera: Tenebrionidae)". *Ohio Journal of Science* 56.6 (1956): 329–337.

Lima, A. R., A. F. Kumagai, and F. C. C. Neto. "Morphological and biological observations on the stick insect Tithonophasma tithonus (Gray, 1835) (Phasmida: Pseudophasmatidae: Pseudophasmatinae)". *Zootaxa* 3700.4 (2013): 588–592.

Lin, C.-S. "Immature stages of four Bombycidae species of Taiwan". *Collection and Research* 18 (2005): 25–31.

Linares, M. A., L. E. Neder, and C. Dietrich. "Description of immature stages and life cycle of the treehopper, Guayaquila projecta". *Journal of Insect Science* 10.199 (2010): 1–9.

Lincoln, D. C. R. "The oxygen and water requirements of the egg of Ocypus olens Müller (Staphylinidae, Coleoptera)". *Journal of Insect Physiology* 7.3 (1961): 265–272.

Linley, J. R., A. H. Benton, and J. F. Day. "Ultrastructure of the eggs of seven flea species (Siphonaptera)". *Journal of Medical Entomology* 31.6 (1994): 813–827.

Linley, J. R., L. P. Lounibos, and J. Conn. "A description and morphometric analysis of the eggs of four South American populations of Anopheles (Nyssorhynchus) aquasalis (Diptera: Culicidae)". *Mosquito Systematics* 25.3 (1993): 198–214.

Linley, J. R., L. P. Lounibos, J. Conn, D. Duzak, and N. Nishimura. "A description and morphometric comparison of eggs from eight geographic populations of the South American malaria vector Anopheles (Nyssorhynchus) nuneztovari (Diptera: Culicidae)". *Journal of the American Mosquito Control Association* 12.2 (1996): 275–292.

Linley, J. R. "Comparative fine structure of the eggs of Aedes albopictus, Ae. aegypti, and Ae. bahamensis (Diptera: Culicidae)". *Journal of Medical Entomology* 26.5 (1989): 510–521.

Linley, J. R. "Scanning electron microscopy of the egg of Aedes (Protomacleaya) triseriatus (Diptera: Culicidae)". *Journal of Medical Entomology* 26.5 (1989): 474–478.

Linley, J. R. and G. B. Craig Jr. "Morphology of long-and short-day eggs of Aedes atropalpus and A. epactius (Diptera: Culicidae)". *Journal of Medical Entomology* 31.6 (1994): 855–867.

Liu, J. P., R. H. Xi, W. B. Li, and Z. Z. Wang. *Illustrated handbook of Chinese acridoid eggs*. Beijing: Tianze Eldonejo, 1990.

Livingstone, D. "On the functional anatomy of the egg and the description of the nymphal instars of Dasytingis rudis Drake & Poor (Heteroptera: Tingidae), a sap sucker on Vitex negundo (Verbinaceae)". *Journal of Natural History* 10.5 (1976): 529–544.

Livingstone, D. "On the functional anatomy of the egg of Tingis buddleiae Drake (Het.: Tingidae)". *Journal of the Zoological Society of India* 19.1 & 2 (1967): 111–119.

Livingstone, D. and M. H. S. Yacoob. "Biosystematics of Tingidae on the basis of the biology and micromorphology of their eggs". *Proceedings of the Indian Academy of Sciences: Animal Sciences* 96.5 (1987): 587–611.

Llácer, E., A. Urbaneja, A. Garrido, and J.-A. Jacas. "Morphology and development of immature stages of Galeopsomyia fausta (Hymenoptera: Eulophidae: Tetrastichinae)". *Annals of the Entomological Society of America* 98.5 (2005): 747–753.

Llórente-Bousquets, J. and J. C. Gerardino. "Estudios en Sistemática de Dismorphiini (Lepidoptera: Pieridae) I: Morfología de huevos y su importancia taxonómica". *Revista de la Academia Colombiana de Ciencias Exactas, Físicas y Naturales* 31 (2007): 145–164.

Londt, J. G. H. "Afrotropical Asilidae (Diptera) 21. Observations on the biology and immature stages of Damalis femoralis Ricardo, 1925 (Trigonomiminae)". *Annals of the Natal Museum* 32 (1991): 149–162.

Lonsdale, O. and S. A. Marshall. "Redefinition of the Clusiinae and Clusiodinae, description of the new subfamily Sobarocephalinae, revision of the genus Chaetoclusia and a description of Procerosoma gen. n. (Diptera: Clusiidae)". *European Journal of Entomology* 103.1 (2006): 163–182.

Lopez O, M. and L. Cervantes P. "Life Histories of Ramosiana insignis (Blanchard) and Vulsirea violacea (F.) (Hemiptera-Heteroptera: Pentatomidae), with Descriptions of Immature Stages". *Proceedings of the Entomological Society of Washington* 112.1 (2010): 81–96.

López-Arroyo, J. I., C. A. Tauber, and M. J. Tauber. "Effects of prey on survival, development, and reproduction of trash-carrying chrysopids (Neuroptera: Ceraeochrysa)". *Environmental Entomology* 28.6 (1999): 1183–1188.

Lounibos, L. P., D. Duzak, J. R. Linley U, and R. Lourenço-de-Oliveira. “Egg Structures of Anopheles fluminensis and Anopheles shannoni”. *Memórias do Instituto Oswaldo Cruz* 92.2 (1997): 221–232.

Lourido, G., N. M. Silva, and C. Motta. “Parâmetros biológicos e injúrias de Macrosoma tipulata Hübner (Lepidoptera: Hedylidae), em cupuaçuzeiro [Theobroma grandiflorum (Wild ex Spreng Schum)] no Amazonas”. *Neotropical Entomology* 36.1 (2007): 102–106.

Lu, W., P. Souphanya, and M. E. Montgomery. “Descriptions of immature stages of Scymnus (Neopullus) sinuanodulus Yu and Yao (Coleoptera: Coccinellidae) with notes on life history”. *The Coleopterists Bulletin* 56.1 (2002): 127–141.

Lubbock, J. *Monograph of the Collembola and Thysanura*. London: Ray Society, 1873.

Lucchi, A. and E. Rossi. “The egg-burster in the Asian planthopper Ricania speculum (Walker) (Hemiptera Ricaniidae)”. *Annals of the Entomological Society of America* 109.1 (2015): 121–126.

Luff, M. L. “Diagnostic characters of the eggs of some Carabidae (Coleoptera)”. *Entomologica Scandinavica Supplement* 15 (1981): 317–327.

Luginbill Jr, P. “A contribution to the embryology of the May beetle”. *Annals of the Entomological Society of America* 46.4 (1953): 505–528.

Lundgren, J. G. “Reproductive ecology of predaceous Heteroptera”. *Biological Control* 59.1 (2011): 37–52.

Ma, N., L. Cai, and B. Hua. “Comparative morphology of the eggs in some Panorpidae (Mecoptera) and their systematic implication”. *Systematics and Biodiversity* 7.4 (2009): 403–417.

Ma, N. and B. Hua. “Fine structure and formation of the eggshell in scorpionfly Panorpa liui Hua (Mecoptera: Panorpidae)”. *Microscopy Research and Technique* 72.7 (2009): 495–500.

Ma, P. W. K., S. Baird, and S. B. Ramaswamy. “Morphology and formation of the eggshell in the tarnished plant bug, Lygus lineloaris (Palisot de Beauvois) (Hemiptera: Miridae)”. *Arthropod Structure & Development* 31.2 (2002): 131–146.

MacDonald, K. E. and S. Caveney. “External morphology and development of immature stages of Elachertus scutellatus (Hymenoptera: Eulophidae) in Florida: the first North American record”. *The Florida Entomologist* 87.4 (2004): 559–565.

MacGillivray, A. D. *Aquatic Chrysomelidae and a table of the families of Coleopterous larvae*. Albany: University of the State of New York, 1903.

Machida, R. “Evidence from embryology for reconstructing the relationships of hexapod basal clades”. *Arthropod Systematics & Phylogeny* 64.1 (2006): 95–104.

Machida, R. “External features of embryonic development of a jumping bristletail, Pedetontus unimaculatus Machida (Insecta, Thysanura, Machilidae)”. *Journal of Morphology* 168.3 (1981): 339–355.

Machida, R., T. Nagashima, and H. Ando. “The early embryonic development of the jumping bristletail Pedetontus unimaculatus Machida (Hexapoda: Microcoryphia, Machilidae)”. *Journal of Morphology* 206.2 (1990): 181–195.

Machida, R. and I. Takahashi. “Rearing technique for proturans (Hexapoda: Protura)”. *Pedobiologia* 48.3 (2004): 227–229.

MacLean, D. B. and R. L. Giese. “The life history of the ambrosia beetle Xyloterinus politus (Coleoptera: Scolytidae)”. *The Canadian Entomologist* 99.3 (1967): 285–299.

Maddox, D. M. “Bionomics of an alligatorweed flea beetle, Agasicles sp. in Argentina”. *Annals of the Entomological Society of America* 61.5 (1968): 1299–1305.

Maddox, D. M. and A. Mayfield. “Biology and life history of Amynothrips andersoni, a thrip for the biological control of alligatorweed”. *Annals of the Entomological Society of America* 72.1 (1979): 136–140.

Maeta, Y., K. Takahashi, and N. Shimada. “Host body size as a factor determining the egg complement of Strepsiptera, an insect parasite”. *International Journal of Insect Morphology and Embryology* 27.1 (1998): 27–37.

Mahmood, F. and J. B. Alexander. “Immature stages of Nemapalpus nearcticus (Diptera: Psychodidae)”. *The Florida Entomologist* 75.2 (1992): 171–178.

Mahr, E. “Normale entwicklung, pseudofurchung und die bedeutung des furchungszentrums im ei des heimchens (Gryllus domesticus)”. *Zeitschrift für Morphologie und Ökologie der Tiere* 49.3 (1960): 263–311.

Majumder, M. Z. R., M. K. Dash, R. A. Khan, and H. R. Khan. "The biology of flesh fly, Boettcherisca peregrina (Robineau-Desvoidy, 1830) (Diptera: Sarcophagidae)". *Bangladesh Journal of Zoology* 40.2 (2013): 189–196.

Maldonado, V., H. J. Finol, and J. C. Navarro. "Anopheles aquasalis eggs from two Venezuelan localities compared by scanning electron microscopy". *Memórias do Instituto Oswaldo Cruz* 92.4 (1997): 487–491.

Malihi, Y., A. Freidberg, and D. Gerling. "Bionomics of the Tamarix leaf beetle, Cryptocephalus sinaita moricei Pic, 1908 (Chrysomelidae: Cryptocephalinae)". *Israel Journal of Entomology* 44.45 (2015): 51–59.

Malipatil, M. B. "Immature stages of Ontiscus Stal (Hemiptera: Lygaeidae: Cyminae)". *Australian Journal of Entomology* 16.3 (1977): 321–326.

Malipatil, M. B. "Immature stages of some Myodochini of the Australian region (Hemiptera: Lygaeidae: Rhyparochrominae)". *Australian Journal of Zoology* 26.3 (1978): 555–584.

Malipatil, M. B. "Immature stages of some New Zealand Rhyparochrominae (Hemiptera: Lygaeidae)". *New Zealand Journal of Zoology* 2.4 (1975): 381–388.

Malipatil, M. B. and R. Kumar. "Biology and immature stages of some Queensland Pentatomomorpha (Hemiptera: Heteroptera)". *Australian Journal of Entomology* 14.2 (1975): 113–128.

Malmqvist, B., P. H. Adler, and D. Strasevicius. "Testing hypotheses on egg number and size in black flies (Diptera: Simuliidae)". *Journal of Vector Ecology* 29.2 (2004): 248–256.

Mamlayya, A. B., S. R. Aland, S. M. Gaikwad, and G. P. Bhawane. "Life History and Diet Breadth of Apoderus tranquebaricus Fab. (Coleoptera: Attelabidae)". *Biological Forum* 2.2 (2011): 46–48.

Manjunatha, H. B. and H. P. Puttaraju. "The egg of Uzi fly, Exorista sorbillans (? E. Bombycis Louis) (Diptera: Tachinidae)". *Applied Entomology and Zoology* 28.4 (1993): 574–577.

Maple, J. D. *The eggs and first instar larvae of Encyrtidae and their morphological adaptations for respiration*. London: Cambridge University Press, 1947.

Marchiondo, A. A., S. M. Meola, K. G. Palma, J. H. Slusser, and R. W. Meola. "Chorion formation and ultrastructure of the egg of the cat flea (Siphonaptera: Pulicidae)". *Journal of Medical Entomology* 36.2 (1999): 149–157.

Marini, M. and G. Campadelli. "Ootaxonomy of Goniini (Diptera Tachinidae) with microtype eggs". *Italian Journal of Zoology* 61.3 (1994): 271–283.

Markow, T. A., S. Beall, and L. M. Matzkin. "Egg size, embryonic development time and ovoviviparity in Drosophila species". *Journal of Evolutionary Biology* 22.2 (2009): 430–434.

Marshall, A. T. and P. M. Marshall. "The life history of a tube-dwelling Cercopoid: Machaerota coronata Maa (Homoptera: Machaerotidae)". *Proceedings of the Royal Entomological Society of London. Series A, General Entomology* 41.1-3 (1966): 17–20.

Marshall, L. D. "Intra-specific variation in reproductive effort by female Parapediasia teterrella (Lepidoptera: Pyralidae) and its relation to body size". *Canadian Journal of Zoology* 68.1 (1989): 44–48.

Martins, F. S. and L. A. Campos. "Morphology and biology of immatures of Euschistus hansi (Hemiptera, Heteroptera, Pentatomidae)". *Iheringia, Série Zoologia* 96.2 (2006): 213–218.

Martins, G. F. and J. Serrão. "A comparative study of the ovaries in some Brazilian bees (Hymenoptera; Apoidea)". *Papéis Avulsos de Zoologia (São Paulo)* 44.3 (2004): 45–53.

Marvaldi, A. E. "Eggs and oviposition habits in Entimini (Coleoptera: Curculionidae)". *The Coleopterists' Bulletin* 53.2 (1999): 115–126.

Mashimo, Y., R. G. Beutel, R. Dallai, M. Gottardo, C.-Y. Lee, and R. Machida. "The morphology of the eggs of three species of Zoraptera (Insecta)". *Arthropod Structure & Development* 44 (2015): 656–666.

Mashimo, Y., R. G. Beutel, R. Dallai, C.-Y. Lee, and R. Machida. "Embryonic development of Zoraptera with special reference to external morphology, and its phylogenetic implications (Insecta)". *Journal of Morphology* 275.3 (2014): 295–312.

Mashimo, Y., M. Fukui, and R. Machida. "Egg structure and ultrastructure of Paterdecolyus yanbarensis (Insecta, Orthoptera, Anostostomatidae, Anabropsinae)". *Arthropod Structure & Development* 45.6 (2016): 637–641.

Mashimo, Y., R. Machida, R. Dallai, M. Gottardo, D. Mercati, and R. G. Beutel. "Egg structure of Zorotypus caudelli Karny (Insecta, Zoraptera, Zorotypidae)". *Tissue and Cell* 43.4 (2011): 230–237.

Maso, A. and V. S. i Monteys. “Confirmación de Cacyreus marshalli Butler, 1898 (Lycaenidae, Polyommatinae) como nueva especie para la fauna europea”. *Boletín de Sanidad Vegetal Plagas* 17.1 (1991): 173–183.

Mason, W. R. M. “Specialization in the egg structure of Exenterus (Hymenoptera: Ichneumonidae) in relation to distribution and abundance”. *The Canadian Entomologist* 99.4 (1967): 375–384.

Masuko, K. “The instars of the ant Amblyopone silvestrii”. *Sociobiology* 17.2 (1990): 221–244.

Materu, M. E. A. “Morphology of adults and description of the young stages of Acanthomia tomentosicollis Stål. and A. horrida Germ. (Hemiptera, Coreidae)”. *Journal of Natural History* 6.4 (1972): 427–450.

Matesco, V. C., B. B. R. J. Fürstenau, and J. L. C. Bernardes. “Morphological features of the eggs of Pentatomidae (Hemiptera: Heteroptera)”. *Zootaxa* 1984 (2009): 1–30.

Matesco, V. C., C. F. Schwertner, and J. Grazia. “Immature stages of Chinavia musiva (Berg, 1878): a unique pattern in the morphology of Chinavia Orian, 1965 (Hemiptera, Pentatomidae)”. *Journal of Natural History* 42.25-26 (2008): 1749–1763.

Matesco, V. C., C. F. Schwertner, and J. Grazia. “Morphology of the immatures and biology of Chinavia longicorialis (Breddin) (Hemiptera: Pentatomidae)”. *Neotropical Entomology* 38.1 (2009): 74–82.

Matesco, V. C., C. F. Schwertner, and J. Grazia. “Description of the immature stages and biology of Chinavia pengue (Rolston) (Hemiptera, Pentatomidae)”. *Revista Brasileira de Entomologia* 51.1 (2007): 93–100.

Matesco, V. C., F. M. Bianchi, L. A. Campos, and J. Grazia. “Egg ultrastructure of two species of Galgupha Amyot & Serville, with a discussion of the eggs and oviposition patterns of thyreocorid and allied groups (Hemiptera: Heteroptera: Pentatomoidea: Thyreocoridae)”. *Zootaxa* 3247 (2012): 43–51.

Matesco, V. C., F. M. Bianchi, B. Fürstenau, P. P. da Silva, L. A. Campos, and J. Grazia. “External egg structure of the Pentatomidae (Hemiptera: Heteroptera) and the search for characters with phylogenetic importance”. *Zootaxa* 3768.3 (2014): 351–385.

Matheny, E. L. and E. A. Heinrichs. “Chorion characteristics of sod webworm eggs”. *Annals of the Entomological Society of America* 65.1 (1972): 238–246.

Matsumoto, R. and T. Saigusa. “The biology and immature stages of Thrybius togashii Kusigemati (Hymenoptera: Ichneumonidae: Cryptinae), with a description of the male”. *Journal of Natural History* 35.10 (2001): 1507–1516.

Matsuo, K. “Scanning electron microscopy of mosquitoes. IV. The egg surface structure of 3 species of Aedes from Japan”. *Japanese Journal of Sanitary Zoology* 26.1 (1975): 49–53.

Matsuura, K. “Termite-egg mimicry by a sclerotium-forming fungus”. *Proceedings of the Royal Society B: Biological Sciences* 273.1591 (2006): 1203–1209.

Matsuura, K. and N. Kobayashi. “Size, hatching rate, and hatching period of sexually and asexually produced eggs in the facultatively parthenogenetic termite Reticulitermes speratus (Isoptera: Rhinotermitidae)”. *Applied Entomology and Zoology* 42.2 (2007): 241–246.

Matsuura, K. and N. Kobayashi. “Termite queens adjust egg size according to colony development”. *Behavioral Ecology* 21.5 (2010): 1018–1023.

Matsuura, K. and T. Yashiro. “Parallel evolution of termite-egg mimicry by sclerotium-forming fungi in distant termite groups”. *Biological Journal of the Linnean Society* 100.3 (2010): 531–537.

Matsuzaki, M., H. Ando, and S. N. Visscher. “Fine structure of oocyte and follicular cells during oogenesis in Galloisiana Nipponensis (Caudell and King) (Grylloblattodea: Grylloblattidae)”. *International Journal of Insect Morphology and Embryology* 8.5-6 (1979): 257–263.

Matthews, R. W. “Nesting biology of the stem-nesting wasp Psenulus interstitialis Cameron (Hymenoptera: Crabronidae: Pemphredoninae) on Magnetic Island, Queensland”. *Australian Journal of Entomology* 39.1 (2000): 25–28.

Matzke, D. and K.-D. Klass. “Reproductive biology and nymphal development in the basal earwig Tagalina papua (Insecta: Dermaptera: Pygidicranidae), with a comparison of brood care in Dermaptera and Embioptera”. *Entomologische Abhandlungen* 62.2 (2005): 99–116.

May, B. M. "The immature stages of Dieuches notatus(Dallas) (Hemiptera: Lygaeidae: Rhyparochrominae)". *New Zealand Journal of Science* 8.3 (1965): 359–367.

May, M. L. "Comparative notes on micropyle structure in "cordulegastroid" and "libelluloid" Anisoptera". *Odonatologica* 24.1 (1995): 53–62.

Mayhew, P. J. and W. R. B. Heitmans. "Life history correlates and reproductive biology of Laelius pedatus (Hymenoptera: Bethylidae) in The Netherlands". *European Journal of Entomology* 97.3 (2000): 313–322.

Mazanec, Z. "Immature stages and life history of Enytus sp. (Hymenoptera: Ichneumonidae), a parasitoid of Perthida glyphopa Common (Lepidoptera: Incurvariidae)". *Australian Journal of Entomology* 29.1 (1990): 57–66.

Mazanec, Z. "The immature stages and life history of Diaulomorpha sp. (Hymenoptera: Eulophidae), a parasitoid of Perthida glyphopa Common (Lepidoptera: Incurvariidae)". *Australian Journal of Entomology* 29.2 (1990): 147–159.

Mazanec, Z. "The immature stages and life history of the jarrah leafminer, Perthida glyphopa Common (Lepidoptera: Incurvariidae)". *Australian Journal of Entomology* 22.2 (1983): 101–108.

Mazzini, M. "Amino acid analysis and morphology of the egg shell of Tettigonia viridissoima L. (Orthoptera: Tettigoniidae)". *International Journal of Insect Morphology and Embryology* 7.3 (1978): 205–214.

Mazzini, M. "Fine structure of the insect micropyle-III. Ultrastructure of the egg of Chrysopa carnea Steph. (Neuroptera: Chrysopidae)". *International Journal of Insect Morphology and Embryology* 5.4 (1976): 273–278.

Mazzini, M. "Overview of Egg Structure in Orthopteroid Insects". *Evolutionary biology of orthopteroid insects*. New York: Halsted Press, 1987. 358–372.

Mazzini, M. "Sulla fine struttura del micropilo negli insetti. IV. Le sculture corionidee come mezzo di identificazione delle uova degli Ortotteri Tettigonioidei". *Redia* 59 (1976): 109–134.

Mazzini, M., G. Callaini, and C. Mencarelli. "A comparative analysis of the evolution of the egg envelopes and the origin of the yolk". *Italian Journal of Zoology* 51.1-2 (1984): 35–101.

Mazzini, M., M. Carcupino, and A. M. Fausto. "Egg chorion architecture in stick insects (Phasmatodea)". *International Journal of Insect Morphology and Embryology* 22.2 (1993): 391–415.

Mazzini, M., M. Carcupino, and L. Santini. "Ootaxonomic investigation of three species of Mycomya (Diptera, Mycetophilidae): a scanning electron microscope study". *Italian Journal of Zoology* 59.1 (1992): 33–39.

Mazzini, M. and E. Gaino. "Fine structure of the egg shells of Habrophlebia fusca (Curtis) and H. consiglioi Biancheri (Ephemeroptera: Leptophlebiidae)". *International Journal of Insect Morphology and Embryology* 14.6 (1985): 327–334.

Mazzini, M. and V. Scali. "Ultrastructure and amino acid analysis of the eggs of the stick insects, Lonchodes pterodactylus Gray and Carausius morosus Br. (Phasmatodea: Heteronemiidae)". *International Journal of Insect Morphology and Embryology* 9.5 (1980): 369–382.

McClendon, J. F. R. "The life history of Ulula hyalina Latreille". *American Naturalist* 36.426 (1902): 421–429.

McClure, R. G. and E. W. Stewart. "Life cycle and production of the mayfly Choroterpes (Neochoroterpes) mexicanus Allen (Ephemeroptera: Leptophlebiidae)". *Annals of the Entomological Society of America* 69.1 (1976): 134–144.

McCorquodale, D. B. "Oocyte development in the primitively social wasp, Cerceris antipodes (Hymenoptera Sphecidae)". *Ethology Ecology & Evolution* 2.4 (1990): 345–361.

McGinley, R. J. and J. G. Rozen Jr. "Nesting Biology, Immature Stages, and Phylogenetic Placement of the Palaearctic Bee Pararhophites (Hymenoptera, Apoidea)". *American Museum Novitates* 2903 (1987): 1–21.

McGuffin, W. C. "Immature stages of some Lepidoptera of Durango, Mexico". *The Canadian Entomologist* 99.11 (1967): 1215–1229.

McGuffin, W. C. "The immature stages of the Canadian species of Pero Herrich-Schaeffer (Lepidoptera: Geometridae)". *The Canadian Entomologist* 95.11 (1963): 1159–1167.

McPherson, J. E. and R. J. Packauskas. "Life history and laboratory rearing of Belostoma lutarium (Heteroptera: Belostomatidae) with descriptions of immature stages". *Journal of the New York Entomological Society* 94.2 (1986): 154–162.

McPherson, J. E. and R. J. Packauskas. “Life history and laboratory rearing of Nepa apiculata (Heteroptera: Nepidae), with descriptions of immature stages”. *Annals of the Entomological Society of America* 80.5 (1987): 680–685.

McPherson, J. E., R. J. Packauskas, and P. P. Korch. “Life history and laboratory rearing of Pelocoris femoratus(Hemiptera: Naucoridae), with descriptions of immature stages”. *Proceedings of the Entomological Society of Washington* 89.2 (1987): 288–295.

McPherson, J. E. and S. M. Paskewitz. “Life history and laboratory rearing of Euschistus ictericus (Hemiptera: Pentatomidae), with descriptions of immature stages”. *Journal of the New York Entomological Society* 92.1 (1984): 53–60.

McPherson, K. R. and S. W. Wilson. “Life history and descriptions of immatures of the dictyopharid planthopper Phylloscelis pallescens (Homoptera: Fulgoroidea)”. *Journal of the New York Entomological Society* 103.2 (1995): 170–179.

Meier, R. “A comparative SEM study of the eggs of the Sepsidae (Diptera) with a cladistic analysis based on egg, larval and adult characters”. *Insect Systematics & Evolution* 26.4 (1995): 425–438.

Melk, J. P. and S. Govind. “Developmental analysis of Ganaspis xanthopoda, a larval parasitoid of Drosophila melanogaster”. *Journal of Experimental Biology* 202.14 (1999): 1885–1896.

Mellanby, H. “Memoirs: the early embryonic development of Rhodnius prolixus (Hemiptera, Heteroptera)”. *Journal of Cell Science* 2.309 (1935): 71–90.

Men, Q. “First record of female Tipula (Formotipula) vindex Alexander with description of eggs, and redescription of male (Diptera: Tipulidae)”. *Entomologica Americana* 120.1 (2014): 1–3.

Mendonça, P. M., R. R. Barbosa, L. B. Cortinhas, J. R. dos Santos-Mallet, and M. M. de Carvalho Queiroz. “Ultrastructure of immature stages of Cochliomyia macellaria (Diptera: Calliphoridae), a fly of medical and veterinary importance”. *Parasitology Research* 113.10 (2014): 3675–3683.

Mendonça, P. M., J. R. dos Santos-Mallet, R. P. de Mello, L. Gomes, and M. M. de Carvalho Queiroz. “Identification of fly eggs using scanning electron microscopy for forensic investigations”. *Micron* 39.7 (2008): 802–807.

Mertins, J. W. “Life history and morphology of the odd beetle, Thylodrias contractus”. *Annals of the Entomological Society of America* 74.6 (1981): 576–581.

Metcalf, C. L. “The Syrphidae of Ohio”. MA thesis. Ohio State University, 1913.

Michaelis, F. B. “The distribution and life history of Rakiura vernale (Trichoptera: Helicopsychidae)”. *Journal of the Royal Society of New Zealand* 3.2 (1973): 295–304.

Michalik, A., M. Miliša, K. Michalik, and E. Rościszewska. “The structure and ultrastructure of the egg capsules of stoneflies of the genus Isoperla (Insecta, Plecoptera, Perlodidae)”. *Microscopy Research and Technique* 80.11 (2017): 1234–1246.

Michalik, A., E. Rościszewska, and M. Miliša. “The structure and ultrastructure of the egg capsule of Brachyptera risi (Plecoptera, Nemouroidea, Taeniopterygidae) with some remarks concerning choriogenesis”. *Microscopy Research and Technique* 78.2 (2015): 180–186.

Michel, B. “Sur la ponte et l’eclosion des Ascalaphes afrotropicaux (Neuroptera, Ascalaphidae)”. *Bulletin de la Societe Entomologique de France* 106.4 (2001): 401–408.

Michelsen, V. “Proposal of Karliella gen. n. for the Afrotropical’Pegomya’sexpunctata Karl, 1935 (Diptera: Anthomyiidae), a Possible Kleptoparasite of Dung-Breeding Beetles”. *African Invertebrates* 54.2 (2013): 335–347.

Michener, C. D. “Size and form of eggs of allodapine bees”. *Journal of the Entomological Society of Southern Africa* 36.2 (1973): 281–285.

Miles, H. W. “Biological Studies of Sawflies infesting Ribes”. *Bulletin of Entomological Research* 23.1 (1932): 1–15.

Miles, M. “On the life-history of Blastodacna atra Haw., the pith moth of the apple”. *Annals of Applied Biology* 17.4 (1930): 775–795.

Miliczky, E. “Observations on the nesting biology of Andrena (Plastandrena) prunorum Cockerell in Washington state (Hymenoptera: Andrenidae)”. *Journal of the Kansas Entomological Society* 81.2 (2008): 110–121.

Miliczky, E. R. “Observations on the nesting biology of Tetralonia hamata Bradley with a description of its mature larva (Hymenoptera: Anthophoridae)”. *Journal of the Kansas Entomological Society* 58.4 (1985): 686–700.

Milléo, J., J. P. Castro, C. S. Ribeiro-Costa, and J. M. T. de Souza. "The first record of Litargus tetraspilotus (Coleoptera, Mycetophagidae) in Brazil, with biological notes and complementary description of the species". *Iheringia, Série Zoologia* 101.1-2 (2011): 24–32.

Miller, A. "The egg and early development of the stonefly, Pteronarcys proteus Newman (Plecoptera)". *Journal of Morphology* 64.3 (1939): 555–609.

Miller, N. C. E. "The developmental stages of some Malayan Rhynchota". *Journal of the Federated Malay States Museums* 17 (1934): 502–525.

Miller, P. L. "Oviposition behaviour and eggshell structure in some libellulid dragonflies, with particular reference to Brachythemis lacustris (Kirby) and Orthetrum coerulescens (Fabricius) (Anisoptera)". *Odonatologica* 16.4 (1987): 361–374.

Milliron, H. E. "Notes on the nesting of Bombus morio (Swederus) (Hymenoptera: Apidae)". *The Canadian Entomologist* 93.11 (1961): 1017–1019.

Milton, M. C. and P. Venkatesan. "Role of fixatives on the egg capsule of the aquatic belostomatid bug, Diplonychus indicus Venk. & Rao (Hemiptera: Belostomatidae)". *Journal of Entomological Research* 23.1 (1999): 35–40.

Minter, L. R. "The egg and larval stages of Nallachius krooni Minter (Insecta: Neuroptera: Dilaridae)". *Current Research in Neuropterology: Proceedings of the Fourth International Symposium on Neuropterology*. Toulouse: Distribution, M. Canard, 1992. 101–113.

Mitchell, R. and B. L. Redmond. "Fine structure and respiration of the eggs of two ephydrid flies (Diptera: Ephydridae)". *Transactions of the American Microscopical Society* 93.1 (1974): 113–118.

Miwa, K. and L. J. Meinke. "Developmental biology and effects of adult diet on consumption, longevity, and fecundity of Colaspis crinicornis (Coleoptera: Chrysomelidae)". *Journal of Insect Science* 15.1 (2015): 1–8.

Miyakawa, K. "Embryology of the Dobsonfly, Protohermes grandis Thunberg (Megaloptera: Corydalidae): I. Changes in External Form of the Embryo during Development". *Kontyû* 47.3 (1979): 367–375.

Miyakawa, K. "The embryology of the caddisfly Stenopsyche griseipennis maclachlan (Trichoptera: Stenopsychidae): I. Early stages and changes in external form of embryo". *Kontyû* 41.4 (1973): 413–425.

Miyamoto, D. M. and J. M. van der Meer. "Early egg contractions and patterned parasynchronous cleavage in a living insect egg". *Wilhelm Roux's Archives of Developmental Biology* 191.2 (1982): 95–102.

Mizutani, K. and Y. Nakashima. "Development and Morphological Characteristics of Immatures of Two Aphid Hyperparasitoids: Dendrocerus carpenteri (Curtis) (Hymenoptera: Megaspilidae) and Asaphes suspensus (Nees) (Hymenoptera: Pteromalidae)". *Annual Report of the Society of Plant Protection of North Japan* 59 (2008): 189–194.

Mohan Rao, H. N. and G. T. Tonapi. "A study on the developmental bionomics of Dineutes indicus Aube (Gyrinidae, Coleoptera)". *Journal of Natural History* 2.2 (1968): 263–271.

Molineri, C. "Phylogeny of the mayfly family Leptohyphidae (Insecta: Ephemeroptera) in South America". *Systematic Entomology* 31.4 (2006): 711–728.

Molineri, C. and E. Domínguez. "Nymph and egg of Melanemerella brasiliana (Ephemeroptera: Ephemerelloidea: Melanemerellidae), with comments on its systematic position and the higher classification of Ephemerelloidea". *Journal of the North American Benthological Society* 22.2 (2003): 263–275.

Molineri, C., J. G. Peters, and M. del Carmen Zuniga de Cardoso. "A new family, Coryphoridae (Ephemeroptera: Ephernerelloidea), and description of the winged and egg stages of Coryphorus". *Insecta Mundi* 15.2 (2001): 117–122.

Monserrat, J. "Morfologia del huevo en los nemopteridos ibericos". *Actas do Congresso Iberico de Entomologia* 2 (1985): 463–474.

Monserrat, V. J. "Larval stages of European Nemopterinae, with systematic considerations on the family Nemopteridae (Insecta, Neuroptera)". *Deutsche Entomologische Zeitschrift* 43.1 (1996): 99–121.

Monserrat, V. J. "Nuevos datos sobre algunas especies de Nemopteridae y Crocidae (Insecta: Neuroptera)". *Heteropterus Revista de Entomología* 8.1 (2008): 1–33.

Montgomery, V. E. and P. R. DeWitt. "Morphological differences among immature stages of three genera of exotic larval parasitoids attacking the cereal leaf beetle in the United States". *Annals of the Entomological Society of America* 68.3 (1975): 574–578.

Morales Agacino, E. "Las ootecas de los Acrididos". *Boletin de Patologia Vegetal y Entomologia Agricola* 18 (1951): 89–109.

Moran, V. C. "The adult and immature stages of a new species in the genus Paurocephala (Homoptera: Psyllidae) from South Africa". *Proceedings of the Royal Entomological Society of London. Series B, Taxonomy* 37.3-4 (1968): 50–56.

Moratorio, M. S. and E. Chiappini. "Biology of Anagrus incarnatosimilis and Anagrus breviphragma (Hymenoptera: Mymaridae)". *Bollettino di Zoologia Agraria e di Bachicoltura* 27.2 (1995): 143–162.

Morgan, N. C. "The biology of Leptocerus aterrimus Steph. with reference to its availability as a food for trout". *The Journal of Animal Ecology* 25.2 (1956): 349–365.

Mori, H. "Abortive anatrepsis and imperfect germ-band formation in the capillary-coated eggs of the waterstrider, Gerris paludum insularis (Hemiptera: Gerridae)". *Annals of the Entomological Society of America* 78.4 (1985): 509–513.

Moritz, G. "Morphogenetic development of some species of the order Thysanoptera (Insecta)". *Thrips Biology and Management*. New York: Plenum Press, 1995. 489–504.

Morrill, A. W. "Notes on the immature stages of some tingitids of the genus Corythuca. Plate 3". *Psyche* 10 (1903): 127–134.

Moscona, A. "Blastokinesis and embryonic development in a phasmid". *Cellular and Molecular Life Sciences* 6.11 (1950): 425–426.

Moscona, A. "Studies of the egg of Bacillus libanicus (Orthoptera, Phasmidae) I. The egg envelopes". *Quarterly Journal of Microscopical Science* 3.14 (1950): 183–193.

Mound, L. A. "The first thrips species (Insecta) inhabiting leaf domatia: Domatiathrips cunninghamii gen. et sp. nov.(Thysanoptera: Phlaeothripidae)". *Journal of the New York Entomological Society* 101.3 (1993): 424–430.

Mouzaki, D. G. and L. H. Margaritis. "The eggshell of the cherry fly Rhagoletis cerasi". *Tissue and Cell* 23.5 (1991): 745–754.

Mouzaki, D. G. and L. H. Margaritis. "The eggshell of the almond wasp Eurytoma amygdali (Hymenoptera, Eurytomidae)-1. Morphogenesis and fine structure of the eggshell layers". *Tissue and Cell* 26.4 (1994): 559–568.

Mouzaki, D. G., F. E. Zarani, and L. H. Margaritis. "Structure and morphogenesis of the eggshell and micropylar apparatus in the olive fly, Dacus oleae (Diptera: Tephritidae)". *Journal of Morphology* 209.1 (1991): 39–52.

Mühlenberg, M. "Die abwandlung des eilegeapparates der bombyliidae (Diptera) Eine funktionsmorphologische studie". *Zoomorphology* 70.1 (1971): 1–72.

Mukerji, D. "Embryology of termites". *Biology of Termites* 2 (1970): 37–72.

Mullins, D. E., K. J. Mullins, and K. R. Tignor. "The structural basis for water exchange between the female cockroach (Blattella germanica) and her ootheca". *Journal of Experimental Biology* 205.19 (2002): 2987–2996.

Munguira, M. L., J. Martin, E. Garcia-Barros, G. Shahbazian, and J. P. Cancela. "Morphology and morphometry of Lycaenid eggs (Lepidoptera: Lycaenidae)". *Zootaxa* 3937.2 (2015): 201–247.

Munyaneza, J. and J. E. McPherson. "Comparative study of life histories, laboratory rearing, and immature stages of Euschistus servus and Euschistus variolarius (Hemiptera: Pentatomidae)". *Great Lakes Entomologist* 26.4 (1994): 253–323.

Murao, R. and O. Tadauchi. "Description of Immature Stages of Colletes esakii (Hymenoptera, Colletidae)". *Esakia* 45 (2005): 55–60.

Murillo, T. and L. F. Jirón. "Egg morphology of Anastrepha obliqua and some comparative aspects with eggs of Anastrepha fraterculus (Diptera: Tephritidae)". *The Florida Entomologist* 77.3 (1994): 342–348.

Murvosh, C. M. "Microdistribution of the water penny Psephenus montanus (Coleoptera: Psephenidae), with notes on life history and zoogeography". *The Southwestern Naturalist* 38.2 (1993): 119–126.

Nacro, S. and J.-P. Nenon. "Anatomy of the Female Reproductive System and the Ultrastructure of". *Journal of Entomology* 3.1 (2006): 16–22.

Nacro, S. and J.-P. Nenon. "Female reproductive biology of Platygaster diplosisae (Hymenoptera: Platygastridae) and Aprostocetus procerae (Hymenoptera: Eulophidae), two parasitoids associated with the African rice gall midge, Orseolia oryzivora (Diptera: Cecidomyiidae)". *Entomological Science* 11.2 (2008): 231–237.

Nagamine, W. T. and M. E. Epstein. "Chronicles of Darna pallivitta (Moore 1877) (Lepidoptera: Limacodidae): biology and larval morphology of a new pest in Hawaii". *The Pan-Pacific Entomologist* 83.2 (2007): 120–135.

Nakamura, I. "Female anal hair tuft in Nordmannia myrtale (Lycaenidae): egg-camouflaging function and taxonomic significance". *Journal of the Lepidopterists' Society* 30.4 (1976): 305–309.

Nakasuji, F. and M. Kimura. "Seasonal polymorphism of egg size in a migrant skipper, Parnara guttata guttata (Lepidoptera, Hesperiidae)". *Kontyû* 52.2 (1984): 253–259.

Nalepa, C. A. and M. Lenz. "The ootheca of Mastotermes darwiniensis Froggatt (Isoptera: Mastotermitidae): homology with cockroach oothecae". *Proceedings of the Royal Society of London B: Biological Sciences* 267.1454 (2000): 1809–1813.

Nápoles, J. R., M. A. D. R. G. Peña, and C. D. Johnson. "Ecology of Stator dissimilis Johnson & Kingsolver (Coleoptera: Chrysomelidae: Bruchinae) in seeds of Lepechinia (Lamiaceae) a new host genus for bruchines, with an ecological comparison to other species of Stator". *The Coleopterists Bulletin* 60.1 (2006): 81–85.

Narayanan, E. S., B. R. S. Rao, and K. R. Thakare. "The biology and some aspects of morphology of the immature stages of Chelonus narayani Subba Rao (Braconidae: Hymenoptera)". *Proceedings of the Indian National Science Academy, Section B* 27 (1961): 68–82.

Nath, M. and I. Rahman. "Biology of bunch caterpillar, Andraca bipunctata walker-A major insect pest of tea in north-east India". *Indian Journal of Entomology* 74.4 (2012): 303–305.

Navasero, M. M., M. V. Navasero, et al. "Biology of the black earwig Chelisoches morio (Fabricius) (Chelisochidae, Dermaptera)". *Philippine Entomologist* 24.2 (2010): 122–136.

Neal, J. W. "Bionomics and instar determination of Synanthedon rhododendri (Lepidoptera: Sesiidae) on rhododendron". *Annals of the Entomological Society of America* 77.5 (1984): 552–560.

Neal, J. W. "Bionomics of immature stages and ethology of Neochlamisus platani (Coleoptera: Chrysomelidae) on American sycamore". *Annals of the Entomological Society of America* 82.1 (1989): 64–72.

Needham, J. G., J. R. Traver, and Y.-C. Hsu. *The biology of mayflies: with a systematic account of North American species*. Ithaca: Comstock Publishing Company, Inc., 1969.

Nelson, C. H. "Note on the phylogenetic systematics of the family Pteronarcyidae (Plecoptera), with a description of the eggs and nymphs of the Asian species". *Annals of the Entomological Society of America* 81.4 (1988): 560–576.

Nelson, G. H., R. L. Westcott, and T. C. MacRae. "Miscellaneous notes on Buprestidae and Schizopodidae occurring in the United States and Canada, including descriptions of previously unknown sexes of six Agrilus Curtis (Coleoptera)". *The Coleopterists' Bulletin* 50.2 (1996): 183–191.

Nénon, J.-P., G. Boivin, and M. R. Allo. "Fine structure of the egg envelopes in Listronotus oregonensis (Leconte) (Coleoptera: Curculionidae) and morphological adaptations to oviposition sites". *International Journal of Insect Morphology and Embryology* 24.3 (1995): 333–342.

Neveu, N., M. R. Allo, J. P. Nénon, X. Langlet, E. Brunel, M. Lahmer, and G. Boivin. "The fine structure of the egg shells of the cabbage maggot, Delia radicum L.(Diptera: Anthomyiidae), and its relation with developmental conditions and oviposition site". *Canadian Journal of Zoology* 75.4 (1997): 535–541.

New, T. R. "Ovariolar dimorphism and repagula formation in some South American Ascalaphidae (Neuroptera)". *Journal of Entomology Series A, General Entomology* 46.1 (1971): 73–77.

New, T. R. "The egg and first instar larva of Stenosmylus (Neuroptera: Osmylidae)". *Australian Entomological Magazine* 2.2 (1974): 24–27.

Nieves-Uribe, S., J. Castro-Gerardino, A. Flores-Gallardo, and J. Llorente-Bousquets. "Corion en los Géneros Anteos y Rhabdodryas 1: su Significado e Implicaciones". *Southwestern Entomologist* 41.2 (2016): 485–504.

Nieves-Uribe, S., J. Castro-Gerardino, A. Flores-Gallardo, and J. Llorente-Bousquets. “Estudio del Corion de Tres Especies del Género Colias Fabricius, 1807 y Zerene cesonia cesonia (Stoll, 1790)”. *Southwestern Entomologist* 41.4 (2016): 1121–1141.

Nieves-Uribe, S., A. Flores-Gallardo, B. C. Hernández-Mejía, and J. Llorente-Bousquets. “Exploración Morfológica del Corion en Biblidinae (Lepidoptera: Nymphalidae): Aspectos Filogenéticos y Clasificatorios”. *Southwestern Entomologist* 40.3 (2015): 589–648.

Niikura, K., K. Hirasawa, T. Inoda, and Y. Kobayashi. “Embryonic Development of a Diving Beetle, Hydaticus pacificus Aubé (Insecta: Coleoptera; Dytiscidae): External Morphology and Phylogenetic Implications”. *Proceedings of Arthropodan Embryological Society of Japan* 48 (2017): 19–32.

Niva, C. C. and M. Becker. “Embryonic external morphogenesis of Rhammatocerus conspersus (Bruner) (Orthoptera: Acrididae: Gomphocerinae) and determination of the diapausing embryonic stage”. *Anais da Sociedade Entomológica do Brasil* 27.4 (1998): 557–583.

Nogueira, G. A. d. L., S. R. Rodrigues, and E. F. Tiago. “Biological aspects of Cyclocephala tucumana Brethes, 1904 and Cyclocephala melanocephala (Fabricius, 1775) (Coleoptera: Scarabaeidae)”. *Biota Neotropica* 13.1 (2013): 86–90.

Nogueira, G., M.-A. Morón, H.-E. Fierros-López, and J.-L. Navarrete-Heredia. “The immature stages of Neoscelis dohrni (Westwood) (Coleoptera: Scarabaeidae: Cetoniinae: Goliathini) with notes on adult behavior”. *The Coleopterists Bulletin* 58.2 (2004): 171–183.

Nolte, U. and T. Hoffmann. “Fast life in cold water: Diamesa incallida (Chironomidae)”. *Ecography* 15.1 (1992): 25–30.

Nonci, N. “Biology and intrinsic growth rate of earwig (Euborellia annulata)”. *Indonesian Journal of Agricultural Science* 6.2 (2013): 69–74.

Nondillo, A., D. R. Solis, E. G. P. Fox, M. L. Rossi, M. Botton, and O. C. Bueno. “Description of the immatures of workers of the ant Linepithema micans forel (Hymenoptera: Formicidae)”. *Microscopy Research and Technique* 74.4 (2011): 337–342.

Nutting, W. L. “Observations on the Reproduction of the Giant Cockroach, Blaberus craniifera Brum”. *Psyche* 60.1 (1953): 6–14.

O’Flynn, M. A. and D. E. Moorhouse. “Identification of early immature stages of some common Queensland carrion flies”. *Australian Journal of Entomology* 19.1 (1980): 53–61.

O’Neill, K. M. and S. W. Skinner. “Ovarian egg size and number in relation to female size in five species of parasitoid wasps”. *Journal of Zoology* 220.1 (1990): 115–122.

O’Neill, K. M. “Egg size, prey size, and sexual size dimorphism in digger wasps (Hymenoptera: Sphecidae) - Canadian Journal of Zoology”. *Canadian Journal of Zoology* 63.9 (1985): 2187–2193.

Obara, M. T., J. A. Da Rosa, N. N. Da Silva, W. Ceretti Jr, P. R. Urbinatti, J. Barata, J. Jurberg, and C. Galvão. “Estudo morfológico e histológico dos ovos de seis espécies do gênero Triatoma (Hemiptera: Reduviidae)”. *Neotropical Entomology* 36.5 (2007): 798–806.

Oetting, R. D. “Biology of the cactus scale, Diaspis echinocacti (Bouche) (Homoptera: Diaspididae)”. *Annals of the Entomological Society of America* 77.1 (1984): 88–92.

Oetting, R. D. and T. R. Yonke. “Immature stages and biology of Podisus placidus and Stiretrus fimbriatus (Hemiptera: Pentatomidae)”. *The Canadian Entomologist* 103.11 (1971): 1505–1516.

Oetting, R. D. and T. R. Yonke. “Immature stages and notes on the biology of Hymenarcys crassa (Hemiptera: Pentatomidae)”. *Annals of the Entomological Society of America* 65.2 (1972): 474–478.

Ogorzałek, A. “Structural and functional diversification of follicular epithelium in Coreus marginatus (Coreidae: Heteroptera)”. *Arthropod Structure & Development* 36.2 (2007): 209–219.

Oishi, M. and H. Sato. “Life history traits, larval habits and larval morphology of a leafminer, Coptotriche japoniella (Tischeriidae), on an evergreen tree, Eurya japonica (Theaceae), in Japan”. *Journal of the Lepidopterists’ Society* 63.2 (2009): 93–99.

Okada, M. “Embryonic development of the rice stemborer, Chilo suppressalis”. *Science Reports of the Tokyo Kyoiku Daigaku* 9.143 (1960): 243–296.

Olivares, T. S., S. A. Torres, and L. A. Zúñiga. “Morfología de huevos de siete especies de nóctuidos de Chile (Lepidoptera: Noctuidae) y clave actualizada para su identificación”. *Revista de Biología Tropical* 53.1-2 (2005): 153–163.

Onagbola, E. O. and H. Y. Fadamiro. “Morphology and development of Pteromalus cerealellae (Ashmead) (Hymenoptera: Pteromalidae) on Callosobruchus maculatus (F.) (Coleoptera: Chrysomelidae)”. *BioControl* 53.5 (2008): 737–750.

Onoyama, K. “Immature stages of the harvester ant Messor aciculatus (Hymenoptera, Formicidae)”. *Kontyû* 50.2 (1982): 324–329.

Onsager, J. A. and G. B. Mulkern. *Identification of eggs and egg-pods of North Dakota grasshoppers*. Fargo: Dept. of Entomology, Agricultural Experiment Station, 1963.

Oseto, C. Y. and T. J. Helms. “Early embryology and histology of Schizaphis graminum (Hemiptera (Homoptera): Aphididae)”. *Annals of the Entomological Society of America* 65.3 (1972): 622–625.

Osmankhil, M. H., A. Mochizuki, K. Hamasaki, and K. Iwabuchi. “Oviposition and larval development of Neochrysocharis formosa (Hymenoptera: Eulophidae) inside the host larvae, Liriomyza trifolii”. *Japan Agricultural Research Quarterly* 44.1 (2010): 33–36.

Osuji, F. N. C. “Some aspects of the biology of Dermestes maculatus DeGeer (Coleoptera, Dermestidae) in dried fish”. *Journal of Stored Products Research* 11.1 (1975): 25–31.

Ovruski, S. M. “Immature stages of Aganaspis pelleranoi (Brethes) (Hymenoptera: cynipoidea: Eucoilidae), a parasitoid of Ceratitis capitata (Wied.) and Anastrepha spp. (Diptera: Tephritidae)”. *Journal of Hymenoptera Research* 3 (1994): 233–239.

Packauskas, R. J. and J. E. McPherson. “Life history and laboratory rearing of Ranatra fusca (Hemiptera: Nepidae) with descriptions of immature stages”. *Annals of the Entomological Society of America* 79.4 (1986): 566–571.

Padilla-Gil, D. N. “Description of the egg and immature stages of Martarega lofoides Padilla-Gil, 2010 (Hemiptera: Heteroptera: Notonectidae)”. *Zootaxa* 3920.4 (2015): 593–599.

Padilla-Gil, D. N. “Description of the egg and immature stages of Potamobates anchicaya J. Polhemus & D. Polhemus, 1995 (Hemiptera: Heteroptera: Gerridae) and intersexual variation in adults”. *Zootaxa* 3745.5 (2013): 524–532.

Padilla-Gil, D. N. “Immature stages of five species of Gerridae (Heteroptera: Gerromorpha) from the Eastern Tropical Pacific”. *International Journal of Tropical Insect Science* 33.2 (2013): 91–98.

Paim, A. C., L. A. Kaminski, and G. R. P. Moreira. “Morfologia externa dos estágios imaturos de heliconíneos neotropicais. IV. Dryas iulia alcionea (Lepidoptera: Nymphalidae: Heliconiinae)”. *Iheringia, Série Zoologia* 94.1 (2004): 25–35.

Painter, R. R. and W. W. Kilgore. “Some physical and chemical characteristics of normal eggs, larvae, and chorions of the house fly, Musca domestica”. *Annals of the Entomological Society of America* 60.6 (1967): 1163–1166.

Paluch, M., M. M. Casagrande, and O. H. H. Mielke. “Estágios imaturos de Actinote carycina Jordan (Lepidoptera, Nymphalidae, Acraeinae)”. *Revista Brasileira de Zoologia* 18.3 (2001): 883–896.

Papp, L. “Description of the immature stages and the adult female of Aulacigaster africana, the first known for the Afrotropical Aulacigastridae (Diptera: Schizophora)”. *African Invertebrates* 49.2 (2008): 227–232.

Parente, E. “Development of Cryoconservation Technology of Lepidoptera: Study of embryonic development and survival after treating with cryoprotective agents”. Diss. Università degli Studi della Basilicata, 2009.

Park, J. Y. and J. E. Lee. “Immature stages of Paracycnotrachelus longiceps (Motschulsky) (Coleoptera: Attelabidae: Apoderinae) from Korea”. *Animal Systematics, Evolution and Diversity* 20.2 (2004): 225–230.

Park, J. Y. and J. E. Lee. “Immature Stages of Tomapoderus (T.) ruficollis Fabricius (Coleoptera: Attelabidae) from Korea”. *Entomological Research* 34.4 (2004): 225–227.

Parker, H. L. “Recherches les formes postembryonaires de chalcidien”. *Annales de la Société Entomologique de France* 93 (1924): 261–379.

Parkin, E. A. “Observations on the biology of the Lyctus powder-post beetles, with special reference to oviposition and the egg”. *Annals of Applied Biology* 21.3 (1934): 495–518.

Parnell, J. R. “Observations on the larval morphology and life history of Olesicoccus costalimai Borgmeier (Diptera: Cecidomyiidae) in Jamaica”. *Physiological Entomology* 41.4-6 (1966): 51–54.

Parnell, J. R. “The biology and morphology of all stages of Toxomyia fungicola Felt (Diptera: Cecidomyiidae) in Jamaica”. *Physiological Entomology* 44.7-9 (1969): 113–122.

Parnell, J. R. “The parasite complex of the two seed beetles Bruchidius ater (Marsham)(Coleoptera: Bruchidae) and Apion fuscirostre Fabricius (Coleoptera: Curculionidae)*”. *Transactions of the Royal Entomological Society of London* 116.4 (1964): 73–88.

Parra, L. E. and H. Ibarra-Vidal. “Taxonomía y antecedentes biológicos de Microdulia mirabilis (Rothschild 1895) (Lepidoptera: Saturniidae)”. *Gayana* 74.1 (2010): 12–18.

Parrella, M. P. “Biology of Liriomyza”. *Annual Review of Entomology* 32.1 (1987): 201–224.

Paskewitz, S. M. and J. E. McPherson. “Descriptions of nymphal instars of Thyanta calceata (Hemiptera: Pentatomidae)[External anatomy, taxonomy, USA]”. *Great Lakes Entomologist* 15.4 (1982): 227–305.

Paskewitz, S. M. and J. E. McPherson. “Life history and laboratory rearing of Arhyssus lateralis (Hemiptera: Rhopalidae) with descriptions of immature stages”. *Annals of the Entomological Society of America* 76.3 (1983): 477–482.

Passoa, S. “A new clear wing moth (Sesiidae) from Central America: A stem borer in Mimosa pigra”. *The Journal of the Lepidopterists Society* 37.3 (1983): 193–206.

Paterson Fox, E. G., D. R. Solis, M. L. Rossi, W. Mackay, and J. Pacheco. “Morphological Notes on the Worker and Queen Larvae of the Thief Ant Solenopsis helena (Hymenoptera, Formicidae, Myrmicinae) from Brazil”. *The Florida Entomologist* 94.4 (2011): 909–915.

Paterson, N. F. “A Contribution to the Empbryological Development of Euyope terminalis Baly, (Coleoptera, Phytophaga, Chrysomelidae.) Part I:—The Early Embryological Development”. *South African Journal of Science* 28 (1931): 344–371.

Paterson, N. F. “Observations on the Embryology of Corynodes Pusis (Coleoptera, Chrysomelidae)”. *Journal of Cell Science* s2-78.309 (1935): 91–131.

Pearse, A. M. “Aspects of the biology of Uropsylla tasmanica Rothschild (Siphonaptera)”. MA thesis. University of Tasmania, 1981.

Peck, S. B. “The eyeless Catopocerus beetles (Leiodidae) of eastern North America”. *Psyche* 81.3-4 (1974): 377–397.

Percival, E. and H. Whitehead. “Observations on the ova and oviposition of certain Ephemeroptera and Plecoptera”. *Proceedings of the Leeds Philosophical Society (Science Section)* 1.6 (1928): 271–288.

Percy, D. M. “Legume-feeding psyllids (Hemiptera, Psylloidea) of the Canary Islands and Madeira”. *Journal of Natural History* 37.4 (2003): 397–461.

Peredo, L. C. and M. B. Baez. “Life cycle of Balboa variabilis Distant (Hemiptera, Heteroptera, Rhyparochromidae, Rhyparochrominae, Ozophorini)”. *Deutsche Entomologische Zeitschrift* 56.2 (2009): 237–242.

Perez-Banon, C. and A. Marcos-Garcia. “Life history and description of the immature stages of Eumerus purpurariae (Diptera: Syrphidae) developing in Opuntia maxima”. *European Journal of Entomology* 95.3 (1998): 373–382.

Perez-Goodwyn, P. J., S. Ohba, and J. A. Schnack. “Chorion morphology of the eggs of Lethocerus delpontei, Kirkaldyia deyrolli, and Horvathinia pelocoroides (Heteroptera: Belostomatidae)”. *Russian Entomological Journal* 15.2 (2006): 151–156.

Pérez-Lachaud, G., J. M. Heraty, A. Carmichael, and J.-P. Lachaud. “Biology and behavior of Kapala (Hymenoptera: Eucharitidae) attacking Ectatomma, Gnamptogenys, and Pachycondyla (Formicidae: Ectatomminae and Ponerinae) in Chiapas, Mexico”. *Annals of the Entomological Society of America* 99.3 (2006): 567–576.

Perondini, A. L. P., H. O. Gutzeit, and L. Mori. “Nuclear division and migration during early embryogenesis of Bradysia tritici Coquillet (syn. Sciara ocellaris) (Diptera: Sciaridae)”. *International Journal of Insect Morphology and Embryology* 15.3 (1986): 155–163.

Petcu, I. and A. N. A. Davideanu. “Some observations upon the eggs of aquatic bugs (Heteroptera, Cryptocerata)”. *Analele Științifice Ale Universității ”Al. I. Cuza”, din Iași: Biologie Animală* 38 (1993): 33–35.

Peter, C. and B. V. David. “Biology of Goniozus sensorius Gordh (Hymenoptera: Bethylidae) a parasitoid of the pumpkin caterpillar, Diaphania indica (Saunders) (Lepidoptera: Pyralidae)”. *International Journal of Tropical Insect Science* 12.4 (1991): 339–345.

Peters, W. and J. Spurgeon. “Biology of the water-boatman Krizousacorixa femorata (Heteroptera: Corixidae)”. *American Midland Naturalist* 86.1 (1971): 197–207.

Peterson, A. “Egg types among moths of the Noctuidae (Lepidoptera)”. *The Florida Entomologist* 47.2 (1964): 71–91.

Peterson, A. “Egg types among moths of the Pyralidae and Phycitidae-Lepidoptera”. *The Florida Entomologist* 46 (1963): 1–14.

Peterson, A. “Eggs of Moths from Additional Species of Geometridae: Lepidoptera”. *The Florida Entomologist* 51.2 (1968): 83–94.

Peterson, A. “Some eggs of moths among the Amatidae, Arctiidae, and Notodontidae: Lepidoptera”. *The Florida Entomologist* 46.2 (1963): 169–182.

Peterson, A. “Some eggs of moths among the Liparidae, Lasiocampidae, and Lacosomidae (Lepidoptera)”. *The Florida Entomologist* 49.1 (1966): 35–42.

Peterson, A. “Some eggs of moths among the Sphingidae, Saturniidae, and Citheroniidae (Lepidoptera)”. *The Florida Entomologist* 48.4 (1965): 213–219.

Peterson, A. “Some eggs of moths from several families of microlepidoptera”. *The Florida Entomologist* 50.2 (1967): 125–132.

Peterson, A. “Some types of eggs deposited by moths, Heterocera-Lepidoptera”. *The Florida Entomologist* 44.3 (1961): 107–114.

Peterson, R. D. and S. M. Newman. “Chorionic structure of the egg of the screwworm, Cochliomyia hominivorax (Diptera: Calliphoridae)”. *Journal of Medical Entomology* 28.1 (1991): 152–160.

Petralia, R. S. and S. B. Vinson. “Developmental morphology of larvae and eggs of the imported fire ant, Solenopsis invicta”. *Annals of the Entomological Society of America* 72.4 (1979): 472–484.

Petrice, T. R., R. A. Haack, J. S. Strazanac, and J. P. Lelito. “Biology and larval morphology of Agrilus subcinctus (Coleoptera: Buprestidae), with comparisons to the emerald ash borer, Agrilus planipennis”. *Great Lakes Entomologist* 43 (2009): 172–184.

Peyron, J. *Zur morphologie der Skandinavischen schmetterlingseier*. Uppsala: Almqvist & Wiksell, 1909.

Pfaffenberger, G. S. “Morphology and biology of larval Gibbobruchus mimus (Say) (Coleoptera: Bruchidae)”. *The Coleopterists' Bulletin* 40.1 (1986): 49–61.

Pfaffenberger, G. S., S. M. De L'Argentier, and A. L. Teran. “Morphological descriptions and biological and phylogenetic discussions of the first and final instars of four species of Megacerus larvae (Coleoptera: Bruchidae)”. *The Coleopterists' Bulletin* 38.1 (1984): 1–26.

Phillips, J. H. H. “Description of the immature stages of Pulvinaria vitis (L.) and P. innumerabilis (Rathvon) (Homoptera: Coccoidea), with notes on the habits of these species in Ontario, Canada”. *The Canadian Entomologist* 94.5 (1962): 497–502.

Picker, M. D. “Embryonic Development of Heel-walkers: Reference to Some Prerevolutionary Stages (Insecta: Mantophasmatodea)”. *Proceedings of Arthropodan Embryological Society of Japan* 39 (2004): 31–39.

Pickford, R. “Life history and behaviour of Scelio calopteni Riley (Hymenoptera: Scelionidae), a parasite of grasshopper eggs”. *The Canadian Entomologist* 96.9 (1964): 1167–1172.

Pietrykowska-Truduj, E. and B. Staniec. “Description of the egg and larva of Philonthus punctus (Gravenhorst, 1802) (Coleoptera, Staphylinidae, Staphylininae)”. *Deutsche Entomologische Zeitschrift* 53.2 (2006): 179–192.

Pietrykowska-Tudruj, E. and B. Staniec. “Morphology of the developmental stages of Hypnogyra angularis (Ganglbauer, 1895) (Coleoptera, Staphylinidae, Staphylininae)”. *Deutsche Entomologische Zeitschrift* 53.1 (2006): 70–85.

Pietrykowska-Tudruj, E., B. Staniec, T. Wojas, and A. Solodovnikov. “Immature stages and phylogenetic importance of Astrapaeus, a rove beetle genus of puzzling systematic position (Coleoptera, Staphylinidae, Staphylinini)”. *Contributions to Zoology* 83.1 (2014): 41–65.

Pietrykowska, E. “Morphology of the egg and first instar larva of Coptocephala rubicunda (Laicharting, 1781) and notes on its biology (Coleoptera: Chrysomelidae)”. *Genus* 11.1 (2000): 37–44.

Pikart, T. G., G. K. Souza, T. V. Zanuncio, J. C. Zanuncio, and J. E. Serrão. “Eggshell structure of the predator Harpactor angulosus (Hemiptera: Reduviidae)”. *Annals of the Entomological Society of America* 105.6 (2012): 896–901.

Pinto, J. D. and M. A. Bologna. “The first-instar larvae of Meloe afer and M. occultus, with a clarification of antennal structure in larval Meloe (Coleoptera: Meloidae)”. *The Coleopterists' Bulletin* 47.4 (1993): 340–348.

Piper, G. L. “Biology and immature stages of Cylindrocopturus quercus (Say) (Coleoptera: Curculionidae)”. *The Coleopterists' Bulletin* 31.1 (1977): 65–72.

Pires, E. M., P. S. F. Ferreira, R. N. C. Guedes, and J. E. Serrão. “Life stages, biological aspects and geographic distribution of Platyscytus decempunctatus (Heteroptera: Miridae: Phylinae)”. *Revista Brasileira de Biociências* 8.2 (2010): 139–148.

Pizarro-Araya, J., V. Jerez, and J. Cepeda-Pizarro. “Reproducción y ultraestructura del huevo y larva de primer estadio de Gyriosomus kingi (Coleoptera: Tenebrionidae) del desierto de Atacama”. *Revista de Biología Tropical* 55.2 (2007): 637–644.

Plachter, H. “Chorionic structures of the eggshells of 15 fungus-and root-gnat species (Diptera: Mycetophiloidea)”. *International Journal of Insect Morphology and Embryology* 10.1 (1981): 43–63.

Plaut, H. N. “On the biology of the immature stages of the almond wasp, Eurytoma amygdali End.(Hym. Eurytomidae) in Israel”. *Bulletin of Entomological Research* 61.4 (1972): 681–687.

Pljushch, I. G. and I. V. Dolinskaya. “Eggshell fine structure of some species of Lithosiinae (Arctiidae) of Far East Russia”. *Nota Lepidopterologica* 23.1 (2000): 50–63.

Pljushch, I. G. and I. V. Dolinskaya. “The features of chorionic surface structures of some species of Saturniidae, Brahmaeidae and Sphingidae of far east Russia (Lepidoptera, Bombycoidea)”. *Lambillionea* 101 (2001): 11–22.

Podoler, H., Z. Mendel, and H. Livne. “Studies on the biology of a bark beetle predator, Aulonium ruficorne (Coleoptera: Colydiidae)”. *Environmental Entomology* 19.4 (1990): 1010–1016.

Poinar Jr, G. “A walking stick, Clonistria dominicana n. sp. (Phasmatodea: Diapheromeridae) in Dominican amber”. *Historical Biology* 23.02-03 (2011): 223–226.

Polidori, C., R. H. L. Disney, and F. Andretti. “Some observations on the reproductive biology of the scuttle fly Megaselia andrenae (Diptera: Phoridae) at the nesting site of its host Andrena agilissima (Hymenoptera: Andrenidae)”. *European Journal of Entomology* 101 (2004): 337–340.

Polilov, A. A. and R. G. Beutel. “Developmental stages of the hooded beetle Sericoderus lateralis (Coleoptera: Corylophidae) with comments on the phylogenetic position and effects of miniaturization”. *Arthropod Structure & Development* 39.1 (2010): 52–69.

Pollo, P., C. Greve, and V. C. Matesco. “Description of the immature stages of Glyphepomis spinosa Campos & Grazia (Hemiptera: Pentatomidae: Pentatominae: Carpocorini)”. *Zootaxa* 3566 (2012): 61–68.

Portman, S. L. “Foraging and Fecundity of Larra bicolor (Hymenoptera: Sphecidae) a Parasitoid of Scapteriscus Mole Crickets”. MA thesis. University of Florida, 2007.

Potter, D. A. “Effect of soil moisture on oviposition, water absorption, and survival of southern masked chafer (Coleoptera: Scarabaeidae) eggs”. *Environmental Entomology* 12.4 (1983): 1223–1227.

Pourian, H.-R., R. Talaei-Hassanloui, A. Ashouri, H.-A. Lotfalizadeh, and J. Nozari. “Ontogeny and reproductive biology of Diadegma semiclausum (Hym.: Ichneumonidae), a larval endoparasitoid of Diamondback Moth, Plutella xylostella (Lep.: Plutellidae)”. *Arthropod Structure & Development* 44.1 (2015): 69–76.

Powell, J. A. “Biology and immature stages of Australian ethmiid moths (Gelechioidea)”. *Journal of Research on the Lepidoptera* 20.4 (1981): 214–234.

Powell, J. A. “Taxonomy and geographical relationships of Australian Ethmiid moths (Lepidoptera: Gelechioidea)”. *Australian Journal of Zoology* 33.112 (1985): 1–58.

Praz, C. J., H. Özbek, A. Monfared, C. Sedivy, J. G. Rozen Jr, J. S. Ascher, A. Müller, and J. G. Rozen Jr. “Nests, petal usage, floral preferences, and immatures of Osmia (Ozbekosmia) avosetta (Megachilidae: Megachilinae: Osmiini), including biological comparisons with other Osmiine bees”. *American Museum Novitates* 3680 (2010): 1–22.

Presser, B. D. and C. W. Rutschky. “The embryonic development of the corn earworm, Heliothis zea (Boddie) (Lepidoptera, Phalaenidae)”. *Annals of the Entomological Society of America* 50.2 (1957): 133–164.

Prezoto, F. and N. Gobbi. “Morfometria dos estágios imaturos de Polistes simillimus Zikán 1951 (Hymenoptera: Vespidae)”. *Revista Brasileira Zoociências* 7.1 (2005): 47–54.

Principi, M. M. “Contributi allo studio dei Neurotteri Italiani. V. Ricerche su Chrysopa formosa Brauer e su alcuni suoi parassiti”. *Bollettino dell'Istituto di Entomologia della R. Universita degli Bologna* 16 (1947): 134–175.

Puchkova, L. V. “Eggs of the true bugs (Hemiptera-Heteroptera). I. Coreidae”. *Entomologicheskoe Obozrenie* 38.3 (1959): 634–648.

Puchkova, L. V. “Eggs of the true bugs (Hemiptera-Heteroptera). III. Coreidae (Supplement) IV. Macrocephalidae”. *Entomologicheskoe Obozrenie* 36.1 (1957): 44–58.

Puchkova, L. V. “Eggs of the true bugs (Hemiptera-Heteroptera). VI. Pentatomoidea, 2 Pentatomidae and Plataspidae”. *Entomologicheskoe Obozrenie* 40.1 (1961): 131–143.

Putshkova, L. V. “Eggs of the true bugs (Hemiptera-Heteroptera). II. Lygaeidae”. *Entomologicheskoe Obozrenie* 35.2 (1956): 262–284.

Quednau, F. W. “Notes on the life history and morphology of Chrysocharis laricinellae (Ratzeburg) (Hymenoptera, Eulophidae), a parasite of the larch casebearer (Coleophora laricella [Hiibner])”. *Annals of the Entomological Society of Quebec* 11 (1966): 200–205.

Quednau, F. W. and J. H. Martin. “Descriptions of two new species of Anomalosiphum (Hemiptera: Aphididae, Greenideinae), including a winged ovipara with pedunculate eggs”. *Zoological Journal of the Linnean Society* 146.2 (2006): 239–249.

Quezada, J. R., C. A. Amaya, and L. H. Herman Jr. “Xanthopygus cognatus Sharp (Coleoptera: Staphylinidae), an enemy of the coconut weevil, Rhynchophorus palmarum L. (Coleoptera: Curculionidae) in El Salvador”. *Journal of the New York Entomological Society* 77.4 (1969): 264–269.

Quicke, D. J. “Biology and immature stages of Panteles schnetzeanus (Hymenoptera: Ichneumonidae), a parasitoid of Lampronia fuscatella (Lepidoptera: Incurvariidae)”. *Journal of Natural History* 39.5 (2005): 431–443.

Qureshi, S. R., W.-L. Quan, R.-Q. Zhou, and X.-P. Wang. “Morphology and Development of Immature Stages of Chelonus Murakatae (Hymenoptera: Braconidae), an Endoparasitoid of Chilo Suppressalis”. *Entomological News* 125.4 (2015): 252–259.

Rahman, K. A. and M. A. Latif. “Description, bionomics and control of the giant mealybug, Drosicha stebbingi, Green (Homoptera: Coccidae)”. *Bulletin of Entomological Research* 35.2 (1944): 197–209.

Raigorodschi, R. S., D. S. Rocha, J. Jurberg, and C. Galvao. “Description and ontogenetic morphometrics of eggs and instars of Triatoma costalimai Verano & Galvão, 1959 (Hemiptera: Reduviidae: Triatominae)”. *Zootaxa* 3062 (2011): 13–24.

Raine, J. “Life history and behavior of the bramble leafhopper, Ribautiana tenerrima (H.-S.) (Homoptera: Cicadellidae)”. *The Canadian Entomologist* 92.1 (1960): 10–20.

Raine, J. “Life History of Dasystoma salicellum Hbn. (Lepidoptera: Oecophoridae), a New Pest of Blueberries in British Columbia”. *The Canadian Entomologist* 98.3 (1966): 331–334.

Rajaei, H., C. Greve, H. Letsch, D. Stüning, N. Wahlberg, J. Minet, and B. Misof. “Advances in Geometroidea phylogeny, with characterization of a new family based on Pseudobiston pinratanai (Lepidoptera, Glossata)”. *Zoologica Scripta* 44.4 (2015): 418–436.

Rajasekhara, K. and S. Chatterji. “Biology of Orius indicus (Hemiptera: Anthocoridae), a predator of Taeniothrips nigricornis (Thysanoptera)”. *Annals of the Entomological Society of America* 63.2 (1970): 364–367.

Rajotte, E. G. “Nesting, foraging and pheromone response of the bee Colletes validus Cresson and its association with lowbush blueberries. (Hymenoptera: Colletidae) (Ericaceae: Vaccinium)”. *Journal of the Kansas Entomological Society* 52.2 (1979): 349–361.

Rakshpal, R. “Morphogenesis and embryonic membranes of Gryllus assimilis (Fabricius) (Orthoptera: Gryllidae)”. *Physiological Entomology* 37.1-3 (1962): 1–12.

Raminani, L. N. and E. W. Cupp. “Early embryology of Aedes aegypti (L.) (Diptera: Culicidae)”. *International Journal of Insect Morphology and Embryology* 4.6 (1975): 517–528.

Ramos Lacau, L. D. S., C. Villemant, O. C. Bueno, J. H. C. Delabie, and S. Lacau. “Morphology of the eggs and larvae of Cyphomyrmex transversus Emery (Formicidae: Myrmicinae: Attini) and a note on the relationship with its symbiotic fungus”. *Zootaxa* 1923 (2008): 37–54.

Ramos, K. S. and J. G. Rozen Jr. “Psaenythisca, a New Genus of Bees from South America (Apoidea: Andrenidae: Protandrenini) with a Description of the Nesting Biology and Immature Stages of One Species”. *American Museum Novitates* 3800 (2014): 1–32.

Rampini, M. and G. Saltini. “Observations on the egg ultrastructure of some Rhaphidophoridae (Orthoptera) of the mediterranean area”. *Italian Journal of Zoology* 61.1 (1994): 1–8.

Ranasinghe, M., H. A. Denmark, and R. C. Wilkinson. “Immature stages of Gnophothrips fuscus and methods for distinguishing its adults from those of Leptothrips pini (Thysanoptera: Phlaeothripidae)”. *The Florida Entomologist* 68.4 (1985): 594–608.

Rasmi, S. and M. H. B. Hamouda. “Life cycle of immature stages of Oryctes agamemnon arabicus Fairmaire (1896) (Coleoptera: Scarabaeidae) under similar natural conditions of southwest Tunisia”. *Pakistan Entomologist* 37.2 (2015): 83–89.

Ratanov, K. N. “Description of the egg-pods of Acrididae”. *Bulletin of the West Siberian Plant Protection Station* 1.9 (1935): 40–70.

Ratcliffe, B. C. “The natural history of Necrodes surinamensis (Fabr.) (Coleoptera: Silphidae)”. *Transactions of the American Entomological Society (1890-)* 98.4 (1972): 359–410.

Rathod, M. K., M. M. Rai, A. M. Khurad, and S. K. Raina. “SEM studies and re-description of Aleurocanthus husaini (Corbett) (Hemiptera: Aleyrodidae) infesting citrus in Central India”. *Entomon* 38.4 (2013): 213–220.

Rattu, R. “Osservazioni sulla biologia di Cebrio sardous perris, 1869 (Insecta, Coleoptera, Elateridae, Cebrioninae)”. *Bollettino del Museo di Storia Naturale di Venezia* 63 (2012): 45–50.

Readio, P. A. “Studies on the biology of the genus Corizus (Coreidae, Hemiptera)”. *Annals of the Entomological Society of America* 21.2 (1928): 189–201.

Rechav, Y. and T. Orion. “The development of the immature stages of Chelonus inanitus”. *Annals of the Entomological Society of America* 68.3 (1975): 457–462.

Reddy, G. V. P., Z. T. Cruz, and R. Muniappan. “Life-history, host preference and establishment status of Melittia oedipus (Lepidoptera: Sesiidae), a biological control agent for Coccinia grandis (Cucurbitaceae) in the Mariana Islands”. *Plant Protection Quarterly* 24.1 (2009): 27–31.

Reed, E. M. and M. F. Day. “Embryonic movement during development of the light brown apple moth”. *Australian Journal of Zoology* 14.2 (1966): 253–263.

Regier, J. C., G. D. Mazur, and F. C. Kafatos. “The silkmoth chorion: morphological and biochemical characterization of four surface regions”. *Developmental Biology* 76.2 (1980): 286–304.

Reinhardt, K. and U. Gerighausen. “Oviposition site preference and egg parasitism in Sympecma paedisca (Odonata: Lestidae)”. *International Journal of Odonatology* 4.2 (2001): 221–230.

Remadevi, O. K., U. V. K. Mohamed, and U. C. Abdurahiman. “Some aspects of the biology of Parasierola nephantidis Mvesebeck (Hymenoptera, Bethylidae), a larval parasitoid of Nephantis serinopa Meyrick (Lepidoptera, Xylorictidae)”. *Polskie Pismo Entomologiczne* 51.4 (1981): 597–604.

Rempel, J. G. “A study of the embryology of Mamestra configurata (Walker) (Lepidoptera, Phalaenidae)”. *The Canadian Entomologist* 83.1 (1951): 1–19.

Rempel, J. G. and N. S. Church. "The embryology of Lytta viridana Le Conte (Coleoptera: Meloidae): I. Maturation, fertilization, and cleavage". *Canadian Journal of Zoology* 43.6 (1965): 915–925.

Ren, S. *An iconography of Hemiptera-Heteroptera eggs in China*. Beijing: Science Press, 1992.

Resh, V. H. "The biology and immature stages of the caddisfly genus Ceraclea in eastern North America (Trichoptera: Leptoceridae)". *Annals of the Entomological Society of America* 69.6 (1976): 1039–1061.

Rhame, R. E. and K. W. Stewart. "Life cycles and food habits of three Hydropsychidae (Trichoptera) species in the Brazos River, Texas". *Transactions of the American Entomological Society* 102.1 (1976): 65–99.

Ribi, W. A. and L. Ribi. "Natural history of the Australian digger wasp Sphex cognatus Smith (Hymenoptera, Sphecidae)". *Journal of Natural History* 13.6 (1979): 693–701.

Richards, A. M. "A comparative study of the biology of the giant wetas Deinacrida heteracantha and D. fallai (Orthoptera: Henicidae) from New Zealand". *Journal of Zoology* 169.2 (1973): 195–236.

Richter, A., W. Osborne, S. Hnatiuk, and A. Rowell. "Moths in fragments: insights into the biology and ecology of the Australian endangered golden sun moth Synemon plana (Lepidoptera: Castniidae) in natural temperate and exotic grassland remnants". *Journal of Insect Conservation* 17.6 (2013): 1093–1104.

Riddick, E. W. and Z. Wu. "Mother-offspring relations: prey quality and maternal size affect egg size of an acariphagous lady beetle in culture". *Psyche* 2012 (2012): 1–7.

Riemann, J. G. "The development of eggs of the screw-worm fly Cochliomyia hominivorax (Coquerel) (Diptera: Calliphoridae) to the blastoderm stage as seen in whole-mount preparations". *The Biological Bulletin* 129.2 (1965): 329–339.

Rivas, N., M. Sánchez, A. Martínez-Ibarra, A. D. Camacho, A. Tovar-Soto, and R. Alejandre-Aguilar. "Morphological study of eggs from five Mexican species and two morphotypes in the genus Triatoma (Laporte, 1832)". *Journal of Vector Ecology* 38.1 (2013): 90–96.

Rivers, R. L., Z. B. Mayo, and T. J. Helms. "Biology, behavior and description of Tiphia berbereti (Hymenoptera: Tiphiidae) a parasite of Phyllophaga anxia (Coleoptera: Scarabaeidae)". *Journal of the Kansas Entomological Society* 52.2 (1979): 362–372.

Rivnay, E. "Physiological and ecological studies on the species of Capnodis in Palestine (Col., Buprestidae)". *Bulletin of Entomological Research* 35.3 (1945): 235–242.

Robinson, W. H. and B. A. Foote. "Some eggs of moths among the Amatidae, Arctiidae, and Notodontidae: Lepidoptera". *Annals of the Entomological Society of America* 61.6 (1968): 1587–1594.

Rocha, K. L., T. Mangine, E. J. Harris, and P. O. Lawrence. "Immature stages of Fopius arisanus (Hymenoptera: Braconidae) in Bactrocera dorsalis (Diptera: Tephritidae)". *The Florida Entomologist* 87.2 (2004): 164–168.

Rocha, T., J. A. d. O. David, and F. H. Caetano. "Ultramorphological features of the egg of Telmatoscopus albipunctatus (Williston) (Diptera, Psychodidae)". *Revista Brasileira de Entomologia* 55.2 (2011): 179–182.

Rodrigues, D. and G. R. P. Moreira. "Comparative description of the immature stages of two very similar leaf footed bugs, Holymenia clavigera (Herbst) and Anisoscelis foliacea marginella (Dallas) (Hemiptera, Coreidae, Anisoscelini)". *Revista Brasileira de Entomologia* 49.1 (2005): 7–14.

Rodrigues, S. R., G. A. L. Nogueira, R. R. Echeverria, and V. S. Oliveira. "Biological aspects of Cyclocephala verticalis burmeister (coleoptera: scarabaeidae)". *Neotropical Entomology* 39.1 (2010): 15–18.

Rodrigues, S. R., G. A. L. Nogueira, and E. S. Gomes. "Biological Aspects of Liogenys bidenticeps Moser, 1919 (Coleoptera: Scarabaeidae)". *The Coleopterists Bulletin* 68.2 (2014): 235–238.

Rodriguez, M. H., B. Chavez, J. E. Hernandez-Avila, A. Orozco, and J. I. Arredondo-Jimenez. "Description and morphometric analysis of the eggs of Anopheles (Anopheles) vestitipennis (Diptera: Culicidae) from southern Mexico". *Journal of Medical Entomology* 36.1 (1999): 78–87.

Rogo, L. M., E. D. Kokwaro, M. J. Mutinga, and C. P. M. Khamala. "Differentiation of vector species of phlebotominae (Diptera: Psychodidae) in Kenya by chorionic sculpturing of their eggs". *Journal of Medical Entomology* 29.6 (1992): 1042–1044.

Román, L. E. N. “Morphology of the immature stages and biological aspects of Tetrastichus sp. (Hymenoptera: Tetrastichinae), parasitoid of Methona confusa psamathe Godm. et Salv. (Lepidoptera: Ithomiidae)”. *Neotropica* 42 (1996): 41–46.

Roonwal, M. L. “On a new phylogenetically significant ratio (width/length) in termite eggs (Isoptera)”. *Zoologischer Anzeiger* 195.1/2 (1975): 43–50.

Roonwal, M. L. “Size, sculpturing, weight and moisture content of the developing eggs of the desert locust, Schistocerca gregaria (Forskal) (Orthoptera, Acrididae)”. *Proceedings National Institute of Sciences India* 20 (1954): 388–398.

Rosa, S. P. “Description of Photuris fulvipes (Blanchard) immatures (Coleoptera, Lampyridae, Photurinae) and bionomic aspects under laboratory conditions”. *Revista Brasileira de Entomologia* 51.2 (2007): 125–130.

Rosa, S. P., C. Costa, and N. Higashi. “New data on the natural history and description of the immatures of Fulgeochlizus bruchi, a bioluminescent beetle from Central Brazil (Elateridae, Pyrophorini)”. *Papéis Avulsos de Zoologia (São Paulo)* 50.41 (2010): 635–641.

Rosel, A. “Oviposition, egg development and other features of the biology of five species of Lyctidae (Coleoptera)”. *Australian Journal of Entomology* 8.2 (1969): 145–152.

Rosen, D. and A. Eliraz. “Biological and systematic studies of developmental stages in Aphytis (Hymenoptera: Aphelinidae). I. Developmental history of Aphytis chilensis Howard”. *Hilgardia* 46.3 (1978): 77–95.

Rosenheim, J. A. “Nesting behavior and bionomics of a solitary ground-nesting wasp, Ammophila dysmica (Hymenoptera: Sphecidae): influence of parasite pressure”. *Annals of the Entomological Society of America* 80.6 (1987): 739–749.

Rosewall, O. W. “Observations on the egg and larval stages of Trogoderma Inclusum Lec. (Dermestidae)”. *Annals of the Entomological Society of America* 31.3 (1938): 381–384.

Ross, D. A. and D. D. Pothecary. “Notes on adults, eggs, and first-instar larvae of Priacma serrata (Coleoptera: Cupedidae)”. *The Canadian Entomologist* 102.3 (1970): 346–348.

Rost, M. M. and I. Poprawa. “Cellularization during embryogenesis in Thermobia domestica (Zygentoma: Lepismatidae)”. *Annals of the Entomological Society of America* 99.3 (2006): 592–597.

Roth, L. M. “Additions to the oöthecae, uricose glands, ovarioles, and tergal glands of Blattaria”. *Annals of the Entomological Society of America* 64.1 (1971): 127–141.

Roth, L. M. “Control of oötheca formation and oviposition in Blattaria”. *Journal of Insect Physiology* 20.5 (1974): 821–844.

Roth, L. M. “Oöthecae of the Blattaria”. *Annals of the Entomological Society of America* 61.1 (1968): 83–111.

Roth, L. M. “Ovarioles of the Blattaria”. *Annals of the Entomological Society of America* 61.1 (1968): 132–140.

Roth, L. M. “Water changes in cockroach oöthecae in relation to the evolution of ovoviviparity and viviparity”. *Annals of the Entomological Society of America* 60.5 (1967): 928–946.

Roth, L. M. and B. Stay. “A comparative study of oöcyte development in false ovoviviparous cockroaches”. *Psyche* 69.4 (1962): 165–208.

Roth, L. M. and B. Stay. “Oöcyte development in Diploptera punctata (Eschscholtz) (Blattaria)”. *Journal of Insect Physiology* 7.3-4 (1961): 186–202.

Rowley, W. A. and D. C. Peters. “Scanning electron microscopy of the eggshell of four species of Diabrotica (Coleoptera: Chrysomelidae)”. *Annals of the Entomological Society of America* 65.5 (1972): 1188–1191.

Roy, A. S. and D. Ghosh. “Scanning electron microscopic and electrophoretic analysis of the eggshell of Gesonula punctifrons (Orthoptera: Acrididae)”. *Entomon* 34.2 (2009): 89–94.

Rozen Jr, J. G. “Biology and immature stages of Moroccan panurgine bees (Hymenoptera, Apoidea)”. *American Museum Novitates* 2457 (1971): 1–37.

Rozen Jr, J. G. “Biology and immature stages of some cuckoo bees belonging to Brachynomadini, with descriptions of two new species (Hymenoptera: Apidae: Nomadinae)”. *American Museum Novitates* 3089 (1994): 1–31.

Rozen Jr, J. G. “Biology and immature stages of the bee Nomioides patruelis (Halictidae: Halictinae: Nomioidini) and of its cleptoparasite, Chiasmognathus pashupati (Apidae: Nomadinae: Ammobatini), with a preliminary phylogeny of the Halictidae based on mature larvae (Apoidea)”. *American Museum Novitates* 3604 (2008): 1–23.

Rozen Jr, J. G. “Biology notes on the bee Andrena accepta Viereck (Hymenoptera, Andrenidae)”. *Journal of the New York Entomological Society* 81.1 (1973): 54–61.

Rozen Jr, J. G. “Biology of the bee Ancylandrena larreae (Andrenidae, Andreninae) and Its cleptoparasite Hexepeolus rhodogyne (Anthophoridae, Nomadinae): With a review of egg deposition in the Nomadinae (Hymenoptera, Apoidea)”. *American Museum Novitates* 3058 (1992): 1–15.

Rozen Jr, J. G. “Biology, immature stages, and phylogenetic relationships of fideline bees, with the description of a new species of Neofidelia (Hymenoptera, Apoidea)”. *American Museum Novitates* 2427 (1970): 1–25.

Rozen Jr, J. G. “Eggs, ovariole numbers, and modes of parasitism of cleptoparasitic bees, with emphasis on Neotropical species (Hymenoptera: Apoidea)”. *American Museum Novitates* 3413 (2003): 1–36.

Rozen Jr, J. G. “Immatures of exomalopsine bees with notes on nesting biology and a tribal key to mature larvae of noncorbiculate, nonparasitic Apinae (Hymenoptera: Apidae)”. *American Museum Novitates* 3726 (2011): 1–52.

Rozen Jr, J. G. “Immatures of the Old World Oil-Collecting Bee Ctenoplectra cornuta (Apoidea: Apidae: Apinae: Ctenoplectrini)”. *American Museum Novitates* 3699 (2010): 1–14.

Rozen Jr, J. G. “Immatures of The Solitary Bee Camptopoeum Friesei (Andrenidae: Panurginae: Panurgini) and of Its Cleptoparasite Parammobatodes Minutus (Apidae: Nomadinae: Ammobatini)”. *American Museum Novitates* 3641 (2009): 1–14.

Rozen Jr, J. G. “Life history and immature stages of the bee Neofidelia (Hymenoptera, Fideliidae)”. *American Museum Novitates* 2519 (1973): 1–14.

Rozen Jr, J. G. “Mature Larvae of Calliopsine Bees: Spinoliella, Callonychium, and Arhysosage Including Biological Notes, and a Larval Key to Calliopsine Genera (Hymenoptera: Apoidea: Andrenidae: Panurginae)”. *American Museum Novitates* 27.3782 (2013): 1–27.

Rozen Jr, J. G. “Nesting biologies and immature stages of the rophitine bees (Halictidae) with notes on the cleptoparasite Biastes (Anthophoridae) (Hymenoptera: Apoidea)”. *American Museum Novitates* 3066 (1993): 1–28.

Rozen Jr, J. G. “Nesting biology and immature stages of a new species in the bee genus Hesperapis (Hymenoptera, Apoidea, Melittidae, Dasypodinae)”. *American Museum Novitates* 2887 (1987): 1–20.

Rozen Jr, J. G. “Nesting biology and immature stages of the panurgine bee genera Rhophitulus and Cephalurgus (Apoidea, Andrenidae, Protandrenini)”. *American Museum Novitates* 3814 (2014): 1–16.

Rozen Jr, J. G. “Ovarian formula, mature oocyte, and egg index of the bee Ctenoplectra (Hymenoptera: Apoidea: Apidae)”. *Journal of the Kansas Entomological Society* 76.4 (2003): 640–642.

Rozen Jr, J. G. “Ovarioles and oocytes of two Old World cleptoparasitic bees with biological notes on Ammobatoides (Hymenoptera, Apidae)”. *American Museum Novitates* 3326 (2001): 1–12.

Rozen Jr, J. G. “Phylogenetic relationships of Euherbstia with other short-tongued bees (Hymenoptera, Apoidea)”. *American Museum Novitates* 3060 (1993): 1–15.

Rozen Jr, J. G. “The Bee Svastra sabinensis: Nesting Biology, Mature Oocyte, Postdefecating Larva, and Association with Triepeolus penicilliferus (Apidae: Apinae: Eucerini and Nomadinae: Epeolini)”. *American Museum Novitates* 3850 (2016): 1–12.

Rozen Jr, J. G. “The biology and description of a new species of African Thyreus, with life history notes on two species of Anthophora (Hymenoptera: Anthophoridae)”. *Journal of the New York Entomological Society* 77.1 (1969): 51–60.

Rozen Jr, J. G. “The ethology and systematic relationships of fideliine bees, including a description of the mature larva of Parafidelia (Hymenoptera, Apoidea)”. *American Museum Novitates* 2637 (1977): 1–15.

Rozen Jr, J. G. “The natural history of the Old World nomadine parasitic bee Pasites maculatus (Anthophoridae, Nomadinae) and its host Pseudapis diversipes (Halictidae, Nomiinae)”. *American Museum Novitates* 2637 (1986): 1–8.

Rozen Jr, J. G. “Two new species and the redescription of another species of the cleptoparasitic bee genus Triepeolus with notes on their immature stages (Anthophoridae, Nomadinae)”. *American Museum Novitates* 2956 (1989): 1–18.

Rozen Jr, J. G. and S. L. Buchmann. “Nesting biology and immature stages of the bees Centris caesalpiniae, C. pallida, and the cleptoparasite Ericrocis lata (Hymenoptera: Apoidea: Anthophoridae)”. *American Museum Novitates* 2985 (1990): 1–30.

Rozen Jr, J. G. and M. S. Favreau. “Biological notes on Dioxys pomonae pomonae and on its host, Osmia nigrobarbata (Hymenoptera: Megachilidae)”. *Journal of the New York Entomological Society* 75.4 (1967): 197–203.

Rozen Jr, J. G. and H. H. Go. “Descriptions of the egg and mature larva of the bee Chelostoma (Prochelostoma) philadelphi with additional notes on nesting Biology (Hymenoptera: Megachilidae: Megachilinae: Osmiini)”. *American Museum Novitates* 3844 (2015): 1–8.

Rozen Jr, J. G. and H. G. Hall. “Nest site selection and nesting behavior of the bee Lithurgopsis apicalis (Megachilidae, Lithurginae).” *American Museum Novitates* 3796 (2014): 1–24.

Rozen Jr, J. G. and H. G. Hall. “Nesting and developmental biology of the cleptoparasitic bee Stelis ater (Anthidiini) and its host, Osmia chalybea (Osmiini) (Hymenoptera: Megachilidae)”. *American Museum Novitates* 3707 (2011): 1–38.

Rozen Jr, J. G. and H. G. Hall. “Nesting biology and immatures of the oligolectic bee Trachusa larreae (Apoidea: Megachilidae: Anthidiini)”. *American Museum Novitates* 3765 (2012): 1–24.

Rozen Jr, J. G. and N. R. Jacobson. “Biology and Immature Stages of Macropis nuda, Including Comparisons to Related Bees (Apoidea, Melittidae)”. *American Museum Novitates* 2702 (1980): 1–11.

Rozen Jr, J. G. and S. M. Kamel. “Hospicidal behavior of the cleptoparasitic bee Coelioxys (Allocoelioxys) coturnix, including descriptions of its larval instars (Hymenoptera: Megachilidae)”. *American Museum Novitates* 3636 (2008): 1–15.

Rozen Jr, J. G. and S. M. Kamel. “Hospicidal behavior of the cleptoparasitic wasp Sapyga luteomaculata and investigation into ontogenetic changes in its larval anatomy (Hymenoptera: Vespoidea: Sapygidae)”. *American Museum Novitates* 3644 (2009): 1–24.

Rozen Jr, J. G. and S. M. Kamel. “Investigations on the biologies and immature stages of the cleptoparasitic bee genera Radoszkowskiana and Coelioxys and their Megachile hosts (Hymenoptera: Apoidea: Megachilidae: Megachilini)”. *American Museum Novitates* 3573 (2007): 1–43.

Rozen Jr, J. G. and S. M. Kamel. “Last larval instar and mature oocytes of the Old World cleptoparasitic bee Stelis murina, including a review of Stelis biology (Apoidea: Megachilidae: Megachilinae: Anthidiini)”. *American Museum Novitates* 3666 (2009): 1–19.

Rozen Jr, J. G. and R. J. McGinley. “Biology and larvae of the cleptoparasitic bee Townsendiella pulchra and nesting biology of its host Hesperapis larreae (Hymenoptera: Apoidea)”. *American Museum Novitates* 3005 (1991): 1–11.

Rozen Jr, J. G., G. A. R. Melo, A. J. C. Aguiar, and I. Alves-dos-Santos. “Nesting biologies and immature stages of the tapinotaspidine bee genera Monoeca and Lanthanomelissa and of their osirine cleptoparasites Protosiris and Parepeolus (Hymenoptera: Apidae: Apinae)”. *American Museum Novitates* 3501 (2006): 1–60.

Rozen Jr, J. G. and C. D. Michener. “Nests and Immature Stages of the Bee Paratetrapedia Swainsonae (Hymenoptera, Anthophoridae)”. *American Museum Novitates* 2909 (1988): 1–17.

Rozen Jr, J. G. and H. Özbek. “Immature stages of the cleptoparasitic bee Dioxys cincta (Apoidea: Megachilidae: Megachilinae: Dioxyini)”. *American Museum Novitates* 3443 (2004): 1–12.

Rozen Jr, J. G. and H. Özbek. “Notes on the egg and egg deposition of the cleptoparasite Thyreus ramosus (Hymenoptera: Apidae: Melectini)”. *Journal of the Kansas Entomological Society* 78.1 (2005): 34–40.

Rozen Jr, J. G. and H. Özbek. “Oocytes, eggs, and ovarioles of some long-tongued bees (Hymenoptera: Apoidea)”. *American Museum Novitates* 3393 (2003): 1–35.

Rozen Jr, J. G., H. Özbek, J. S. Ascher, and M. G. Rightmyer. "Biology of the bee Hoplitis (Hoplitis) monstrabilis Tkalců and descriptions of its egg and larva (Megachilidae, Megachilinae, Osmiini)". *American Museum Novitates* 3645 (2009): 1–12.

Rozen Jr, J. G. and F. D. Parker. "Nesting biology of the bee Ashmeadiella holtii and its cleptoparasite, a new species of Stelis (Apoidea, Megachilidae)". *American Museum Novitates* 2900 (1987): 1–10.

Rozen Jr, J. G. and A. Roig-Alsina. "Biology, larvae, and oocytes of the parasitic bee tribe Caenoprosopidini (Hymenoptera, Anthophoridae, Nomadinae)". *American Museum Novitates* 3004 (1991): 1–10.

Rozen Jr, J. G., A. Roig-Alsina, and B. A. Alexander. "The cleptoparasitic bee genus Rhopalolemma, with reference to other Nomadinae (Apidae), and biology of its host Protodufourea (Halictidae: Rophitinae)". *American Museum Novitates* 3194 (1997): 1–28.

Rozen Jr, J. G. and L. Ruz. "South American panurgine bees (Andrenidae, Panurginae). Part II. Adults, immature stages, and biology of Neffapis longilingua, a new genus and species with an elongate glossa". *American Museum Novitates* 3136 (1995): 1–15.

Rozen Jr, J. G., J. Straka, and K. Rezkova. "Oocytes, larvae, and cleptoparasitic behavior of Biastes emarginatus (Hymenoptera: Apidae: Nomadinae: Biastini)". *American Museum Novitates* 3667 (2009): 1–15.

Rozen Jr, J. G., S. B. Vinson, R. Coville, and G. Frankie. "Biology and morphology of the immature stages of the cleptoparasitic bee Coelioxys chichimeca (Hymenoptera: Apoidea: Megachilidae)". *American Museum Novitates* 3679 (2010): 1–26.

Rozen Jr, J. G., S. B. Vinson, R. Coville, and G. Frankie. "Biology of the Cleptoparasitic Bee Mesoplia sapphirina (Ericrocidini) and Its Host Centris flavofasciata (Centridini) (Apidae: Apinae)". *American Museum Novitates* 3723 (2011): 1–36.

Rozen Jr, J. G. and E. S. Wyman. "Early nesting biology of the wood-nesting adventive bee, Lithurgus chrysurus Fonscolombe (Apoidea: Megachilidae: Lithurginae)". *American Museum Novitates* 3804 (2014): 1–12.

Rozen Jr, J. G. and E. S. Wyman. "The Chilean Bees Xeromelissa nortina and X. sielfeldi: Their Nesting Biologies and Immature Stages, Including Biological Notes on X. rozeni (Colletidae: Xeromelissinae)". *American Museum Novitates* 3838 (2015): 1–20.

Rozen Jr, J. G., D. Yanega, G. W. Byers, R. H. Hagen, and R. W. Brooks. "Nesting biology and immature stages of the South American bee genus Acamptopoeum (Hymenoptera: Andrenidae: Panurginae)". *Entomological Contributions in Memory of Byron A. Alexander*. Lawrence: Natural History Museum, the University of Kansas, 1999. 59–67.

Ruberson, J. R., J. R. Larsen, and C. D. Jorgensen. "Embryogenesis of the codling moth, Cydia pomonella (Lepidoptera: Tortricidae)". *Annals of the Entomological Society of America* 80.5 (1987): 561–570.

Ruberson, J. R., C. A. Tauber, and M. J. Tauber. "Development and survival of Telenomus lobatus, a parasitoid of chrysopid eggs: effect of host species". *Entomologia Experimentalis et Applicata* 51.2 (1989): 101–106.

Ruf, M. L. L. "Notas sobre Naucoroidea (Hemiptera: Naucoridae). 3ra. Serie. Estudios con microscopio electrónico de barrido: corion de los huevos de Ambrysus (Ambrysus) attenuatus Montandon, Ambrysus (Ambrysus) acutangulus Montandon y Ambrysus (Ambrysus) stali La Rivers". *Lundiana* 8.1 (2007): 9–12.

Rugg, D. and A. Rose. "Reproductive Biology of Some Australian Cockroaches (Blattodea: Blaberidae)". *Journal of the Australian Entomological Society* 23.2 (1984): 113–117.

Rust, R. W. and R. W. Thorp. "The biology of Stelis chlorocyanea, a parasite of Osmia nigrifrons (Hymenoptera: Megachilidae)". *Journal of the Kansas Entomological Society* 46.4 (1973): 548–562.

Rust, R., P. Torchio, and G. Trostle. "Late embryogenesis and immature development of Osmia rufa cornigera (Rossi) (Hymenoptera: Megachilidae)". *Apidologie* 20.4 (1989): 359–367.

Ruz, L. and J. G. Rozen. "South American panurgine bees (Apoidea, Andrenidae, Panurginae). Part 1, Biology, mature larva, and description of a new genus and species". *American Museum Novitates* 3057 (1993): 1–12.

Ruzicka, J. "The immature stages of central European species of Nicrophorus (Coleoptera, Silphidae)". *Acta Entomologica Bohemoslovaca* 89 (1992): 113–135.

Ryan, R. B. “Contribution to the embryology of Coeloides brunneri (Hymenoptera: Braconidae)”. *Annals of the Entomological Society of America* 56.5 (1963): 639–648.

Saakyan-Baranova, A. A. “Morphological study of preimaginal stages of six species of the genus Trichogramma Westwood (Hymenoptera, Trichogrammatidae)”. *Entomologicheskoe Obozrenie* 69.2 (1990): 257–263.

Sacchi, L., C. A. Nalepa, E. Bigliardi, M. Lenz, C. Bandi, S. Corona, A. Grigolo, S. Lambiase, and U. Laudani. “Some aspects of intracellular symbiosis during embryo development of Mastotermes darwiniensis (Isoptera: Mastotermitidae)”. *Parassitologia* 40.3 (1998): 309–316.

Sagliocco, J.-L. and J. B. Coupland. “Biology and host specificity of Chamaesphecia mysiniformis (Lepidoptera: Sesiidae), a potential biological control agent of Marrubium vulgare (Lamiaceae) in Australia”. *Biocontrol Science and Technology* 5.4 (1995): 509–516.

Sahlén, G. “Transmission electron microscopy of the eggshell in five damselflies (Zygoptera: Coenagrionidae, Megapodagrionidae, Calopterygidae)”. *Odonatologica* 24.3 (1995): 311–318.

Sahlén, G. and F. Suhling. “Relationships between egg size and clutch size among European species of Sympetrinae (Odonata: Libellulidae)”. *International Journal of Odonatology* 5.2 (2002): 181–191.

Saini, E. D. “Identificación de los huevos de pentatomidos (Heteroptera) encontrados en cultivos de soja”. *Idia* 425/428 (1984): 79–84.

Saito, S. “A study on the development of the tusser worm, Antheraea pernyi Guer”. *Journal of the Faculty of Agriculture, Hokkaido Imperial University* 33.4 (1934): 249–266.

Salas-Araiza, M. D., W. P. Mackay, J. Valdez-Carrasco, E. Salazar-Solís, and O. A. Martínez-Jaime. “Characterization and comparison of the eggs of seven species of Mexican grasshoppers”. *Southwestern Entomologist* 38.2 (2013): 267–274.

Salkeld, E. H. “A catalogue of the eggs of some Canadian Geometridae (Lepidoptera), with comments”. *Memoirs of the Entomological Society of Canada* 115.S126 (1983): 3–271.

Salkeld, E. H. “Biosystematics of the genus Euxoa (Lepidoptera: Noctuidae): IV. Eggs of the subgenus Euxoa Hbn.” *The Canadian Entomologist* 107.11 (1975): 1137–1152.

Salkeld, E. H. “Microtype eggs of some Tachinidae (Diptera)”. *The Canadian Entomologist* 112.1 (1980): 51–83.

Salkeld, E. H. “The chorionic architecture of Zelus exsanguis (Hemiptera: Reduviidae)”. *The Canadian Entomologist* 104.3 (1972): 433–442.

Salkeld, E. H. “The chorionic structure of the eggs of some species of bumblebees (Hymenoptera: Apidae: Bombinae), and its use in taxonomy”. *The Canadian Entomologist* 110.1 (1978): 71–83.

Sallum, M. A. M. and D. C. Flores. “Ultrastructure of the eggs of two species of Anopheles (Anopheles) Meigen (Diptera, Culicidae)”. *Revista Brasileira de Entomologia* 48.2 (2004): 185–192.

San Blas, G. and D. R. Davis. “Redescription of Dicranoses capsulifex Kieffer and Jörgensen (Lepidoptera: Cecidosidae) with description of the immature stages and biology”. *Zootaxa* 3682.2 (2013): 371–384.

Sanchez, M. and A. C. Hendricks. “Life history and secondary production of Cheumatopsyche spp. in a small Appalachian stream with two different land uses on its watershed”. *Hydrobiologia* 354.1-3 (1997): 127–139.

Sánchez, S. L., A. De los Santos, and C. Montes. “Estudio morfológico de los estados preimaginales de Micrositus ulyssiponensis Germ. 1824 (Coleoptera: Tenebrionidae)”. *Anales de Biología* 3 (1985): 95–102.

Sander, K. “Analyse des ooplasmatischen Reaktionssystems von Euscelis plebejus Fall.(Cicadina) durch Isolieren und Kombinieren von Keimteilen”. *Development Genes and Evolution* 151.4 (1959): 430–497.

Sandoval, C. M., E. Nieves, V. M. Angulo, J. A. Rosa, and E. Aldana. “Morphology of the eggs of the genus Belminus (Hemiptera: Reduviidae: Triatominae) by optical and scanning electron microscopy”. *Zootaxa* 2970 (2011): 33–40.

Sanford, K. H. “Eggs and oviposition sites of some predacious mirids on apple trees (Miridae: Hemiptera)”. *The Canadian Entomologist* 96.9 (1964): 1185–1189.

Sanit, S., P. Sribanditmongkol, K. L. Sukontason, K. Moophayak, T. Klong-Klaew, T. Yasanga, and K. Sukontason. “Morphology and identification of fly eggs: application in forensic entomology”. *Tropical Biomedicine* 30.2 (2013): 325–337.

Santana, F. J. “The biology of immature Diptera associated with bacterial decay in the giant saguaro cactus, (Cereus giganteus Engelmann)”. MA thesis. University of Arizona, 1961.

Sarto i Monteys, V., L. Aguilar, M. Saiz-Ardanaz, D. Ventura, and M. Martí. “Comparative morphology of the egg of the castniid palm borer, Paysandisia archon (Burmeister, 1880) (Lepidoptera: Castniidae)”. *Systematics and Biodiversity* 3.2 (2005): 179–201.

Sartori, M., J. G. Peters, and M. D. Hubbard. “A revision of Oriental Teloganodidae (Insecta, Ephemeroptera, Ephemerelloidea)”. *Zootaxa* 1957 (2008): 1–51.

Saska, P. and A. Honek. “Development of the beetle parasitoids, Brachinus explodens and B. crepitans (Coleoptera: Carabidae)”. *Journal of Zoology* 262.1 (2004): 29–36.

Satar, A. and C. Özbay. “Eggs, first instar larvae and distribution of the neuropterids Lertha extensa and L. sheppardi (Neuroptera: Nemopteridae) in south-eastern Turkey”. *Zoology in the Middle East* 32.1 (2004): 91–96.

Satar, A., Z. Suludere, S. Candan, and S. Canbulat. “Morphology and surface structure of eggs and first instar larvae of Croce schmidti (Navás, 1927) (Neuroptera: Nemopteridae)”. *Zootaxa* 1554 (2007): 49–55.

Satchell, G. H. “On the early stages of Bruchomyia argentina Alexander (Diptera: Psychodidae)”. *Proceedings of the Royal Entomological Society of London. Series A, General Entomology* 28.1-3 (1953): 1–12.

Sathish, R., D. J. Naik, and K. Deepika. “Biology of sapota midrib folder, Banisia myrsusalis elearalis Walker (Thyrididae: Lepidoptera) infesting sapota under hill zone of Karnataka”. *Pest Management In Horticultural Ecosystems* 19.2 (2013): 160–163.

Savino, V., C. E. Coviella, and M. G. Luna. “Reproductive biology and functional response of Dineulophus phtorimaeae, a natural enemy of the tomato moth, Tuta absoluta”. *Journal of Insect Science* 12.1 (2012): 1–14.

Scali, V. “Revision of the Iberian stick insect genus Leptynia Pantel and description of the new genus Pijnackeria”. *Italian Journal of Zoology* 76.4 (2009): 381–391.

Scali, V., B. Mantovani, and O. Marescalchi. “Identity between Ramulus libanicus (Uvarov) and Ramulus turcus (Karabağ) (Insecta, Phasmatodea): body, egg and chromosome analysis”. *Zoologica Scripta* 19.1 (1990): 65–72.

Scali, V., B. Mantovani, M. Mazzini, G. Nascetti, and L. Bullini. “Intraspecific ootaxonomy of Bacillus rossius (Rossi) (Insecta Phasmatodea)”. *Italian Journal of Zoology* 54.1 (1987): 41–47.

Scali, V. and M. Mazzini. “Fine morphology and amino acid analysis of the egg capsule of the stick insect, Clonopsis gallica (Charp.) (Cheleutoptera: Bacillinae)”. *International Journal of Insect Morphology and Embryology* 6.5 (1977): 255–264.

Scali, V. and M. Mazzini. “L'uovo di Palophus rothschildi Bolivar al microscopio electtronico a scansiore”. *Bollettino della Società Entomologica Italiana, Genova* 122.2 (1990): 91–101.

Schaefer, C. W. and K. W. Wolf. “Notes on the harpactocorine genus Sinea (Hemiptera: Heteroptera: Reduviidae)”. *Journal of the New York Entomological Society* 111.4 (2003): 227–234.

Schaefer, C. H. “Life history of Conophthorus radiatae (Coleoptera: Scolytidae) and its principal parasite, Cephalonomia utahensis (Hymenoptera: Bethylidae)”. *Annals of the Entomological Society of America* 55.5 (1962): 569–577.

Schmidt-Ott, U., A. M. Rafiqi, K. Sander, and J. S. Johnston. “Extremely small genomes in two unrelated dipteran insects with shared early developmental traits”. *Development Genes and Evolution* 219.4 (2009): 207–210.

Schmidt, D. A. “Description of the immatures of Erichsonius alumnus Frank and E. pusio (Horn) (Coleoptera: Staphylinidae)”. *The Coleopterists' Bulletin* 50.3 (1996): 205–215.

Schmidt, D. A. “Notes on the biology and a description of the egg, third instar larva and pupa of Neobisnius sobrinus (Coleoptera: Staphylinidae)”. *Transactions of the Nebraska Academy of Sciences* 21 (1994): 55–61.

Schmidt, D. A. “Notes on the biology and a description of the egg, third instar larva and pupa of Platydracus tomentosus (Gravenhorst) (Coleoptera: Staphylinidae)”. *The Coleopterists' Bulletin* 48.4 (1994): 310–318.

Schulte, G. G., M. A. Elnitsky, J. B. Benoit, D. L. Denlinger, and R. E. Lee. “Extremely large aggregations of collembolan eggs on Humble Island, Antarctica: a response to early seasonal warming?” *Polar Biology* 31.7 (2008): 889–892.

Schwalm, F. E. and H. A. Bender. "Early development of the kelp fly, Coelopa frigida (Diptera). II. Morphology of cleavage and blastoderm formation". *Journal of Morphology* 141.2 (1973): 235–255.

Schwertner, C. F., G. S. Albuquerque, and J. Grazia. "Description of the Immature Stages of Acrosternum (Chinavia) ubicum Rolston (Heteroptera: Pentatomidae) and Effect of the Host Plant on Size and Coloration of Nymphs". *Neotropical Entomology* 31.4 (2002): 571–579.

Scoble, M. J. and A. Aiello. "Moth-like butterflies (Hedylidae: Lepidoptera): a summary, with comments on the egg". *Journal of Natural History* 24.1 (1990): 159–164.

Scott, R. R. "A study of the biology and population dynamics of Synanthedon tipuliformis (Clerck) (Lepidoptera: Sesiidae) in Canterbury, New Zealand". Diss. University of Canterbury, 1975.

Scott, R. R. and R. A. Harrison. "The biology and life history of currant clearwing, Synanthedon tipuliformis (Lepidoptera: Sesiidae), in Canterbury". *New Zealand Journal of Zoology* 6.1 (1979): 145–163.

Scudder, G. G. E., J. P. E. C. Darlington, and S. B. Hill. "A new species of Lygaeidae (Hemiptera) from the Tamana Caves, Trinidad". *Annales de Speleologie* 22 (1967): 465–469.

Segoli, M., A. Bouskila, A. R. Harari, and T. Keasar. "Developmental patterns in the polyembryonic parasitoid wasp Copidosoma koehleri". *Arthropod Structure&Development* 38.1 (2009): 84–90.

Sehl, A. "Furchung und bildung der keimanlage bei der mehlmotte ephestia kuehniella zell. Nebst einer allgemeinen übersicht uber den verlauf der embryonalentwicklung". *Zeitschrift für Morphologie und Ökologie der Tiere* 20.2-3 (1931): 533–598.

Seidel, F. "Untersuchungen über das Bildungsprinzip der Keimanlage im Ei der Libelle Platycnemis pennipes I–V". *Wilhelm Roux'Archiv für Entwicklungsmechanik der Organismen* 119.1 (1929): 322–440.

Seifert, H. F. "Embryological studies of Thysanoptera". MA thesis. University of Illinois, 1917.

Seko, T. and F. Nakasuji. "Adaptive significance of egg size plasticity in response to temperature in the migrant skipper, Parnara guttata guttata (Lepidoptera: Hesperiidae)". *Population Ecology* 48.2 (2006): 159–166.

Selivon, D. and A. L. P. Perondini. "Description of Anastrepha sororcula and A. serpentina (Diptera: Tephritidae) eggs". *The Florida Entomologist* 82.2 (1999): 347–353.

Sellick, J. T. C. "Descriptive terminology of the phasmid egg capsule, with an extended key to the phasmid genera based on egg structure". *Systematic Entomology* 22.2 (1997): 97–122.

Sellick, J. T. C. "The micropylar plate of the eggs of Phasmida, with a survey of the range of plate form within the order". *Systematic Entomology* 23.3 (1998): 203–228.

Sellick, J. T. C. "The range of egg capsule morphology within the Phasmatodea and its relevance to the taxonomy of the order". *Italian Journal of Zoology* 64.1 (1997): 97–104.

Semelbauer, M. and M. Kozánek. "Immature stages of Meiosimyza Hendel 1925 and related genera (Diptera, Lauxaniidae)". *Organisms Diversity & Evolution* 14.1 (2014): 89–103.

Setty, L. R. "Biology and morphology of some north American Bittacidae (Order Mecoptera)". *The American Midland Naturalist* 23.2 (1940): 257–353.

Setty, L. R. "The Biology of Bittacus Stigmaterus Say (Mecoptera, Bittacusidæ)". *Annals of the Entomological Society of America* 24.3 (1931): 467–484.

Sforza, R., T. Bourgoin, S. W. Wilson, and E. Boudon-Padieu. "Field observations, laboratory rearing and descriptions of immatures of the planthopper Hyalesthes obsoletus (Hemiptera: Cixiidae)". *European Journal of Entomology* 96.4 (1999): 409–418.

Sharma, S., J. S. Tara, and S. Bhatia. "Bionomics of Hyblaea Puera (Lepidoptera: Hyblaeidae), a Serious Pest of Teak (tectona Grandis) from Jammu (india)". *Munis Entomology & Zoology* 8.1 (2013): 139–147.

Sharp, D. "Account of the Phasmidae, with notes on the eggs". *Zoological Results* 2 (1898): 75–94.

Shazli, A. and T. M. Mustafa. "Studies on the morphology and life cycle of Thomasiniana oleisuga Targ. (Dipt., Cecidomyiidae) in Jordan". *Zeitschrift für Angewandte Entomologie* 88.1-5 (1979): 80–87.

She, H. D. N., J. A. Odebiyi, and H. R. Herren. "The biology of Hyperaspis jucunda [Col.: Coccinellidae] an exotic predator of the cassava mealybug Phenacoccus manihoti [Hom.: Pseudococcidae] in southern Nigeria". *Entomophaga* 29.1 (1984): 87–93.

Sheehan, W. “Nesting biology of the sand wasp Stictia heros (Hymenoptera: Sphecidae: Nyssoninae) in Costa Rica”. *Journal of the Kansas Entomological Society* 57.3 (1984): 377–386.

Shelford, V. E. “Life histories and larval habits of the tiger beetles (Cicindelidae)”. *Journal of the Linnean Society of London, Zoology* 30 (1908): 157–184.

Shepard, W. D. and R. W. Baumann. “Calileuctra, a new genus, and two new species of stoneflies from California (Plecoptera: Leuctridae)”. *The Great Basin Naturalist* 55.2 (1995): 124–134.

Shepard, W. D. “Neoeubria inbionis Shepard & Barr, a new genus and new species of Neotropical water penny beetle (Coleoptera: Psephenidae: Eubriinae), with a key to the adult Eubriinae of the Neotropic Zone”. *Zootaxa* 3811.4 (2014): 553–568.

Shields, K. S. and R. J. Pupedis. “Morphology and surface structure of Mantispa sayi (Neuroptera: Mantispidae) eggs”. *Annals of the Entomological Society of America* 90.6 (1997): 810–813.

Shimizu, S. and R. Machida. “Reproductive biology and postembryonic development in the basal earwig Diplatys flavicollis (Shiraki) (Insecta: Dermaptera: Diplatyidae)”. *Arthropod Systematics and Phylogeny* 69.2 (2011): 83–97.

Shin, C., H. Jin, and C. S. Chaboo. “Biology and Morphology of Neochlamisus gibbosus (Fabricius, 1777) (Coleoptera: Chrysomelidae: Cryptocephalinae: Fulcidacini)”. *Journal of the Kansas Entomological Society* 85.2 (2012): 116–134.

Shipp, J. L. “Classification system for embryonic development of Simulium arcticum Malloch (IIS-10.11) (Diptera: Simuliidae)”. *Canadian Journal of Zoology* 66.1 (1988): 274–276.

Shirlee, M., K. Palma, and R. W. Meola. “Flea eggs: Target of the new IGR on–animal treatments”. *Proceedings of the First International Conference on Urban Pests*. Exeter: BPCC Wheatons Ltd, 1993.

Shorthouse, J. D. and J. J. Leggo. “Immature stages of the galler Diplolepis triforma (Hymenoptera: Cynipidae) with comments on the role of its prepupa”. *The Canadian Entomologist* 134.4 (2002): 433–446.

Shuzhi, R. “Fine surface structure of eggs and classification of five species of Coptosoma laporte”. *La Animala Mondo* 2.3-4 (1985): 235–243.

Silva, L., G. Markin, and J. Tavares. “Argyresthia atlanticella Rebel (Insecta: Lepidoptera) an excluded agent for Myrica faya Aiton (Myricaceae) biocontrol”. *Arquipélago: Ciências Biológicas e Marinhas* 30.A (1995): 105–113.

Silvestri, F. “Descrizione di due specie neotropicali di Zorotypus”. *Bollettino del Laboratoria di Entomologia Agraria di Portici* 7 (1947): 1–12.

Simpson, G. B. “Immature stages of Nala lividipes (Dufour) (Dermaptera: Labiduridae)”. *Austral Entomology* 32.1 (1993): 51–57.

Simpson, G. B. “Immature stages of Protaetia fusca (Herbst) (Coleoptera: Scarabaeidae: Cetoniinae) with notes on biology”. *Australian Journal of Entomology* 29.1 (1990): 67–73.

Singh, P., J. Madan, and N. Gupta. “Egg shell morphology of an amblyceran louse, Hohorstiella rampurensis (Phthiraptera) infesting ring dove, Streptopelia decaocta”. *Journal of Applied and Natural Science* 8.1 (2016): 469–472.

Sites, R. W. “Egg ultrastructure and descriptions of nymphs of Pelocoris poeyi (Guérin Méneville) (Hemiptera: Naucoridae)”. *Journal of the New York Entomological Society* 99.4 (1991): 622–629.

Situmorang, J. and B. P. Gabriel. “Biology of two species of predatory earwigs Nala lividipes (Dufour) (Dermaptera: Labiduridae) and Euborellia (Euborellia) annulata (Fabricius) (Dermaptera: Carcinophoridae)”. *Philippine Entomologist* 7.3 (1988): 215–238.

Skrzypczyńska, M. “Megastigmus suspectus Borries, 1895 (Hymenoptera, Torymidae), its morphology, biology and economic significance”. *Zeitschrift für angewandte Entomologie* 85.1-4 (1978): 204–215.

Slater, J. A. “A contribution to the biology of the subfamily Cyminae (Heteroptera: Lygaeidae)”. *Annals of the Entomological Society of America* 45.2 (1952): 315–326.

Slater, J. A. “The immature stages of Lygaeidae (Hemiptera: Heteroptera) of southwest Australia”. *Australian Journal of Entomology* 15.1 (1976): 101–126.

Smereka, E. P. “The life history and habits of Chrysomela crotchi Brown (Coleoptera: Chrysomelidae) in Northwestern Ontario”. *The Canadian Entomologist* 97.5 (1965): 541–549.

Smith, C. C. "The life-history and galls of a spruce gall midge, Phytophaga piceae Felt (Diptera: Cecidomyiidae)". *The Canadian Entomologist* 84.9 (1952): 272–275.

Smith, E. S. C. "Studies on Amblypelta theobromae Brown (Hepteroptera: Coreidae) in Papua New Guinea. I. Descriptions of the immature and adult stages". *Bulletin of Entomological Research* 74.3 (1984): 541–547.

Smith, K. G. V. "The biology and taxonomy of the genus Stylogaster Macquart, 1835 (Diptera: Conopidae, Stylogasterinae) in the Ethiopian and Malagasy regions". *Transactions of the Royal Entomological Society of London* 119.2 (1967): 47–69.

Smith, R. H. "A technique for studying the oviposition habits of the southern lyctus beetle and its egg and early larval stages". *Journal of Economic Entomology* 49.2 (1956): 263–264.

Smith, S. A. and M. E. Clay. "Biological and morphological studies on the bat flea, Myodopsylla insignis (Siphonaptera: Ischnopsyllidae)". *Journal of Medical Entomology* 25.5 (1988): 413–424.

Smith, T. R. "Life History and Pesticide Susceptibility of Cybocephalus Nipponicus Endrödy-Younga (Coleoptera: Cybocephalidae) and a Taxonomic Revision of the Cybocephalidae of North America and the West Indies". Diss. University of Florida, 2006.

Smoleński, M. "Immature stages of Rugilus rufipes Germar (Coleoptera, Staphylinidae), with notes on biology". *Annales Zoologici* 46.3-4-11 (1997): 233–243.

Snyder, T. E. "Egg and manner of oviposition of Lyctus planicollis". *Journal of Agricultural Research* 6.7 (1916): 273–276.

Socha, R. "Altered anteroposterior polarity of micropyle ring formation in the eggs of Pyrrhocoris apterus L. (Heteroptera: Pyrrhocoridae)". *International Journal of Insect Morphology and Embryology* 17.2 (1988): 135–143.

Solis, D. R., E. G. P. Fox, M. L. Rossi, and O. C. Bueno. "Description of the immatures of Linepithema humile Mayr (Hymenoptera: Formicidae)". *Biological Research* 43.1 (2010): 19–30.

Solis, D. R., N. B. Dias, and E. G. P. Fox. "External Morphology of the Immatures of Polybia paulista (Hymenoptera: Vespidae)". *The Florida Entomologist* 95.4 (2012): 890–899.

Solis, D. R., E. G. P. Fox, M. Ceccato, I. C. Reiss, P. Decio, N. Lorenzon, N. G. Da Silva, and O. C. Bueno. "On the morphology of the worker immatures of the leafcutter ant Atta sexdens Linnaeus (Hymenoptera: Formicidae)". *Microscopy Research and Technique* 75.8 (2012): 1059–1065.

Solis, D. R., E. G. P. Fox, L. M. Kato, C. M. de Jesus, A. T. Yabuki, A. E. de Carvalho Campos, and O. C. Bueno. "Morphological description of the immatures of the ant, Monomorium floricola". *Journal of Insect Science* 10.1 (2010): 1–17.

Solis, D. R., E. G. P. Fox, M. L. Rossi, and O. C. Bueno. "Description of the immatures of workers of the Weaver Ant, Camponotus textor (Hymenoptera: Formicidae)". *Sociobiology* 54.2 (2009): 541–559.

Solis, D. R., E. G. P. Fox, M. L. Rossi, and O. C. Bueno. "Compared morphology of the immatures of males of two urban ant species of Camponotus". *Journal of Insect Science* 12.1 (2012): 1–12.

Solis, D. R., E. G. P. Fox, M. L. Rossi, T. D. C. Moretti, and O. C. Bueno. "Description of the immatures of workers of the ant Camponotus vittatus (Hymenoptera: Formicidae)". *The Florida Entomologist* 93.2 (2010): 265–276.

Solis, D. R., M. A. Nakano, E. G. P. Fox, M. L. Rossi, R. M. Feitosa, O. C. Bueno, and M. S. de Castro Morini. "Description of the immatures of the ant, Myrmelachista catharinae". *Journal of Insect Science* 11.1 (2011): 1–9.

Sonan, J. "On the life-history of Hybris subjacens Walker". *Transactions of the Natural History Society of Formosa* 20 (1938): 273–275.

Sonnenblick, B. P. "The early embryology of Drosophila melanogaster". *Biology of Drosophila*. New York: Hafner Pub. Co., 1950. 62–167.

Sosa, A. J., A. M. M. D. R. Lenicov, R. Mariani, and H. A. Cordo. "Life history of Megamelus scutellaris with description of immature stages (Hemiptera: Delphacidae)". *Annals of the Entomological Society of America* 98.1 (2005): 66–72.

Sota, T. and M. Mogi. "Interspecific variation in desiccation survival time of Aedes (Stegomyia) mosquito eggs is correlated with habitat and egg size". *Oecologia* 90.3 (1992): 353–358.

Sottile, L. “Il Corion Delle Uova di Epilachna chrysomelina F. (Coleoptera Coccinellidae)”. *Italian Journal of Zoology* 45.S1 (1978): 49–49.

Sousa, J. M. “Development of Tiphodytes gerriphagus (Hymenoptera: Scelionidae) in Limnoporus dissortis eggs (Hemiptera: Gerridae)”. *The Canadian Entomologist* 131.2 (1999): 219–228.

Southwood, T. R. E. “The structure of the eggs of the terrestrial heteroptera and its relationship to the classification of the group”. *Transactions of the Royal Entomological Society of London* 108.6 (1956): 163–221.

Souza, G. K. “Morfologia de ovos, glândulas salivares e sistemas digestivo e reprodutor de Thaumastocoris peregrinus (Hemiptera: Thaumastocoridae)”. MA thesis. Universidade Federal de Viçosa, 2012.

Souza, T. B., A. C. D. Maia, C. M. R. Albuquerque, and L. Iannuzzi. “Biology and management of the masked chafer Cyclocephala distincta Burmeister (Melolonthidae, Dynastinae, Cyclocephalini)”. *Revista Brasileira de Entomologia* 59.1 (2015): 37–42.

Spangler, P. J. and J. L. Cross. “Description of the egg case and larva of the water scavenger beetle, Helobata striata (Coleoptera: Hydrophilidae)”. *Proceedings of the Biological Society of Washington* 85.35 (1972): 413–418.

Spangler, P. J. “Notes on the biology and distribution of Sperchopsis tessellatus (Ziegler) (Coleoptera: Hydrophilidae)”. *The Coleopterists' Bulletin* 15.4 (1961): 105–112.

Spooner, G. M. “The british species of psenine wasps (Hymenoptera: Sphecidae)”. *Transactions of the Royal Entomological Society of London* 99.3 (1948): 129–172.

Spradbery, J. P. “The biology of Stenogaster concinna Van der Vecht with comments on the phylogeny of Stenogastrinae (Hymenoptera: Vespidae)”. *Australian Journal of Entomology* 14.3 (1975): 309–318.

Spradbery, J. P. “The nesting of Anischnogaster irzdipennzs (Smith) (Hymenoptera: Vespidae) in New Guinea”. *Australian Journal of Entomology* 28.4 (1989): 225–228.

Sreedevi, K. and S. Tyagi. “Diagnostic characters of immature stages of flower chafer beetle, Chiloloba acuta (Wiedemann) (Coleoptera: Scarabaeidae: Cetoniinae): Taxonomic importance”. *Pest Management In Horticultural Ecosystems* 19.2 (2013): 185–190.

Sreedevi, K., S. Tyagi, and V. V. Ramamurthy. “Egg Morphology of Twelve Species of Melolonthinae and Rutelinae (Coleoptera: Scarabaeidae)”. *The Coleopterists Bulletin* 69.3 (2015): 426–434.

Sruoga, V. and A. Diškus. “Stephensia brunnichella (Lepidoptera: Elachistidae) new species for Lithuania”. *Acta Zoologica Lituanica* 11.1 (2001): 73–77.

St. George, R. A. “Egg and first-stage larva of Tarsostenus univittatus (Rossi), a beetle predacious on powder-post beetles”. *Journal of Agricultural Research* 29.1 (1924): 49–51.

Stairs, G. R. “On the embryology of the spruce budworm, Choristoneura fumiferana (Clem.) (Lepidoptera, Tortricidae)”. *The Canadian Entomologist* 92.2 (1960): 147–154.

Staniec, B. “Comparative morphology of eggs and notes on the reproductive period of the fifteen Bledius species collected in Poland (Coleoptera: Staphylinidae)”. *Polskie Pismo Entomologiczne* 69.1 (2000): 31–46.

Staniec, B. and E. Pietrykowska-Tudruj. “Morphology of developmental stages of Philonthus fumarius (Gravenhorst, 1806) (Coleoptera, Staphylinidae) with notes on biology”. *Acta Zoologica Academiae Scientiarum Hungaricae* 54.3 (2008): 213–234.

Staniec, B. “A description of the developmental stages of Acylophorus wagenschieberi Kiesenwetter, 1850 (Coleoptera, Staphylinidae), with comments on its biology, egg parasite and distribution in Poland”. *Deutsche Entomologische Zeitschrift* 52.1 (2005): 97–113.

Staniec, B. “A description of the developmental stages of Aploderus caelatus (Gravenhorst, 1802) (Coleoptera: Staphylinidae)”. *Deutsche Entomologische Zeitschrift* 44.2 (1997): 203–230.

Staniec, B. “A description of the egg and mature larva (L3) of Aploderus caesus (Erichson, 1839) (Coleoptera: Staphylinidae)”. *Genus* 10.3 (1999): 361–370.

Staniec, B. “A Description of the Preimaginal Stages and Notes on the Biology of Bledius nanus Erichson, 1840 (Coleoptera, Staphylinidae)”. *Deutsche Entomologische Zeitschrift* 45.1 (1998): 95–109.

Staniec, B. "Description of the developmental stages of Atanygnathus terminalis (Erichson, 1839) (Coleoptera, Staphylinidae, Staphylininae), with comments on its biology". *Deutsche Entomologische Zeitschrift* 52.2 (2005): 173–190.

Staniec, B. "Description of the developmental stages of Hesperus rufipennis (Gravenhorst, 1802) (Coleoptera: Staphylinidae), with comments on its biology". *Annales Zoologici* 54.3 (2004): 529–539.

Staniec, B. "Description of the egg, larva and pupa of Platystethus alutaceus (Thomson, 1861) (Coleoptera: Staphylinidae)". *Genus* 14.1 (2003): 27–41.

Staniec, B. "Developmental stages of Platystethus nitens (CR Sahlberg, 1832) (Coleoptera: Staphylinidae)". *Genus* 14.3 (2003): 345–355.

Staniec, B. and E. Pietrykowska-Tudruj. "Comparative morphology of the eggs of sixteen Central European species of Staphylininae (Coleoptera, Staphylinidae)". *Deutsche Entomologische Zeitschrift* 54.2 (2007): 235–252.

Staniec, B. and E. Pietrykowska-Tudruj. "Developmental stages of Philonthus rubripennis Stephens, 1832 (Coleoptera, Staphylinidae, Staphylininae) with comments on its biology". *Deutsche Entomologische Zeitschrift* 54.1 (2007): 95–113.

Staniec, B. and E. Pietrykowska-Tudruj. "Immature stages of Rabigus tenuis (Fabricius, 1792) (Coleoptera, Staphylinidae, Staphylininae) with observation on its biology and taxonomic comments". *Belgian Journal of Zoology* 139.1 (2009): 22–39.

Staniec, B. and E. Pietrykowska-Tudruj. "Morphology of the immature stages and notes on biology of Philonthus nigrita (Gravenhorst, 1806) (Coleoptera, Staphylinidae) a stenotopic species inhabiting Sphagnum peatbogs". *Deutsche Entomologische Zeitschrift* 55.1 (2008): 167–183.

Staniec, B., E. Pietrykowska-Tudruj, and D. Sałapa. "Description of the egg and larva of Paederidus Mulsant & Rey, 1878 (Coleoptera, Staphylinidae, Paederinae) based on the two European species". *Zootaxa* 2888 (2011): 39–56.

Staniec, B., J. Pilipczuk, and E. Pietrykowska-Tudruj. "Morphology of immature stages and notes on biology of Ocypus fulvipennis Erichson, 1840 (Coleoptera: Staphylinidae)". *Annales Zoologici* 59.1 (2009): 47–66.

Stanley, M. S. M. and A. W. Grundmann. "The embryonic development of Tribolium confusum". *Annals of the Entomological Society of America* 63.5 (1970): 1248–1256.

Stark, B. P. and S. W. Szczytko. "Egg morphology and classification of Perlodinae (Plecoptera: Perlodidae)". *Annales de Limnologie* 20.1-2 (1984): 99–103.

Stark, B. P. and S. Green. "Eggs of western Nearctic Acroneuriinae (Plecoptera: Perlidae)". *Illiesia* 7.17 (2011): 157–166.

Stark, B. P. and D. L. Lentz. "Morphology of the egg capsule in Megaphasma dentricus (Phasmatodea: Heteronemiidae)". *Journal of the Kansas Entomological Society* 59.2 (1986): 398–401.

Stark, B. P. and S. W. Szczytko. "Egg morphology and phylogeny in Arcynopterygini (Plecoptera: Perlodidae)". *Journal of the Kansas Entomological Society* 61.2 (1988): 143–160.

Stark, B. P. and S. W. Szczytko. "Egg morphology and phylogeny in Pteronarcyidae (Plecoptera)". *Annals of the Entomological Society of America* 75.5 (1982): 519–529.

Starmer, W. T., M. Polak, S. Pitnick, S. F. McEvey, J. S. F. Barker, and L. L. Wolf. "Phylogenetic, geographical, and temporal analysis of female reproductive trade-offs in Drosophilidae". *Evolutionary Biology, Vol. 33*. Boston: Springer, 2003. 139–171.

Starzyk, J. R. and M. Partyka. "Study on the morphology, biology and distribution of Obrium cantharinum (L.) (Col., Cerambycidae)". *Journal of Applied Entomology* 116.1-5 (1993): 333–344.

Stathas, G. J. "Studies on morphology and biology of immature stages of the predator Rhyzobius lophanthae Blaisdell (Col.: Coccinellidae)". *Anzeiger für Schädlingskunde* 74.5 (2001): 113–116.

Stechauner-Rohringer, R. and L. C. Pardo-Locarno. "Redescripción de inmaduros, ciclo de vida, distribución e importancia agrícola de Cyclocephala lunulata Burmeister (Coleóptera: Melolonthidae: Dynastinae) en Colombia". *Boletín Científico Centro de Museos - Museo de Historia Natural* 14.1 (2010): 203–220.

Steiner, F. M., B. C. Schlick-Steiner, H. Höttinger, A. Nikiforov, K. Moder, and E. Christian. "Maculinea alcon and M. rebeli (Insecta: Lepidoptera: Lycaenidae) – one or two alcon blues? Larval cuticular compounds and egg morphology of East Austrian populations". *Annalen des Naturhistorischen Museums in Wien, Serie B für Botanik und Zoologie* 107 B (2005): 165–180.

Stiling, P. D. and D. R. Strong. "A leaf miner (Diptera: Ephydridae) and its parasitoids on Spartina alterniflora in northwest Florida". *The Florida Entomologist* 64.4 (1981): 468–471.

Stokkebo, S. and I. C. W. Hardy. "The importance of being gravid: egg load and contest outcome in a parasitoid wasp". *Animal Behaviour* 59.6 (2000): 1111–1118.

Straka, J. and J. G. Rozen Jr. "First observations on nesting and immatures of the bee genus Ancyla (Apoidea: Apidae: Apinae: Ancylaini)". *American Museum Novitates* 3749 (2012): 1–24.

Strebel, O. "Beiträge zur biologie, ökologie und physiologie einheimischer collembolen". *Zoomorphology* 25.1 (1932): 31–153.

Strickman, D. "Observations on adults and eggs of chaoborus flavicans (Diptera: Chaoboridae)". *Hydrobiologia* 74.3 (1980): 195–197.

Striebel, V. H. "Zur Embrionalentwicklung der Termiten". *Acta Tropica* 17 (1960): 193–260.

Stroyan, H. L. G. "Notes on the early stages of Rhopalus parumpunctatus Schill. (Hemiptera: Coreidae)". *Proceedings of the Royal Entomological Society of London. Series A, General Entomology* 29.1-3 (1954): 32–38.

Suazo, A., D. P. Pacheco, R. D. Cave, and J. H. Frank. "Longevity and fecundity of Metamasius quadrilineatus Champion (Coleoptera: Dryophthoridae) on a natural bromeliad host in the laboratory". *The Coleopterists Bulletin* 60.3 (2006): 264–270.

Sukontason, K. L., N. Bunchu, T. Chaiwong, B. Kuntalue, and K. Sukontason. "Fine structure of the eggshell of the blow fly, Lucilia cuprina". *Journal of Insect Science* 7.1 (2007): 1–8.

Sukontason, K. L., P. Sribanditmongkol, T. Chaiwong, R. C. Vogtsberger, S. Piangjai, and K. Sukontason. "Morphology of immature stages of Hemipyrellia ligurriens (Wiedemann) (Diptera: Calliphoridae) for use in forensic entomology applications". *Parasitology Research* 103.4 (2008): 877–887.

Sukontason, K. L., K. Sukontason, N. Boonchu, T. Chaiwong, and S. Piangjai. "Ultrastructure of eggshell of Chrysomya nigripes Aubertin (Diptera: Calliphoridae)". *Parasitology Research* 93.2 (2004): 151–154.

Sukontason, K. L., K. Sukontason, R. C. Vogtsberger, S. Piangjai, N. Boonchu, and T. Chaiwong. "Ultramorphology of eggshell of flesh fly Liosarcophaga dux (Diptera: Sarcophagidae)". *Journal of Medical Entomology* 42.1 (2005): 86–88.

Sukontason, K., K. L. Sukontason, S. Piangjai, N. Boonchu, H. Kurahashi, M. Hope, and J. K. Olson. "Identification of forensically important fly eggs using a potassium permanganate staining technique". *Micron* 35.5 (2004): 391–395.

Suksuwan, W., X. Cai, L. Ngernsiri, and S. Baumgartner. "Segmentation gene expression patterns in Bactrocera dorsalis and related insects: regulation and shape of blastoderm and larval cuticle". *International Journal of Developmental Biology* 61.6-7 (2017): 439–450.

Suludere, Z. "Description of the eggs of Rhodostrophia meonaria Guenée from North Pakistan (Geometridae: Lepidoptera)". *Communucations, Faculty of Sciences, University of Ankara, Series C* 6 (1988): 47–52.

Suludere, Z. "Studies on the external morphology of the eggs of some Argynninae species (Satyridae: Lepidoptera)". *Communucations, Faculty of Sciences, University of Ankara, Series C* 6 (1988): 9–28.

Suludere, Z. "Studies on the external morphology of the eggs of some Melitaea species (Satyridae: Lepidoptera)". *Communucations, Faculty of Sciences, University of Ankara, Series C* 6 (1988): 73–84.

Suludere, Z., S. Canbulat, and S. Candan. "External morphology of eggs of Macronemurus bilineatus and Megistopus flavicornis (Neuroptera, Myrmeleontidae): a scanning electron microscopy study". *Turkish Journal of Zoology* 33.4 (2009): 387–392.

Suludere, Z., S. Candan, Y. Kalender, and A. Hasbenli. "Ultrastructure of the chorion of Machimus rusticus (Meigen, 1820) (Diptera, Asilidae)". *Journal of the Entomological Research Society* 2.2 (2000): 63–71.

Suludere, Z., A. Satar, S. Candan, and S. Canbulat. "Morphology and surface structure of eggs and first instar larvae of Dielocroce baudii (Neuroptera: Nemopteridae) from Turkey". *Entomological News* 117.5 (2006): 521–530.

Suman, D. S., A. R. Shrivastava, B. D. Parashar, S. C. Pant, O. P. Agrawal, and S. Prakash. “Variation in morphology and morphometrics of eggs of Culex quinquefasciatus mosquitoes from different ecological regions of India”. *Journal of Vector Ecology* 34.2 (2009): 191–199.

Suzuki, M. and T. Tanaka. “Development of Meteorus pulchricornis and regulation of its noctuid host, Pseudaletia separata”. *Journal of Insect Physiology* 53.10 (2007): 1072–1078.

Suzuki, N. “Embryology of the Mecoptera (Panorpidae, Panorpodidae, Bittacidae and Boreidae)”. *Bulletin of the Sugadaira Montane Research Center University of Tsukuba* 11 (1990): 1–87.

Suzuki, N., S. Shimizu, and H. Ando. “Early embryology of the alderfly, Sialis mitsuhashii Okamoto (Megaloptera: Sialidae)”. *International Journal of Insect Morphology and Embryology* 10.5 (1981): 409–418.

Švácha, P. “Bionomics, behaviour and immature stages of Pelecotoma fennica (Paykull) (Coleoptera: Rhipiphoridae)”. *Journal of Natural History* 28.3 (1994): 585–618.

Svihla, A. “The life history of Tanypteryx hageni Selys (Odonata)”. *Transactions of the American Entomological Society (1890-)* 85.3 (1959): 219–232.

Swadener, S. O. and T. R. Yonke. “Immature stages and biology of Sinea complexa with notes on four additional reduviids (Hemiptera: Reduviidae)”. *Journal of the Kansas Entomological Society* 46.1 (1973): 123–136.

Swadener, S. O. and T. R. Yonke. “Immature stages and biology of Apiomerus crassipes (Hemiptera: Reduviidae)”. *Annals of the Entomological Society of America* 66.1 (1973): 188–196.

Swadener, S. O. and T. R. Yonke. “Immature stages and biology of Zelus socius (Hemiptera: Reduviidae)”. *The Canadian Entomologist* 105.2 (1973): 231–238.

Swaminathan, S. and V. Sriramulu. “Embryogenesis of Chrysocoris purpureus (Westw.) (Hemiptera: Pentatomidae)”. *Proceedings: Plant Sciences* 81.2 (1975): 75–82.

Swammerdam, J. *The Book of Nature; or, The History of Insects: Reduced to distinct Classes, confirmed by particular Instances, Displayed in the Anatomical Analysis of many Species, and Illustrated with Copper-plates including the Generation of the Frog, the History of the Ephemerus, the Changes of Flies, Butterflies, and Beetle; with the Original Discovery of the Milk-Vessels of the Cuttle-Fish, and many other curious Particulars*. London: printed for C. G. Seyffert, Bookseller, in Dean-Street, Soho, 1758.

Swan, D. I. “The common nemobiine field crickets of New Zealand (Orthoptera: Gryllidae)”. *Journal of the Royal Society of New Zealand* 2.4 (1972): 533–539.

Taber, S. W. “A new Nearctic species of Micropsectra Kieffer midge (Diptera: Chironomidae)”. *Southwestern Entomologist* 37.1 (2012): 61–71.

Takada, Y., S. Kawamura, and T. Tanaka. “Biological characteristics. Growth and development of the egg parasitoid Trichogramma dendrolimi (Hymenoptera: Trichogrammatidae) on the cabbage armyworm Mamestra brassicae (Lepidoptera: Noctuidae)”. *Applied Entomology and Zoology* 35.3 (2000): 369–379.

Tan, J.-L., M.-J. Duan, L.-F. Yin, H.-W. Hao, and X.-x. Chen. “The pre-overwintering nests and the immature stages of the hornet Vespa fumida van der Vecht (Hymenoptera: Vespidae)”. *Journal of Natural History* 47.19-20 (2013): 1325–1337.

Tan, J.-L. and B. Hua. “Description of the immature stages of Bittacus planus Cheng (Mecoptera: Bittacidae) with notes on its biology”. *Proceedings of the Entomological Society of Washington* 111.1 (2009): 111–121.

Tanaka, M. “Early embryonic development of amata fortunei (Lepidoptera, Amatidae)”. *Recent Advances in Insect Embryology in Japan*. ISEBU Co.: Tsukuba Science City, 1985. 139–155.

Tanaka, M. “Early embryonic development of the parasitic wasp, Trichogramma chilonis (Hymenoptera, Trichogrammatidae)”. *Recent Advances in Insect Embryology in Japan*. ISEBU Co.: Tsukuba Science City, 1985. 171–179.

Tauber, C. A., M. J. Tauber, and M. J. Tauber. “Egg size and taxon: their influence on survival and development of chrysopid hatchlings after food and water deprivation”. *Canadian Journal of Zoology* 69.10 (1991): 2644–2650.

Tauber, M. J., C. A. Tauber, and T. W. Hilton. “Life history and reproductive behavior of the endemic Hawaiian Anomalochrysa hepatica (Neuroptera: Chrysopidae): A comparative approach”. *European Journal of Entomology* 103.2 (2006): 327–336.

Tavares, M., L. A. Kaminski, and G. R. P. Moreira. "External morphology of the immature stages of neotropical heliconians: II. Dione juno juno (Cramer) (Lepidoptera, Nymphalidae, Heliconiinae)". *Revista Brasileira de Zoologia* 19.4 (2002): 961–976.

Tawfik, M. F. S., M. Hafez, and A. A. Ibrahim. "Immature stages of Microplitis rufiventris Kok. (Hym., Braconidae)". *Deutsche Entomologische Zeitschrift* 27.1-3 (1980): 39–50.

Tawfik, M. F. S., S. I. El-Sherif, and A. F. Lutfallah. "On the life-history of the giant water-bug Limnogeton fieberi Mayr (Hemiptera: Belostomatidae), predatory on some harmful snails". *Zeitschrift für Angewandte Entomologie* 86.1-4 (1978): 138–145.

Tawfik, M. F. S., S. I. El-Sherif, and A. F. Lutfallah. "The biology of Sphaerodema urinator Duf.(Hemiptera, Belostomatidae)". *Zeitschrift für Morphologie und Ökologie der Tiere* 86.1-4 (1978): 266–273.

Taylor, G. S. "The structure of the eggs of some Australian Psylloidea (Hemiptera)". *Australian Journal of Entomology* 31.2 (1992): 109–117.

Taylor, M. E., C. S. Bundy, and J. E. Mcpherson. "Life History and Laboratory Rearing of Bagrada hilaris (Hemiptera: Heteroptera: Pentatomidae) with Descriptions of Immature Stages". *Annals of the Entomological Society of America* 108.4 (2015): 536–551.

Teixeira, É. P. and S. A. Casari. "Descriptions and biological notes of immatures of Microctenochira Difficilis (Coleoptera, Chrysomelidae, Hispinae, Cassidini)". *Iheringia, Série Zoologia* 93.1 (2003): 23–30.

Tewari, S. K., V. Kumar, A. K. Awasthi, and R. K. Datta. "Surface morphology of egg chorion of the Uzi fly, Exorista bombycis (Louis),(Diptera: Tachinidae)-an endoparasite of the silkworm, Bombyx mori Linn." *Zoological Studies* 34.1 (1995): 62–66.

Thiery, A., C. Martinz, C. Malosse, and D. Thiery. "Morphology and chemical characterization of the egg chorion in tingids: a case study of the plane tree Corythucha ciliata (Hemiptera: Tingidae)". *Entomological Problems* 30.1 (1999): 73–82.

Thireau, J. C., J. Régnière, and C. Cloutier. "Biology and morphology of immature stages of Meteorus trachynotus Vier. (Hymenoptera: Braconidae)". *Canadian Journal of Zoology* 68.5 (1990): 1000–1004.

Thomas, J. A., M. L. Munguira, J. Martin, and G. W. Elmes. "Basal hatching by Maculinea butterfly eggs: a consequence of advanced myrmecophily?" *Biological Journal of the Linnean Society* 44.2 (1991): 175–184.

Thompson, P. B., M. P. Parrella, B. C. Murphy, and M. L. Flint. "Life history and description of Dasineura gleditchiae (Diptera: Cecidomyiidae) in California". *The Pan-Pacific Entomologist* 74.2 (1998): 85–98.

Thyssen, P. J. and A. X. Linhares. "First description of the immature stages of Hemilucilia segmentaria (Diptera: Calliphoridae)". *Biological Research* 40.3 (2007): 271–280.

Tian, J., B.-z. Hua, and H.-j. Zhang. "Morphology of Eogystia sibirica (Alphéraky) (Lepidoptera: Cossidae) attacking Asparagus officinalis in northern China with descriptions of its immature stages". *Journal of Natural History* 44.43-44 (2010): 2581–2595.

Tiegs, O. W. and F. V. Murray. "Memoirs: the embryonic development of Calandra oryzae". *Journal of Cell Science* 2.318 (1938): 159–273.

Tiwari, N. K. "Eupelmus tenuicornis Kieffer (Hymenoptera: Chalcidoidea), a parasite of Bimba toombii Grover (Diptera: Cecidomyiidae)". *Journal of Applied Entomology* 74.1-4 (1973): 384–388.

Togashi, K. and M. Itabashi. "Maternal size dependency of ovariole number in Dastarcus helophoroides (Coleoptera: Colydiidae)". *Journal of Forest Research* 10.5 (2005): 373–376.

Tojo, K. and R. Machida. "Early embryonic development of the mayfly Ephemera japonica McLachlan (Insecta: Ephemeroptera, Ephemeridae)". *Journal of Morphology* 238.3 (1998): 327–335.

Tojo, K. and K. Matsukawa. "A description of the second species of the family Dipteromimidae (Insecta, Ephemeroptera), and genetic relationship of two dipteromimid mayflies inferred from mitochondrial 16S rRNA gene sequences". *Zoological Science* 20.10 (2003): 1249–1259.

Tonapi, G. T. "A note on the eggs and larva of Dineutes indicus Aube (Coleoptera, Gyrinidae)". *Current Science* 28 (1959): 158–159.

Torchio, P. F. “The biology of Perdita nuda and descriptions of its immature forms and those of its Sphecodes parasite (Hymenoptera: Apoidea)”. *Journal of the Kansas Entomological Society* 48.3 (1975): 257–279.

Torchio, P. F. “The nesting biology of Hylaeus bisinuatus Forster and development of its immature forms (Hymenoptera: Colletidae)”. *Journal of the Kansas Entomological Society* 57.2 (1984): 276–297.

Torchio, P. F., J. G. Rozen Jr, G. E. Bohart, and M. S. Favreau. “Biology of Dufourea and of its cleptoparasite, Neopasites (Hymenoptera: Apoidea)”. *Journal of the New York Entomological Society* 75.3 (1967): 132–146.

Torchio, P. F. and N. N. Youssef. “The biology of Anthophora (Micranthophora) flexipes and its cleptoparasite, Zacosmia maculata, including a description of the immature stages of the parasite (Hymenoptera: Apoidea, Anthophoridae)”. *Journal of the Kansas Entomological Society* 41.3 (1968): 289–302.

Torchio, P. F. “In-nest biologies and development of immature stages of three Osmia species (Hymenoptera: Megachilidae)”. *Annals of the Entomological Society of America* 82.5 (1989): 599–615.

Torchio, P. F. “Late embryogenesis and egg eclosion in Triepeolus and Anthophora with a prospectus of nomadine classification (Hymenoptera: Anthophoridae)”. *Annals of the Entomological Society of America* 79.4 (1986): 588–596.

Torchio, P. F. “The ethology of the wasp, Pseudomasaris edwardsii (Cresson), and a description of its immature forms (Hymenoptera: Vespoidea, Masaridae)”. *Contributions in Science, Los Angeles County Museum* 202 (1970): 1–32.

Torchio, P. F. and D. J. Burdick. “Comparative notes on the biology and development of Epeolus compactus Cresson, a cleptoparasite of Colletes kincaidii Cockerell (Hymenoptera: Anthophoridae, Colletidae)”. *Annals of the Entomological Society of America* 81.4 (1988): 626–636.

Torchio, P. F. and G. E. Trostle. “Biological notes on Anthophora urbana urbana and its parasite, Xeromelecta californica (Hymenoptera: Anthophoridae), including descriptions of late embryogenesis and hatching”. *Annals of the Entomological Society of America* 79.3 (1986): 434–447.

Tormos, J., F. Beitia, E. A. Böckmann, and J. D. Asís. “The preimaginal stages and development of Spalangia cameroni Perkins (Hymenoptera: Pteromalidae) on Ceratitis capitata (Wiedemann) (Diptera: Tephritidae)”. *Micron* 40.5 (2009): 646–658.

Tormos, J., F. Beitia, E. A. Böckmann, J. D. Asís, and S. Fernández. “The preimaginal phases and development of Pachycrepoideus vindemmiae (Hymenoptera, Pteromalidae) on mediterranean fruit fly, Ceratitis capitata (Diptera, Tephritidae)”. *Microscopy and Microanalysis* 15.5 (2009): 422–434.

Tormos, J., L. de Pedro, F. Beitia, B. Sabater, J. D. Asís, and C. Polidori. “Development, preimaginal phases and adult sensillar equipment in Aganaspis parasitoids (Hymenoptera: Figitidae) of fruit flies”. *Microscopy and Microanalysis* 19.6 (2013): 1475–1489.

Torréns, J. and J. M. Heraty. “A new genus of Eucharitidae (Hymenoptera: Chalcidoidea), with notes on life history and immature stages”. *Zootaxa* 3630.2 (2013): 347–358.

Torréns, J. and J. M. Heraty. “Description of the species of Dicoelothorax Ashmead (Chalcidoidea, Eucharitidae) and biology of D. platycerus Ashmead”. *ZooKeys* 165 (2012): 33–46.

Torréns, J., J. M. Heraty, and P. Fidalgo. “Biology and description of a new species of Lophyrocera Cameron (Hymenoptera: Eucharitidae) from Argentina”. *Zootaxa* 1871 (2008): 56–62.

Torres, P. L. M., M. C. Michat, and M. Archangelsky. “Description of the preimaginal stages of three species of the genus Tropisternus Solier, subgenus Strepitornus Hansen (Coleoptera: Hydrophilidae), with emphasis on morphometry and chaetotaxy”. *Zootaxa* 1702 (2008): 1–25.

Torretta, J. P., S. P. Durante, M. G. Colombo, and A. M. Basilio. “Nesting biology of the leafcutting bee Megachile (Pseudocentron) gomphrenoides (Hymenoptera: Megachilidae) in an agro-ecosystem”. *Apidologie* 43.6 (2012): 624–633.

Toschi, C. A. “The taxonomy, life histories, and mating behavior of the green lacewings of Strawberry Canyon (Neuroptera: Chrysopidae)”. *Hilgardia* 36.11 (1964): 391–431.

Trehan, K. N. “Studies on the british white-flies (Homoptera, Aleyrodidae)”. *Transactions of the Royal Entomological Society of London* 90.22 (1940): 575–616.

Triggerson, C. J. "A study of Dryophanta erinacei (Mayr) and its gall". *Annals of the Entomological Society of America* 7.1 (1914): 1–46.

Tripp, H. A. "Descriptions and Habits of Cecidomyiidae (Diptera) from White Spruce Cones". *The Canadian Entomologist* 87.6 (1955): 253–263.

Tripp, H. A. "The biology of Perilampus hyalinus Say (Hymenoptera: Perilampidae), a primary parasite of Neodiprion swainei Midd. (Hymenoptera: Diprionidae) in Quebec, with descriptions of the egg and larval stages". *The Canadian Entomologist* 94.12 (1962): 1250–1270.

Trostle, G. E. and P. F. Torchio. "Notes on the nesting biology and immature development of Euparagia scutellaris Cresson (Hymenoptera: Masaridae)". *Journal of the Kansas Entomological Society* 59.4 (1986): 641–647.

Trotta-Moreu, N., E. Montes de Oca, and I. M. MartÍnez. "Ecological and Reproductive characteristics of Geotrupes (Halffterius) rufoclavatus Jekel 1865 (Coleoptera: Geotrupidae: Geotrupinae) on the Cofre de Perote Volcano (Veracruz, Mexico)". *The Coleopterists Bulletin* 61.3 (2007): 435–446.

Trueman, J. W. H. "Egg chorionic structures in Corduliidae and Libellulidae (Anisoptera)". *Odonatologica* 20.4 (1991): 441–452.

Trueman, J. W. H. "Eggshells of Australian Gomphidae: Plastron respiration in eggs of stream-dwelling Odonata (Anisoptera)". *Odonatologica* 19 (1990): 395–401.

Tsai, J. H. and S. W. Wilson. "Biology of Peregrinus maidis with descriptions of immature stages (Homoptera: Delphacidae)". *Annals of the Entomological Society of America* 79.3 (1986): 395–401.

Tschudi-Rein, K. and S. Dorn. "Reproduction and immature development of Hyssopus pallidus (Hymenoptera: Eulophidae), an ectoparasitoid of the codling moth". *European Journal of Entomology* 98.1 (2001): 41–46.

Tsui, P. T. P. and W. L. Peters. "Embryonic development, early instar morphology, and behavior of Tortopus incertus (Ephemeroptera: Polymitarcidae)". *The Florida Entomologist* 57.4 (1974): 349–356.

Tsuneki, K. "Ethological studies on the Japanese species of Pemphredon (Hymenoptera, Sphecidae), with notes on their parasites, Ellampus spp. (Hym., Chrysididae) (With 5 Text-figures)". *Journal of the Faculty of Science Hokkaido University, Series VI Zoology* 11.1 (1952): 57–75.

Tuck, J. B. and R. C. Smith. "Identification of the eggs of mid-western grasshoppers by the chorionic sculpturing". *Agricultural Experiment Station Technical Bulletin* 48 (1939): 1–39.

Turillazzi, S. "Brood rearing behaviour and larval development in Parischnogaster nigricans serrei (Du Buysson) (Hymenoptera Stenogastrinae)". *Insectes Sociaux* 32.2 (1985): 117–127.

Turillazzi, S. and M. H. Hansell. "Biology and social behaviour of three species of Anischnogaster (Vespidae, Stenogastrinae) in Papua New Guinea". *Insectes Sociaux* 38.4 (1991): 423–437.

Turillazzi, S. "Egg deposition in the genus Parischnogaster (Hymenoptera: Stenogastrinae)". *Journal of the Kansas Entomological Society* 58.4 (1985): 749–752.

Ubero-Pascal, N. and M. A. Puig. "Egg morphology update based on new chorionic data of Potamanthus luteus (Linnaeus), Ephemera danica Müller and Oligoneuriella rhenana (Imhoff) (Insecta, Ephemeroptera) obtained by scanning electron microscopy". *Zootaxa* 1465 (2007): 15–29.

Uchifune, T. and R. Machida. "Embryonic development of Galloisiana yuasai Asahina, with special reference to external morphology (Insecta: Grylloblattodea)". *Journal of Morphology* 266.2 (2005): 182–207.

Uemiya, H. and H. Ando. "Blastodermic cuticles of a springtail, Tomocerus ishibashii Yosii (Collembola: Tomoceridae)". *International Journal of Insect Morphology and Embryology* 16.5-6 (1987): 287–294.

Uemiya, H. and H. Ando. "Embryogenesis of a springtail, Tomocerus ishibashii (Collembola, Tomoceridae): external morphology". *Journal of Morphology* 191.1 (1987): 37–48.

Ullmann, S. L. "The Origin and Structure of the Mesoderm and the Formation of the Coelomic Sacs in Tenebrio molitor L. [Insecta, Coleoptera]". *Philosophical Transactions of the Royal Society of London B: Biological Sciences* 248.747 (1964): 245–277.

Urban, J. "A contribution to the knowledge of biology and harmfulness of Deporaus betulae (L.) (Coleoptera, Attelabidae)". *Acta Universitatis Agriculturae et Silviculturae Mendelianae Brunensis* 60.6 (2012): 317–338.

Urban, J. “Apoderus coryli (L.)–a Biologically Little Known Species of the Attelabidae (Coleoptera)”. *Acta Universitatis Agriculturae et Silviculturae Mendelianae Brunensis* 62.5 (2014): 1141–1160.

Urban, J. “Biology of Byctiscus populi (L.) (Coleoptera, Attelabidae). Part II. Leafrolls, larvae and this year’s imagoes”. *Acta Universitatis Agriculturae et Silviculturae Mendelianae Brunensis* 60.1 (2013): 155–166.

Urban, J. “Occurrence, Biology and Harmfulness of Byctiscus betulae (L.) (Coleoptera, Rhynchitidae)”. *Acta Universitatis Agriculturae et Silviculturae Mendelianae Brunensis* 63.5 (2015): 1601–1624.

Urban, J. “Occurrence, development and harmfulness of the bark anobiid Ernobius mollis (L.) (Coleoptera: Anobiidae)”. *Journal of Forest Science* 51.8 (2005): 327–347.

Urbaneja, A., H. Montón, and O. Mollá. “Suitability of the tomato borer Tuta absoluta as prey for Macrolophus pygmaeus and Nesidiocoris tenuis”. *Journal of Applied Entomology* 133.4 (2009): 292–296.

Vala, J.-C., C. Casc, G. Gbedjissi, and C. Dossou. “Life history, immature stages and sensory receptors of Sepedon (Parasepedon) trichrooscelis an Afrotropical snail-killing fly (Diptera: Sciomyzidae)”. *Journal of Natural History* 29.4 (1995): 1005–1014.

Valim, M. P. and A. C. Cicchino. “Immature stages of chewing lice (Insecta: Phthiraptera) from Neotropical Icteridae (Aves: Passeriformes), and descriptions of three new species”. *Annales Zoologici* 65.3 (2015): 491–521.

Valim, M. P. and A. C. Cicchino. “Six new species of Myrsidea Waterston, 1915 (Phthiraptera: Menoponidae) from New World jays of the genus Cyanocorax Boie (Passeriformes: Corvidae), with notes on the chorionic structure of eggs”. *Systematic Parasitology* 90.2 (2015): 191–211.

Valley, K. and A. G. Wheeler. “Biology and Immature Stages of Stomopteryx palpilineella (Lepidoptera: Gelechiidae), a Leaf miner and Leaf tier of Crownvetch”. *Annals of the Entomological Society of America* 69.2 (1976): 317–324.

Van der Starre-van der Molen, L. G. “Embryogenesis of Calliphora erythrocephala Meigen. I. Morphology”. *Netherlands Journal of Zoology* 22.2 (1971): 119–182.

Van Emden, F. I. “Mormotomyia hirsuta Austen (Diptera) and its systematic position”. *Proceedings of the Royal Entomological Society of London. Series B, Taxonomy* 19.7-8 (1950): 121–128.

Van Veen, J. C. and M. L. E. Wijk. “The unique structure and functions of the ovipositor of the non-paralyzing ectoparasitoid Colpoclypeus florus Walk. (Hym., Eulophidae) with special reference to antennal sensilla and immature stages”. *Zeitschrift für angewandte Entomologie* 99.1-5 (1985): 511–531.

Vårdal, H., G. Sahlén, and F. Ronquist. “Morphology and evolution of the cynipoid egg (Hymenoptera)”. *Zoological Journal of the Linnean Society* 139.2 (2003): 247–260.

Vargas, H. A., F. X. Oyarzún, and L. E. Parra. “Egg and First Instar of the Neotropical Geometrid Moth Pero obtusaria Prout (Geometridae: Ennominae: Azelinini)”. *The Journal of the Lepidopterists’ Society* 71.1 (2017): 50–56.

Vargas, H. A., R. Brito, D. S. Basilio, and G. R. P. Moreira. “A morphological reappraisal of the immature stages and life history of Elachista synethes Meyrick (Lepidoptera, Elachistidae), an Australian leaf miner alien to Chile”. *Revista Brasileira de Entomologia* 59.4 (2015): 265–273.

Vaught, G. L. and K. W. Stewart. “The life history and ecology of the stonefly Neoperla clymene (Newman) (Plecoptera: Perlidae)”. *Annals of the Entomological Society of America* 67.2 (1974): 167–178.

Venkatesha, M. G. and K. Gopinath. “Description of immature stages of a species of Glyptapanteles (Hymenoptera: Braconidae), a gregarious endoparasitoid of Amata passalis (Fabricius) (Lepidoptera: Arctiidae), a defoliator of sandalwood, Santalum album L.” *International Journal of Tropical Insect Science* 15.2 (1994): 161–165.

Vennison, S. J. and D. P. Ambrose. “Diversity of eggs and ovipositional behaviour in Reduviids (Insecta, Heteroptera, Reduviidae) of South India”. *Mitteilungen aus dem Museum für Naturkunde in Berlin. Zoologisches Museum und Institut für Spezielle Zoologie (Berlin)* 66.2 (1990): 319–331.

Vilimova, J. and M. Rohanova. “The external morphology of eggs of three Rhopalidae species (Hemiptera: Heteroptera) with a review of the eggs of this family”. *Acta Entomologica Musei Nationalis Pragae* 50.1 (2010): 75–95.

Villalobos, G., J. A. Martínez-Ibarra, F. Martínez-Hernández, S. López-Alcaide, and R. Alejandre-Aguilar. "The morphological variation of the eggs and genital plates of two morphotypes of Triatoma protracta Uhler, 1894". *Journal of Vector Ecology* 37.1 (2012): 179–186.

Villet, M. "Qualitative relations of egg size, egg production and colony size in some ponerine ants (Hymenoptera: Formicidae)". *Journal of Natural History* 24.5 (1990): 1321–1331.

Vincini, A. M., A. N. López, P. L. Manetti, H. Alvarez-Castillo, and D. Mabel Carmona. "Description of the immature stages of Dyscinetus rugifrons (Burmeister, 1847) (Coleoptera: Scarabaeidae: Dynastinae)". *Elytron* 14 (2000): 91–98.

Visciarelli, E., A. Ferrero, and S. R. Costamagna. "Exochorial aspects of eggs of Triatoma patagonica Del Ponte, 1929 shown by scanning electron microscopy". *Entomología y Vectores* 11.4 (2004): 653–668.

Visscher, P. K. and R. S. Vetter. "Annual and multi-year nests of the western yellowjacket, Vespula pensylvanica, in California". *Insectes Sociaux* 50.2 (2003): 160–166.

Vogelgesang, M. and T. Szklarzewicz. "Formation and structure of egg capsules in scale insects (Hemiptera, Coccinea): I. Ortheziidae". *Arthropod Structure & Development* 30.1 (2001): 63–68.

Volkoff, A.-N., J. Daumal, P. Barry, M.-C. François, N. Hawlitzky, and M. M. Rossi. "Development of Trichogramma cacoeciae Marchal (Hymenoptera: Trichogrammatidae): time table and evidence for a single larval instar". *International Journal of Insect Morphology and Embryology* 24.4 (1995): 459–466.

Volkoff, N. and S. Colazza. "Growth patterns of teratocytes in the immature stages of Trissolcus basalis (Woll.) (Hymenoptera: Scelionidae), an egg parasitoid of Nezara viridula (L.) (Heteroptera: Pentatomidae)". *International Journal of Insect Morphology and Embryology* 21.4 (1992): 323–336.

Von Tschirnhaus, M. "4.3. 01 Acartophthalmidae, Borboropsidae, Chyromyidae, Micropezidae, Odiniidae, Opetiidae, Periscelididae, Pseudopomyzidae, and Tanypezidae". *A dipterological perspective on a changing alpine landscape. Results from a survey of the biodiversity of Diptera (Insecta) in the Stilfserjoch National Park (Italy).* Vol. Supplement. 2008. 65–97.

Al-Wahaibi, A. K. and J. G. Morse. "Egg morphology and stages of embryonic development of the glassy-winged sharpshooter (Hemiptera: Cicadellidae)". *Annals of the Entomological Society of America* 102.2 (2009): 241–248.

Wall, C. "Embryonic development in two species of Chesias (Lepidoptera: Geometridae)". *Journal of Zoology* 169.1 (1973): 65–84.

Wallace, M. M. H. "The biology of the jarrah leaf miner, Perthida glyphopa Common (Lepidoptera: Incurvariidae)". *Australian Journal of Zoology* 18.1 (1970): 91–104.

Wallin, H., P. A. Chiverton, B. S. Ekbom, and A. Borg. "Diet, fecundity and egg size in some polyphagous predatory carabid beetles". *Entomologia Experimentalis et Applicata* 65.2 (1992): 129–140.

Waloff, N. "The effect of the number of queens of the antLasius flavus (Fab.) (Hym., Formicidae) on their survival and on the rate of development of the first brood". *Insectes Sociaux* 4.4 (1957): 391–408.

Waloff, N. "The egg pods of british short-horned grasshoppers (Acrididae)". *Proceedings of the Royal Entomological Society of London. Series A, General Entomology* 25.10-12 (1950): 115–126.

Walsh, D. B., M. P. Bolda, R. E. Goodhue, A. J. Dreves, J. Lee, D. J. Bruck, V. M. Walton, S. D. O'Neal, and F. G. Zalom. "Drosophila suzukii (Diptera: Drosophilidae): invasive pest of ripening soft fruit expanding its geographic range and damage potential". *Journal of Integrated Pest Management* 2.1 (2011): 1–7.

Walton, G. A. "The egg of Agraptocorixa gestroi Kirkaldy (Hemiptera-Heteroptera: Corixidae)". *Proceedings of the Royal Entomological Society of London. Series A, General Entomology* 37.7-9 (1962): 104–106.

Wang, Y.-K. "Life History of Mayatrichia Ponta Ross (Trichoptera: Hydroptilidae) in Honey Creek, Turner Falls Park, Oklahoma". MA thesis. University of North Texas, 1997.

Wang, L. Y. "Eggs and oviposition of some Sphingidae". *Acta Entomologica Sinica* 27.4 (1984): 478–479.

Wang, Y., M. Shi, X. Hou, S. Meng, F. Zhang, and J. Ma. "Adaptation of the egg of the desert beetle, Microdera punctipennis (Coleoptera: Tenebrionidae), to arid environment". *Journal of Insect Science* 14.1 (2014): 1–8.

Wang, Z.-P., Y.-S. Liu, X.-H. He, F. Lv, and H. He. "Morphology and biology of Carabus smaragdinus". *Chinese Bulletin of Entomology* 5 (2008): 814–817.

Wang, Z., M. A. Alonso-Zarazaga, D. Zhou, and R. Zhang. "A description of preimaginal stages of Pseudaspidapion botanicum Alonso-Zarazaga & Wang, 2011 (Apionidae, Curculionoidea)". *ZooKeys* 260 (2013): 49–59.

Ward, R. D. "Some observations on the biology and morphology of the immature stages of Psychodopygus wellcomei fraiha, shaw and lainson, 1971: (Diptera: psychodidae)". *Memórias do Instituto Oswaldo Cruz* 70.1 (1972): 15–28.

Ward, R. D. "The immature stages of some phlebotomine sandflies from Brazil (Diptera: Psychodidae)". *Systematic Entomology* 1.3 (1976): 227–240.

Warne, A. C. "Embryonic development and the systematics of the Tettigoniidae (Orthoptera: Saltatoria)". *International Journal of Insect Morphology and Embryology* 1.3 (1972): 267–287.

Wąsowska, M. "Morphology of the first instar larva and of the egg of Labidostomis longimana (Linnaeus, 1761) and of Labidostomis tridentata (Linnaeus, 1758) (Coleoptera, Chrysomelidae, Clytrinae), with a key to clytrine genera with the first instar larva known". *Deutsche Entomologische Zeitschrift* 54.1 (2007): 51–67.

Watson, J. A. L. and C. D. Howick. "The rediscovery of Mastopsenius australis Seevers (Coleoptera: Staphylinidae)". *Australian Journal of Entomology* 14.1 (1975): 19–21.

Watt, J. C. "Entomology of the Aucklands and other islands south of New Zealand: Coleoptera: Scarabaeidae, Byrrhidae, Ptinidae, Tenebrionidae". *Pacific Insects Monograph* 27 (1971): 193–224.

Wegner, A. M. R. "Biological notes on Megacrania wegneri Willemse and M. Alpheus. Westwood (Orthoptera, Phasmidae)". *Treubia* 23.1 (2016): 47–52.

Weigensberg, I., Y. Carriere, and D. A. Roff. "Effects of male genetic contribution and paternal investment to egg and hatchling size in the cricket, Gryllus firmus". *Journal of Evolutionary Biology* 11.2 (1998): 135–146.

Weinstein, P. and A. D. Austin. "Primary parasitism, development and adult biology in the wasp Taeniogonalos venatoria Riek (Hymenoptera: Trigonalyidae)". *Australian Journal of Zoology* 43.6 (1995): 541–555.

Weintraub, P. G. and A. R. Horowitz. "The newest leafminer pest in Israel, Liriomyza huidobrensis". *Phytoparasitica* 23.2 (1995): 177–184.

Weng, J.-L., K. Nishida, P. Hanson, and L. LaPierre. "Biology of Lissoderes Champion (Coleoptera, Curculionidae) in Cecropia saplings inhabited by Azteca ants". *Journal of Natural History* 41.25-28 (2007): 1679–1695.

Wengrat, A. P. G. D. S., V. C. Matesco, K. R. Barão, J. Grazia, and V. Pietrowski. "External morphology of the immature stages of Vatiga manihotae (Hemiptera: Tingidae) with comments on ontogenesis". *The Florida Entomologist* 98.2 (2015): 626–632.

West, J. A., G. E. Cantwell, and T. J. Shortino. "Embryology of the house fly, Musca domestica (Diptera: Muscidae), to the blastoderm stage". *Annals of the Entomological Society of America* 61.1 (1968): 13–17.

Wheeler Jr, A. G. and E. R. Hoebeke. "Biology and seasonal history of Rhopalus (Brachycarenus) tigrinus, with descriptions of immature stages (Heteroptera: Rhopalidae)". *Journal of the New York Entomological Society* 96.4 (1988): 381–389.

Wheeler Jr, A. G. and S. W. Wilson. "Life history of the issid planthopper Thionia elliptica (Homoptera: Fulgoroidea) with description of a new Thionia species from Texas". *Journal of the New York Entomological Society* 95.3 (1987): 440–451.

Wheeler, A. G. and G. L. Miller. "Leptoglossus fulvicornis (Heteroptera: Coreidae), a specialist on magnolia fruits: seasonal history, habits, and descriptions of immature stages". *Annals of the Entomological Society of America* 83.4 (1990): 753–765.

Wheeler, G. C. and J. Wheeler. "Larvae of the formicine ant genus Polyrhachis". *Transactions of the American Entomological Society* 116.3 (1990): 753–767.

Wheeler, G. C. and J. Wheeler. "Notes on ant larvae". *Transactions of the American Entomological Society* 115.4 (1989): 457–473.

Wheeler, G. C. and J. Wheeler. "Supplementary studies on ant larvae: Formicinae (Hymenoptera: Formicidae)". *Journal of the New York Entomological Society* 94.3 (1986): 331–341.

Wheeler, G. C. and J. Wheeler. "Young larvae of Eciton (Hymenoptera: Formicidae: Dorylinae)". *Psyche* 93.3-4 (1986): 341–350.

Wheeler, G. C. and J. Wheeler. “Young larvae of Veromessor pergandei (Hymenoptera: Formicidae)”. *Psyche* 94.3-4 (1987): 303–307.

Wheeler, W. M. *The embryology of Blatta germanica and Doryphora decemlineata*. Boston: Ginn and Company, 1889.

Wheeler, W. M. “A contribution to insect embryology”. *Journal of Morphology* 8.1 (1893): 1–160.

Wiesenborn, W. D. “The thrips (Thysanoptera) Liothrips xanthocerus (Phlaeothripidae) and Neohydatothrips catenatus (Thripidae) inhabit leaf clusters on Pluchea sericea (Asteraceae)”. *The Florida Entomologist* 94.3 (2011): 706–708.

Wigglesworth, V. B. and J. W. L. Beament. “The respiratory mechanisms of some insect eggs”. *Quarterly Journal of Microscopical Science* 3.16 (1950): 429–452.

Wilkinson, J. D. and D. M. Daugherty. “The Biology and Immature Stages of Bradysia impatiens (Diptera: Sciaridae)”. *Annals of the Entomological Society of America* 63.3 (1970): 656–660.

Williams, F. X. “Notes on the life-history of some North American Lampyridae”. *Journal of the New York Entomological Society* 25.1 (1917): 11–33.

Williams, J. R. “The sugar-cane Delphacidae and their natural enemies in Mauritius”. *Transactions of the Royal Entomological Society of London* 109.2 (1957): 65–110.

Williams, L., M. C. Coscaró, P. M. Dellapé, and T. M. Roane. “The shield-backed bug, Pachycoris stallii: Description of immature stages, effect of maternal care on nymphs, and notes on life history”. *Journal of Insect Science* 5.1 (2005): 1–13.

Wilson, D. D. and R. L. Ridgway. “Morphology, development, and behavior of the immature stages of the parasitoid, Campoletis sonorensis (Hymenoptera: Ichneumonidae)”. *Annals of the Entomological Society of America* 68.2 (1975): 191–196.

Wilson, L. F. “Life history, habits, and damage of a gall midge, Oligotrophus papyriferae (Diptera: Cecidomyiidae), injurious to paper birch in Michigan”. *The Canadian Entomologist* 100.6 (1968): 663–669.

Wilson, L. F. “Life history, habits, and damage of the boxelder leaf gall midge, Contarinia negundifolia Felt (Diptera: Cecidomyiidae) in Michigan”. *The Canadian Entomologist* 98.7 (1966): 777–784.

Wilson, L. F. and G. C. Heaton. “Life history, damage, and gall development of the gall midge, Neolasioptera brevis (Diptera: Cecidomyiidae), injurious to honeylocust in Michigan”. *Great Lakes Entomologist* 20.3 (1987): 111–118.

Wilson, S. W. and J. E. McPherson. “Descriptions of the immature stages of Bruchomorpha oculata with notes on laboratory rearing”. *Annals of the Entomological Society of America* 74.4 (1981): 341–344.

Wilson, S. W. and J. H. Tsai. “Descriptions of the immature stages of Myndus crudus (Homoptera: Fulgoroidea: Cixiidae)”. *Journal of the New York Entomological Society* 90.3 (1982): 166–175.

Windsor, D. M., D. W. Trapnell, and G. Amat. “The egg capitulum of a Neotropical walkingstick, Calynda bicuspis, induces aboveground egg dispersal by the ponerine ant, Ectatomma ruidum”. *Journal of Insect Behavior* 9.3 (1996): 353–367.

Winterbourn, M. J. and N. H. Anderson. “The life history of Philanisus plebeius Walker (Trichoptera: Chathamiidae), a caddisfly whose eggs were found in a starfish”. *Ecological Entomology* 5.3 (1980): 293–304.

Wipfler, B., M. Bai, S. Schoville, R. Dallai, T. Uchifune, R. Machida, Y. Cui, and R. G. Beutel. “Ice Crawlers (Grylloblattodea)–the history of the investigation of a highly unusual group of insects”. *Journal of Insect Biodiversity* 2.2 (2014): 1–25.

Withycombe, C. L. “XV. Some Aspects of the Biology and Morphology of the Neuroptera. With special reference to the immature stages and their possible phylogenetic significance”. *Transactions of the Royal Entomological Society of London* 72.3-4 (1925): 303–411.

Woglum, R. S. and E. A. McGregor. “Observations on the life history and morphology of Agulla astuta (Banks) (Neuroptera: Raphidiodea: Raphidiidae)”. *Annals of the Entomological Society of America* 52.5 (1959): 489–502.

Woglum, R. S. and E. A. McGregor. “Observations on the life history and morphology of Agulla bractea Carpenter (Neuroptera: Raphidiodea: Raphidiidae)”. *Annals of the Entomological Society of America* 51.2 (1958): 129–141.

Wolf, K. W., C. Murphy, W. Reid, and E. Garraway. “Fine structure of the eggshell in Utetheisa ornatrix (Lepidoptera: Arctiidae)”. *Invertebrate Reproduction & Development* 38.2 (2000): 85–94.
Wolf, K. W., W. Reid, and D. A. Rider. “Eggs of the stink bug Acrosternum (Chinavia) marginatum (Hemiptera: Pentatomidae): a scanning electron microscopy study”. *Journal of Submicroscopic Cytology and Pathology* 34.2 (2002): 143–150.
Wolf, K. W. and W. Reid. “Egg morphology and hatching in Mormidea pictiventris (Hemiptera: Pentatomidae)”. *Canadian Journal of Zoology* 79.4 (2001): 726–736.
Wolf, K. W. and W. Reid. “The architecture of the anterior appendage in the egg of the assassin bug, Zelus longipes (Hemiptera: Reduviidae)”. *Arthropod Structure & Development* 29.4 (2000): 333–341.
Wolf, K. W., W. Reid, and M. Schrauf. “Optical illusions in scanning electron micrographs: the case of the eggshell of Acrosternum (Chinavia) marginatum (Hemiptera: Pentatomidae)”. *Micron* 34.1 (2003): 57–62.
Wolfe, K. L. and M. A. Balcazar-Lara. “Chile’s Cercophana venusta and its immature stages”. *Tropical Lepidoptera* 5.1 (1994): 35–42.
Wood, J. R., V. H. Reshm, and E. M. McEwan. “Egg masses of Nearctic sericostomatid caddisfly genera (Trichoptera) [Fattigia, Gumaga, ecology, taxonomy]”. *Annals of the Entomological Society of America* 75.4 (1982): 430–434.
Woodley, N. E. and D. D. Judd. “Notes on the host, egg, and puparium of Stylogaster biannulata (Say) (Diptera: Conopidae)”. *Proceedings of the Entomological Society of Washington* 100.4 (1998): 658–664.
Woodroffe, G. E. “A life-history study of Endrosis lactella (Schiff.) (Lep. Oecophoridae)”. *Bulletin of Entomological Research* 41.4 (1951): 749–760.
Woodroffe, G. E. “A life-history study of the brown house moth, Hofmannophila pseudospretella (Staint.) (Lep., Oecophoridae)”. *Bulletin of Entomological Research* 41.3 (1951): 529–553.
Woodward, T. E. “On Australian and New Zealand Peloridiidae (Homoptera: Coleorrhyncha)”. *University of Queensland Papers* 1.3 (1956): 31–56.
Wright, E. J. “Immature stages of Encyrtus saliens (Hymenoptera: Encyrtidae), an imported parasite of ice plant scales (Homoptera: Coccidae) in California”. *Annals of the Entomological Society of America* 79.2 (1986): 273–279.
Xu, P., Z.-w. Wan, X.-x. Chen, S. Liu, and M.-g. Feng. “Immature morphology and development of Opius caricivorae (Hymenoptera: Braconidae), an endoparasitoid of the leafminer Liriomyza sativae (Diptera: Agromyzidae)”. *Annals of the Entomological Society of America* 100.3 (2007): 425–432.
Yamada, Y. “Characteristics of the oviposition of a parasitoid, Chrysis shanghaiensis (Hymenoptera: Chrysididae)”. *Applied Entomology and Zoology* 22.4 (1987): 456–464.
Yan, J., X. Qin, G. Shaanxi, S. Henan, J. Shandong, J. Anhui, and H. Hubei. “Anoplophora glabripennis (Motsch.)” *Forest Insects of China*. Beijing: China Forestry Publishing House, 1992. 455–457.
Yang, C. T. and C. I. Tsay. “Immature stages of three species of the genus Epipaylla (Homoptera: Psyllidae)”. *Proceedings of the National Science Council, Republic of China* 4.4 (1980): 418–423.
Yano, T. “The developmental stages of four species of the Japanese Pentatomidae (Hemiptera)”. *Transactions of the Shikoku Entomological Society* 2.1 (1951): 7–16.
Yen, A. L. “The immature stages of Psylla acaciaepycnanthae Froggatt and Psylla uncatoides Ferris and Klyver (Hemiptera: Psylloidea)”. *Australian Entomological Magazine* 11.4, 5 (1984): 69–74.
Yeo, Y. S., Y. D. Chang, and H. G. Hoh. “A morphological observation of an egg parasitoid, Anagrus incarnatus Haliday (Hymenoptera: Mymaridae), of the rice planthoppers”. *Korean Journal of Applied Entomology* 29.1 (1990): 1–5.
Yonke, T. R. and J. T. Medler. “Description of immature stages of Coreidae. 1. Euthochtha galeator”. *Annals of the Entomological Society of America* 62.3 (1969): 469–473.
Yonke, T. R. and J. T. Medler. “Description of immature stages of Coreidae. 2. Acanthocephala terminalis”. *Annals of the Entomological Society of America* 62.3 (1969): 474–476.
Yonke, T. R. and J. T. Medler. “Description of immature stages of Coreidae. 3. Archimerus alternatus”. *Annals of the Entomological Society of America* 62.3 (1969): 477–480.

Yonke, T. R. and D. L. Walker. “Description of the egg and nymphs of Harmostes reflexulus (Hemiptera: Rhopalidae)”. *Annals of the Entomological Society of America* 63.6 (1970): 1749–1754.

Youssef, N. N. and G. E. Bohart. “The nesting habits and immature stages of Andrena (Thysandrena) candida Smith (Hymenoptera, Apoidea)”. *Journal of the Kansas Entomological Society* 41.4 (1968): 442–455.

Yu, R.-X., M. Shi, F. Huang, and X.-X. Chen. “Immature development of Cotesia vestalis (Hymenoptera: Braconidae), an endoparasitoid of Plutella xylostella (Lepidoptera: Plutellidae)”. *Annals of the Entomological Society of America* 101.1 (2008): 189–196.

Zacharuk, R. Y. “Distribution, habits, and development of Ctenicera destructor (Brown) in western Canada, with notes on the related species C. aeripennis (Kby.) (Coleoptera: Elateridae)”. *Canadian Journal of Zoology* 40.4 (1962): 539–552.

Zanuncio, T. V., J. C. Zanuncio, G. P. Santos, M. C. Q. Fialho, and A. S. Bernardino. “Aspectos biológicos e morfológicos de Mimallo amilia (Lepidoptera: Mimallonidae) em folhas de Eucalyptus urophylla”. *Revista Árvore* 29.2 (2005): 321–326.

Zawadzka, M., W. Jankowska, and S. M. Biliński. “Egg shells of mallophagans and anoplurans (Insecta: Phthiraptera): morphogenesis of specialized regions and the relation to F-actin cytoskeleton of follicular cells”. *Tissue and Cell* 29.6 (1997): 665–673.

Zenker, M. M., A. Specht, and E. Corseuil. “Immature stages of Spodoptera cosmioides (Walker) (Lepidoptera, Noctuidae)”. *Revista Brasileira de Zoologia* 24.1 (2007): 99–107.

Zhang, L.-J. and X.-k. Yang. “Description of the immature stages of Ophrida xanthospilota (Baly) (Coleoptera: Chrysomelidae: Alticinae) from China”. *Proceedings of the Entomological Society of Washington* 110.3 (2008): 693–700.

Zhao, W., S. Dong, M. Shi, and X.-X. Chen. “Morphology and Development of Immature Stage of Diadromus collaris (Hymenoptera: Ichneumonidae), an Important Endoparasitoid of Plutella xylostella (Lepidoptera: Plutellidae)”. *Annals of the Entomological Society of America* 107.1 (2014): 234–241.

Zilahi-Balogh, G. M. G., L. M. Humble, L. T. Kok, and S. M. Salom. “Morphology of Laricobius nigrinus (Coleoptera: Derodontidae), a predator of the hemlock woolly adelgid”. *The Canadian Entomologist* 138.5 (2006): 595–601.

Zimin, L. S. “Les pontes des acridiens: morphologie, classification et écologie”. *Tableaux Analytiques de la Fauna de l’URSS* 23 (1938): 1–83.

Zimmerman, J. H., H. D. Newson, G. R. Hooper, and H. A. Christensen. “A comparison of the egg surface structure of six anthropophilic phlebotomine sandflies (Lutzomyia) with the scanning electron microscope (Diptera: Psychodidae)”. *Journal of Medical Entomology* 13.4-5 (1977): 574–579.

Zissler, D. and K. Sander. “The cytoplasmic architecture of the egg cell of Smittia spec. (Diptera, Chironomidae)”. *Development Genes and Evolution* 172.3 (1973): 175–186.

Zompro, O. “Redescription and new synonymies of Heteronemia Gray, 1835 (Insecta: Phasmatodea) transferred to the suborder Areolatae”. *Studies on Neotropical Fauna and Environment* 36.3 (2001): 221–225.

Zompro, O., J. Adis, and W. Weitschat. “A Review of the Order Mantophasmatodea (Insecta)”. *Zoologischer Anzeiger* 241.3 (2002): 269–279.

Zongo, J. O., C. Vincent, and R. K. Stewart. “Biology of Trichogrammatoidea simmondsi (Hym.: Trichogrammatidae) on sorghum shoot fly, Atherigona soccata (Dipt.: Muscidae) eggs”. *Entomophaga* 38.2 (1993): 267–272.

Zwölfer, H. “Investigations on Sphenoptera (Chilostetha) jugoslavica Obenb.(Col. Buprestidae), a possible biocontrol agent of the weed Centaurea diffusa Lam. (Compositae) in Canada”. *Zeitschrift für angewandte Entomologie* 80.1-4 (1976): 170–190.
